# *ob*ABPP-HT*: A Precision-Engineered Activity Proteomics Pipeline for the Streamlined Discovery of Deubiquitinase Inhibitors

**DOI:** 10.1101/2025.05.27.656269

**Authors:** Hannah B. L. Jones, Simeon D. Draganov, Sofia Schönbauer Huamán, Peter A. C. Wing, Chuong Nguyen, Zhu Liang, Julia Dörner, Jasper Lithgow, Emma Murphy, Alice Beard, Andrea Pierangelini, Jack W. Houghton, Mackenzie E. Robert, Sarah Flannery, Anthony Tumber, Iolanda Vendrell, Edward W. Tate, Paul R. Elliot, Graham M. West, Darragh P. O’Brien, Eidarus Salah, Andrew P. Turnbull, Christopher J. Schofield, Benedikt M. Kessler, Lennart Brewitz, Adán Pinto-Fernández

## Abstract

Deubiquitinases (DUBs), and the dysregulation thereof, are implicated in human disease. The recent inclusion of selective DUB inhibitors in clinical trials has heightened interest in DUB-focused drug discovery. Current DUB screening methods remain constrained, however, as they often rely on recombinant proteins that are truncated or derived from non-human sources, typically necessitating extensive optimisation of initial hits. We introduce a high-throughput, endogenous human DUB-focused activity proteomics workflow designed for the simultaneous screening and profiling of small, targeted libraries of catalytic group-reactive compounds. In a proof-of-concept screen, this innovative platform expanded the repertoire of electrophilic groups targeting DUBs, leading to the discovery of potent and selective inhibitors for USP47, OTUD7B, and USP5. Remarkably, these inhibitors required minimal or no optimisation to confirm the previously reported biological roles of the three DUBs, underscoring the advantages of this methodology for drug discovery applications.

## Main

Post-translational ubiquitylation regulates key cellular processes, including protein and organelle homeostasis, through the interplay of ubiquitinating and deubiquitinating machineries^1–4^. Deubiquitinases (DUBs), primarily nucleophilic cysteine enzymes, cleave Ub from biomolecules, mainly proteins, modulating their fate^5, 6^. Dysregulated DUB functions are linked to diseases and are tractable therapeutic targets^1^. Small-molecule USP1 and USP30 inhibitors are in clinical trials for cancer^7, 8^, chronic kidney disease^9^, and Parkinson’s disease, for example^3, 10^.

Activity-based probes (ABPs) targeting nucleophilic cysteine DUBs have been developed^11–14^. We and others have designed assays using a haemagglutinin (HA)-tagged, ubiquitin-based DUB ABP with a propargylamine (PA) warhead (HA-Ub-PA) to assess inhibitor potency and selectivity across endogenous DUBs, whilst also providing information on cellular permeability, and stability^15–17^. DUB activity proteomics, or activity-based protein profiling (ABPP), uses affinity purification in combination with liquid chromatography-tandem mass spectrometry (LC-MS/MS) to analyse DUB–ABP complexes. Unlike recombinant enzyme assays, ABPP evaluates ligand binding to endogenous, unmodified proteins in a cellular environment, supporting the discovery of selective inhibitors, including those for challenging targets^18, 19^.

ABPP’s broader applicability was historically limited by low sensitivity and throughput, due to labour-intensive workflows encompassing ABP labelling, purification, sample preparation, MS analysis, and data processing. To enhance its utility for DUB inhibitor discovery, we previously increased throughput by adapting the probe purification step to a 96-well plate format, integrating an automated liquid handling system (Agilent Bravo) with ultra-fast, sensitive LC– MS/MS instrumentation (Evosep-timsTOF). This method (ABPP-HT) facilitated analysis of ∼100 samples daily^20^. We subsequently enhanced the sensitivity of ABPP-HT by incorporating data-independent acquisition LC-MS/MS (DIA), enabling deeper and more comprehensive DUBome profiling (ABPP-HT*)^21^. Further improvements aimed at enhancing sensitivity included using diverse Ub probe approaches, high-pH pre-fractionation^22^, and isobaric labelling (Extended Data Fig. 1 a–e).

Here, we present an on-bench version of the ABPP-HT* workflow, named “*ob*ABPP-HT*”, which maintains the throughput and sensitivity of ABPP-HT* without requiring automated platforms for ABP immunoprecipitation. We used *ob*ABPP-HT* to screen a curated small-molecule library enriched in cysteine-reactive scaffolds and substrate mimetics. This resulted in the discovery of selective inhibitors for USP47, OTUD7B, and USP5, which were subsequently validated in biochemical and cellular assays. Our findings showcase *ob*ABPP-HT* as a widely accessible platform and the utility of using advanced activity proteomics to accelerate DUB inhibitor discovery and development.

## Results

### Development of a Flexible ABPP-HT Methodology for Broad Applicability

To enhance accessibility, the *ob*ABPP-HT* workflow incorporates magnetic beads and a multi-well magnetic rack, circumventing the need for highly specialised equipment (Fig. 1a). Compared to our Bravo robot protocol, *ob*ABPP-HT* enables greater DUBome depth (Fig. 1b), enriching in excess of 40 DUBs in the same MCF-7 cell line (Fig. 1c-e). This may be a result of extended binding times during affinity purification. The high-throughput approach uses a single-shot DIA method with 11 min gradients on an Evosep-timsTOF LC-MS/MS, enabling analysis of approximately 100 samples per day. For laboratories lacking elements of this setup, increased throughput can also be achieved through TMT-based multiplexing, which has been previously incorporated into DUB ABPP^18^ (Extended Data Fig. a-c).

**Fig. 1.**
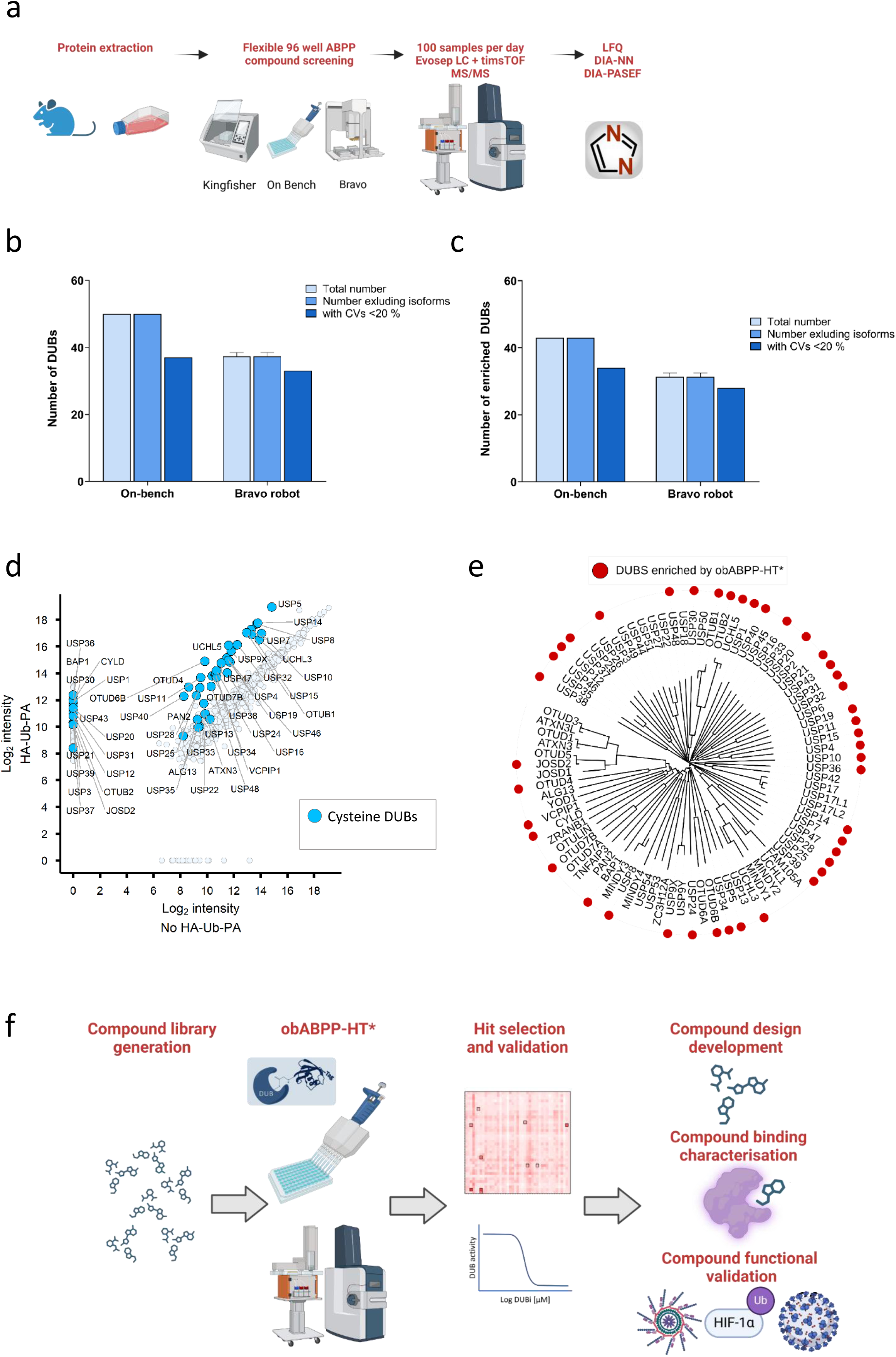
High-throughput and sensitive DUB activity proteomics methodology: *ob*ABPP-HT*. **a**, HA-Ub-PA high-throughput workflows. Created in BioRender. Jones, H. (2025) https://BioRender.com/yl1zds3. **b-c**, The number of cysteine-active DUBs identified from MCF-7 cell lysates by ABPP in total (**b**), or enriched > 5-fold compared to a negative control (**c**) using the high-throughput workflows outlined in **a**, with on-bench or Bravo robot 96-well immunoprecipitation (error bars = standard deviation (SD)). **d**, Enrichment of cysteine-active DUBs with the obABPP-HT* methodology. **e**, Phylogenetic tree of cysteine-active DUBs, with those enriched by the obABPP-HT* methodology denoted with red circles. The phylogenetic tree was generated with clustal omega multiple sequence alignment^84^, and visualised using iTOL^85^. **f**, obABPP-HT* DUB inhibitor screen workflow. Created in BioRender. Jones, H. (2025) https://BioRender.com/gkv1hyo. LFQ = Label-free quantitation, DIA-PASEF = data-independent acquisition-Parallel Accumulation–Serial Fragmentation.

We further validated the versatility of *ob*ABPP-HT* by translating it onto a Thermo KingFisher Apex purification system (Fig. 1a), using two different ABPs (HA-Ub-PA and bio-ISG15-PA), in a second cell line (HAP1)^20, 23, 24^. In this configuration, the Ub probe captured DUB profiles (Extended Data Fig. 1f, 1g) comparable to the on-bench method, while the ISG15 probe selectively reacted with known deISGylases, including USP5, USP14, USP16, USP18 and USP36^23, 25^ (Extended Data Fig. 1h).

### *ob*ABPP-HT* for DUB Inhibitor Identification

We demonstrated the utility of the *ob*ABPP-HT* workflow for the discovery of DUB inhibitors by investigating a curated library of small-molecules capable of reacting with nucleophilic cysteine enzymes (Fig. 1f). These included αβ,α′β′-diepoxyketones (DEKs), some known to covalently react with nucleophilic cysteine enzymes^26^, γ-lactams, with examples reported to covalently react with the SARS-CoV-2 main protease^27^, and Ub-derived oligopeptides specifically designed for this study, bearing either a C-terminal nitrile or alkyne group (Table S1-3).

Small-molecules (25 µM) were incubated with MCF-7 breast cancer cell lysates in a 96 well plate, enabling the profiling of 41 compounds and FT827, a well-characterised DUB inhibitor^28^, in technical duplicates. The plate also included negative/positive controls (−/+ HA-Ub-PA). Across replicates, 38 DUBs were consistently enriched >5-fold, with an average coefficient of variation (CV) of 16%, confirming robustness (Extended Data Fig. 2a). Reassuringly, *ob*ABPP-HT* replicated the known potent and selective USP7 inhibition by FT827^28^. Eight compounds significantly inhibited at least one DUB (>20% intensity loss, *p* < 0.05; Fig. 2a). Of these, **Compounds 4** and **27** were excluded due to a lack of selectivity. Selective hits included: **Compound 3** (USP47 > USP7), **Compound 6** (OTUD7B), **Compound 7** (USP30/USP47/VCPIP1), **Compound 25** (USP5), and **Compounds 26** and **32** (USP33) (Fig. 2b, 2c). Notably, USP5-selective **Compound 25** is a diastereomer of the inactive **Compound 27**, highlighting the workflow’s ability to resolve isomer-specific effects. (Extended Data Fig. 2b, 2c).

**Fig. 2.**
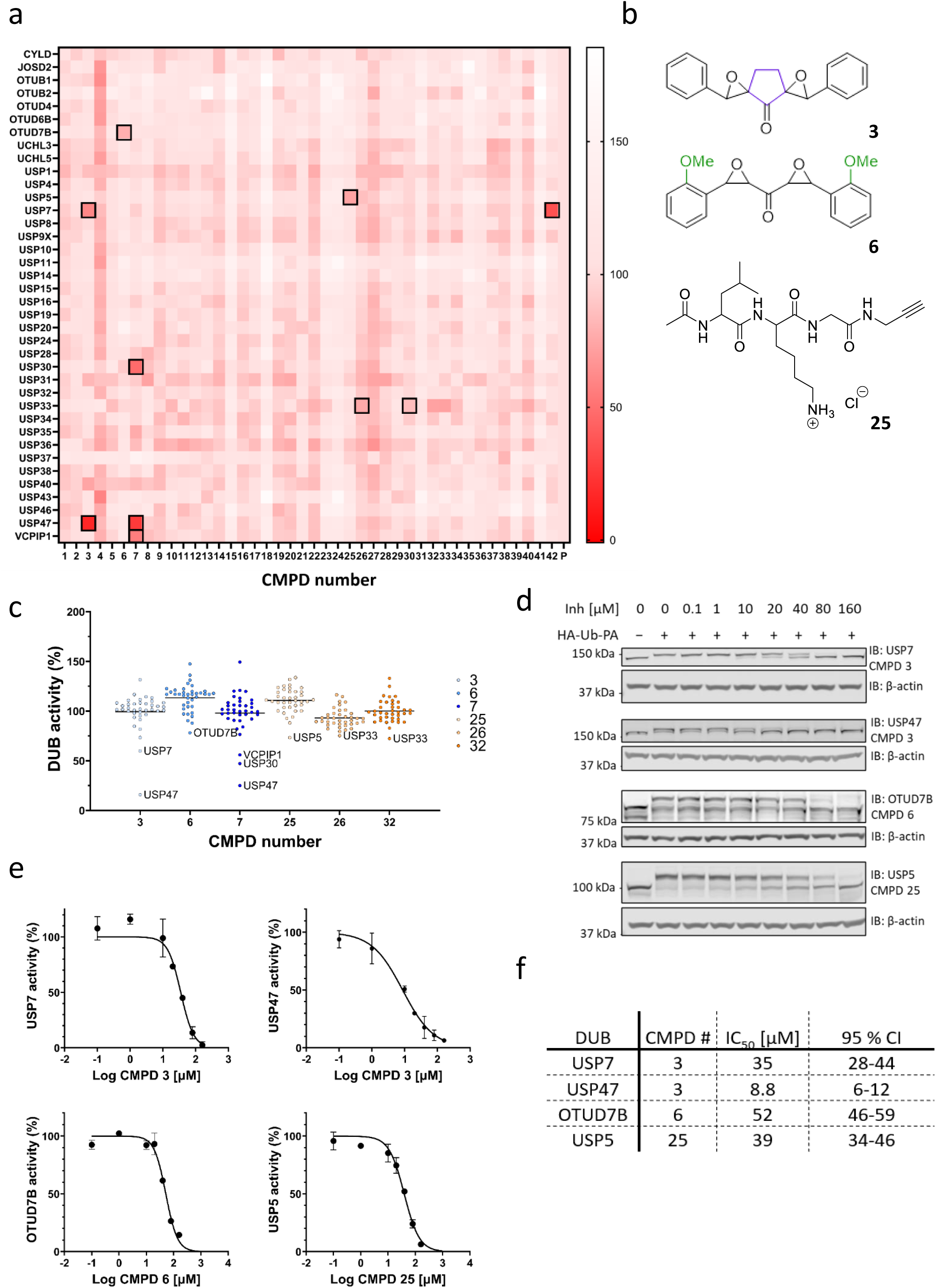
DUB inhibitor screen using *ob*ABPP-HT*. **a**, Heatmap of DUBs identified from *ob*ABPP-HT* screen, performed using MCF-7 lysates. DUB activity in the presence of 25 µM of **Compounds 1-41** (**Compound 42** = established USP7 inhibitor FT827) as a % of positive control activity (n = 2, controls n = 4). Cells outlined with black squares indicate > 20 % inhibition (p < 0.05 - two tailed unpaired t-test, (**Compound 6** OTUD7B p= 0.066)), with significant inhibition from **Compounds 4** and **27** not highlighted due to lack of selectivity. **b**, Compound structures of positive hits. **c,** Significant hits from **a** taken forward for validations, showing reduction in activity relative to other DUBs. **d**, HA-Ub-PA ABP validations in MCF-7 cell lysates showing concentration-dependent inhibition of DUBs by positive hits identified from *ob*ABPP-HT* screen. **e**, Densitometric quantitation from western blots of positive hits (n = 2) showing log inhibitor concentration *vs* normalised DUB activity, fit to Y=100/(1+10^((LogIC50-X)*HillSlope)) (error bars = SD). **f**, IC50 values extracted from fits in e, (CI= Confidence interval).

### Hit Validation and Dose-Response Analysis

Western blots confirmed inhibition of USP47/USP7 by **Compound 3**, OTUD7B by **Compound 6**, and USP5 by **Compound 25** (Fig. 2d). DUB inhibition was concentration-dependent (Fig. 2e) and IC₅₀ values revealed the preference of **Compound 3** for USP47 (8.8 μM) over USP7 (36 μM) (Fig. 2f), which was consistent with screening data. **Compounds 26** and **32** were deemed false positives (Extended Data Fig. 2d), while **Compound 7** induced protein crosslinking and/or aggregation, likely due to its bifunctional epoxide structure (Extended Data Fig. 2e)^26^. To assess the stability of binding, **Compounds 3**, **6**, and **25** were pre-incubated with MCF-7 lysates before probe addition and incubated for 5, 15, or 45 min. Minimal inhibitor displacement by HA-Ub-PA was observed (Extended Data Fig. 2f–i), suggesting irreversible binding at DUB active sites.

### DUB Inhibitor Selectivity Assessment by *ob*ABPP-HT*

Due to its increased throughput capabilities, the *ob*ABPP-HT* workflow was also used to assess the selectivity of inhibitors **3**, **6**, and **25** across several concentrations (Fig. 3a-f). The dose-dependent inhibition curves and IC50 values for the three compounds were comparable to those generated by immunoblotting (Extended Data Fig. 3 a-d)^20^. **Compound 3** exclusively inhibited USP47 and USP7 up to 50 µM, with weak USP40 inhibition being observed at higher doses (Fig. 3a, 3d). The inhibition profile of **Compound 3** was consistent when tested in HAP1 cells using the KingFisher purification system (Extended Data Fig. 3e). **Compound 6** was selective for OTUD7B up to 100 µM but also affected USP16 and USP28 at 250 µM (Fig. 3b, 3e). Notably, **Compound 25** was highly selective, only inhibiting USP5 across the tested concentration range (Fig. 3c, 3f).

**Fig. 3.**
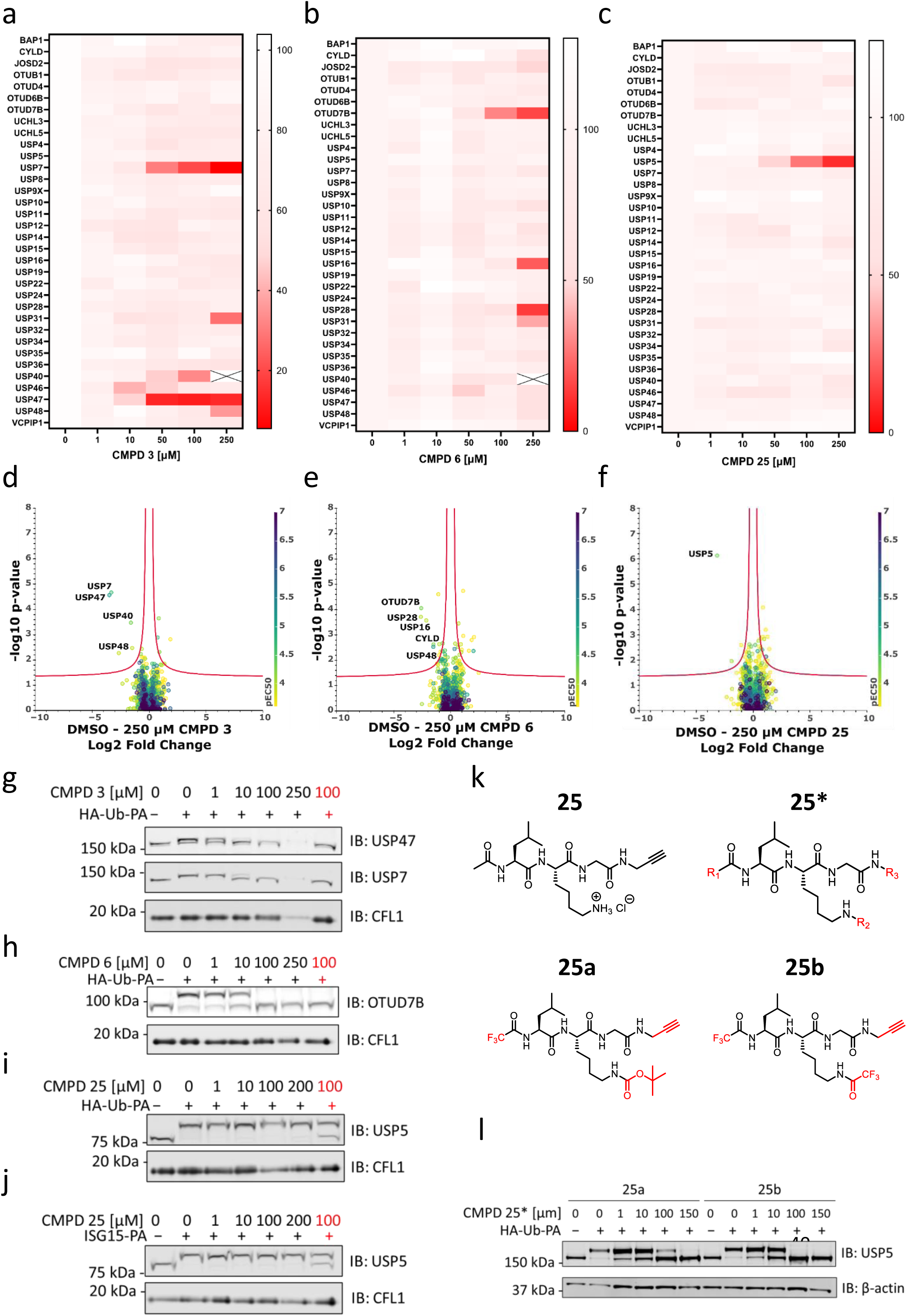
Hit profiling using *ob*ABPP-HT* and cellular permeability. **a-c**, *ob*ABPP-HT* DUB activity with concentration-dependence of **Compounds 3** (**a**), **6** (**b**) and **25** (**c**) (n = 2, controls n = 3). **d-f**, Curve curator volcano plots^75^, showing Log2 Fold change decrease in DUB intensity with 250 µM of **Compound 3** (**d**), **6** (**e**), and **25** (**f**). Colour is reflective of pEC50 values across concentration-dependence shown in in a-c. **g-j**, Cell permeability concentration-dependence of **Compounds 3**, **6** and **25** after incubation with MCF-7 cells for 4 h, lysis and subsequent labelling with HA-Ub-PA or ISG15-PA. **k**, Varying the R1 and R2 groups of **Compound 25** enhances cellular permeability. **l**, Cellular permeability concentration-dependence of USP5 inhibition with **Compounds 25a** and **25b** after incubation with MCF-7 cells for 4 h, lysis and subsequence labelling with HA-Ub-PA.

The active sites of DUBs and deISGylases are structurally similar and most deISGylating enzymes are cross-reactive DUBs, cleaving both Ub and ISG15, with the notable exception of USP18^23–25, 29^. Given USP5’s dual Ub/ISG15 specificity, we evaluated the effect of **Compound 25** on deISGylases by immunoblotting, using the bio-ISG15-PA ABP. Remarkably, **Compound 25** inhibited USP5 to a similar extent as with the Ub ABP, and did not cross-react with USP18 or any of the other detected deISGylases in the tested concentration range (up to 125 μM), further demonstrating its high selectivity (Extended Data Fig. 3f). It is important to note that both **Compounds 3** and **6** are also reported to weakly inhibit the mycobacterial L,D-transpeptidase LdtMt2 and the SARS-CoV-2 main protease^26^, implying that they may also react with additional human nucleophilic cysteine enzymes other than DUBs.

### Cell Permeability and Inhibitor Optimisation

We assessed cell permeability of the identified DUB inhibitors by treating intact MCF-7 cells prior to probe labelling. **Compounds 3** and **6** were cell-permeable and inhibited USP47/USP7 and OTUD7B, respectively, in a concentration-dependent manner (Fig. 3g, 3h). **Compound 3** induced toxicity at the highest concentration (250 µM), suggesting off-target effects. **Compound 25**, however, was inactive in intact cells (Fig. 3i, 3j). To optimise **Compound 25** for cellular activity, we modified the alkyne and amine protecting groups. Substituting the terminal alkyne reduced potency (Fig. 3k; Extended Data Table 1; synthesis detailed in SI), supporting that **Compound 25** covalently reacts with the USP5 nucleophilic cysteine via its alkyne group. By contrast, altering amine-protecting groups at the N-terminus (R1) and/or at the *N*^ε^-lysine amino group (R2), led to two active derivatives, **25a** and **25b**, with trifluoromethyl groups at R1 and trifluoroacetyl (**25a**) or *tert*-butyloxycarbonyl (Boc; **25b**) at R2 (Fig. 3l). Both derivatives retained the potency and selectivity observed in lysates (Extended Data Fig. 4a), offering promising leads for functional studies or therapeutic development.

### Molecular Basis of Compound 3 Preference for USP47

Reported USP47 inhibitors inhibit USP7 with comparable potency^30, 31^, likely reflecting their highly related phylogeny and homology^29, 32^. Given the preference of **Compound 3** for USP47 over USP7, we decided to use it as a tool to further investigate its mode of action and elucidate molecular differences between the two DUBs. In fluorogenic ubiquitin-rhodamine (Ub-Rho) biochemical assays, **Compound 3** showed ∼10-fold more activity for recombinant USP47 over recombinant USP7 after 30 min preincubation (IC₅₀: 0.48 vs. 4.3 μM; Fig. 4a, Extended Data Fig. 5a–d). Time-dependency experiments indicated covalent or slow tight-binding inhibition for USP47 (Fig. 4c), while USP7 inhibition was weaker and time-independent (Fig. 4b). For both enzymes, progress curves in the presence of **Compound 3** were non-linear indicating slow initial binding (Extended Data Fig. 5e, 5f).

**Fig. 4.**
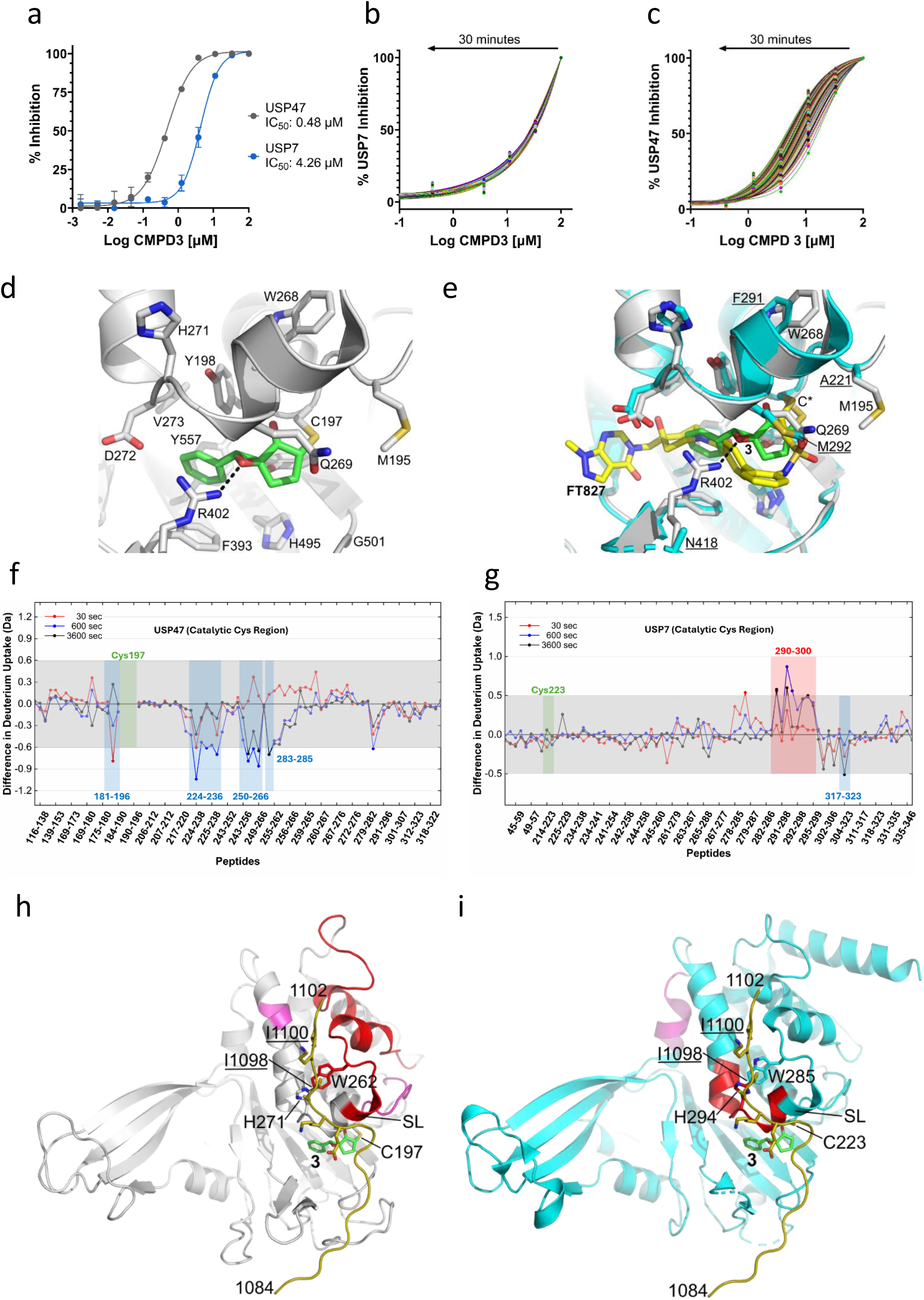
Molecular Basis for Selective Inhibition of USP47 by Compound 3. **a**, Inhibition of purified USP47 and USP7 determined from Ub-Rho cleavage after a 30 min pre-incubation with varying concentrations of **Compound 3**. IC50 curves fit to: Y=Bottom + (Top-Bottom)/(1+10^((LogIC50-X)*HillSlope)) (Error bars = SD). **b**, Inhibition of USP47 with **Compound 3** without preincubation, determined from Ub-Rho cleavage. Each curve represents inhibition data at an individual incubation time from 18 to 1782 s. **c**, Inhibition of USP7 with **Compound 3** without pre-incubation, determined from Ub-Rho cleavage. Each curve represents inhibition data at an individual incubation time from 18 to 1782 s. **d**, Modelled structure of human USP47 in complex with **Compound 3**, shown as a stick representation with carbon atoms coloured green. Residues within 5 Å of the **Compound 3**-C197 adduct are highlighted, and the key hydrogen-bonding interaction between the epoxide moiety of **Compound 3** and the side chain of R402 is represented as a dotted line. Figure prepared using PyMOL (The PyMOL Molecular Graphics System, version 2.5.8; Schrödinger, LLC). **e**, Superposition of the modelled structure of human USP47 (grey) in complex with **Compound 3** (green carbon atoms) on the X-ray structure of human USP7 (cyan) in complex with the covalent inhibitor, FT827 (yellow carbon atoms; PDB: 5NGF). C* denotes the catalytic cysteine (corresponding to C197 in human USP47 and C223 in human USP7). **Compound 3** is predicted to bind within the thumb-palm cleft in a similar fashion to FT827, with FT827 extending out further towards the fingers subdomain. Sequence differences are labelled for human USP47 and the corresponding residues are labelled and underlined for human USP7. R402 in USP47 is predicted to hydrogen bond directly to **Compound 3** (represented as a dotted line). R402 in USP47 corresponds to N418 (disordered side chain in PDB: 5NGF) in USP7 with a shorter side chain that cannot interact with the compound, which may explain the higher potency of **Compound 3** for USP47. Residual ΔHDX plots showing differences in deuterium uptake between holo- and apo-states for USP47 (**f**) and USP7 (**g**), across all indicated peptide fragments and time points (30 sec to 1 h). These zoomed-in views highlight the peptides surrounding only those regions of the full-length protein (residues 116-323 of USP47 (**f**) and 45-347 of USP7 (**g**)). These regions also encompass the catalytic cysteine residue of each DUB. Positive ΔHDX values indicate increased solvent exposure or flexibility in the presence of **Compound 3**, whilst negative values suggest protection or structural stabilisation. A grey shaded area marks the significance threshold for ΔHDX applied in each study. The green boxes indicate those peptides which encompass the catalytic cysteine. Due to the disappearance of these peptides in the holo-state of USP47, these peptides could not be visualised on the plot. This is further evidence of the covalent attachment of **Compound 3** to the catalytic cysteine in USP47. **h**, Modelled structure of human USP47 catalytic domain (grey) in complex with **Compound 3** (carbon atoms in green) highlighting regions identified in the HDX-MS analysis coloured red (mean perturbation over 60 min of 5–20%) and magenta (mean perturbation over 60 min of <60%). The 19-residue C-terminal activation peptide of human USP7 (PDB: 5JTV; corresponding to residues 1084 to 1102) is shown in gold with key residues labelled and underlined. The activation peptide of USP47 (corresponding to residues 1345 to 1362)^33^ is proposed to play an important role in activation and regulation analogous to the activation peptide of USP7, which maintains the switching loop (SL; especially W285 and H294) in the active “out” conformation^34^. W285 and H294 in human USP7 correspond to W262 and H271 in human USP47. Regions identified in the USP47 HDX-MS analysis correlate with a conformational shift in the SL and potential engagement of the activation peptide upon **Compound 3** binding. **i**, Human USP7 catalytic domain (cyan; from PDB: 5NGF) with the superimposed modelled position of **Compound 3** in human USP47 (green carbon atoms) highlighting regions identified in the HDX-MS analysis coloured red (mean perturbation over 60 min of 5–20%) and magenta (mean perturbation over 60 min of <60%). The 19-residue C-terminal activation peptide of human USP7 (PDB: 5JTV; corresponding to residues 1084 to 1102) is shown in gold with key residues labelled and underlined. Analysis of human USP7 HDX-MS data reveals that the region of highest confidence directly flanks the proposed compound binding site and differences are not observed in structural elements implicated in activation peptide binding, suggesting that **Compound 3** does not trigger an equivalent shift in SL to the active “out” conformation and subsequent binding of the activation peptide as predicted for USP47.

Human USP47 shares 43% sequence identity to human USP7 within the catalytic domain. In the absence of a human USP47 Protein Data Bank (PDB) structure, the structure of human USP7 in complex with the covalent inhibitor, FT827 (PDB: 5NGF), was used as the template to generate a homology model of human USP47 with an appropriate conformational state for covalent inhibitor docking studies. **Compound 3** most likely forms a covalent adduct with the catalytic cysteine of human USP47 (C197) via 1,2-carbonyl attack and retro-aldol fragmentation based on previously reported work (Extended Data Fig. 5g)^26^.

Docking studies suggest that **Compound 3** binds in the thumb-palm cleft (Extended Data Fig. 5h) and sterically blocks Ub binding (Extended Data Fig. 5i), analogous to the covalent USP7 inhibitor, FT827 (PDB: 5NGF; Extended Data Fig. 5j). The binding site is flanked by 14 residues including R402, which is predicted to form a key hydrogen bonding interaction with the epoxide moiety of **Compound 3** (Fig. 4d). In human USP7, the substitution of R402 with an asparagine residue (N418), which has a shorter side chain and cannot interact with the compound, may explain the higher potency of **Compound 3** for USP47 (Fig. 4e).

USP47 apoform (PDB: 8ITN) and Ub-bound (PDB: 8ITP) structures exist for the closely related catalytic domain of *C. elegans* USP47 (44% sequence identity). A comparison of the human USP47-**Compound 3** model with apoform *C.elegans* USP47 suggests that **Compound 3** binding disrupts hydrogen bonding between R402 and H271, which is involved in stabilising the switching loop “in”, catalytically incompetent conformation, thereby promoting an active, switching loop “out” conformation compatible with activation peptide binding as observed in USP7’s regulatory mechanism (Extended Data Fig. 5o, 5p)^33, 34^.

Hydrogen-deuterium exchange mass spectrometry (HDX-MS) analysis confirmed differential binding modes (Fig. 4f, 4g, and Extended Data Fig. 5k, 5l), revealing conformational shifts indicative of a switching loop rearrangement in USP47 (Extended Data Fig. 5m) and potential engagement of the activation peptide (Fig. 4h)^34^. Notably, HDX-MS for USP7 showed no such distal conformational changes, indicating that **Compound 3** does not promote equivalent switching loop reorganisation or activation peptide binding in USP7 (Fig. 4i and Extended Data Fig. 5n). These differences likely underpin **Compound 3**’s selectivity for USP47.

### Compound 3 Inhibits the NLRP3 Inflammasome

Both USP7 and USP47 are reported to regulate the NLRP3 inflammasome^32^. When inducing inflammasome activation with lipopolysaccharide (LPS) and nigericin in human monocyte cells (THP-1), treatment with **Compound 3** at low concentrations increased viability and reduced inflammasome-mediated pyroptosis (Extended Data Fig. 6a, 6b), suggesting inflammasome suppression. In contrast, high concentrations and prolonged nigericin exposure induced toxicity, consistent with additional targets beyond DUBs^26^. Importantly, at shorter nigericin incubation times, **Compound 3** inhibited inflammasome activation, as evidenced by reduced Caspase-1 activity (Fig. 5a, Extended Data Fig. 6 c-e), blocked oligomerisation of apoptosis-associated speck-like protein containing a CARD (ASC), and prevented cleavage of Caspase-1, IL-1β, and gasdermin D (GSDMD) (Fig. 5a, 5b). Intriguingly, while we observed a decrease in NLRP3 levels, the treatment of cells with **Compound 3** triggered the appearance of a high molecular weight NLRP3 smear, consistent with ubiquitylation (Fig. 5b).

**Fig. 5.**
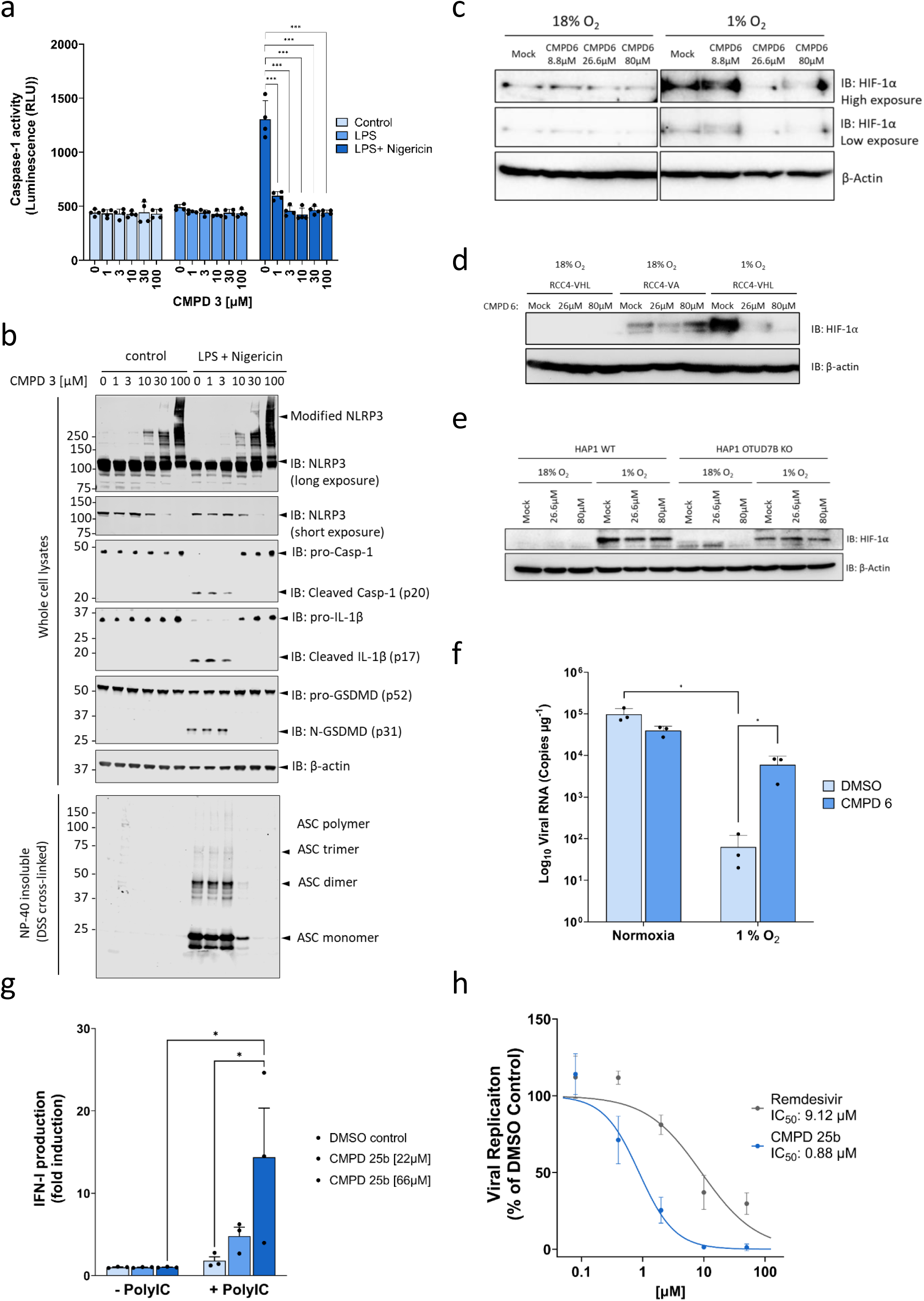
Functional validation of compounds 3, 6 and 25b. **a**, Caspase-1 activity in the media (supernatant) of PMA-differentiated THP-1 cells treated with indicated concentrations of **Compound 3**, determined using Caspase-Glo 1 luminescence assay. **Compound 3** incubation for 4 h, ± 1 µg/mL LPS incubation for 4 h and 10 µM Nigericin incubation for 45 min (error bars = SD). One-way ANOVA was used. ***P < 0.001. **b**, Immunoblots of NLRP3, Caspase-1 (Casp-1), and gasdermin d (GSDMD) from whole cell lysates of PMA-differentiated THP-1 cells treated with indicated concentrations of **Compound 3**. Immunoblot of ASC oligomerisation of the NP40-insoluble fraction of PMA-differentiated THP-1 cells after treatment with the indicated concentrations of **Compound 3** and DSS cross-linking. **Compound 3** incubation for 4 h, ± 1 µg/mL LPS incubation for 4 h and 10 µM Nigericin incubation for 45 min. **c**, Immunoblot of HIF-1a in Calu-3 cells incubated in 18% or 1% O2 for 24 h followed by treatment with increasing doses of **Compound 6** for 1h. **d**, expression of HIF-1a in RCC4-VA (Mutant VHL) or –VHL (Wt VHL) cells treated with increasing doses of **Compound 6**. Lysates from hypoxic RCC4-VHL cells treated with **Compound 6** were also performed to induce HIF-1a expression. **e**, HAP1-WT and OTUD7B KO cells were incubated in 18% or 1% O2 for 24 h followed by treatment with indicated doses of **Compound 6** for 1 h. HIF-1α expression was then determined by immunoblot in WT and KO cells. **f,** Calu-3 cells were infected with SARS-CoV-2 (VIC02/20) at a multiplicity of infection (MOI) of 0.01 for 1h. The inoculum was removed and cells incubated at 18% or 1% O2 for 24 h treated with or without 26.6 µM of **Compound 6**. Viral replication was quantified by qPCR of intracellular viral RNA and expressed as copies µg-1 of total cellular RNA. Data is representative of three biological replicates and data presented as mean -/+ S.D. Significance from Mann-Whitney test, * = p < 0.0332. **g**, Interferon I (IFN-I) production by HAP1 cells treated with **Compound 25b**/DMSO for 1 hour, followed by poly(I:C) (0.5 μg/mL) for 48 h. HEK293 IFN reporter cells were incubated with HAP1 media for 24 h, and luminescence determined with the One-Glo luciferase assay system. Data is representative of three replicates with data presented as mean -/+ SEM. Significance from 2-way ANOVA, with Tukey’s multiple comparisons test, * = p < 0.0332 **h**, Calu-3 cells were infected with SARS-CoV-2-mNeonGreen reporter virus at a MOI of 0.1 and treated with a range of doses of the USP5 inhibitor **Compound 25b** or Remdesivir^86^. Data is representative of three biological replicates and data presented as mean -/+ SEM. Dose response curves were determined using a non-linear regression analysis to calculate IC50 values for each compound.

### Compound 6 Induces HIF-1α Degradation via OTUD7B Inhibition

OTUD7B has been shown to regulate Hypoxia-Inducible Factor 1 (HIF-1α) protein homeostasis in hypoxia in a von Hippel-Lindau disease tumour suppressor (VHL)- and proteasome-dependent manner^35^. In hypoxic Calu-3 cells, **Compound 6** reduced HIF-1α in a dose-dependent manner (Fig. 5c). In renal carcinoma cells lacking active VHL (RCC4-VA)^36^, we observed that treatment with **Compound 6** had no impact on HIF-1α expression. When wild-type VHL (RCC4-VHL) expression was restored, however, a dose-dependent reduction in hypoxic HIF-1α expression was observed (Fig. 5d). In HAP1 WT cells, treatment with **Compound 6** also resulted in a dose-dependent reduction in HIF-1α expression. Importantly, HIF-1α levels were markedly reduced in OTUD7B KO cells compared to WT, consistent with published findings^35^. Treatment of KO cells with **Compound 6** did not further reduce HIF-1α, supporting an on-target mechanism (Fig. 5e). In line with reported work^37^, we demonstrated that incubation of infected Calu-3 cells in hypoxia restricted replication of SARS-CoV-2, yet treatment with **Compound 6** limited the antiviral impact of hypoxia (Fig. 5f), consistent with limited HIF-1α expression through inhibition of OTUD7B.

### Compound 25b is a Potent Anti-Viral Against SARS-CoV-2

The functions of USP5 in viral replication have been linked to type I interferon (IFN-I) production by regulating the ubiquitylation of different innate immune factors such as RIG-I and IRF3^38, 39^. In agreement with these observations, treatment of cells with the synthetic RNA Poly I:C induced IFN-I production, which was significantly amplified by co-treatment with **Compound 25b** (Fig. 5g)^39, 40^. **Compound 25b** exhibited a dose-dependent reduction in viral replication against SARS-CoV-2 in infected Calu-3 cells. In comparison to remdesivir, a guanosine nucleoside analogue which targets the viral replicase complex and is in clinical use as a broad-spectrum antiviral medication, **Compound 25b** showed a ∼10-fold increase in potency (Fig. 5h). Interestingly, **Compound 25b** did not show any significant toxicity at concentrations as high as 200 µM (Extended Data Fig. 7a).

## Discussion

As interest in DUBs as drug targets grows^6, 40, 41^, so does the need for better methodologies to identify DUB inhibitors. High-throughput screening (HTS) has been instrumental in DUB drug discovery by enabling the rapid evaluation of extensive compound libraries^28, 42^. However, HTS is typically limited by high costs, low hit rates, reliance on less disease-relevant recombinant proteins, and *in vitro* assays that may not reflect physiological conditions. It also typically requires extensive compound optimisation and follow-up validation due to identification of false and/or not selective hits.

To address these challenges, artificial intelligence (AI) and activity-based proteomics have emerged as promising alternatives. AI-enabled virtual screening approaches, exemplified by AtomNet, streamline early-stage discovery by computationally assessing compound-target interactions prior to synthesis, even in the absence of prior ligand data^43^. Similarly, using dedicated activity proteomics strategies for the unbiased identification and mapping of reactive cysteine residues in the human proteome has allowed directed, more efficient drug design for cysteine-containing proteins, including DUBs^44–47^. Importantly, recent progress in DUB ABPP has produced high-throughput-ready platforms, unlocking new opportunities for small-molecule screening^18, 20, 48^.

Here, we describe the *ob*ABPP-HT* activity proteomics workflow. *ob*ABPP-HT* is cost-effective, requiring no specialised equipment, however, it can seamlessly integrate with automation systems like the Thermo KingFisher Apex and Agilent Bravo, and is compatible with different probe types, enabling profiling of enzyme classes other than DUBs. This high-throughput and highly sensitive method enables the analysis of ∼100 samples per day using EvoSep timsTOF LC-MS/MS.

As a proof-of-concept, we screened for mechanistically novel DUB inhibitors by combining *ob*ABPP-HT* with a curated small-molecule library composed of molecules that bear electrophilic warheads for potential covalent reaction with cysteine residues, including αβ,α′β′-diepoxyketones (DEKs), γ-lactams, and ubiquitin-derived peptide inhibitors specifically designed for this study. This screen showed a hit rate ∼10-times higher than traditional DUB HTS and, after hit validation, identified new inhibitors against USP47, OTUD7B, and USP5. The potential of these DUBs as drug targets has not yet been fully explored; therefore, the potency, cellular permeability, selectivity, and functional effects of the identified DUB antagonists was further characterised.

USP47 is an attractive therapeutic target^49–51^. However, reported USP47 inhibitors also inhibit USP7 with comparable potency^30, 31^, attributable to their high homology^29, 32^. We identified **Compound 3**, a first-in-class USP47 inhibitor with preference for USP47 over USP7, an observation which was confirmed using orthogonal kinetic studies, and characterised through HDX-MS-based mapping and molecular docking studies, revealing the molecular basis for selective USP47 inhibition. Importantly, USP47/7 inhibition and knockdown has previously been demonstrated to prevent inflammasome activation^32^, a finding that is recapitulated by **Compound 3** treatment.

Although roles of OTUD7B in disease remain to be comprehensively investigated^52^, evidence suggests that OTUD7B is a potential therapeutic target^35, 53–55^. This interest is highlighted by recent reports detailing the development of OTUD7B inhibitors^48, 56^. We identified **Compound 6**, a potent, permeable, stable and selective OTUD7B inhibitor which was able to reduce HIF-1α levels under hypoxic conditions^37^, phenocopying OTUB7B knockdown studies^35^. Recently, hypoxia has been described to impact the ability of SARS-CoV-2 to infect cells in a HIF-1α-dependent manner^37, 57^. In line with its effects on HIF-1α stability, **Compound 6** rescued SARS-CoV-2 infectivity in hypoxia^57^.

USP5 catalyses hydrolysis of free poly-ubiquitin chains, with its zinc finger ubiquitin binding domain (ZnF UBP) containing a pocket to bind the C-terminal di-glycine motif on ubiquitin^58^. Although USP5 is a potential target for treatment of cancer, pain, and inflammation^59^, only limited structure-activity relationship studies on USP5 inhibitors have been reported (Fig. S1) and lack characterisation in a cellular context^60^. Existing USP5 inhibitors were designed based on the Ub C-terminus and thus contain a carboxylate group to mediate binding to the USP5 ZnF^60, 61^. By contrast, we achieved selective USP5 inhibition using Ub-derived inhibitors in which the C-terminal glycine is substituted for a propargyl group, based on the observation that Ub-PA can covalently react with nucleophilic DUB cysteine thiols^54^. The identified **Compound 25** exhibits high levels of selectivity for USP5 inhibition over related DUBs USP16 and USP22^62^. That relatively simple substrate-derived molecules like **Compound 25** exhibit high selectivity for individual DUBs is remarkable, highlighting subtle but critical differences in active site architecture. **Compound 25** likely covalently modifies the nucleophilic cysteine of USP5 via its alkyne warhead, as supported by reported crystal structure of USP5 in complex with a Ub-derived probe (PDB: 3IHP^63^), the structures of related USPs with Ub-PA, and the observed reduced inhibition upon steric hindrance by introducing sterically bulky substituents near the alkyne. The selectivity of **Compound 25** for USP5 may arise from USP5’s unique domain architecture. A disulphide bridge between C793 (BL2) and C195 (cUBP) in the USP5-Ub complex may limit BL2 flexibility, positioning cUBP near the binding site and enabling additional compound interactions. C195 may also affect dimerization, further shaping compound engagement. The use of a latent electrophile for covalent reaction with the active site cysteine may also result in improved USP5 selectivity, potentially due to the differential binding kinetics of **Compound 25** and derivatives with DUBs^64–67^. Note that small molecules bearing electronically activated alkynes, e.g., propiolamides, for selective covalent reaction with a cysteine residue of Bruton’s tyrosine kinase (Btk) are in therapeutic use^68, 69^, highlighting the potential of our USP5 inhibitor design. Structure–activity studies of **Compound 25** yielded fluorinated, cell-permeable analogues **25a** and **25b**, active in cells and suitable for USP5 inhibition studies. **Compound 25b** boosts type I interferon production and shows potent antiviral activity, supporting known USP5 functions.

The combined results manifest the potential of combining cysteine-reactive, warhead-containing compound sets with the *ob*ABPP-HT* workflow to streamline development of selective DUB inhibitors for drug development and for use as biochemical tools in functional assignment and mechanistic studies.

## Online Methods

### Small-molecule synthesis

αβ,α′β′-Diepoxyketones (DEKs) (**Compounds 1-8**) and γ-lactams (**Compounds 9-23**) were synthesised as reported^26^. The synthesis of substrate-based small-molecules (**25**-**31 and 25a-k**) is described in the Supplementary Information. The structures of **Compounds 1-31** are shown in the Supplementary Information (Table S1-3) and the structures of **Compounds 25a- k** are detailed in the Extended Data Table 1.

### Cell culture

MCF-7 cells were cultured in high glucose DMEM supplemented with 10% fetal bovine serum (FBS) (v/v) and L-glutamine (2 mM). Cells were maintained at 37 °C, 5% CO2. MCF-7 cells were washed with phosphate buffered saline (PBS) and collected in PBS by scraping and centrifugation at 200 *xg* for 10 min. For data shown in Extended data Figure 1, MCF-7 cells were collected by the addition of TrypLE followed by media and centrifuged at 1600 rpm for 5 min with pellets being resuspended in PBS (3 cycles). HAP1 cells were cultured in IMDM media (#21980065) supplemented with 10% FBS (v/v), and maintained at 37 °C, 5% CO2. ∼3 × 10^6^ cells were seeded per 10 cm plate in 10 mL media overnight. After seeding, 4 mL of media were aspirated from each plate, and recombinant human interferon-alpha, IFN-α2 (α2b; PBL Assay Science; Cat. No. 11105-1), was added to the remaining media (1000 units/mL final concentration). IFN-α treatment was performed for 24 h (to induce USP18), media from each plate was then aspirated and cells were washed once with PBS (5 mL per plate). Cells were subsequently stored at –80 °C until lysis. THP-1 cells were cultured in RPMI 1640 supplemented with 10% FBS (v/v), 0.05 mM β-mercaptoethanol, GlutaMAX Supplement, and Penicillin-Streptomycin. Cells were maintained at 37 °C 5% CO2. RCC4-VHL and VA cells were cultured in DMEM media supplemented with 10% fetal bovine serum (FBS), GlutaMAX, and Penicillin-Streptomycin^36^. Calu-3 cells were obtained from Prof Nicole Zitzmann’s lab and maintained in Advanced DMEM media, 10% FCS, L-glutamine, and Penicillin-Streptomycin.

### In-depth ABPP workflows

#### Sample preparation

Cell lysis and DUB Activity-Based Enrichment: Cells were lysed in lysis buffer (50 mM Tris, pH 7.5, 5 mM MgCl2, 0.5 mM EDTA, 1 mM DTT and 250 mM sucrose) containing acid-washed glass beads (Sigma Aldrich #G4649). Glass beads made up approximately one third of the total lysis buffer volume. Cell lysates were vortexed and centrifuged (14K x g, 4 °C, 25 min) to separate the pellet, glass beads, nuclei, and membranes from supernatant. The supernatant was transferred to Eppendorf tubes and the pellets were discarded.

Proteins extracted from ∼1·10^7^ cells were used as starting materials. When profiling a DUB inhibitor, PR-619 (Sigma-Aldrich # 662141) was first incubated and mixed at 1,000 rpm at 37 °C for 1 h. 25 μg of a DUB probe (HA-Ahx-Ahx-Ub-PA (UbiQ-078), HA-Ahx-Ahx-Ub-VME (UbiQ-035), HA-Ahx-Ahx-Ub-VS (UbiQ-178)) or probe mixtures were added to each of the samples and mixed at 1,000 rpm at 37 °C for 1 h. The reaction was quenched by the addition of SDS to 0.4% (v/v) and NP-40 to 0.5% (v/v). Lysate mixtures were then diluted to 1 mL, 0.5 mg/mL with NP-40 lysis buffer (pH 7.4, 50 mM Tris, 0.5% (v/v) NP-40, 150 mM NaCl, and 20 mM MgCl2). 200 μL of anti-HA-Agarose (Pierce # 26182) slurry, previously washed three times with NP-40 lysis buffer, was added to the samples and incubated on a rotator overnight at 4 °C. After a first centrifugation step (2,000 g, 4 °C, 1 min), beads were washed four times with 500 μL of NP-40 lysis buffer. Protein complexes were eluted by boiling beads in 250 μL SDS Laemmli 2X sample buffer.

Methanol-chloroform precipitation was performed prior to protease digestion. For each sample, a 4x volume of methanol was added and vortexed for 5 s. A 1x volume of chloroform was added to the samples, which were then vortexed for 5 s. A 3x volume of water was added, and samples were vortexed for 5 s. Samples were then centrifuged for 10 min at 10,000 rpm. The aqueous and organic phases were removed to leave only the protein pellet. The pellets were subsequently washed with 4x volume of methanol and centrifuged for 10 min at 10,000 rpm. The supernatant was discarded, and pellets were re-dissolved in 100 μL of water. Protein precipitation was repeated for a second time to ensure sufficient removal of detergent.

Sample digestion and TMT labelling were performed using the IST-NHS 96x kit (PreOmics # P.O.00030) according to the manufacturer’s instructions. Briefly, 100 µL lysis buffer was added to each sample and heated to 95 °C while mixing at 1,000 rpm for 10 min. Lyophilised digestion enzymes were dissolved in 100 µL water before being added to the samples. The samples were digested at 37 °C for 2 h. For samples to be analysed by a Label-Free quantitation MS method, digestion was stopped, and the sample was transferred to the cartridge in a centrifuge at 2000 g for 1 min before proceeding the washing steps with wash solvent 1 and 2 provided with the kit. For the remaining samples, 200 µg TMTpro 18 Plex reagents (ThermoFischer Scientific # A52045) were added to each of the samples and incubated for 1 h while shaking at 1000 rpm. A total of 10 µL of 5% (v/v) aqueous hydroxylamine was added to each of the samples. A total of 100 µL of stop reagent was also added to the samples before transferring to cartridge for washing with washing solvent 1 and 2 provided with the kit. Samples were eluted with 250 µL of 50% (v/v) acetonitrile in 0.01% (v/v) aqueous formic acid. Samples labelled with TMT reagents were combined, dried and subjected to high pH fractionation using HPLC (Agilent). Peptides were loaded into a XBridge C18 column 4.6 x 250 mm (Waters # PN186003117) in buffer A (H2O, pH 10) and eluted with increasing buffer B (90% (v/v) aqueous acetonitrile, pH 10) at a 0.50 mL/min flowrate over a 96 min gradient. Eluted samples were collected every minute and concatenated to give a total of 24 fractions. Desalting was performed by solid phase extraction (SPE) employing reversed-phase C18 cartridges (Sep-Pak C18 light).

#### LC-MS/MS methodology

For LFQ Data Dependent Acquisition (DDA) and Data independent Acquisition (DIA) workflows, peptides were analysed by liquid chromatography tandem mass spectrometry (LC-MS/MS) using an Ultimate 3000 UHPLC coupled to an Orbitrap Fusion Lumos Mass Spectrometer (both from ThermoFisher). Tryptic peptides were separated on an EASY-spray PepMap column (2um, 75um x 50cm; ThermoScientific) using a 60-minute linear gradient from 2 to 35% buffer B (5% DMSO, 100% acetonitrile, 0.1% TFA) with a 250nl/min flow and analysed on the Orbitrap Fusion Lumos (Thermo Scientific). Data were acquired in DDA and DIA as previously described^16, 70, 71^. In both cases the advance peak detection (ADP) was enabled. For DDA, survey scans were acquired in the Orbitrap at 120 k resolution over a m/z range 400 -1500, AGC target of 4e5 and S-lens RF of 30. Fragment ion spectra (MS/MS) were obtained in the Ion trap (rapid scan mode) with a Quad isolation window of 1.6, 40% AGC target and a maximum injection time of 35 ms, with HCD activation and 28% collision energy. For DIA, MS1 scans were acquired in the Orbitrap with 120k resolution (m/z 350-1650), an AGC target of 5e5 and a maximum injection time of 20ms. The full MS events were followed by 40 DIA scan windows per cycle (with variable isolation width) covering the m/z range from 350 – 1650. The MS/M S scans were acquired in the orbitrap at 30k resolution and normalised HCD set up at 30%.

For TMT labelled samples, mass spectrometric analyses were carried out with a Proxeon Easy Nano-LC fitted with a PepMap C18 column 75 µm x 50 cm (Thermos Scientific #PN ES903) and connected to a Lumos Orbitrap mass spectrometer (Thermo Scientific). Peptides were eluted at 450 nL flowrate during 120 min gradient. MS data was acquired with SPS M3 approach. Each full orbitrap scan (R=60K, m/z scan range: 385-1350, AGC target: 100k, max. injection time: 100 ms) was acquired and followed with dependent IT MS/MS scans (isolation window: 0.5, CID collision energy (%): 35, AGC target: 30k ions, max. injection time: 175 ms) for a 2.7S cycle time. SPS MS3 scans were acquired with the following parameters (R: 50k, isolation window = 1.5, AGC target: 120k, HCD collision energy (%): 55, max. injection time: 100 ms).

#### LC-MS/MS data analysis Single shot LFQ

Data was searched using a Uniprot *Homo sapiens* database containing isoforms (42,469 entries, retrieved on 15/08/2023). Software versions: DIA-NN 1.8.1, Fragpipe 22.0, MaxQuant/MaxDIA 2.6.1.0. All search settings were left as default, with match between runs enabled and cysteine carbamidomethylation included as a fixed modification. Each search was 3 x HA-Ub-* and 3 x HA-Ub-*+ 50 µM PR619. The PA warhead alone was searched separately from the PA/VME/VS cocktail of warheads. The intensities output ‘report.pg_matrix’ was used from DIA-NN, the LFQ intensities from ‘combined_protein’ output was used from Fragpipe, and the LFQ intensities from the ‘protein groups’ was used from Maxquant for both DDA and DIA searches.

#### LC-MS/MS data analysis TMT fractions

MS data processing was performed with Proteome Discoverer Version 2.4.0.305 (Thermo Scientific) with UniProtKB Swiss-Prot (TaxID = 9606, *Homo sapiens*) fasta. Peptide identifications were carried out using the Sequest™ search engine with the following parameters: precursor tolerance: 10 ppm, fragment tolerance = 0.6 Da, max. missed cleavage sites: 3, dynamic modifications: M oxidation and N/Q deamination, fixed modifications: PreOMics cysteine modification (+113.084 Da) and TMTpro (+304.207 N-terminus and K).

### High-throughput ABPP workflows

#### Sample preparation for Agilent Bravo ABPP high-throughput screen

The data presented here on the Agilent Bravo ABPP immunoprecipitation is from published work,^20^ and is included for comparison.

#### Sample preparation for *ob*ABPP-HT* ABPP high-throughput screen and compound concentration-dependence validations

HA-Ub-PA was synthesised as reported.^20, 22^ MCF-7 cells were washed with PBS, scraped in PBS and centrifuged at 200 *g* for 10 min. Pellets were then resuspended in lysis buffer (50 mM Tris, 5 mM MgCl2, 0.5 mM EDTA, 250 mM sucrose, 1 mM dithiothreitol (DTT), pH 7.5), and lysed using glass beads (2:1 ratio lysate to beads) with vortexing (10 cycles of 30 s vortexing followed by 2 min incubation on ice). The MCF-7 lysate was then clarified at 600 *x g* for 10 min at 4 °C. Lysate protein concentration was determined by BCA, and 250 µg was aliquoted per well, into a 96 well plate, at a concentration of ∼4.7 mg/mL.

The compound library was diluted to 1 mM DMSO stock solutions in a 96 well plate and multichannel-pipetted with mixing into the lysate plate to a final compound concentration of 25 µM in duplicate for the screen. For the concentration-dependence validations, the indicated inhibitor concentrations were added in duplicate. Additional control wells (4 x negative, 4 x positive for the screen, 3 x negative and 3 x positive for the concentration-dependence validations) were treated with DMSO. The plate was then incubated at 37 °C for 1 h on a thermomixer and without shaking.

A total of 2 µg of HA-Ub-PA was multichannel-pipetted with mixing into to the positive control wells and all inhibitor-treated wells, with an equivalent volume of buffer added to the negative control wells. The lysates were then incubated with the HA-Ub-PA at 37 °C for 45 min on a thermomixer without shaking. The final reaction volume with the MCF-7 lysate, inhibitor/DMSO and HA-Ub-PA/buffer was 60 µL. Reactions were quenched by the addition of SDS and NP-40 (IGEPAL CA-630) to final concentrations of 0.4 % (v/v) and 0.5 % (v/v) respectively. Reactions were then diluted by addition of 200 µL NP40 buffer (50 mM Tris, 0.5% NP-40 (IGEPAL CA-630), 150 mM NaCl, and 20 mM MgCl2·6 H2O, pH 7.4).

75 µL of anti-HA magnetic beads were aliquoted into a separate 96 well plate and washed 5x with 200 µL of NP40 buffer (50 mM Tris Base, 0.5% NP-40 (IGEPAL CA-630), 150 mM NaCl, and 20 mM MgCl2·6 H2O, pH 7.4), using a magnetic rack for bead separation and multichannel pipetting with cut tips for mixing. Reaction mixtures were then added to the washed magnetic beads, mixed by multichannel-pipetting, and incubated in a thermomixer overnight at 4 °C with gentle agitation (350 rpm). The beads were then washed 5x with 200 µL of NP40 buffer as before.

Proteins were eluted from the beads and reduced by addition of 100 µL of 2 x Laemmli buffer supplemented with 20 mM DTT using a multichannel pipette, and incubated for 10 min at 90 °C with 350 rpm on a thermomixer. Eluates were then removed from the beads using a magnetic rack and transferred to a fresh 96 well plate. Eluates were alkylated by the addition of iodoacetamide (IAA) to a final concentration of 40 mM. Proteins were then digested using trypsin and cleaned up using a 96 well S-trap plate according to the manufacturer’s instructions^72^, dried down in a speedvac and resuspended in 0.1% (v/v) aqueous formic acid.

#### Sample preparation for the ABPP Thermo KingFisher Apex workflow

The bio-ISG15-PA probe was synthesised as previously described^73^. Cell lysis, subsequent labelling with HA-Ub-PA or bio-ISG15-PA probes, immunoprecipitation and protein enrichment, and processing of samples for analysis by western blot and LC-MS/MS was performed as described^74^. Briefly, HAP1 cells were treated and prepared as described in the subsection ‘*Cell culture’*. Cells were resuspended in lysis buffer (50 mM Tris, 5 mM MgCl2, 0.5 mM EDTA, 2.5% (v/v) glycerol, 1 mM dithiothreitol (DTT), pH 7.5), and lysed using glass beads (2:1 ratio lysate to beads) with vortexing (10 x 30 s vortexing followed by 1 min incubation on ice). The HAP1 lysates were then centrifuged (600 x *g* for 5 min at 4 °C; ∼80% of supernatant was transferred into clean tubes, then the remaining ∼20% of supernatant was spinned at 14 000 x *g* for 10 min at 4 °C, and then the two fractions from each spin were combined). The protein concentration in the lysates was estimated using the BCA assay kit (#23227), and 270 µg were aliquoted per well into a 96-deep well plate. The total volume in each well was made up to 143 μL for samples to be labelled with HA-Ub-PA, or 147 μL for samples to be labelled with bio-ISG15-PA, with lysis buffer.

For **Compound 3**, stock solutions (100×) were prepared at a concentration of 0.46, 1.39, 4.16 or 12.5 mM in DMSO in a 96-well plate, and 1.5 μL of each stock were transferred into the respective wells using a multichannel pipette (resulting in a final concentration of 4.6, 13.9, 41.6 or 125 μM) in duplicates. Negative and probe-only positive controls were treated with 1.5 μL of DMSO in triplicates. The plate was then incubated at rt for 1 h without shaking.

Following compound incubation, 5.4 µL of HA-Ub-PA or 2.35 µL of bio-ISG15-PA (volumes determined from probe titration assays for protein labelling) were transferred into the respective wells using a multichannel pipette, with an equivalent volume of buffer added to the corresponding negative control wells, and incubated at 37 °C for 45 min without shaking. The final reaction volume with the HAP1 lysate, inhibitor or DMSO, and HA-Ub-PA or bio-ISG15-PA or buffer was 150 µL. Reactions were quenched by addition of SDS and NP-40 (IGEPAL CA-630) to final concentrations of 0.4% (v/v) and 0.5% (v/v), respectively, from a master mixture. The master mixture was prepared immediately before quenching by mixing a 5% (v/v) SDS stock with a 10% (v/v) NP-40 stock in a ratio of 1.6:1 and adding 21.5 μL of the mixture to each well. 20 μL of each reaction mixture was kept as ‘input’ samples (to evaluate efficiency of probe labelling), and the remaining ∼150 μL of reaction volume was diluted by addition of 250 µL NP-40 buffer (50 mM Tris Base, 0.5% (v/v) NP-40 (IGEPAL CA-630), 150 mM NaCl, and 20 mM MgCl2, pH 7.4) to a total volume of 400 μL.

For immunoprecipitation/pull-down steps, 50 µL of anti-HA magnetic beads or 37.5 µL of streptavidin magnetic Sepharose beads were used per sample. The appropriate total amount of anti-HA magnetic beads or streptavidin magnetic Sepharose beads were split into 450 µL aliquots, washed with 4 × 1.6 mL of NP-40 buffer (50 mM Tris, 0.5% NP-40 (IGEPAL CA-630), 150 mM NaCl, and 20 mM MgCl2, pH 7.4) using a magnetic rack for bead separation, and transferred into the respective reaction wells using a multichannel pipette with cut tips. The 96-deep well plate was then incubated for 4 h. at 4 °C on the Thermo KingFisher Apex system, with mixing speed set to ‘medium’. The beads were then washed with 4 × 500 μL NP-40 buffer, as described above, and proteins were eluted from the beads twice with 2× Laemmli buffer supplemented with 50 mM DTT (2 × 55 μL, 2 × 5 min at 95 °C with mixing speed set to ‘slow’). Beads were removed from the eluates by programming the system to capture and release the beads into an empty 96-deep well plate. Eluates were reduced with DTT to a final concentration of 10 mM (15 min, rt), then alkylated by the addition of iodoacetamide (IAA) to a final concentration of 20 mM (10 min, rt). Proteins were then digested using trypsin and cleaned up using a 96-well S-trap plate according to the manufacturer’s instructions^72^, dried down in a speedvac and resuspended in 0.1% (v/v) aqueous formic acid.

#### LC-MS/MS timsTOF *ob*ABPP-HT* methodology

Peptide samples were loaded onto Evotips (Evosep) desalting columns according to the manufacturer’s instructions, then chromatographically separated on an 8 cm x 150 µm analytical column with bead size 1.5 µm (EV1109, Evosep), using an Evosep One liquid chromatography (LC) system with the 100 spd standard method. Peptides were eluted onto a TimsTOF Pro mass spectrometer (Bruker) operated in diaPASEF mode using 8 diaPASEF scans per TIMS-MS scan. The ion mobility range was set to 0.85-1.3 Vs/cm^2^. Each mass window isolated was 25 m/z wide, ranging from 475-1000 m/z with an ion mobility-dependent collision energy that increased linearly from 20 eV to 59 eV between 0.6-1.6 Vs/cm^2^.

#### LC-MS/MS for timsTOF Bravo Agilent robot methodology

The previously reported timsTOF data collected after immunoprecipitation using the Agilent robot sample prep methodology is identical to that detailed above^21^, other than a different ion mobility range (0.6–1.6 Vs/cm^2^), and mass range (400–1000 m/z).

#### LC-MS/MS for timsTOF for KingFisher Apex robot methodology

Peptides were analysed by nanoLC-MS/MS using an Evosep One LC system coupled with a timsTOF HT (Bruker) equipped with an 8 cm x 150 µm, 1.5 µm analytical column (Evosep). 200 ng peptides were separated using the Evosep 60SPD workflow (Analytical solvents A: 0.1% FA and B: acetonitrile plus 0.1% FA). Column was held at 40 °C. Data were acquired in data-independent acquisition PASEF mode with the following settings: m/z range from 100 m/z to 1700 m/z, ion mobility range from 1/K0 = 1.30 to 0.85 Vs/cm2 using equal ion accumulation and ramp times in the dual TIMS analyser of 100 ms each. Each cycle consisted of 8 PASEF ramps covering 21 mass steps each with 25 Da windows each with 2/3 non-overlapping ion mobility windows covering the 475 to 1000 m/z range and 0.85 and 1.26 Vs/cm2 ion mobility range. The collision energy was lowered as a function of increasing ion mobility from 59 eV at 1/K0 = 1.6 Vs/cm2 to 20 eV at 1/K0 = 0.6 Vs/cm2.

#### ABPP-HT* *vs ob*ABPP-HT* comparison LC-MS/MS data analysis

Data was searched in DIA-NN 1.8.1 with search settings left as default and match between runs enabled. Data was searched using a Uniprot *Homo sapiens* database containing isoforms (42,469 entries, retrieved on 15/08/2023). N=3 for HA-Ub-PA positive controls, N=1 for negative controls with no HA-Ub-PA present. The ‘report.pg_matrix’ output was used for analysis.

#### ABPP Thermo KingFisher Apex workflow LC-MS/MS data analysis

Data was searched in DIA-NN 1.8.1 with search settings left as default and match between runs enabled. Data was searched using a Uniprot H*omo sapiens* database containing isoforms (20412 entries, retrieved 13/12/2023). The ‘report.pg_matrix’ output was used (identical number of DUBs were identified in the ‘report.unique_genes_matrix’ output). DUBs were filtered to select hits >1.5-fold enriched in the positive HA-Ub-PA control *vs* the negative control without HA-Ub-PA. Volcano plots were generated/calculated using GraphPad Prism (Version 10.2.3). For the **Compound 3** concentration-dependence heatmap DUBs were filtered for >5-fold enrichment in the positive HA-Ub-PA control *vs* the negative control without HA-Ub-PA. USP6 was discounted due to it’s unreliable detection across replicates.

#### *ob*ABPP-HT* screen and concentration-dependence data analysis

Data was searched in DIA-NN 1.8.1 with search settings left as default and match between runs enabled. Data was searched using a Uniprot *Homo sapiens* database, not containing isoforms (20,416 entries, retrieved on 15/02/2023). The ‘report.unique_genes_matrix’ output was used to ensure inhibitor identification unique to specific DUBs. DUBs were filtered to remove those that were not >5-fold enriched in the positive HA-Ub-PA control *vs* the negative control without HA-Ub-PA. For the concentration-dependent data the following DUBs were discounted due to their unreliable detection across replicates, likely attributable to their low intensities: OTUD1, USP42, OTUD5, USP1, USP30, USP33, USP37, USP43 and OTUB2. Intensities were normalised as a percentage of the positive HA-Ub-PA control and for the screen multiple two-tailed unpaired (homoscedastic) T-tests were carried out (Microsoft Excel) to identify DUBs with significantly different intensities relative to the positive control. The resultant data were then filtered to select hits with a >20% average reduction in DUB activity. The concentration-dependence ABPP outputs were processed using CurveCurator (Version 0.3.0),^75^ with the alpha asymptote set to 5%.

### ABP western blot compound validations

#### *ob*ABPP-HT* screen hit validations

For positive hit validations by western blot the *ob*ABPP-HT* screen and Thermo KingFisher Apex methodologies were used with the concentrations of compounds indicated in figures. For the *ob*ABPP-HT* screen, following compound incubation and reaction with HA-Ub-PA, the reaction was quenched directly with Laemmli buffer supplemented with 100 mM DTT and analysed by western blot. IC50 values were extracted by quantifying the HA-Ub-PA labelled band by densitometry (Image studio lite) normalised to the β-actin signal and subsequently normalising the signal of the DUB activity to the negative and positive HA-Ub-PA controls. For the Thermo KingFisher Apex workflow, following compound incubation and reaction with HA-Ub-PA, the reaction was quenched directly with 2× Laemmli buffer supplemented with 50 mM DTT and analysed by western blots. Concentration-dependencies were fit to Y=100/(1+10^((LogIC50-X)*HillSlope)) in Graphpad Prism (Version 10.1.1).

#### Compound *vs* HA-Ub-PA time dependence competition studies

Samples were processed as for the ABP western blot compound concentration-dependence validations, with the HA-Ub-PA incubated for the indicated time.

#### ABP compound cellular permeability studies

MCF-7 cells were cultured in 6-well plates to 90% confluence, with a media change to 1 mL prior to DMSO/compound treatment at the indicated concentration for 4 h. Cells were then washed 3 x with PBS, and collected by scraping in PBS and centrifuged (200 x *g* for 10 min). Pellets were then processed as outlined above in the ABPP screen sample preparation section, with 0.4 µg of HA-Ub-PA being used per 50 µg of lysate protein. The mixture was incubated for 45 min at 37 °°C, before being quenched by addition of Laemmli buffer supplemented with 100 mM of DTT; the mixture was then analysed by western blotting.

### Enzyme kinetics

Fluorescence intensity measurements were used to monitor the cleavage of a ubiquitin-rhodamine substrate. All activity assays were performed in black 384-well plates in assay buffer (20 mM Tris, pH 8.0, 150 mM potassium glutamate, 0.1 mM TCEP and 0.03% Bovine Gamma Globulin) with a final assay volume of 20 μL. USP7, 1 nM (USP7 (1-1102), DU15644, MRC Protein Phosphorylation and Ubiquitylation Unit) or USP47, 10 nM (USP47 (N-terminal His), DU15682, MRC Protein Phosphorylation and Ubiquitylation Unit) was added and preincubated with **Compound 3** for 30 mins. 180 nM ubiquitin-rhodamine 110 (Ubiquigent) was added to initiate the reaction and the fluorescence intensity was recorded for 30 mins on a PherastarFSX (BMG Labtech) with an Ex485/Em520 optic module. Initial rates were plotted against compound concentration to determine IC50. To investigate the time-dependence of the potency of **Compound 3**, activity assays were repeated without a preincubation step.

### HDX-MS

#### Sample Preparation

Hydrogen Deuterium eXchange Mass Spectrometry (HDX-MS) experiments were performed on the same stock of recombinant USP7 and USP47 protein constructs as those used in the *in vitro* enzyme kinetics experiments. USP7 was provided at a concentration of 25.3 μM and USP47 at 16.5 μM, respectively. Both proteins were prepared in 50 mM HEPES, pH 7.5, 10% glycerol, 150 mM NaCl, 1 mM DTT. The initial stock concentration of **Compound 3** was 30 mM in DMSO. In solution HDX-MS was performed by preparing a volume-to-volume mixture at a molar ratio of 1:2 of protein:**Compound 3** (*i.e*., the holo-state). Samples were diluted to achieve a final concentration of 16 pmol on column per reaction for USP7 and 21 pmol on column per reaction for USP47. Equivalent control samples (*i.e*., the apo-state) were prepared where DMSO was supplemented in place of the compound.

#### Data Acquisition

HDX-MS was performed as previously described^76^. Briefly, an identical labeling buffer was prepared as that of the protein stocks except in deuterium oxide D2O (99+ %D, Cambridge Isotope Laboratories, Tewksbury, MA) and with the 10% glycerol was removed (a final composition of 50 mM HEPES, pH 7.5, 150 mM NaCl, 1 mM DTT). The pH of the labeling buffer was measured and corrected to pD (pD=pH+0.4). The quenching buffer comprised of 2M Guanidine HCl, 100 mM citric acid, pH 2.3 in H2O. Proteins +/- **Compound 3** were preincubated for 30 min at room temperature to allow complexes to form. Samples were then diluted with the labeling buffer in a 1:20 ratio, achieving an excess D2O concentration of 95%). Several labeling time points were sampled at 20 ᵒC, more specifically, at 30, 600, and 3600 s). All analyses were acquired in triplicate. Non-deuterated controls were prepared in an identical fashion, with H2O in place of D2O. At the end of each labeling time point, samples were quenched upon addition of quench buffer (1:1 ratio), resulting in a final pH of 2.5. Samples were subsequently digested with a dual pepsin/proteaseXIII column (2.1 x 3.0 mm; NovaBioAssays, MA) at 8 ᵒC for 3 min. Peptides were trapped on a 1.0 mm x 5.0 mm, 5.0 µm trap cartridge (Thermo Scientific™ Acclaim PepMap100) for desalting. The flow rate was maintained at 150 µL/min. Peptides were then separated on a Thermo Scientific™ Hypersil Gold^™^, 50 x 1 mm, 1.9 um, C18 column increasing hydrophobicity achieved by a linear gradient of 10% to 40% Buffer B (A: water, 0.1% FA; B: ACN, 0.1% FA). The flow rate was maintained at 40 µL/min. A protease wash (2M guanidine, 100 mM citric acid, pH 2.3 in H2O) was performed for each run to limit carry-over. To minimise back-exchange, the quenching, trapping, and separation steps were performed at 1.5ᵒC. Labeling, quenching, and online digestion steps were performed with the aid of an automated HDX robot (Trajan Scientific and Medical). Sample preparation was managed in Chronos (version 5.4.1). All samples were acquired in MS1 mode on Thermo Scientific^™^ Orbitrap Exploris^™^ 480 Hybrid^™^ mass spectrometer.

#### Data Analysis

Before each HDX-MS experiment, an unspecific digested peptide database was created in BioPharma Finder (version 5.2) for non-deuterated USP7 and USP47 proteins through a data-dependent and targeted HCD-MS^2^ acquisition regime^76^. Labeling data were processed and manually curated in HDExaminer version 3.4.2 (Trajan Scientific and Medical). The charge state with the highest quality spectra for all replicates for each peptide across all HDX-MS labeling times was used in the final analysis. The significant differences observed at each residue was used to map HDX-MS consensus effects (based on overlapping peptides) onto the AlphaFold model.

### Caspase-1 assay

PMA-differentiated THP-1 cells were treated as indicated, and the caspase-1 activity in the culture medium (supernatant) was measured using the Caspase-Glo 1 Inflammasome Assay (Promega) according to the manufacturer’s instruction. Briefly, the Caspase-Glo 1 Reagent was prepared by resuspending the Z-WEHD substrate in Caspase-Glo 1 buffer and addition of MG132 inhibitor to a final concentration of 60 μM. Cell culture medium (50 μL) was then added to the Caspase-Glo 1 Reagent (50 μL) in a 96-well plate and the resultant mixture was incubated at room temperature for 1 h. Luminescence was measured using a FLUOstar Omega multi-mode microplate reader (BMG LABTECH) according to the manufacturer’s instructions.

### MTS cell viability assay

Cell viability was assessed using the CellTiter 96 AQueous One Solution Cell Proliferation Assay (MTS, Promega). The One Solution Reagent (20 μL) was added to each well of a 96-well plate containing 100 μL of sample in culture medium. Following incubation at 37 °C for 1 h, absorbance was measured at the wavelength of 490 nm using a FLUOstar Omega multi-mode microplate reader (BMG LABTECH). Cell viability was calculated according to the manufacturer’s instruction.

### ASC speck oligomer crosslinking

ASC oligomer chemical crosslinking was performed as previously described^70^. Briefly, PMA-differentiated THP-1 cells were primed with LPS (1 μg/mL) for 4 h, followed by stimulation with nigericin (10 μM) for 45 min and indicated concentrations of **Compound 3**. Cells were lysed on ice in Buffer A (20 mM HEPES, pH 7.5, 10 mM KCl, 1.5 mM MgCl₂, 1 mM EDTA, 1 mM EGTA, 320 mM sucrose, and protease inhibitors), and homogenised using a 21-gauge needle. Lysates were centrifuged at 300 × g for 8 min at 4 °C. The supernatants were collected, mixed 1:1 with CHAPS buffer (20 mM HEPES, pH 7.5, 5 mM MgCl₂, 0.5 mM EGTA, 0.1% CHAPS, and protease inhibitors), and centrifuged at 2,400 × g for 8 min to pellet crude inflammasome complexes. Pellets were washed twice with ice-cold PBS, resuspended in CHAPS buffer, and cross-linked with 2 mM DSS (prepared in DMSO) at 37 °C for 20 min. The reaction was quenched with NuPAGE™ LDS Sample Buffer (4X) and subjected to Western blot analysis.

### Alignment and structure prediction of *homo sapiens* and Pan *troglodytes* USP47

For kinetic and HDX MS studies of **Compound 3**, full-length *Homo Sapiens* USP7 and full-length *Pan troglodytes* isoform 2 were used. Sequences of full-length *Homo sapiens* canonical USP47 and full-length USP47 from *Pan troglodytes* isoform 2 were aligned using T-coffee multiple sequence alignment with differences reported at Extended Data Fig. 8a^77^. The sequences, including the His-tag of the recombinant *Pan troglodytes* isoform 2, were then used to predict the structures using AlphaFold 3^78^, with the highest-ranking structures being superposed using PyMOL (version 2.5.0) with an RMSD of 2.511 (Extended Data Fig. 8b and 8c).

### HIF-1α hypoxia conditions and Covid-19 infection assay

SARS-CoV-2 Australia/VIC01/2020 virus and cells were provided by the Peter Doherty Institute for Infection and Immunity, Melbourne, Australia at P1 and passaged twice in Vero/hSLAM cells (Cat#04091501) - obtained from the European Collection of Cell Cultures (ECACC), UK. Virus infectivity was determined by plaque assay on Vero-TMPRSS2 cells as previously reported^79^. Calu-3 cells were infected with the above strain of SARS-CoV-2 at an MOI of 0.01 for 2 h. Viral inocula were removed, cells washed three times in PBS and maintained in growth media until harvest. Cell lines were maintained at 37 °C and 5% CO2 in a standard culture incubator and exposed to hypoxia using an atmosphere-regulated workstation set to 37 °C, 5% CO2:1%–5% O2:balance N2 (InvivO2 200, Baker-Ruskinn Technologies). For quantification of viral RNA, total cellular RNA was extracted using the RNeasy kit (Qiagen) according to manufacturer’s instructions. Equal amounts of RNA, as determined by Nanodrop analysis, were used in a one-step RT-qPCR using the Takyon-One Step RT probe mastermix (Eurogentec) and run on a Roche Light Cycler 96. For quantification of viral copy numbers, qPCR runs contained serial dilutions of viral RNA standards. Total SARS-CoV-2 RNA was quantified using: 2019-nCoV_N1-F: 5’-GAC CCC AAA ATC AGC GAA AT-3’, 2019-nCoV_N1-R: 5’-TCT GGT TAC TGC CAG TTG AA TCT G-3’, 2019nCoV_N1-Probe: 5’-FAM-ACC CCG CAT TAC GTT TGG TGG ACC-BHQ1-3’. For assaying the antiviral response against **Compound 25b**, Calu-3 cells were infected with a SARS-CoV-2 reporter virus containing a mNeonGreen fluorescent reporter as previously described at a MOI of 0.1^80^. Infected cells were treated with serial dilutions of the compound for 24 h followed by fixation in 4% PFA. Cells were stained with DAPI and mNeonGreen fluorescence was quantified by Clariostar. Fluorescence data was normalised to the DAPI signal and plotted as relative to the DMSO control.

### Type I interferon production

Type I interferon production was performed as previously described^81^, using HEK293 cells transduced with a pGreenFire-ISRE reporter. In brief, HAP1 cells were seeded at 5,000 cells per well in a 96-well plate in DMEM containing 10% FBS and 2 mM glutamine. After attachment, cells were treated with **Compound 25b** for 1 h, followed by poly(I:C) (0.5mg/mL) for 48 h. HAP1 media was then collected, centrifuged, and transferred to HEK293 IFN reporter cells, which were seeded at 25,000 cells per well in a 96-well plate and allowed to attach. Recombinant human IFNα2 (PBL assay science) was used for standards. After 24 h, the One-Glo Luciferase Assay System (Promega) was used to quantify luciferase expression following manufacturer’s instructions.

### Cell growth and proliferation assays

A total of 5,000 cells were seeded into either 6-well or 12-well plates and imaged using an IncuCyte Zoom Imager (Sartorius) for live-cell imaging. Imaging was performed at the specified time points using the phase-contrast channel, as previously described^73^. Cell growth was assessed by measuring the percentage confluence.

### Molecular docking

A model for human USP47 catalytic domain (UniProt: Q96K76), which is representative of a covalently bound inhibitor conformational state, was generated using the structure of human USP7 in complex with the covalent inhibitor, FT827 (PDB: 5NGF; Turnbull *et al*., 2017), as the template in SwissModel^82^. Coordinates for **Compound 3** were generated using ChemDraw Prime 23.0.1.10 and molecular docking was performed using CovDock^83^ implemented in Maestro version 13.8.135 (Schrödinger Release 2024-3: Maestro, Schrödinger, LLC, New York, NY, 2024). The best docking pose for **Compound 3** corresponded to a docking score of -5.704.

### Antibodies

List of antibodies used for immunoblotting in this study:

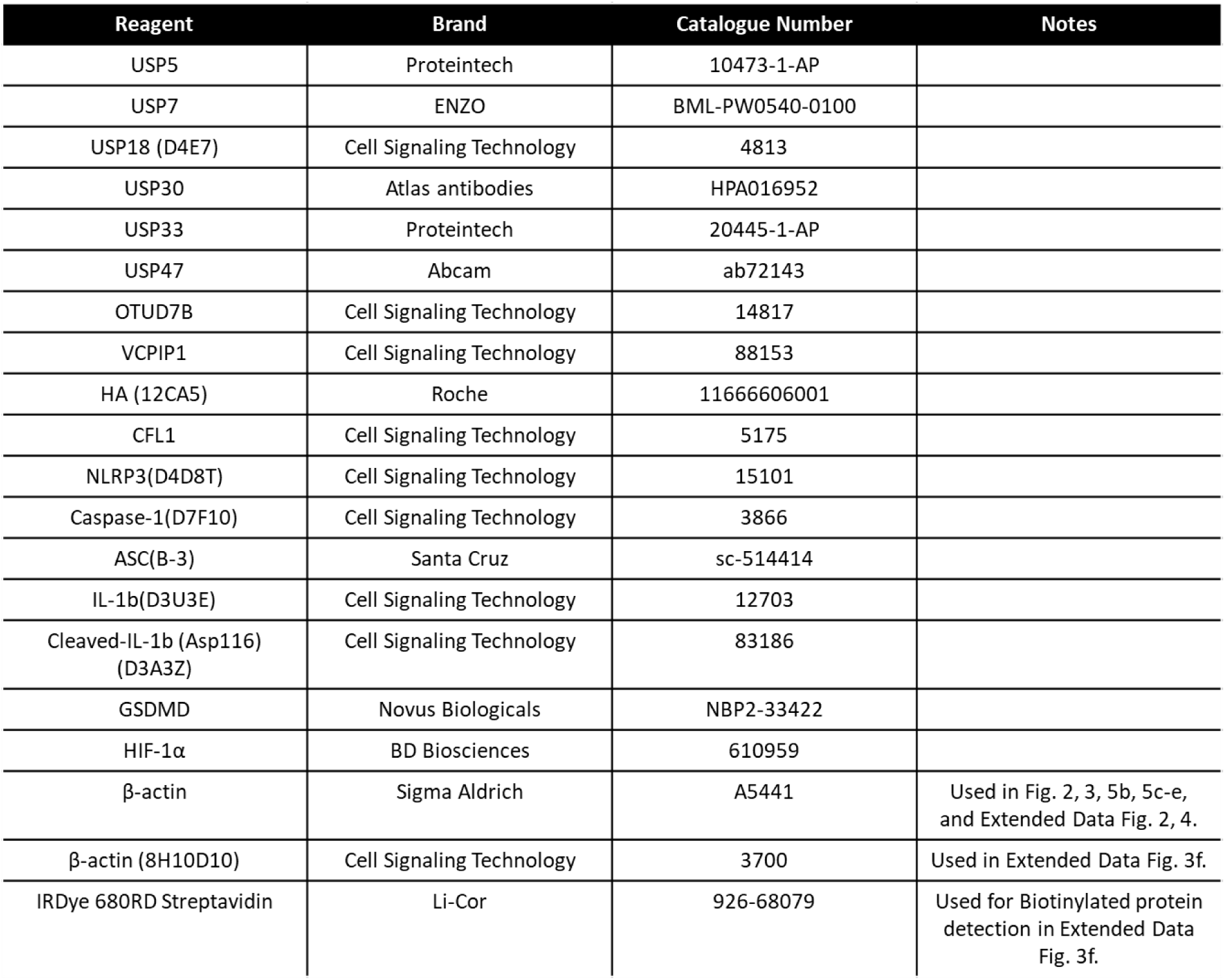

## Data availability

The mass spectrometry proteomics data have been deposited to the ProteomeXchange Consortium via the PRIDE partner repository with the dataset identifier PXD064209. The mass spectrometry HDX data have been deposited to the ProteomeXchange Consortium via the PRIDE partner repository with the dataset identifier PXD064060.

## Acknowledgements

The A.P.-F. and B.M.K. labs were supported by the Chinese Academy of Medical Sciences Innovation Fund for Medical Science, China (grant number: 2018-I2M-2-002) and by Pfizer Inc. The A.P.-F. lab was also supported by CRUK, Ono Pharma, and Boehringer Ingelheim. We would like to thank Prof. Jan Rehwinkel for kindly supplying the type I interferon reporter cells. H.B.L.J. was supported by a Bristol-Myers Squibb fellowship.

## Author contributions

H.B.L.J., S.D., L.B., and A.P.-F. designed research; H.B.L.J., S.D., S.S.H., P.A.C.W., C.N., Z.L., J.D., J.L., E.M., A.B., A.P., J.W.H., M.E.R., S.F., A.T., D.O.B., E.S., and A.P.T. performed research; P.R.E. contributed new reagents/analytic tools; I.V., E.W.T., D.O.B., G.M.W., B.M.K., C.J.S., L.B., A.P.-F. provided supervison; H.B.L.J., L.B., and A.P.-F. wrote the paper, and D.O.B., S.D., S.S.H., A.P.T., C.J.S., P.R.E., and B.M.K. provided feedback on the manuscript.

## Ethics declaration

Non applicable

## Corresponding authors

Hannah B.L. Jones (H.B.L.J.), Lennart Brewitz (L.B.), and Adán Pinto-Fernández (A.P.-F.)

## Extended Data

**Extended Data Fig. 1:**
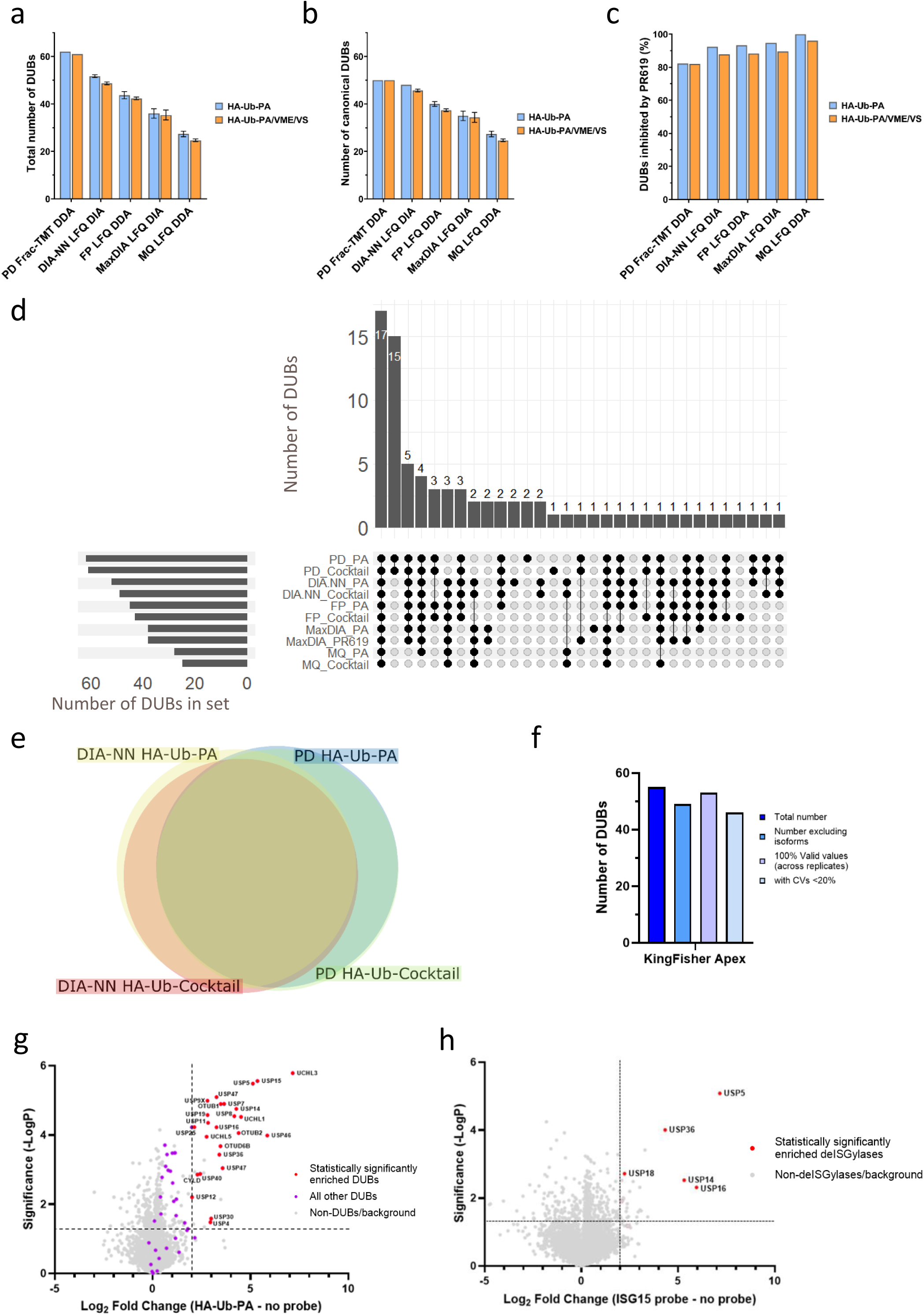
ABPP optimisation and flexibility. **a-b,** Comparison of the total number of DUBs, including isoforms (a) and canonical DUBs (b), identified from MCF-7 lysates by ABPP using either a propargylamine warhead (HA-Ub-PA) or a mixture of propargylamine, vinyl methyl ester (VME) and vinyl sulfone (VS) warheads, using ultra-sensitive ABPP workflows (error bars = SD). **c**, Percentage of DUBs inhibited >50% by PR619 from each ABPP workflow. **d**, Upset plot showing the commonality of DUBs identified from MCF-7 cell lysates by each high DUBome depth workflow including isoforms. **e**, Comparison of the identities of canonical DUBs identified from MCF-7 lysates using proteome discoverer (PD) from data-dependent acquisition (DDA) tandem mass tagged (TMT) labelled fractionated ABPP samples, or data-independent acquisition (DIA)-NN DIA non-fractionated label-free ABPP samples, using HA-Ub-PA or HA-Ub-PA/VME/VS (cocktail). Frac: high pH fractionated; LFQ: Label-free quantitation; FP: FragPipe; MQ: Maxquant. **f-g,** The number of DUBs identified from HAP1 lysates by ABPP in total (f), and in terms of statistical significance as illustrated on a volcano plot (g) using the ABPP-HT* high-throughput workflow with KingFisher Apex 96 well automated affinity purification. **h,** Enrichment of DUBs in HAP1 lysates by bio-ISG15-PA ABPP using the ABPP-HT* workflow with Kingfisher Apex 96 well automated affinity purification.

**Extended Data Fig. 2:**
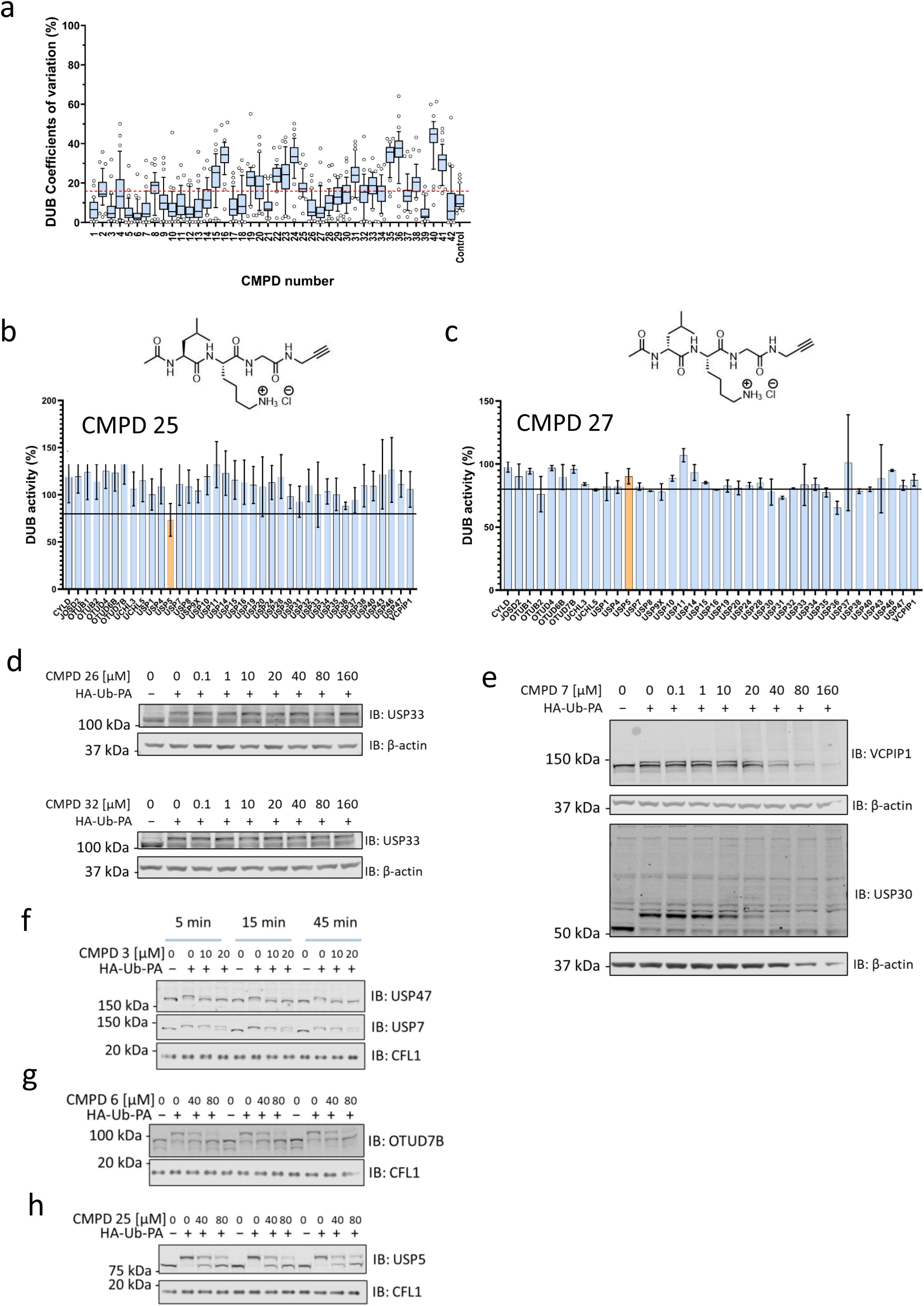
Hit Validation and Dose-response Analysis. **a**, Coefficient of variation (CV) for DUBs identified across the *ob*ABPP-HT* screen for each of the tested compounds. Box and whiskers are at 10-90 percentile. Red dashed line: average CV for DUBs across all conditions. **b-c**, HA-Ub-PA ABPP of **Compounds 25** and **27** at 25 µM in MCF-7 cell lysates (error bars = SD). **d**, Concentration-dependent binding of **Compounds 26** and **32** to USP33 in MCF-7 cell lysates. Data was obtained using with HA-Ub-PA as ABP. **e**, Concentration-dependent for VCPIP1 in MCF-7 lysates and USP30 in SH-SY5Y lysates by **Compound 7** with HA-Ub-PA labelling. The observation of high-molecular weight smearing and reduced intensity of DUBs suggests that **Compound 7** mediated DUB cross-linking. **f-h**, Time-dependent incubation of MCF-7 lysates at the indicated compound concentrations using HA-Ub-PA labelling.

**Extended Data Fig. 3:**
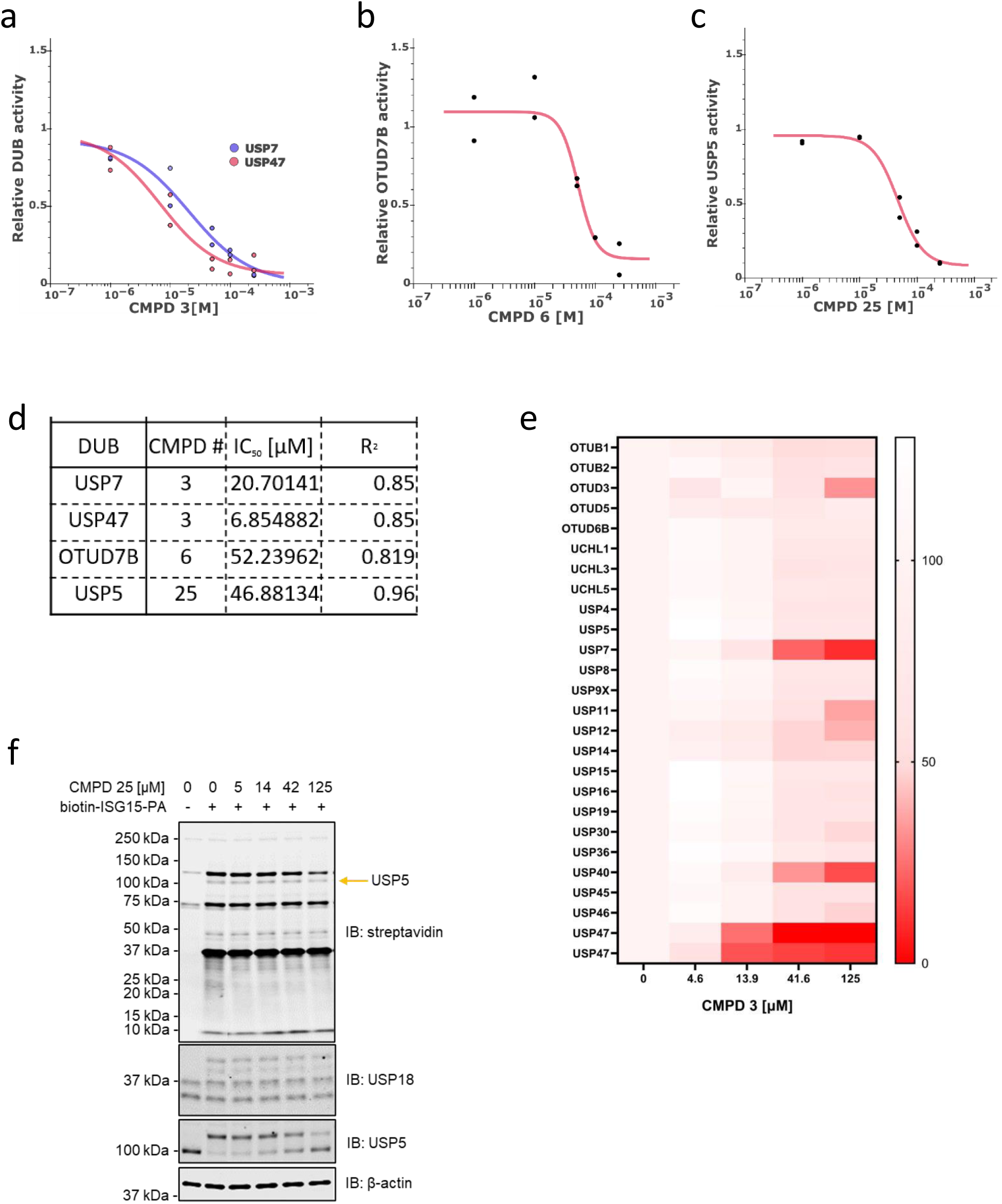
DUB Inhibitor Selectivity Assessment by *ob*ABPP-HT*. **a-c,** Concentration-dependent compound binding to DUBs identified using *ob*ABPP-HT*, with HA-Ub-PA. Data were analysed using curve curator^75^. **d,** IC50 values obtained using curve curator from the curves shown in a-c. **e,** Concentration-dependent binding of **Compound 3** to DUBs in HAP-1 lysates determined using high-throughput KingFisher Apex ABPP with HA-Ub-PA. **f, Compound 25** concentration dependence bio-ISG15-PA ABP with USP5 and USP18 in lysates from HAP1 cells treated with 1000 U/mL of human interferon α 2 (α2b) for 24 h.

**Extended Data Fig. 4:**
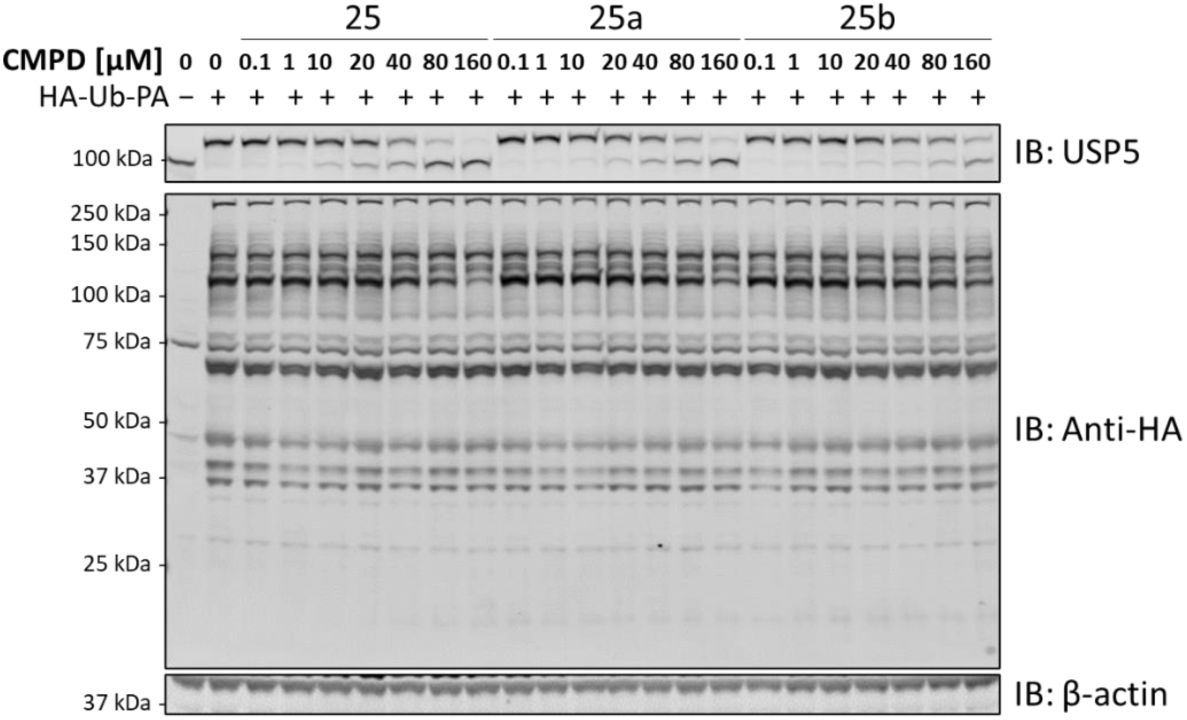
Selective inhibition of USP5 by compounds 25, 25a and 25b. HA immunoblot showing concentration-dependent selective binding of **Compounds 25**, **25a** and **25b** to USP5 in MCF-7 cell lysates using HA-Ub-PA ABP.

**Extended Data Fig. 5.**
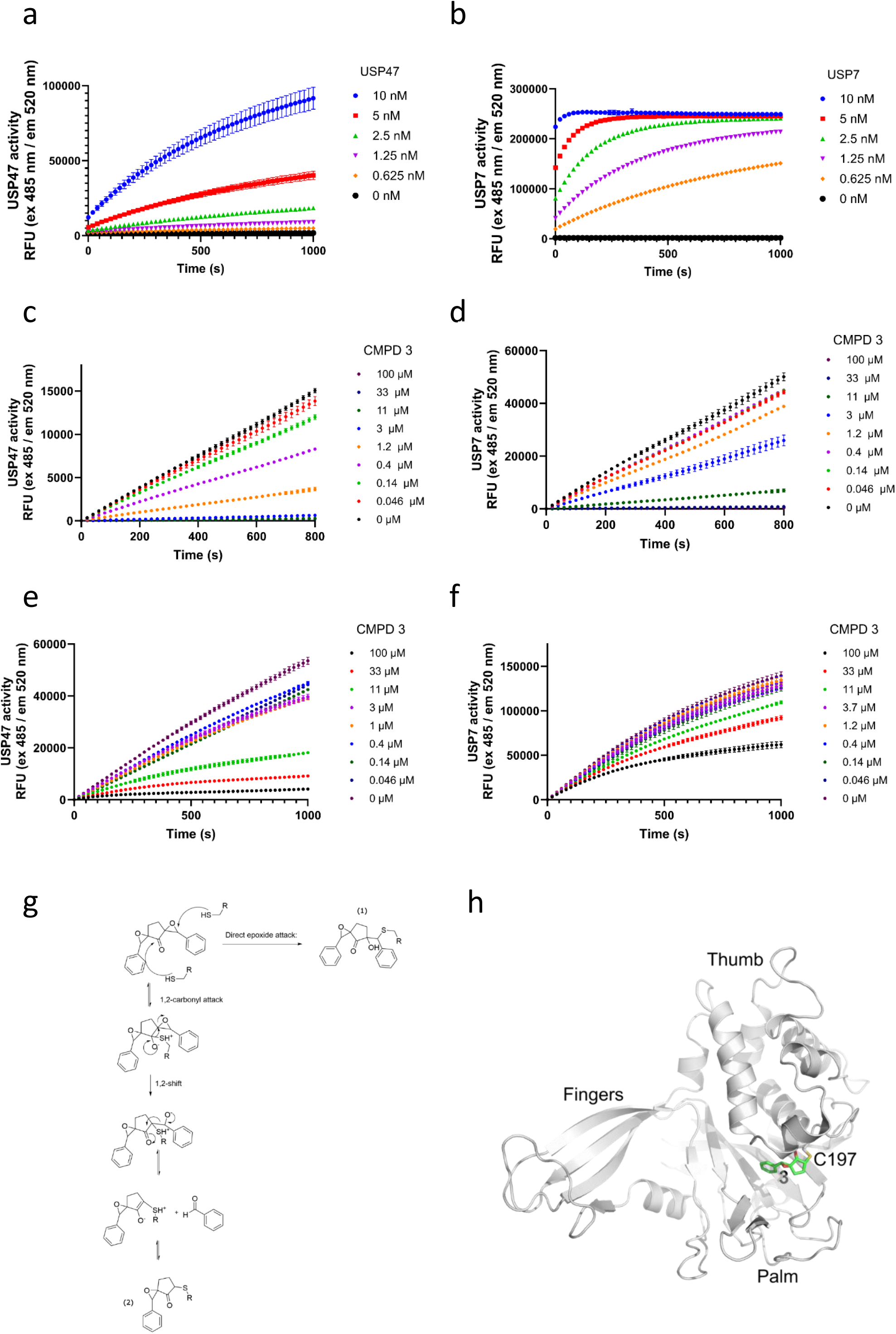
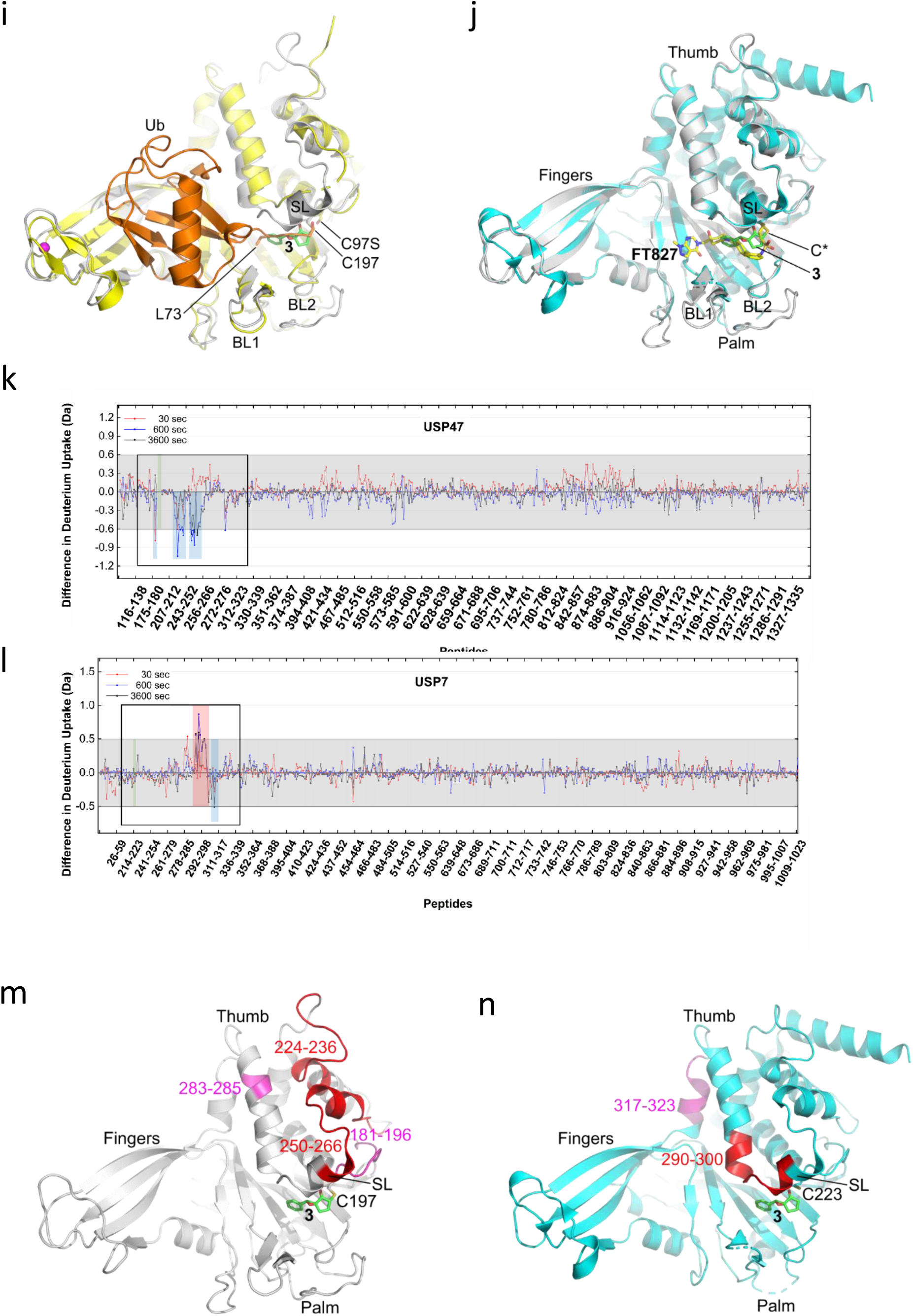
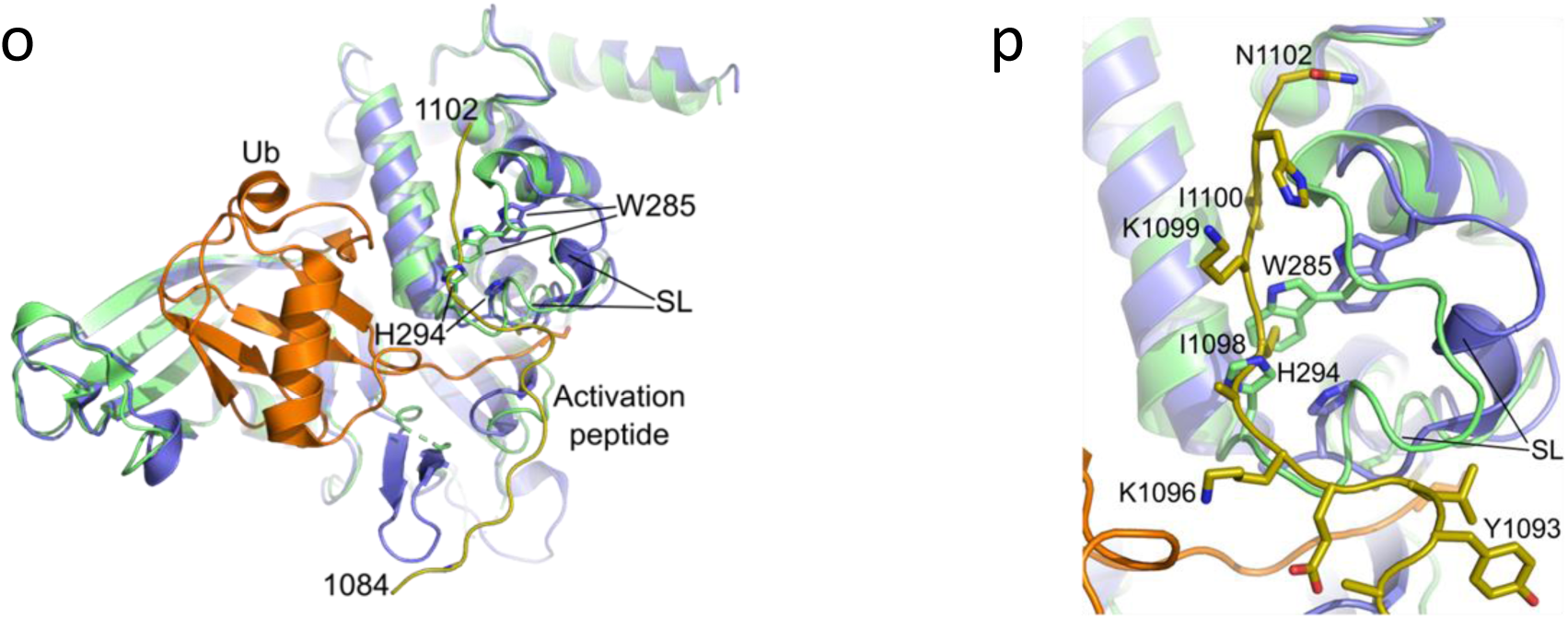
Molecular Basis for Selective Inhibition of USP47 by Compound 3. **a-b**, Progress curves showing (a) USP47- and (b) USP7-catalysed hydrolysis of Ub-Rho over time (error bars represent SD). **c-d**, Progress curves showing (c) USP47- and (d) USP7-catalysed hydrolysis of Ub-Rho in the presence of varying concentrations of **Compound 3**, following preincubation of USP47 or USP7 with **Compound 3** for 30 min (error bars: standard error of the mean (SEM)). **e-f**, Progress curves showing (e) USP47- and (f) USP7-catalysed hydrolysis of Ub-Rho in the presence of varying concentrations of **Compound 3**, without preincubation (error bars: SEM). **g,** Proposed mechanism of the covalent reaction of **Compound 3** with the catalytic cysteine C197 of human USP47. At least two alternative mechanisms for the reaction of **Compound 3** with the catalytic cysteine are plausible, with the reaction proceeding via 1,2-carbonyl attack and retro-aldol fragmentation to form (2) being the most probable based on previous observations using a related diepoxide^26^. **h,** Modelled structure of human USP47 in complex with **Compound 3** shown as a stick representation with carbon atoms colored green. The thumb, palm and fingers subdomains of the catalytic domain and the catalytic cysteine, Cys197, are highlighted. Figure prepared using PyMOL (The PyMOL Molecular Graphics System, version 2.5.8; Schrödinger, LLC). **i,** Superposition of the modelled structure of human USP47 in complex with **Compound 3** on the X-ray structure of *C.elegans* USP47 in complex with ubiquitin (PDB code: 8ITP^33^). The thumb, palm and fingers subdomains, the *C.elegans* USP47 catalytic cysteine point mutation (C97S), and the catalytic cysteine in the human USP47 model (Cys197) are highlighted. The catalytic domain of the modelled structure of human USP47 in complex with **Compound 3** is colored grey, and **Compound 3** is shown with carbon atoms colored green. The catalytic domain of *C.elegans* USP47 in complex with Ub is colored yellow with Ub shown in orange. **Compound 3** is predicted to bind in the thumb-palm cleft that is reported to guide the ubiquitin C-terminus into the active site. The positions of BL1, (BL2), and SL are highlighted. **j,** Superposition of the modelled structure of human USP47 (grey) in complex with **Compound 3** (green carbon atoms) on the X-ray structure of human USP7 (cyan) in complex with the covalent inhibitor **FT827** (yellow carbon atoms; PDB code: 5NGF^28^). The thumb, palm and fingers subdomains, and positions of blocking loops 1 and 2 (BL1, BL2) and switching loop (SL) are highlighted. C* denotes the catalytic cysteine (corresponding to C197 in human USP47 and C223 in human USP7). **Compound 3** is predicted to bind within the thumb-palm cleft in a similar manner as **FT827** with **FT827** extending out further towards the fingers subdomain. **k-l,** Differential HDX-MS analysis of USP47 and USP7 in apo- and holo-states. A total of 491 peptides covering 88.2% of the full-length USP47 protein sequence were identified, with an average residue redundancy of 4.2. Similarly, a total of 536 peptides covering 84.2% of the full-length USP7 sequence were identified, with an average residue redundancy of 5.9. Residual ΔHDX plots showing differences in deuterium uptake between holo- and apo-states for USP47 (k) and USP7 (l), across all mapped peptide fragments and time points (30 s to 1 h). Positive ΔHDX values indicate increased solvent exposure or flexibility in the presence of **Compound 3**, whilst negative values suggest protection or structural stabilisation. A grey shaded area marks the significance threshold for ΔHDX applied in each study. **m,** Modelled structure of human USP47 in complex with **Compound 3** (carbon atoms in green) highlighting regions identified in the HDX-MS analysis colored red (mean perturbation over 60 min of 5– 20%) and magenta (mean perturbation over 60 min of <60%). The thumb, palm and fingers subdomains, catalytic cysteine (C197), and SL are labelled **n,** Human USP7 catalytic domain (from PDB code: 5NGF^28^) colored cyan with the superimposed modelled position of **Compound 3** in human USP47 (green carbon atoms) highlighting regions identified in the HDX-MS analysis colored red (mean perturbation over 60 min of 5–20%) and magenta (mean perturbation over 60 min of <60%). The thumb, palm and fingers subdomains, catalytic cysteine (C223), and SL are labelled. **o,** Structure of apoform human USP7 catalytic domain (PDB code: 1NB8^87^; lime) superimposed on the structure of the human USP7 catalytic domain from the USP7-catalytic domain-UBL45-ubiquitin complex (PDB code: 5JTV^34^; catalytic domain in violet, the C-terminal activation peptide in gold and ubiquitin in orange). The 19-amino acid activation peptide at the C-terminus of full length human USP7 stabilizes the SL (especially H294 and W285) in an active “out” conformation. **p,** Close-up view of (o) highlighting the conformational shift in SL between the inactive “in” conformation in apoform USP7 and active “out” conformation in its complex with ubiquitin.

**Extended Data Fig. 6:**
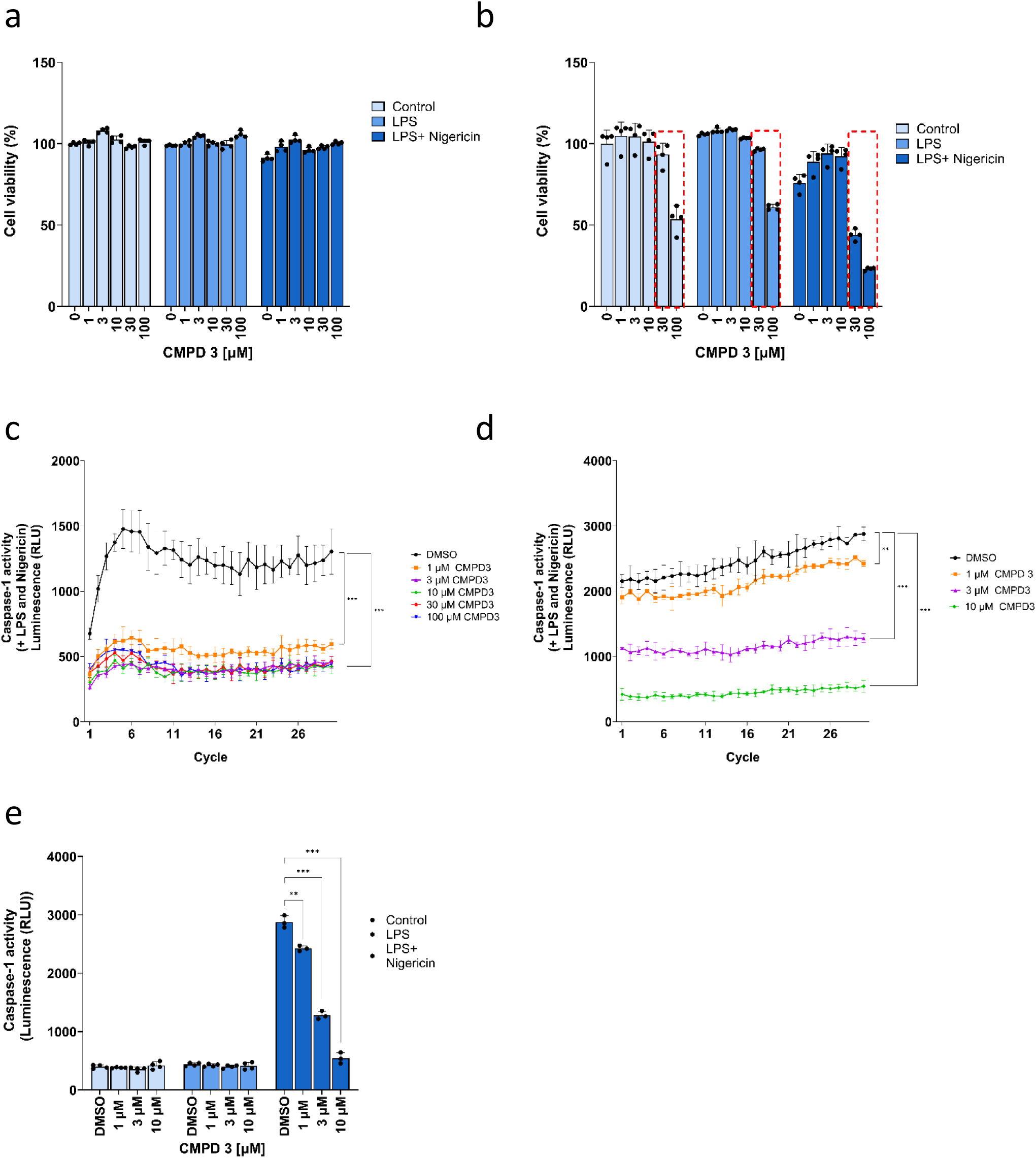
Compound 3 inflammasome activation and toxicity. **a-b**, MTS cell viability of THP-1 cells treated with indicated concentrations of **Compound 3** ± LPS or LPS/Nigericin. Compound treatment for 4 h, 1 µg/mL LPS treatment for 4 h, 10 µM Nigericin treatment for 45 min (a) and 2 h (b) (error bars = SD). **c-d**, Caspase-1 activity over time in the media (supernatant) of PMA-differentiated THP-1 cells treated with indicated concentrations of **Compound 3**, determined using Caspase-Glo 1 luminescence assay. **Compound 3** incubation for 4 h, 1 µg/mL LPS incubation for 4 h and 10 µM Nigericin incubation for 45 min (c) and 2 h (d) (error bars = SD). Two-way ANOVA with Dunn’s multiple comparison test was used. ***P < 0.001 **e**, Caspase-1 activity in the media (supernatant) of PMA-differentiated THP-1 cells treated with indicated concentrations of **Compound 3**, determined using Caspase-Glo 1 luminescence assay. **Compound 3** incubation for 4 h, ± 1 µg/mL LPS incubation for 4 h and 10 µM Nigericin incubation for 2 h (error bars = SD).

**Extended Data Fig. 7:**
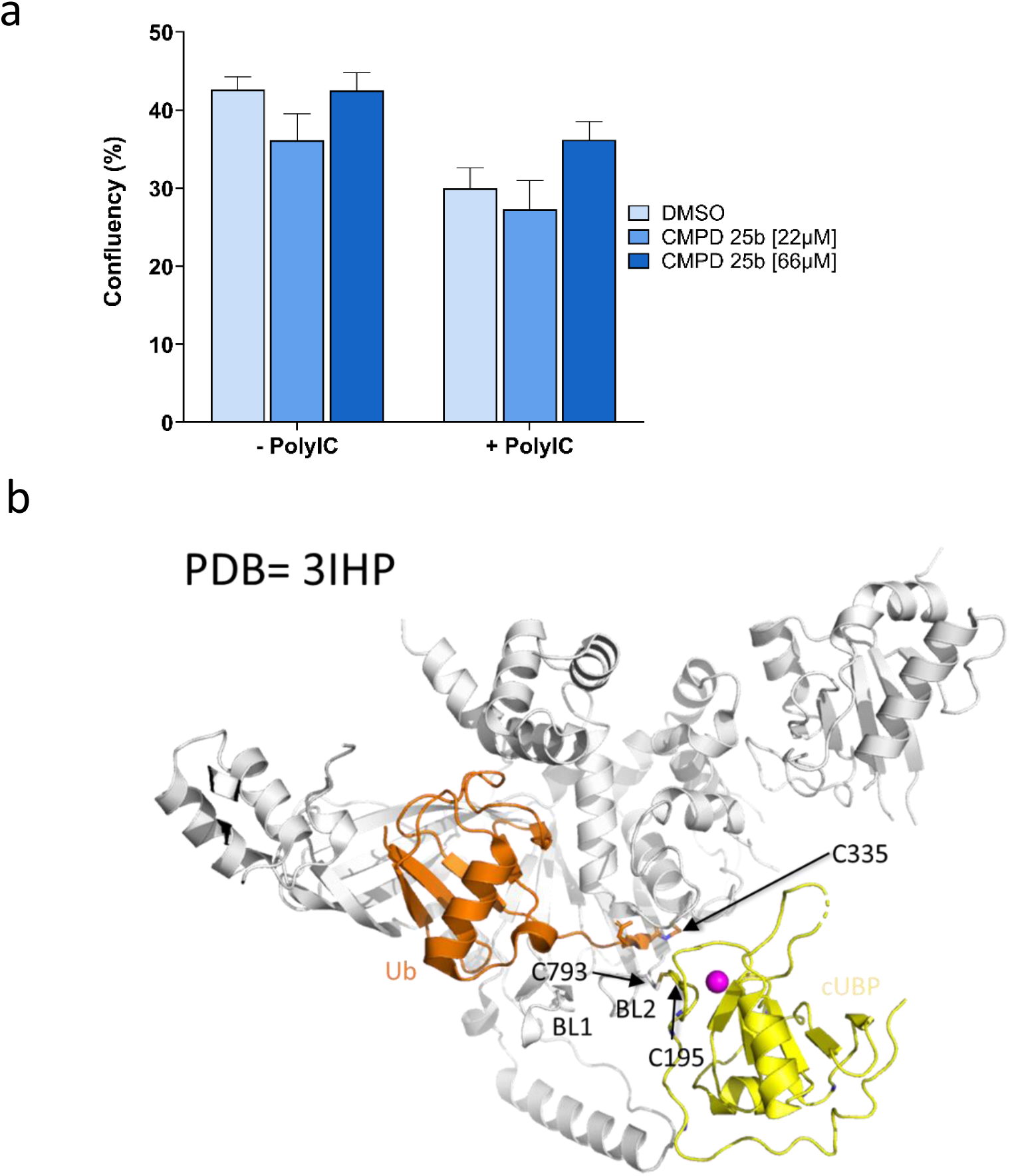
Studies on cellular toxicity and molecular basis for selectivity of Compound 25b a, Compound 25b cellular toxicity. **a,** HAP1 cells were seeded in triplicate at 5,000 cells per well in a 96-well plate, and treated with **Compound 25b** ± Poly(I:C) 0.5 µg/mL for 48 h. Confluence at 48 h was determined by live-cell imaging using an IncuCyte SX5 machine (Sartorius) (error bars = SEM). **b**, Structure of the reported covalent USP5–Ub complex (PDB ID: 3IHP^63^) highlighting a disulphide bridge between C793 of the BL2 loop and C195 of the cUBP domain, observed in chains A and B. This tethering may restrict BL2 flexibility and position cUBP near the putative **Compound 25** binding site, potentially enabling additional interactions. As C195 is also implicated in USP5 dimerisation, the functional relevance of this disulphide bond warrants further investigation.

**Extended Data Fig. 8:**
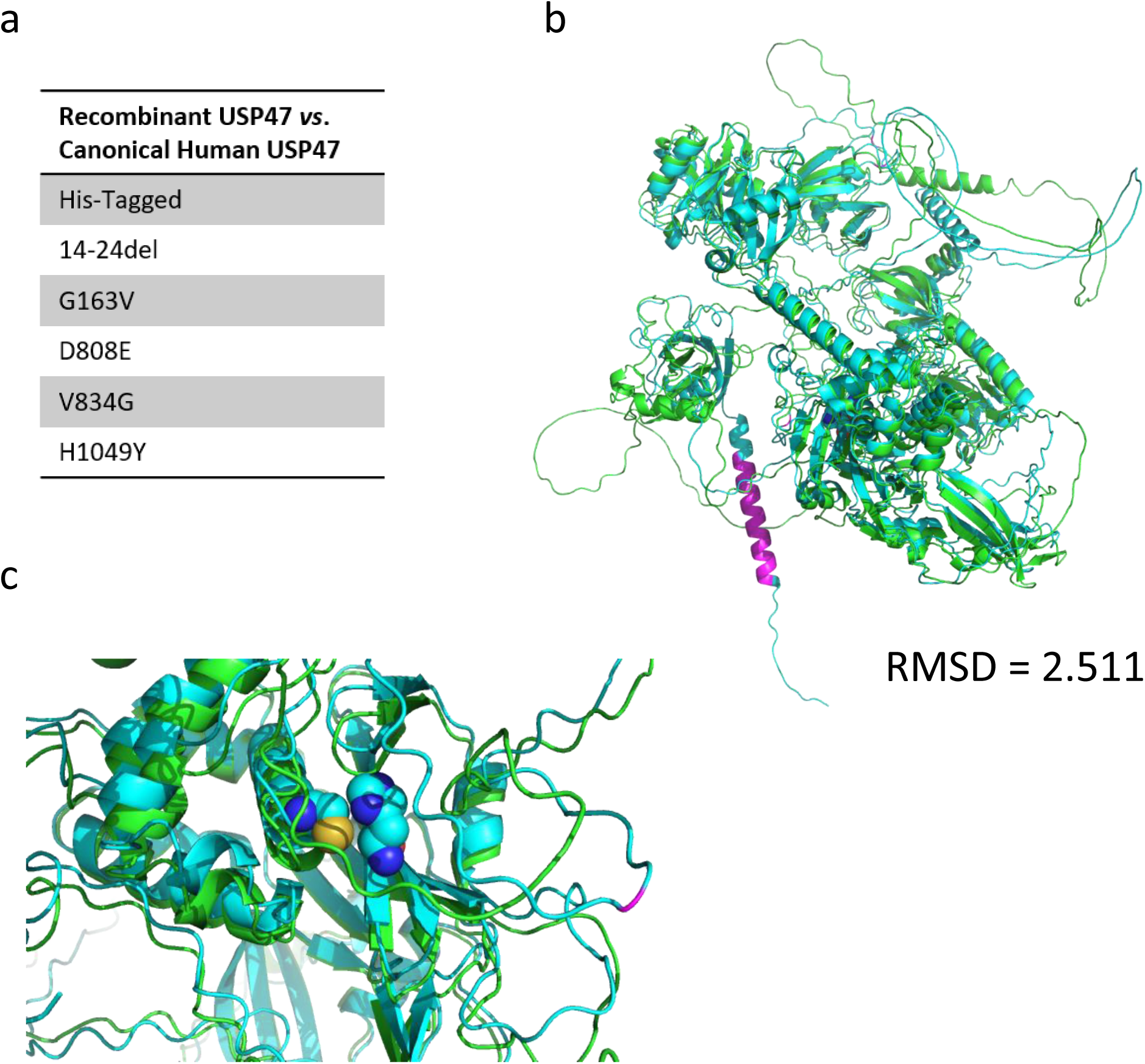
USP47 protein *Pan troglodytes* isoform 2 vs *Homo sapiens* isoform 1. **a**, Sequence differences between USP47 of *Pan troglodytes* isoform 2 and canonical human USP47. **b**, Alignment of AlphaFold structures of USP47 based on *Pan troglodytes* isoform 2 used in kinetic and HDX-MS studies and canonical human USP47. Sequence differences are in pink, the catalytic cysteine 197 and histidine 503 residues are as spheres. **c**, Zoom in of alignment showing the catalytic cysteine 197 and histidine 503 residues as spheres and the nearest sequence difference in pink.

**Extended Data Table 1:**
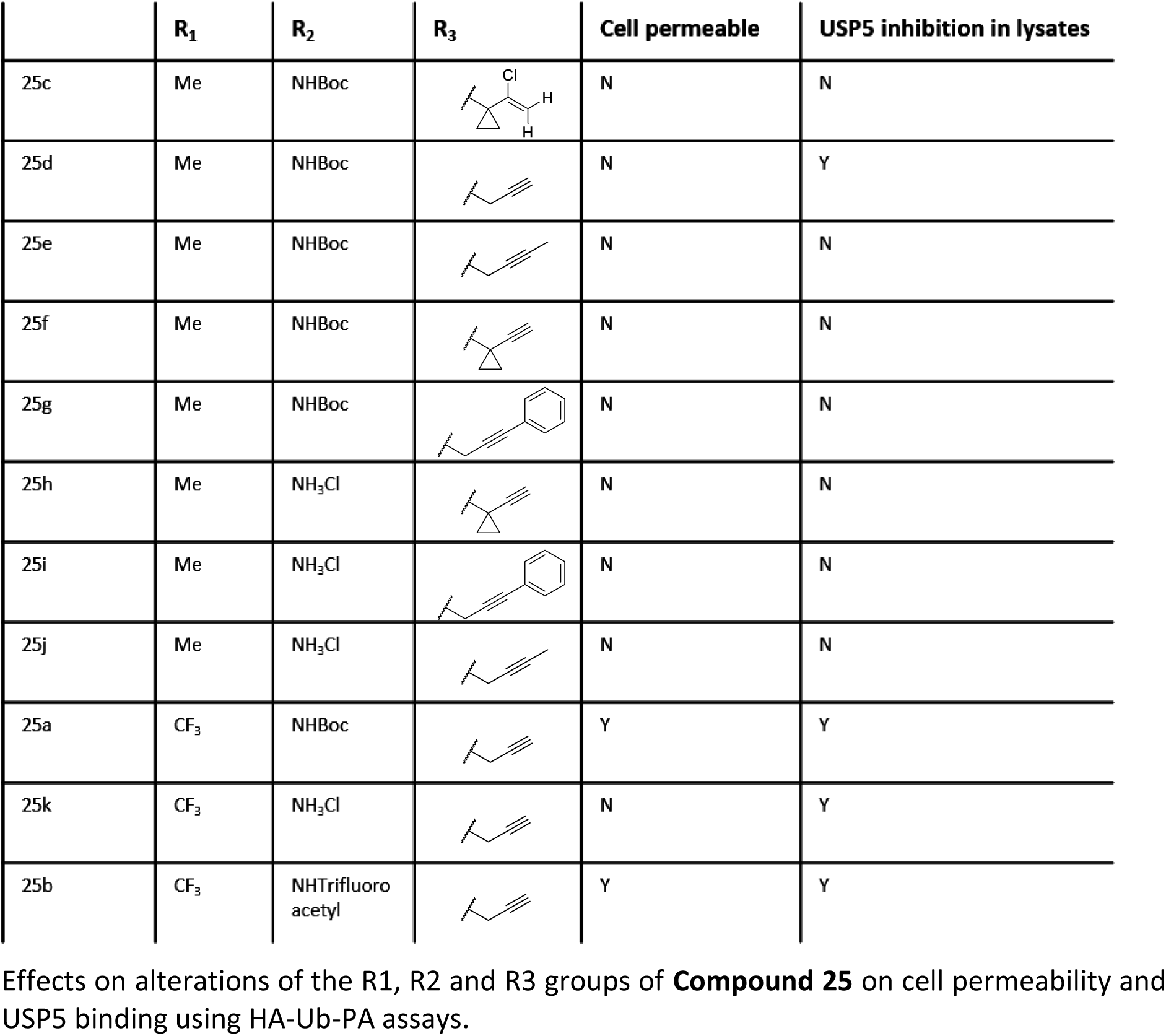
Structure-activity relationship (SAR) of Compound 25 to increase cellular permeability. Effects on alterations of the R1, R2 and R3 groups of Compound 25 on cell permeability and USP5 binding using HA-Ub-PA assays.

|  | R<sub>1</sub> | R<sub>2</sub> | R<sub>3</sub> | Cell permeable | USP5 inhibition in lysates |
| --- | --- | --- | --- | --- | --- |
| 25c | Me | NHBoc |  | N | N |
| 25d | Me | NHBoc |  | N | Y |
| 25e | Me | NHBoc |  | N | N |
| 25f | Me | NHBoc |  | N | N |
| 25g | Me | NHBoc |  | N | N |
| 25h | Me | NH<sub>3</sub>Cl |  | N | N |
| 25i | Me | NH<sub>3</sub>Cl |  | N | N |
| 25j | Me | NH<sub>3</sub>Cl |  | N | N |
| 25a | CF<sub>3</sub> | NHBoc |  | Y | Y |
| 25k | CF<sub>3</sub> | NH<sub>3</sub>Cl |  | N | Y |
| 25b | CF<sub>3</sub> | NHTrifluoroacetyl |  | Y | Y |

