## Supplementary Information for "*ob*ABPP-HT*: A Precision-Engineered Activity Proteomics Pipeline for the Streamlined Discovery of Deubiquitinase Inhibitors"

**Supplementary Information Data Table 1.**

| | |
|---|---|
| 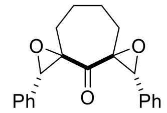 **1** | 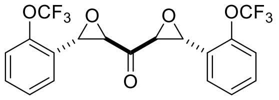 **5** |
| 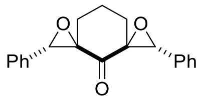 **2** | 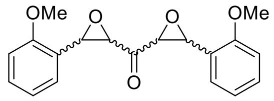 **6** (2:1 *trans,trans*:*cis,cis*) |
| 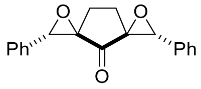 **3** | 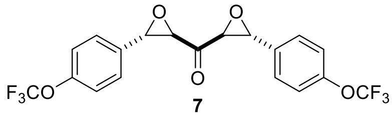 **7** |
| 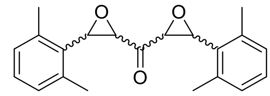 **4** (1.2:1 *trans,trans*:*cis,cis*) | 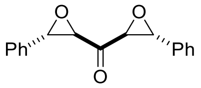 **8** |

Structures of reported αβ,α'β'-diepoxyketones (DEKs) tested using *ob*ABPP-HT*[1].

**Supplementary Information Data Table 2.**

| | | | |
|---|---|---|---|
| 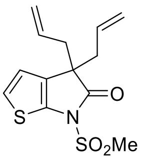 **9** | 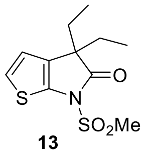 **13** | 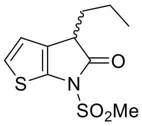 **17** | 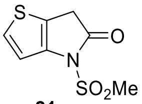 **21** |
| 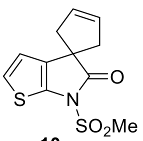 **10** | 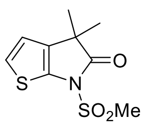 **14** | 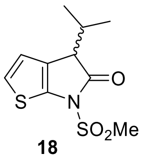 **18** | 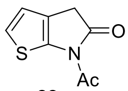 **22** |
| 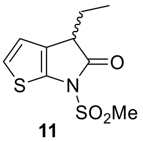 **11** | 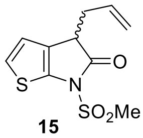 **15** | 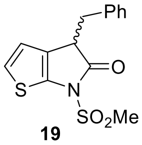 **19** | 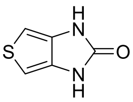 **23** |
| 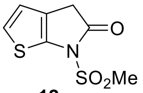 **12** | 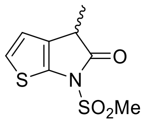 **16** | 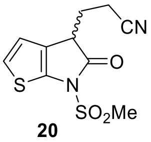 **20** | |

Structures of reported γ-lactams tested using *ob*ABPP-HT*[2].

**Supplementary Information Data Table 3.**

| | | |
|---|---|---|
| 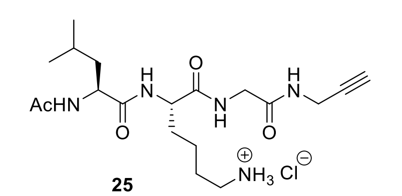 **25** | 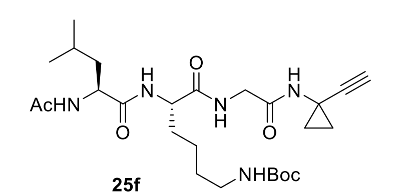 **25f** | 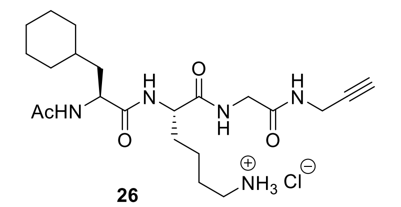 **26** |
| 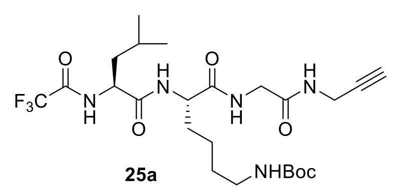 **25a** | 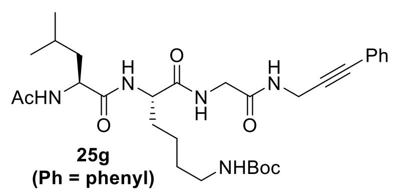 **25g (Ph = phenyl)** | 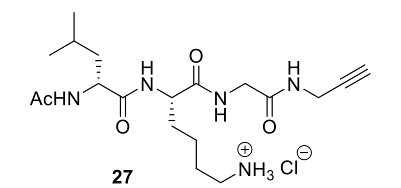 **27** |
| 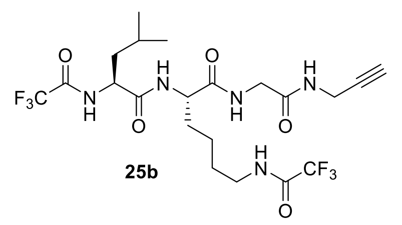 **25b** | 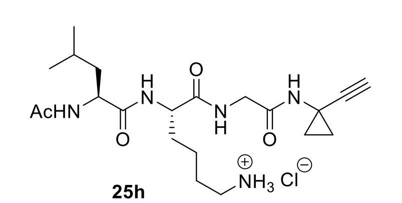 **25h** | 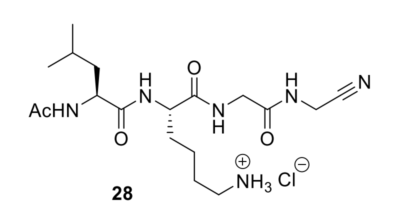 **28** |
| 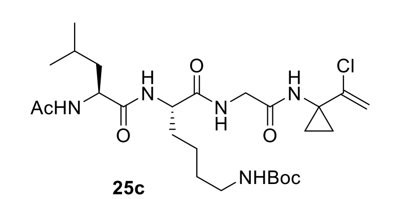 **25c** | 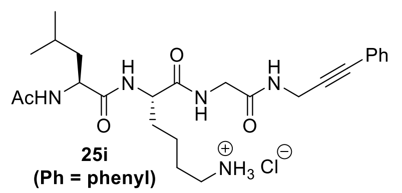 **25i (Ph = phenyl)** | 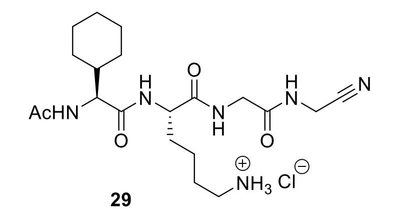 **29** |
| 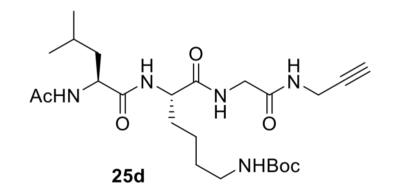 **25d** | 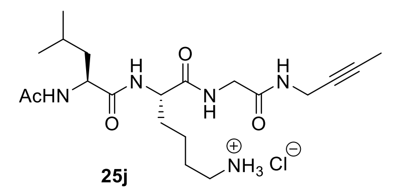 **25j** | 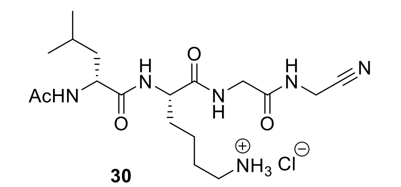 **30** |
| 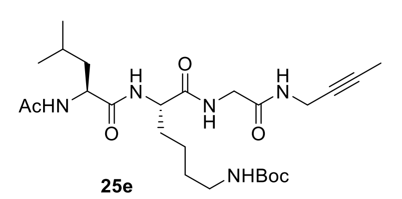 **25e** | 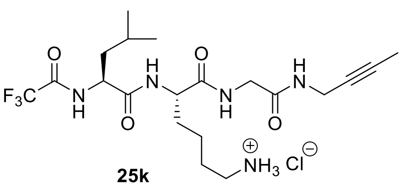 **25k** | 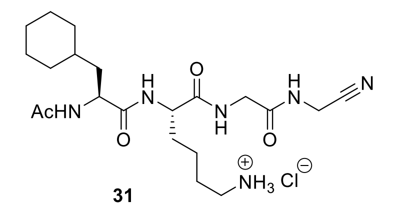 **31** |

Structures of Ub-derived peptides designed for this work that were tested using *ob*ABPP-HT*.

**Supplementary Information Figure 1: Reported small-molecule USP5 inhibitors.**

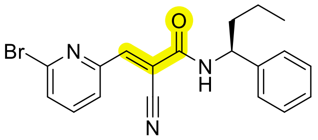

**Degrasyn (WP1130)**

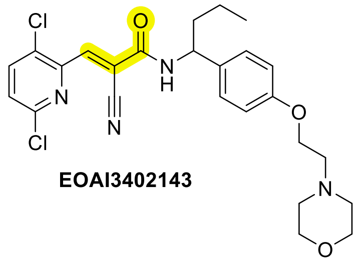

**EOAI3402143**

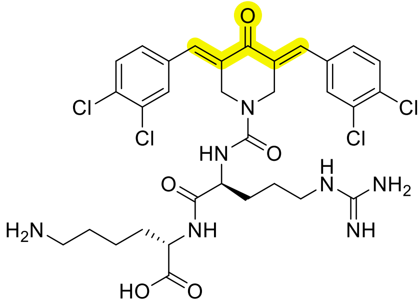

**RA-9**

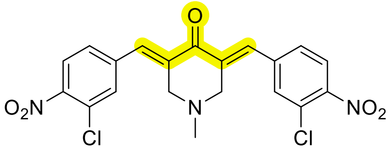

**RA-14**

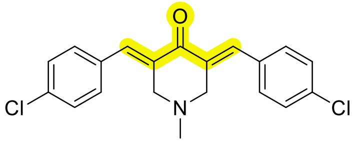

**AM146**

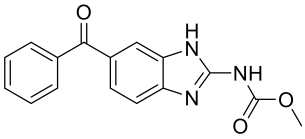

**Mebendazole**

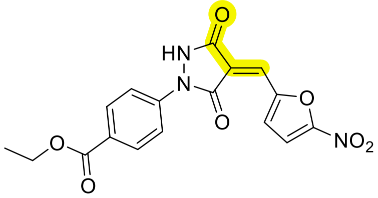

**PYR-41**

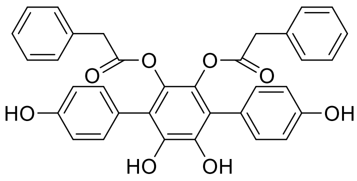

**Vialinin A**

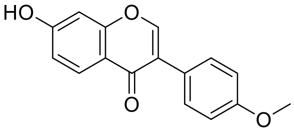

**Formononetin**

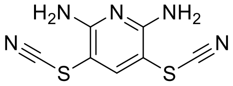

**PR 619**

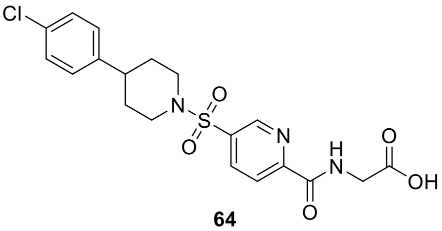

**64**

Several small-molecule USP5 inhibitors are reported, which, however, to our knowledge, have not yet advanced into clinical testing[3]. Some of the reported small-molecule USP5 inhibitors likely work via covalently reacting with USP5. **Degrasyn** (WP1130): Degrasyn is a pan-DUB inhibitor, which in addition to USP5, is also reported to inhibit USP9x, USP14, USP37, and UCHL1, as well as other nucleophilic cysteine enzymes[4]. It likely inhibits USP5 via covalent reaction. **EOAI3402143**: Like the structurally related degrasyn, EOAI3402143 is a pan-DUB inhibitor, which in addition to USP5 is reported to also inhibit USP9x and USP24[5,6]. **PR619**: Like degrasyn and EOAI3402143, PR619 is a pan-DUB inhibitor, which in addition to USP5 is reported to also inhibit e.g., USP2, USP4, USP7, USP8, USP15, USP20, USP28.[7] It likely inhibits via cyanide transfer[8]. **Dibenzylideneacetones RA-9, RA-14, AM146**: These dibenzylideneacetone derivatives are cell-permeable USP5 inhibitors, which likely inhibit via covalent reaction. However, they display imperfect selectivity and efficiently inhibit UCHL1, UCHL3, USP2, and USP8, in addition to USP5[9]. **PYR-41**: Inhibits various DUBs including USP5 (USP9x, USP14, UCH37, UCHL3)[10]. **Vialinin A**: Inhibits USP5, USP4 and UCHL1, not much else has been tested so far[11]. **Mebendazole**: Mebendazole is clinically used to treat parasite infections[12]. Notably, it is reported to also efficiently inhibit USP5 and USP7 suppressing c-Maf transcription. Note, however, that its effects on other DUBs have, to our knowledge, not yet been described. **Formononetin** inhibits USP5 and structurally most similar USP13, however no mechanistic studies have been conducted[13]. **Cmpd 64** Potentially selective, tested HDAC, USP3, USP13, USP16, USP33, USP39, USP49, USP51, BRAP, and was selective for USP5[8,14].

**Supplementary Information Data Scheme 1: Synthesis of substrate-based small-molecules tested for DUB inhibition (25-31).**

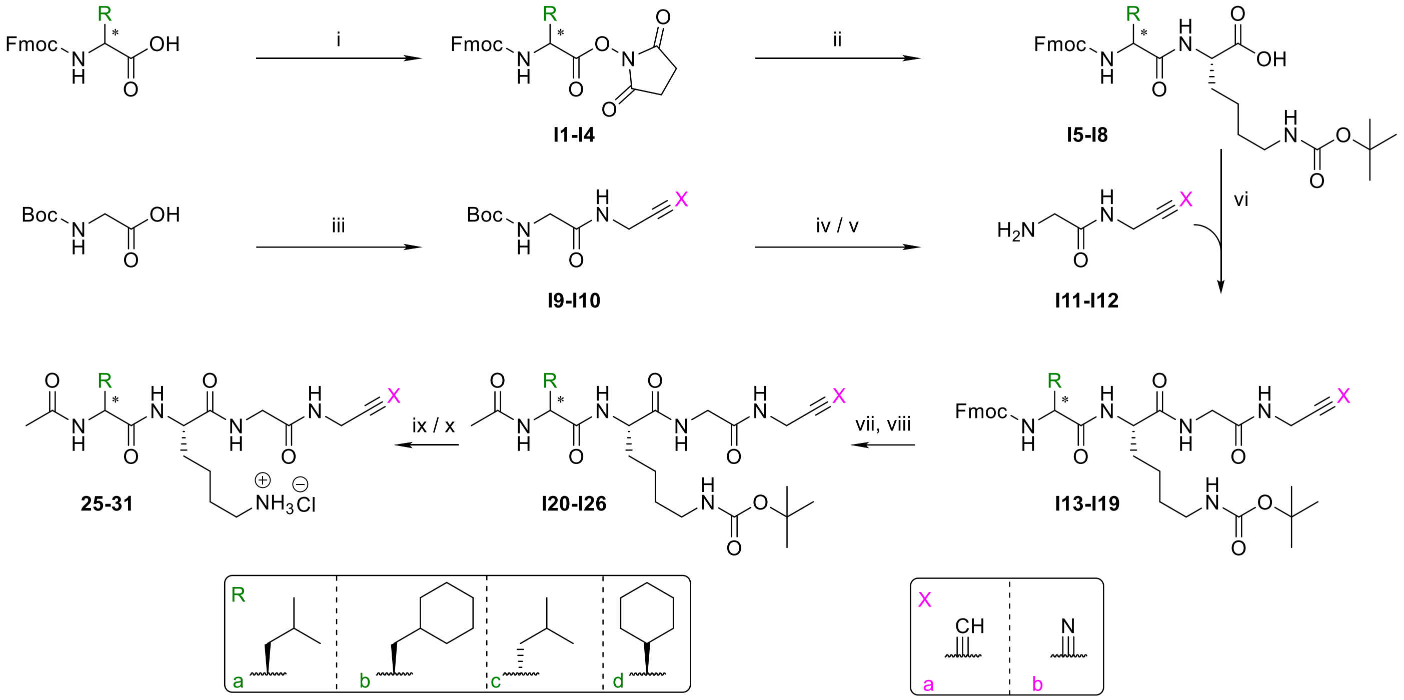


Reagents and Conditions: i) *N*-hydroxysuccinimide (NHS), *N*,*N*'-dicyclohexylcarbodiimide (DCC), tetrahydrofuran, 0 °C to rt, 89%-app. quant.; ii) Lys(Boc)-OH, $Na_2CO_3$, tetrahydrofuran:water ($2{:}1_{v/v}$), 0 °C, 87-98%; iii) amine salt, *N*-(3-dimethylaminopropyl)-*N*'-ethylcarbodiimide (EDC), 1-hydroxybenzotriazole hydrate (HOBt), *N*-methylmorpholine (NMM), DCM, rt, 50-60%; iv) HCl in dioxane (4M), DCM, rt, app. quant.; v) formic acid, rt, app. quant.; vi) NMM, isobutyl chloroformate (IBCF), tetrahydrofuran:water ($3{:}1_{v/v}$), –15 °C to rt, 33-97%; vii) *N*,*N*-diethylamine, MeCN, rt, app. quant.; viii) *N,N*-dimethylpyridin-4-amine (DMAP), $Ac_2O$, tetrahydrofuran, reflux, 23-61%; ix) formic acid, rt; then: HCl in dioxane (4M), 66%-app. quant.; x) HCl in dioxane (4M), 2-propanol, rt, 78%-app. quant.

**Supplementary Information Data Scheme 2: Synthesis of substrate-based small-molecules tested for DUB inhibition (25a-25k).**

| Compound | R | X | Conditions |
|---|---|---|---|
| **25** | a | a | iv, x |
| **26** | b | a | iv, x |
| **27** | c | a | iv, x |
| **28** | a | b | v, ix |
| **29** | d | b | v, ix |
| **30** | c | b | v, ix |
| **31** | b | b | v, ix |

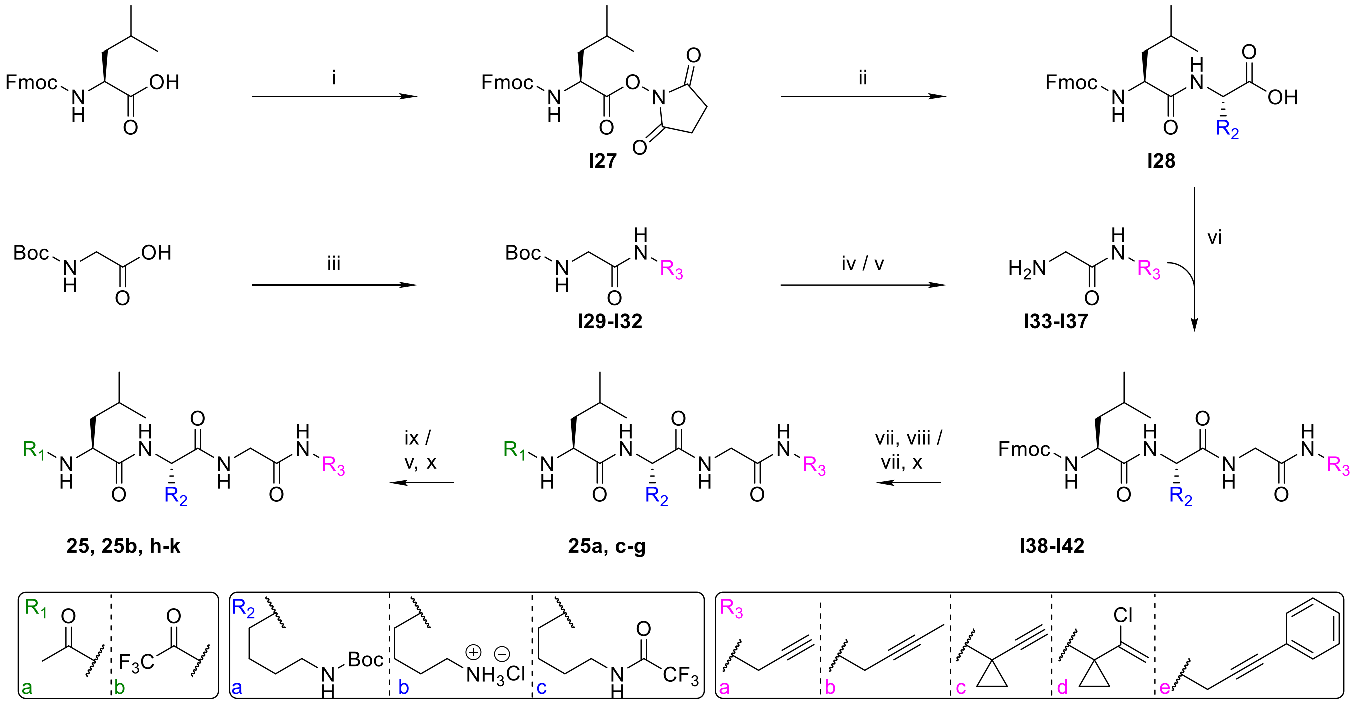


Reagents and Conditions: i) *N*-hydroxysuccinimide (NHS), *N,N'*-dicyclohexylcarbodiimide (DCC), tetrahydrofuran, 0 °C to rt, 66%; ii) Lys(Boc)-OH, $Na_2CO_3$, tetrahydrofuran:water ($2:1_{v/v}$), 0 °C, app. quant.; iii) amine salt, *N*-(3-dimethylaminopropyl)-*N'*-ethylcarbodiimide (EDC), 1-hydroxybenzotriazole hydrate (HOBt), *N*-methylmorpholine (NMM), DCM, rt, 54-85%; iv) HCl in dioxane (4M), DCM, rt, app. quant.; v) formic acid, rt, app. quant.; vi) NMM, isobutyl chloroformate (IBCF), tetrahydrofuran:water ($3:1_{v/v}$), –15 °C to rt, 91-100%; vii) *N,N*-diethylamine, MeCN, rt, app. quant.; viii) *N,N*-dimethylpyridin-4-amine (DMAP), $Ac_2O$, tetrahydrofuran, reflux, 1-21%; ix) formic acid, rt; then: HCl in dioxane (4M), 58-87%; x) DMAP, trifluoroacetic anhydride (TFAA), tetrahydrofuran, reflux, 1-11%.

| Compound | $R_1$ | $R_2$ | $R_3$ | Conditions | Compound | $R_1$ | $R_2$ | $R_3$ | Conditions |
|---|---|---|---|---|---|---|---|---|---|
| **I28** | - | a | - | | **I42** | - | a | e | |
| **I29** | - | - | a | | **25d** | a | a | a | vii, viii |
| **I30** | - | - | b | | **25e** | a | a | b | vii, viii |
| **I31** | - | - | c | | **25f** | a | a | c | vii, viii |
| **I32** | - | - | e | | **25c** | a | a | d | vii, viii |
| **I33** | - | - | a | iv | **25g** | a | a | e | vii, viii |
| **I34** | - | - | b | iv | **25a** | b | a | a | vii, x |
| **I35** | - | - | c | v | **25** | a | b | a | ix |
| **I36** | - | - | d | iv | **25j** | a | b | b | ix |
| **I37** | - | - | e | iv | **25h** | a | b | c | ix |
| **I38** | - | a | a | | **25i** | a | b | e | ix |
| **I39** | - | a | b | | **25k** | b | b | a | ix |
| **I40** | - | a | c | | **25b** | b | c | a | v, x |
| **I41** | - | a | d | | | | | | |

### Synthesis

#### General information

All reagents were purchased from commercial suppliers (Merck, Thermo Fisher Scientific, Enamine). Anhydrous solvents were under an atmosphere of $N_2$ or Ar gas. All solvents, liquids and solutions were transferred using stainless steel needles and syringes or gas-tight micro-syringes. All solvents used in extractions, workup and purifications were of HPLC grade.

Column chromatography was performed using an automated Biotage® Selekt purification machine using Biotage® Sfär Silica D Duo chromatography cartridges. Cartridge size, solvent gradients, and column volumes (CV) are specified in individual experimental procedures.

High-performance liquid chromatography (HPLC) purification was performed using a Shimadzu Nexera semi-preparative machine, with an Avantor® ACE® 10 AQ preparative column (250 × 21.2 mm) at flow rate of 20 mL/min, using a linear gradient of 2% (v/v) to 98% (v/v) acetonitrile in water (each containing 0.1% formic acid) over 45 or 30 min.

Thin layer chromatography (TLC) analysis was performed on Merck silica gel 60 $F_{254}$ TLC plates; compounds were visualized under UV light followed by ninhydrin stain.

High resolution mass spectrometry (HRMS) data was recorded in the positive ionization mode via flow injection analysis on an ACQUITY I-Class PLUS UPLC System (Waters, Milford, MA, USA) coupled to an ACQUITY RDa mass spectrometer (Waters, Milford, MA, USA) equipped with an electrospray ionization (ESI) probe.

NMR spectra were recorded using a Bruker AVIII HD 400, AVIII HD 600 or AVIII 700 instrument. For $^1$H NMR, chemical shifts are reported in ppm downfield from tetramethylsilane, referenced to residual protium in the NMR solvent (DMSO-$d_6$: δ = 2.50 ppm). For $^{13}$C NMR, chemical shifts are reported in the scale relative to the NMR solvent (DMSO-$d_6$: δ = 39.52 ppm). Coupling constants are accurate to 0.1 Hz. $^{19}$F NMR chemical shifts are reported in ppm in absolute reference to their respective $^1$H NMR spectra taken at the same probe under identical conditions.

IR spectroscopy was performed using a Bruker Tensor-27 Fourier transform infrared (FT-IR) spectrometer. Only purified samples were analyzed.

Optical Rotation (α) was determined using a Unipol (Schmidt Haensch) polarimeter at a sample concentration as specified in the individual experimental procedures (in g·100mL$^{-1}$). Results are reported as $[\alpha]_{D}^{T}$ values in deg·cm$^2$·g$^{-1}$ at a water bath temperature of T = 25 °C and at the wavelength of the D sodium line (λ = 589 nm).

#### General Synthetic Procedures

##### General Procedure A

A solution of *N,N′*-dicyclohexylcarbodiimide (DCC) (1.08 equiv.) in anhydrous tetrahydrofuran (0.68 M) was added dropwise to a solution of an *N*-Fmoc-protected amino acid (1.0 equiv.) and *N*-hydroxysuccinimide (NHS; 1.0 equiv.) in anhydrous tetrahydrofuran (0.19 M) at 0 °C under an atmosphere of $N_2$ gas. The reaction mixture was stirred while warming to ambient temperature overnight (14-17 h). Then, the resultant suspension was cooled to –20 °C for 1.5 h and filtered through celite®; the filtrate was concentrated under reduced pressure. The crude residue was redissolved in a minimal amount of acetone, cooled to –20 °C for 3 h, and filtered through celite®. The filtrate was concentrated under reduced pressure to afford the corresponding purified NHS-ester, which was used in the following step without further purification.

##### General Procedure B

A solution of a commercially sourced L-Lys(Boc)-OH amine (1.5 equiv.) and anhydrous $Na_2CO_3$ (1.5 equiv.) in a tetrahydrofuran:water mixture (2:1$_{v/v}$, 0.10 M) was added dropwise to a solution of an NHS-ester (1.0 equiv.) in tetrahydrofuran (0.17 M) at 0 °C. The reaction mixture was stirred until TLC analysis indicated complete conversion (~1-6 h), before being concentrated under reduced pressure. The crude residue was suspended in ethyl acetate and was subsequently washed with aqueous $NaHSO_4$ (10%$_{w/v}$; 3 times) and brine. The organic layer was dried over anhydrous $Na_2SO_4$, filtered and concentrated under reduced pressure to afford an amide product which was used in the following step without further purification.

##### General Procedure C

An amine hydrochloride salt (1.0 equiv.) was added to a solution of *N*-Boc-glycine (1.1 equiv.) in anhydrous dichloromethane (0.24 M) at ambient temperature. Sequentially, *N*-methylmorpholine (NMM) (1.1 equiv.), 1-hydroxybenzotriazole hydrate (HOBt) (1.1 equiv.) and *N*-(3-dimethylaminopropyl)-*N′*-ethylcarbodiimide (EDCl) (1.1 equiv.) were added to the reaction mixture, which was then stirred until TLC analysis indicated complete conversion (~3-6 h). The reaction mixture was concentrated under reduced pressure; the crude residue was suspended in ethyl acetate and was sequentially washed with aqueous $NaHSO_4$ (10%$_{w/v}$; 4 times), saturated aqueous

$NaHCO_3$ (3 times), and brine (3 times). The organic layer was dried over anhydrous $Na_2SO_4$, filtered and concentrated under reduced pressure to give an amide product, which was used in the following step without further purification.

**General Procedure D**

HCl in dioxane (4 M; 4.0 equiv.) was added to a solution of an *N*-Boc-protected amine (1.0 equiv.) in anhydrous dichloromethane (0.2 M) at ambient temperature under an atmosphere of $N_2$ gas. The reaction mixture was stirred overnight (15-19 h), before being concentrated under reduced pressure. The crude amine product was used immediately, without further purification, in the following reaction.

**General Procedure E**

NMM (1.5 equiv.) and isobutyl chloroformate (IBCF) (1.0 equiv.) were sequentially added to a solution of an *N*-Fmoc α-amino acid (1.0 equiv.) in tetrahydrofuran (0.06 M) at –15 °C under an atmosphere of $N_2$ gas. The reaction mixture was stirred vigorously for 10 min, before a solution of an amine hydrochloride (1.0 equiv.) in tetrahydrofuran:water ($3{:}1_{v/v}$, 0.13 M) was added. Additional NMM (1.5 equiv.) was added to the reaction mixture, which was then stirred while warming to ambient temperature over 2 h, before being concentrated under reduced pressure. The crude residue was suspended in water and the suspension was acidified by addition of aqueous $H_2SO_4$ (2 N) to pH 1 before being extracted with ethyl acetate (3 times); the combined organic extracts were sequentially washed with saturated aqueous $NaHCO_3$ and brine, dried over anhydrous $Na_2SO_4$, filtered and concentrated under reduced pressure. The crude amide product was used in the following step without further purification.

**General Procedure F**

An excess of *N*,*N*-diethylamine (~72 equiv.) was added to a suspension of an *N*-Fmoc-protected amine (1.0 equiv.) in acetonitrile (0.13 M). The reaction mixture was stirred until TLC analysis indicated complete conversion (~1-1.5 h), before being concentrated under reduced pressure and being azeotroped with acetonitrile to remove residual *N*,*N*-diethylamine (3 times). Then, 4-(*N,N*-dimethylamino)pyridine (DMAP) (1.05 equiv.), anhydrous tetrahydrofuran (0.09 M) and acetic anhydride (1.05 equiv.) or trifluoroacetic anhydride (1.05 equiv.) were added sequentially to the crude residue, and the resultant mixture was stirred under reflux for 1 h before being cooled to ambient temperature. The reaction mixture was concentrated under reduced pressure, and the crude residue was redissolved in ethyl acetate before being sequentially washed with aqueous HCl (1 M), saturated aqueous $NaHCO_3$ and brine, dried over anhydrous $Na_2SO_4$, filtered, and concentrated under reduced pressure. The residue was purified via flash column chromatography to give the corresponding amine.

**General Procedure G**

An *N*-Boc-protected amine (1.0 equiv.) was diluted in formic acid (0.04 M); the resultant mixture was stirred at ambient temperature until TLC analysis indicated complete conversion (~2.5-3 h). The reaction mixture was then concentrated under reduced pressure; the crude residue was suspended in acetonitrile (0.03 M). HCl in dioxane (4 M; 2.0 equiv.) was added to the mixture, which was then sonicated and allowed to sediment. The supernatant was removed before the residue was sonicated in acetonitrile (0.03 M) and allowed to sediment twice more. The supernatant was removed both times, and the resultant solid was dried under reduced pressure and used without further purification as the final *N*-Boc-deprotected peptide.

**General Procedure H**

To a solution or suspension of a $N^{\alpha}$-acetylated, $N^{\varepsilon}$-Boc-protected tetrapeptide alkyne (1.0 equiv.) in 2-propanol (0.05 M) was added a commercially sourced solution of 4M HCl in dioxane (~ 80 equiv.) at ambient temperature under an atmosphere of nitrogen. The reaction mixture was stirred at ambient temperature for 1 h before being concentrated under reduced pressure. The solid residue was suspended in acetonitrile (~ 0.03 M) and sonicated. The dispersed solid was allowed to sediment before the supernatant was removed. The precipitate was dried *in vacuo* to afford the corresponding $N^{\varepsilon}$-deprotected tetrapeptide alkyne as hydrochloride adduct, which was > 95% pure as judged by $^{1}H$ and $^{13}C$ NMR analysis.

#### Procedures and Characterization

##### 2,5-Dioxopyrrolidin-1-yl (((9*H*-fluoren-9-yl)methoxy)carbonyl)-*L*-leucinate (I1)

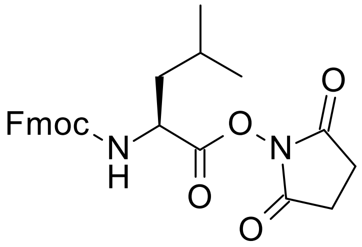


According to General Procedure A, NHS-ester **I1** (3.22 g, apparent quant.) was obtained from Fmoc-L-Leu-OH (2.50 g, 7.1 mmol). Note that $^{1}H$ NMR spectroscopy revealed the presence of minor amounts of BHT (<1%), resulting from the use of THF during the workup procedure that contained BHT as an inhibitor; the product was used without further purification.

White foam; $^{1}$**H NMR** (500 MHz, DMSO-$d_6$): $\delta$ = 8.12 (d, $J$ = 8.1 Hz, 1H), 7.89 (d, $J$ = 7.5 Hz, 2H), 7.72-7.69 (m, 2H), 7.42 (t, $J$ = 7.4 Hz, 2H), 7.32 (tt, $J$ = 7.4, 1.5 Hz, 2H), 4.42 (ddd, $J$ = 10.4, 8.0, 4.5 Hz, 1H), 4.36 (d, $J$ = 7.0 Hz, 2H), 4.24 (t, $J$ = 6.9 Hz, 1H), 2.80 (s, 4H), 1.81-1.69 (m, 2H), 1.68-1.59 (m, 1H), 0.93 (d, $J$ = 6.3 Hz, 3H), 0.88 ppm (d, $J$ = 6.3 Hz, 3H); $^{13}$**C NMR** (125 MHz, DMSO-$d_6$): $\delta$ = 170.0 (2C), 169.1, 156.0, 143.8, 143.6, 140.8 (2C), 127.7 (2C), 127.0(7), 127.0(6), 125.2 (2C), 120.2, 120.1, 65.8, 50.5, 46.6, 39.3 (determined by HSQC), 25.5 (2C), 24.1, 22.7, 21.0 ppm; **IR** (film): ṽ = 2957, 2926, 2870, 2853, 1816, 1786, 1740, 1522, 1449, 1362, 1325, 1257, 1206, 1066, 1046 $cm^{-1}$; **HRMS** (ESI): *m/z* calculated for $C_{25}H_{26}O_6N_2Na$ $[M+Na]^+$: 473.1683, found: 473.1681; $[\alpha]_D^{25}$ = –31.5 (c = 0.50, $CHCl_3$).

**2,5-Dioxopyrrolidin-1-yl (((9*H*-fluoren-9-yl)methoxy)carbonyl)-*D*-leucinate (I2)**

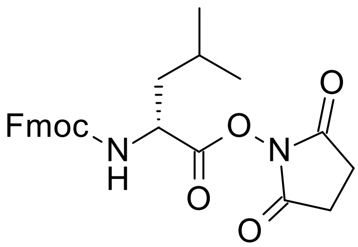


According to General Procedure A, NHS-ester **I2** (1.70 g, 95%) was obtained from Fmoc-D-Leu-OH (1.41 g, 4.0 mmol). Note that $^{1}$H NMR spectroscopy revealed the presence of minor amounts of BHT (<2%), resulting from the use of THF during the workup procedure that contained BHT as an inhibitor; the product was used without further purification.

White foam; $^{1}$**H NMR** (500 MHz, DMSO-$d_6$): $\delta$ = 8.12 (d, $J$ = 8.1 Hz, 1H), 7.89 (d, $J$ = 7.6 Hz, 2H), 7.72-7.69 (m, 2H), 7.42 (t, $J$ = 7.5 Hz, 2H), 7.32 (tt, $J$ = 7.4, 1.5 Hz, 2H), 4.43 (ddd, $J$ = 10.4, 8.0, 4.5 Hz, 1H), 4.36 (d, $J$ = 7.0 Hz, 2H), 4.24 (t, $J$ = 6.9 Hz, 1H), 2.80 (s, 4H), 1.79-1.71 (m, 2H), 1.65-1.60 (m, 1H), 0.93 (d, $J$ = 6.3 Hz, 3H), 0.88 ppm (d, $J$ = 6.3 Hz, 3H); $^{13}$**C NMR** (125 MHz, DMSO-$d_6$): $\delta$ = 170.0 (2C), 169.1, 156.0, 143.8, 143.6, 140.8 (2C), 127.7 (2C), 127.0(7), 127.0(6), 125.2 (2C), 120.2, 120.1, 65.8, 50.5, 46.6, 39.3 (determined by HSQC), 25.5 (2C), 24.1, 22.7, 21.0 ppm; **IR** (film): ṽ = 2956, 2925, 2870, 2852, 1816, 1786, 1739, 1524, 1450, 1361, 1327, 1258, 1206, 1065, 1046 $cm^{-1}$; **HRMS** (ESI): *m/z* calculated for $C_{25}H_{26}O_6N_2Na$ $[M+Na]^+$: 473.1683, found: 473.1681; $[\alpha]_D^{25}$ = +34.6 (c = 0.50, $CHCl_3$).

**2,5-Dioxopyrrolidin-1-yl (*S*)-2-((((9*H*-fluoren-9-yl)methoxy)carbonyl)amino)-3-cyclohexylpropanoate (I3)**

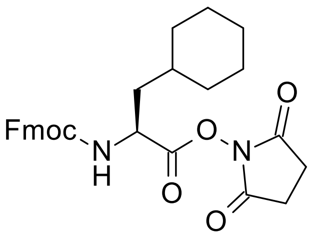


According to General Procedure A, NHS-ester **I3** (1.24 g, 99%) was obtained from Fmoc-L-Cha-OH (1.00 g, 2.6 mmol). Note that $^{1}$H NMR spectroscopy revealed the presence of minor amounts of BHT (<2%), resulting from the use of THF during the workup procedure that contained BHT as an inhibitor; the product was used without further purification.

White foam; **$^1$H NMR** (600 MHz, DMSO-$d_6$): $\delta$ = 8.11 (d, $J$ = 8.0 Hz, 1H), 7.90 (d, $J$ = 7.5 Hz, 2H), 7.71-7.70 (m, 2H), 7.42 (t, $J$ = 7.4 Hz, 2H), 7.33 (t, J = 7.5, 2H), 4.45 (ddd, $J$ = 10.3, 8.0, 4.6 Hz, 1H), 4.38-4.32 (m, 2H), 4.25 (t, $J$ = 6.9 Hz, 1H), 2.80 (s, 4H), 1.76-1.61 (m, 7H), 1.49-1.42 (m, 1H), 1.24-1.09 (m, 3H), 1.00-0.93 (m, 1H), 0.90-0.84 ppm (m, 1H); **$^{13}$C NMR** (150 MHz, DMSO-$d_6$): $\delta$ = 170.0 (2C), 169.2, 155.9, 143.8, 143.6, 140.7 (2C), 127.7 (2C), 127.1, 127.0, 125.2 (2C), 120.2, 120.1, 65.9, 49.9, 46.6, 38.0, 33.3, 32.9, 31.2, 25.9, 25.7, 25.4(9), 25.4(6) (2C) ppm; **IR** (film): ṽ = 2923, 2851, 1816, 1786, 1740, 1522, 1449, 1361, 1329, 1259, 1205, 1065, 1045 cm$^{-1}$; **HRMS** (ESI): *m/z* calculated for $C_{28}H_{30}O_6N_2Na$ [$M$+Na]$^+$: 513.1996, found: 513.1994; $[\alpha]_D^{25}$ = –22.4 (c = 0.50, $CHCl_3$).

**2,5-Dioxopyrrolidin-1-yl (*S*)-2-((((9*H*-fluoren-9-yl)methoxy)carbonyl)amino)-2-cyclohexylacetate (I4)**

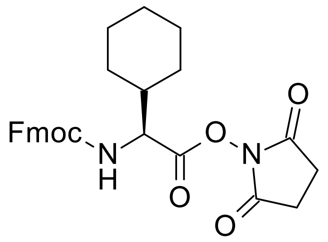


According to General Procedure A, NHS-ester **I4** (1.15 g, 89%) was obtained from Fmoc-L-Chg-OH (1.03 g, 2.7 mmol). Note that $^1$H NMR spectroscopy revealed the presence of minor amounts of BHT (<2%), resulting from the use of THF during the workup procedure that contained BHT as an inhibitor; the product was used without further purification.

White foam; **$^1$H NMR** (600 MHz, DMSO-$d_6$): $\delta$ = 8.12 (d, $J$ = 8.4 Hz, 1H), 7.89 (d, $J$ = 7.4 Hz, 2H), 7.75 (d, $J$ = 7.5 Hz, 1H), 7.73 (d, $J$ = 7.5 Hz, 1H), 7.41 (t, $J$ = 7.4 Hz, 2H), 7.36-7.29 (m, 2H), 4.36-4.30 (m, 3H), 4.24 (t, $J$ = 7.2 Hz, 1H), 2.81 (s, 4H), 1.89-1.84 (m, 1H), 1.80-1.78 (m, 1H), 1.72-1.71 (m, 3H), 1.63-1.61 (m, 1H), 1.28-1.08 (m, 5H); **$^{13}$C NMR** (150 MHz, DMSO-$d_6$): $\delta$ = 170.0 (2C), 167.8, 156.2, 143.8, 143.6, 140.7 (2C), 127.7 (2C), 127.1, 127.0, 125.3(2), 125.2(6), 120.1(1), 120.1(0), 66.0, 57.4, 46.6, 39.1 (determined by HSQC), 28.4, 27.7, 25.5 (2C), 25.4 (3C) ppm; **IR** (film): ṽ = 2925, 2853, 1815, 1785, 1740, 1527, 1515, 1450, 1358, 1331, 1297, 1247, 1205, 1064, 1033 cm$^{-1}$; **HRMS** (ESI): *m/z* calculated for $C_{27}H_{28}O_6N_2Na$ [$M$+Na]$^+$: 499.1840, found: 499.1840; $[\alpha]_D^{25}$ = –8.8 (c = 0.50, $CHCl_3$).

**$N^2$-((((9*H*-Fluoren-9-yl)methoxy)carbonyl)-*L*-leucyl)-$N^6$-(*tert*-butoxycarbonyl)-*L*-lysine (I5)**

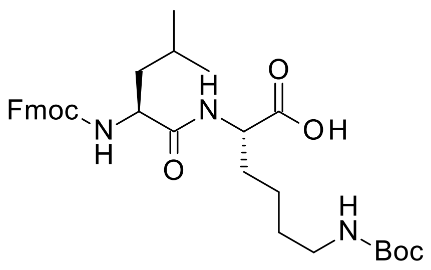

According to General Procedure B, dipeptide **I5** (1.54 g, 98%) was obtained from NHS-ester **I1** (1.22 g, 2.71 mmol). Note that partial deprotection (<2%) of the Fmoc group was observed in using $^1$H NMR spectroscopy.

White foam; **$^1$H NMR** (500 MHz, DMSO-$d_6$): $\delta$ = 12.50 (br, 1H), 8.04 (d, *J* = 7.6 Hz, 1H), 7.89 (d, *J* = 7.6 Hz, 2H), 7.72 (t, *J* = 7.4 Hz, 2H), 7.48 (d, *J* = 8.6 Hz, 1H), 7.41 (t, *J* = 7.4 Hz, 2H), 7.33-7.30 (m, 2H), 6.74 (t, *J* = 5.4 Hz, 1H), 4.31-4.19 (m, 3H), 4.15-4.06 (m, 2H), 2.89-2.85 (app.q, 2H), 1.72-1.53 (m, 3H), 1.50-1.39 (m, 2H), 1.38-1.21 (m, 13H), 0.89 (d, *J* = 6.6 Hz, 3H), 0.85 ppm (d, *J* = 6.6 Hz, 3H); **$^{13}$C NMR** (125 MHz, DMSO-$d_6$): $\delta$ = 173.5, 172.4, 155.8, 155.5, 144.0, 143.7, 140.7 (2C), 127.6 (2C), 127.0 (2C), 125.3 (2C), 120.1(0), 120.0(7), 77.31, 65.52, 52.82, 51.78, 46.69, 40.73, 39.5 (determined by HSQC), 30.7, 29.1, 28.3 (3C), 24.1, 23.1, 22.8, 21.5 ppm; **IR** (film): ṽ = 3312, 3066, 3009, 2956, 2927, 2869, 1708, 1665, 1532, 1451, 1407, 1394, 1367, 1338, 1319, 1251, 1171, 1122, 1079, 1044 cm$^{-1}$; **HRMS** (ESI): *m/z* calculated for $C_{32}H_{44}O_7N_3$ $[M+H]^+$: 582.3174, found: 582.3171; $[\alpha]_D^{25}$ = –0.2 (c = 0.50, $CHCl_3$).

**$N^2$-((((9*H*-Fluoren-9-yl)methoxy)carbonyl)-*D*-leucyl)-$N^6$-(*tert*-butoxycarbonyl)-*L*-lysine (I6)**

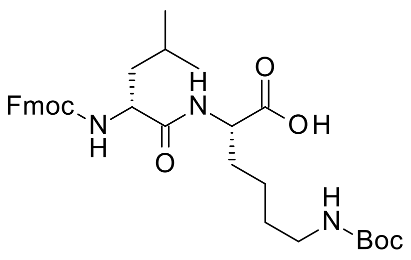

According to General Procedure B, dipeptide **I6** (605 mg, 94%) was obtained from NHS-ester **I2** (500 mg, 1.1 mmol).

White foam; **$^1$H NMR** (600 MHz, DMSO-$d_6$): $\delta$ = 12.56 (br, 1H), 8.12 (d, *J* = 7.9 Hz, 1H), 7.89 (d, *J* = 7.5 Hz, 2H), 7.73 (t, *J* = 7.6 Hz, 2H), 7.45 (d, *J* = 8.8 Hz, 1H), 7.41 (t, *J* = 7.4 Hz, 2H), 7.33-7.30 (m, 2H), 6.73 (t, *J* = 5.5 Hz, 1H), 4.29-4.19 (m, 3H), 4.16-4.11 (m, 2H), 2.90-2.81 (m, 2H), 1.70-1.64 (m, 1H), 1.64-1.53 (m, 2H), 1.51-1.46 (m, 1H), 1.42-1.30 (m, 12H), 1.27-1.18 (m, 2H), 0.89 (d, *J* = 6.6 Hz, 3H), 0.86 ppm (d, *J* = 6.6 Hz, 3fH); **$^{13}$C NMR** (150 MHz, DMSO-$d_6$): $\delta$ = 173.5, 172.3, 155.8, 155.5, 143.9, 143.7, 140.7 (2C), 127.6 (2C), 127.0(4), 127.0(1), 125.4, 125.3, 120.1 (2C), 77.3, 65.6, 52.9, 51.7, 46.7, 41.2, 39.4 (determined by HSQC), 30.8, 28.9, 28.3 (3C), 24.2, 23.0, 22.6, 21.5 ppm; **IR** (film): ṽ = 3311, 3069, 3019, 2954, 2927, 2864, 2854, 1712, 1662, 1529, 1479, 1451, 1412, 1367, 1319, 1252, 1172, 1122, 1079, 1044 cm$^{-1}$; **HRMS** (ESI): *m/z* calculated for $C_{32}H_{42}O_7N_3$ $[M-H]^-$: 580.3028, found: 580.3022; $[\alpha]_D^{25}$ = +26.2 (c = 0.50, $CHCl_3$).

**$N^2$-((*S*)-2-((((9*H*-Fluoren-9-yl)methoxy)carbonyl)amino)-3-cyclohexylpropanoyl)-$N^6$-(*tert*-butoxycarbonyl)-*L*-lysine (I7)**

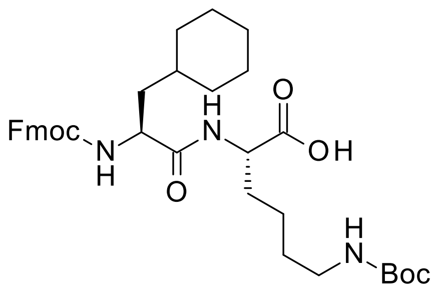

According to General Procedure B, dipeptide **I7** (823 mg, 92%) was obtained from NHS-ester **I3** (706 mg, 1.4 mmol).

White foam; **$^1$H NMR** (500 MHz, DMSO-$d_6$): $\delta$ = 12.50 (br, 1H), 8.03 (d, *J* = 7.7 Hz, 1H), 7.89 (d, *J* = 7.6 Hz, 2H), 7.73-7.71 (m, 2H), 7.48 (d, *J* = 8.5 Hz, 1H), 7.42 (t, *J* = 7.3 Hz, 2H), 7.33-7.30 (m, 2H), 6.73 (t, J = 5.4 Hz, 1H), 4.31-4.27 (m, 1H), 4.24-4.20 (m, 2H), 4.15-4.08 (m, 2H), 2.89-2.85 (app.q, 2H), 1.72-1.51 (m, 7H), 1.50-1.41 (m, 2H), 1.37-1.16 (m, 15H), 1.16-1.07 (m, 2H), 0.93-0.81 ppm (m, 2H); **$^{13}$C NMR** (125 MHz, DMSO-$d_6$): $\delta$ = 173.5, 172.5, 155.8, 155.5, 143.9, 143.8, 140.72 (2C), 127.6 (2C), 127.0 (2C), 125.3 (2C), 120.1 (2C), 77.3, 65.6, 52.2, 51.8, 46.7, 39.5 (determined by HSQC), 39.0 (determined by HSQC), 33.4, 33.2, 31.8, 30.7, 29.1, 28.3 (3C), 26.1, 25.9, 25.7, 22.7 ppm; **IR** (film): ṽ = 3309, 3063, 2925, 2852, 1701, 1669, 1533, 1524, 1478, 1450, 1407, 1395, 1367, 1277, 1249, 1171, 1134, 1107, 1082, 1043 cm$^{-1}$; **HRMS** (ESI): *m/z* calculated for $C_{35}H_{46}O_7N_3$ [*M*-H]$^-$: 620.3341, found: 620.3332; $[\alpha]_D^{25}$ = +1.7 (c = 0.50, $CHCl_3$).

**$N^2$-((*S*)-2-((((9*H*-Fluoren-9-yl)methoxy)carbonyl)amino)-2-cyclohexylacetyl)-$N^6$-(*tert*-butoxycarbonyl)-*L*-lysine (I8)**

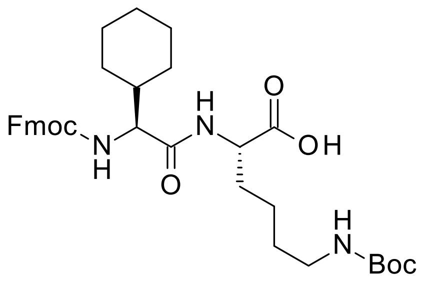

According to General Procedure B, dipeptide **I8** (435 mg, 87%) was obtained from NHS-ester **I4** (390 mg, 0.8 mmol). Note that partial deprotection (<2%) of the Fmoc group was observed using $^1$H NMR spectroscopy.

White amorphous solid; **$^1$H NMR** (500 MHz, DMSO-$d_6$): $\delta$ = 12.48 (br, 1H), 8.08 (d, *J* = 7.5 Hz, 1H), 7.89 (d, *J* = 7.6 Hz, 2H), 7.76-7.72 (app.t, 2H), 7.41 (t, *J* = 7.5 Hz, 2H), 7.36-7.30 (m, 3H), 6.73 (t, J = 5.4 Hz, 1H), 4.32-4.26 (m, 1H), 4.24-4.19 (m, 2H), 4.15-4.11 (m, 1H), 3.92 (dd, *J* = 9.1, 7.1 Hz, 1H), 2.89-2.85 (app.q, 2H), 1.71-1.53 (m, 8H), 1.37-1.21 (m, 13H), 1.19-0.93 ppm (m, 5H); **$^{13}$C NMR** (125 MHz, DMSO-$d_6$): $\delta$ = 173.5, 171.1, 156.0, 155.5, 144.0, 143.8, 140.72, 140.70, 127.6 (2C), 127.0 (2C), 125.4 (2C), 120.0(9), 120.0(6), 77.3, 65.6, 59.2, 51.8, 46.7, 39.6 (determined by HSQC), 39.5 (determined by HSQC), 30.6, 29.1, 29.0, 28.3 (3C), 28.1, 25.9, 25.6(1), 25.5(6), 22.7 ppm; **IR** (film): ṽ = 3308, 3069, 3017, 2927, 2853, 1712, 1663, 1526, 1478, 1450, 1411, 1366, 1335, 1288, 1252, 1171, 1139, 1105, 1083, 1057, 1030 cm$^{-1}$; **HRMS** (ESI): *m/z* calculated for $C_{34}H_{44}O_7N_3$ [*M*-H]$^-$: 606.3185, found: 606.3177; $[\alpha]_D^{25}$ = +8.3 (c = 0.50, $CHCl_3$).

***tert*-Butyl (2-((cyanomethyl)amino)-2-oxoethyl)carbamate** (**I9**)

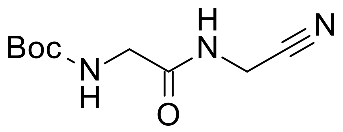

According to General Procedure C, to a solution of Boc-Gly-OH (3.00 g, 17.1 mmol, 1.0 equiv.) in THF (45 mL) were sequentially added *N*-methyl morpholine (1.88 mL, 17.1 mmol, 1.0 equiv.) and isobutyl chloroformate (2.22 mL, 17.1 mmol, 1.0 equiv.) at -25 °C under an atmosphere of nitrogen. Immediately after the formation of a white precipitate, a solution of aminoacetonitrile sulfate (1.99 g, 9.5 mmol, 1.1 equiv. of the free amine) in precooled water (1.5 mL) and 1 N NaOH (18.9 mL) was added to the mixture. The reaction mixture was allowed to warm to ambient temperature and stirred for 2h before THF was removed under reduced pressure. The residue was diluted with water, adjusted to pH 1 with 2N $H_2SO_4$ and extracted three times with ethyl acetate. The combined organic layers were washed with water (1x), a saturated aqueous solution of $NaHCO_3$ (2x) and brine (1x), dried over $Na_2SO_4$ and filtered. After evaporation of the solvent, the solid residue was recrystallized from cyclohexane / ethyl acetate to afford *tert*-butyl (2-((cyanomethyl)amino)-2-oxoethyl)carbamate **I9** (2.18 g, 60%).

White solid; **m.p.**: 113-115 °C; **$^1$H NMR** (600 MHz, DMSO-*d6*): $\delta$ = 8.50 (t, *J* = 5.4 Hz, 1H), 7.06 (t, *J* = 6.0 Hz, 1H), 4.12 (d, *J* = 5.6 Hz, 2H), 3.57 (d, *J* = 6.1 Hz, 2H), 1.38 ppm (s, 9H); **$^{13}$C NMR** (150 MHz, DMSO-*d6*): $\delta$ = 170.2, 155.8, 117.6, 78.2, 43.0, 28.2 (3C), 27.0 ppm; **IR** (solid): $\tilde{\nu}$ = 3354, 3319, 2995, 2979, 2942, 1706, 1688, 1671, 1543, 1522, 1432, 1391, 1366, 1350, 1288, 1254, 1239, 1223, 1160, 1081, 1055, 1034 cm$^{-1}$; **HRMS** (ESI): *m/z* calculated for $C_9H_{15}O_3N_3Na$ [*M*+Na]$^+$: 236.1006, found: 236.1004.

***tert*-Butyl (2-oxo-2-(prop-2-yn-1-ylamino)ethyl)carbamate (I10)**

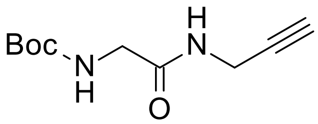

According to General Procedure C, to a solution of Boc-Gly-OH (350 mg, 2.0 mmol, 1.1 equiv.) in anhydrous dichloromethane (7.5 mL) were sequentially added propargylamine (0.12 mL, 1.8 mmol, 1.0 equiv.), N-methyl morpholine (0.22 ml, 2.0 mmol, 1.1 equiv.), HOBt hydrate (306 mg, 2.0 mmol, 1.1 equiv.) and EDCI•HCl (383 mg, 2.0 mmol, 1.1 equiv.) at ambient temperature under an atmosphere of nitrogen. The resulting suspension turned into a solution which was stirred overnight (16 h). The reaction mixture was diluted with ethyl acetate and the resulting suspension was sequentially washed with a 10% aqueous solution of $NaHSO_4$ (3x), a saturated aqueous solution of $NaHCO_3$ (3x) and brine (3x). The organic layer was dried over $Na_2SO_4$, filtered and evaporated to afford *tert*-butyl (2-oxo-2-(prop-

2-yn-1-ylamino)ethyl)carbamate **I10** (193 mg, 50%) which was used in the following step without further purification.

Off-white solid, **m.p.**: 108.0-111.5 °C; **$^{1}$H NMR** (600 MHz, DMSO-*d*$_6$): *δ* = 8.21 (t, *J* = 5.6 Hz, 1H), 6.94 (t, *J* = 6.2 Hz, 1H), 3.85 (dd, *J* = 5.5 Hz, 2.5 Hz, 2H), 3.51 (d, *J* = 6.2 Hz, 2H), 3.09 (t, *J* = 2.6 Hz, 1H), 1.38 ppm (s, 9H); **$^{13}$C NMR** (150 MHz, DMSO-*d*$_6$): *δ* = 169.1, 155.8, 81.1, 78.0, 72.9, 43.0, 28.2 (3C), 27.8 ppm; **IR** (solid): ṽ = 3314, 3297, 3059, 3009, 2981, 2931, 2919, 2850, 1704, 1677, 1677, 1542, 1473, 1454, 1420, 1389, 1368, 1293, 1255, 1235, 1172, 1071, 1050, 1031, 1018 cm$^{-1}$; **HRMS** (ESI): *m/z* calculated for $C_{10}H_{16}O_3N_2Na$ [*M*+Na]$^+$: 235.1053, found: 235.1052.

**(9*H*-Fluoren-9-yl)methyl ((*S*)-1-(((*S*)-1-cyano-15,15-dimethyl-3,6,13-trioxo-14-oxa-2,5,12-triazahexadecan-7-yl)amino)-4-methyl-1-oxopentan-2-yl)carbamate (I13)**

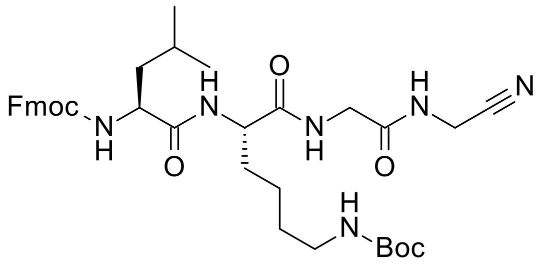


According to General Procedure E, tetrapeptide **I13** (2.01 g, 91%) along with minor impurities was obtained from dipeptide **I5** (1.91 g, 3.3 mmol) and was used in the next step without further purification. Note that partial deprotection (~5%) of the Fmoc group was observed using $^{1}$H NMR and $^{13}$C NMR spectroscopy.

Off-white amorphous solid; **$^{1}$H NMR** (600 MHz, DMSO-*d*$_6$): *δ* = 8.52 (t, *J* = 5.3 Hz, 1H), 8.29 (t, *J* = 5.8 Hz, 1H), 8.00 (d, *J* = 7.3 Hz, 1H), 7.89 (d, *J* = 7.5 Hz, 2H), 7.73-7.70 (m, 2H), 7.50 (d, *J* = 8.5 Hz, 1H), 7.41 (t, *J* = 7.4 Hz, 2H), 7.33-7.31 (m, 2H), 6.71 (t, *J* = 5.5 Hz, 1H), 4.36-4.17 (m, 4H), 4.17-4.00 (m, 3H), 3.76 (dd, *J* = 16.8, 6.1 Hz, 1H), 3.70 (dd, *J* = 16.8, 5.7 Hz, 1H), 2.88-2.84 (m, 2H), 1.73- 1.57 (m, 2H), 1.56-1.50 (m, 1H), 1.49-1.39 (m, 2H), 1.38-1.29 (m, 11H), 1.29-1.17 (m, 2H, overlap with an impurity signal at 1.23 ppm), 0.88 (d, *J* = 6.6 Hz, 3H), 0.85 ppm (d, *J* = 6.5 Hz, 3H); **$^{13}$C NMR** (150 MHz, DMSO-*d*$_6$): *δ* = 172.6, 172.0, 169.6, 155.9, 155.5, 143.9, 143.7, 140.7 (2C), 127.6 (2C), 127.0 (2C), 125.3 (2C), 120.1, 120.08, 117.5, 77.3, 65.5, 52.9, 52.8, 46.7, 41.7, 40.6, 39.5 (determined by HSQC), 31.4, 29.2, 28.3 (3C), 27.0, 24.2, 23.1, 22.5, 21.4 ppm; **IR** (solid): ṽ = 3293, 3062, 2956, 2936, 2868, 1688, 1659, 1646, 1530, 1479, 1450, 1404, 1392, 1367, 1336, 1320, 1250, 1171, 1120, 1079, 1044 cm$^{-1}$; **HRMS** (ESI): *m/z* calculated for $C_{36}H_{49}O_7N_6$ [*M*+H]$^+$: 677.3657, found: 677.3658; $[\alpha]_D^{25}$ = –17.2 (c = 0.50, MeOH).

**(9*H*-Fluoren-9-yl)methyl ((*R*)-1-(((*S*)-1-cyano-15,15-dimethyl-3,6,13-trioxo-14-oxa-2,5,12-triazahexadecan-7-yl)amino)-4-methyl-1-oxopentan-2-yl)carbamate (I14)**

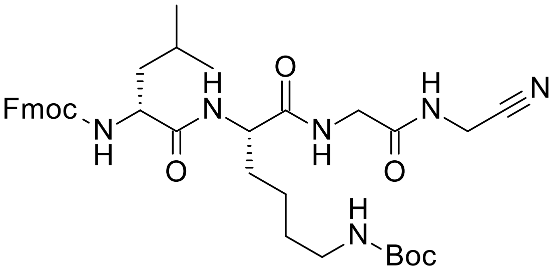


According to General Procedure E, tetrapeptide **I14** (378 mg, 61%) along with minor impurities was obtained from dipeptide **I6** (535 mg, 0.9 mmol), following column chromatography (25 g Sfär; 80 mL/min; initially, 100% dichloromethane (3 CV), followed by a linear gradient (15 CV): $0\%_{v/v} \rightarrow 50\%_{v/v}$ methanol in dichloromethane). Note that partial deprotection (~5%) of the Fmoc group was using $^1$H NMR spectroscopy.

Off-white amorphous solid; **$^1$H NMR** (600 MHz, DMSO-*d*$_6$): $\delta$ = 8.47 (t, *J* = 5.6 Hz, 1H), 8.23 (t, *J* = 6.0 Hz, 1H), 8.20 (d, *J* = 7.7 Hz, 1H), 7.89 (d, *J* = 7.6 Hz, 2H), 7.73-7.70 (app.t, 2H), 7.54 (d, *J* = 8.0 Hz, 1H), 7.41 (t, *J* = 7.4 Hz, 2H), 7.33-7.30 (m, 2H), 6.71 (t, *J* = 5.3 Hz, 1H), 4.32-4.17 (m, 4H), 4.15-4.08 (m, 3H), 3.75 (dd, *J* = 16.7, 6.0 Hz, 1H), 3.70 (dd, *J* = 16.7, 5.8 Hz, 1H), 2.89-2.81 (m, 2H), 1.70-1.62 (m, 1H), 1.61-1.56 (m, 1H), 1.55-1.40 (m, 3H), 1.39-1.27 (m, 11H), 1.26-1.15 (m, 2H, overlap with an impurity signal at 1.23 ppm), 0.89 (d, *J* = 6.5 Hz, 3H), 0.86 ppm (d, *J* = 6.5 Hz, 3H); **$^{13}$C NMR** (150 MHz, DMSO-*d*$_6$): $\delta$ = 172.7, 172.0, 169.5, 156.1, 155.5, 143.9, 143.7, 140.7(2), 140.7(0), 127.6 (2C), 127.1, 127.0, 125.3 (2C), 120.1, 120.0(8), 117.5, 77.3, 65.6, 53.1, 52.6, 46.7, 41.7, 40.7, 39.5 (determined by HSQC), 31.3, 29.1, 28.3 (3C), 27.0, 24.2, 22.9, 22.6, 21.6 ppm; **IR** (solid): $\tilde{\nu}$ = 3294, 3067, 2930, 2854, 1689, 1639, 1532, 1450, 1391, 1366, 1342, 1283, 1254, 1234, 1171, 1137, 1085, 1032 cm$^{-1}$; **HRMS** (ESI): *m/z* calculated for $C_{36}H_{49}O_7N_6$ [*M*+H]$^+$: 677.3657, found: 677.3660; $[\boldsymbol{\alpha}]_{\boldsymbol{D}}^{\mathbf{25}}$ = +12.3 (c = 0.50, MeOH).

**(9*H*-Fluoren-9-yl)methyl ((*S*)-1-(((*S*)-1-cyano-15,15-dimethyl-3,6,13-trioxo-14-oxa-2,5,12-triazahexadecan-7-yl)amino)-3-cyclohexyl-1-oxopropan-2-yl)carbamate (I15)**

According to General Procedure E, tetrapeptide **I15** (554 mg, 95%) was obtained from dipeptide **I7** (497 mg, 0.8 mmol) and was used in the next step without further purification. Note that partial deprotection (~5%) of the Fmoc group was observed using $^1$H NMR spectroscopy.

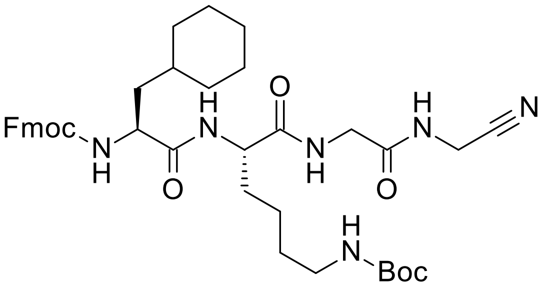


Off-white amorphous solid; **$^1$H NMR** (600 MHz, DMSO-*d*$_6$): $\delta$ = 8.51 (t, *J* = 5.6 Hz, 1H), 8.27 (t, *J* = 5.8 Hz, 1H), 7.98 (d, *J* = 7.3 Hz, 1H), 7.89 (d, *J* = 7.6 Hz, 2H), 7.73-7.70 (app.t, 2H), 7.51 (d, *J* = 8.4 Hz, 1H), 7.42 (t, *J* = 7.4 Hz, 2H), 7.34-7.30 (m, 2H), 6.71

(t, $J$ = 5.3 Hz, 1H), 4.35-4.27 (m, 1H), 4.26-4.16 (m, 3H), 4.14 (d, $J$ = 5.6 Hz, 2H), 4.11-4.07 (m, 1H), 3.76 (dd, $J$ = 16.8, 5.9 Hz, 1H), 3.70 (dd, $J$ = 16.8, 5.5 Hz, 1H), 2.88-2.83 (m, 2H), 1.72-1.56 (m, 6H), 1.55-1.42 (m, 3H), 1.39-1.28 (m, 12H), 1.27-1.14 (m, 3H, overlap with an impurity signal at 1.23 ppm), 1.14-1.07 (m, 2H), 0.93-0.81 ppm (m, 2H); **$^{13}$C NMR** (150 MHz, DMSO-$d_6$): $\delta$ = 172.7, 172.0, 169.5, 155.9, 155.5, 143.9, 143.7, 140.7 (2C), 127.6 (2C), 127.0 (2C), 125.3 (2C), 120.1 (2C), 117.5, 77.3, 65.5, 52.7, 52.3, 46.6, 41.7, 39.5 (determined by HSQC), 39.0 (determined by HSQC), 33.5, 33.3, 31.6, 31.4, 29.2, 28.2 (3C), 27.0, 26.1, 25.8, 25.6, 22.5 ppm; **IR** (solid): ṽ = 3307, 3066, 2925, 2852, 1648, 1525, 1479, 1449, 1404, 1393, 1366, 1336, 1274, 1248, 1170, 1133, 1104, 1082, 1042 cm$^{-1}$; **HRMS** (ESI): *m/z* calculated for $C_{39}H_{53}O_7N_6$ [$M$+H]$^+$: 717.3970, found: 717.3972; $[\alpha]_D^{25}$ = –12.4 (c = 0.50, MeOH).

**(9*H*-Fluoren-9-yl)methyl ((*S*)-2-(((*S*)-1-cyano-15,15-dimethyl-3,6,13-trioxo-14-oxa-2,5,12-triazahexadecan-7-yl)amino)-1-cyclohexyl-2-oxoethyl)carbamate (I16)**

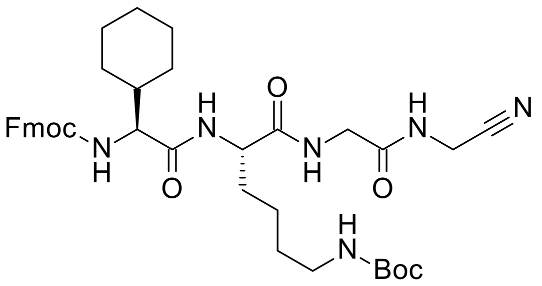


According to General Procedure E, tetrapeptide **I16** (140 mg, 33%) along with minor impurities was obtained from dipeptide **I8** (365 mg, 0.6 mmol) following column chromatography (10 g Sfär; 35 mL/min; initially, 100% ethyl acetate (7 CV), followed by a linear gradient (15 CV): 0%$_{v/v}$ → 100%$_{v/v}$ acetone in ethyl acetate). Note that partial deprotection (~15%) of the Fmoc group was observed by $^1$H NMR and $^{13}$C NMR spectroscopy.

White amorphous solid; **$^1$H NMR** (600 MHz, DMSO-$d_6$): $\delta$ = 8.53 (t, $J$ = 5.7 Hz, 1H), 8.30 (t, $J$ = 5.6 Hz, 1H), 8.06 (d, 6.9 Hz, 1H), 7.89 (d, $J$ = 7.8 Hz, 2H), 7.74 (d, $J$ = 7.7 Hz, 1H), 7.72 (d, $J$ = 7.9 Hz, 1H), 7.43-7.39 (m, 3H), 7.33-7.31 (m, 2H), 6.71 (t, $J$ = 5.4 Hz, 1H), 4.33 (dd, $J$ = 9.8, 6.7 Hz, 1H), 4.26-4.21 (m, 2H), 4.19-4.13 (m, 3H), 3.93-3.90 (app.t, 1H), 3.77 (dd, $J$ = 16.9, 6.2 Hz, 1H), 3.68 (dd, $J$ = 16.8, 5.6 Hz, 1H), 2.88-2.84 (app.q, 2H), 1.87-1.47 (m, 8H), 1.39-1.29 (m, 11H), 1.29-1.18 (m, 2H, overlap with an impurity signal at 1.23 ppm), 1.18-1.03 (m, 3H), 1.01-0.92 ppm (m, 2H); **$^{13}$C NMR** (150 MHz, DMSO-$d_6$): $\delta$ = 172.0, 171.4, 169.6, 156.0, 155.5, 143.9, 143.7, 140.7(3), 140.7(1), 127.6 (2C), 127.0 (2C), 125.3 (2C), 120.1, 120.0, 117.5, 77.3, 65.5, 59.3, 52.9, 46.7, 41.7, 39.7 (determined by HSQC), 39.6 (determined by HSQC), 31.2, 29.2, 29.0, 28.3 (3C), 28.0, 27.0, 25.8, 25.6 (2C), 22.5 ppm; **IR** (solid): ṽ = 3294, 3066, 2929, 2854, 1690, 1641, 1531, 1450, 1392, 1366, 1335, 1284, 1254, 1233,

1170, 1138, 1103, 1085, 1054, 1030 cm$^{-1}$; **HRMS** (ESI): *m/z* calculated for $C_{38}H_{50}O_7N_6Na$ [*M*+Na]$^+$: 725.3633, found: 725.3629; $[\alpha]_D^{25}$ = –7.2 (c = 0.50, MeOH).

**(9*H*-Fluoren-9-yl)methyl ((*S*)-1-(((*S*)-2,2-dimethyl-4,11,14-trioxo-3-oxa-5,12,15-triazaoctadec-17-yn-10-yl)amino)-4-methyl-1-oxopentan-2-yl)carbamate (I17)**

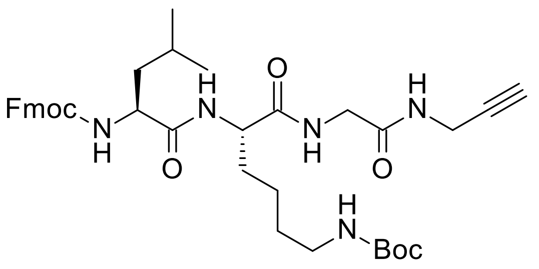


According to General Procedure E, tetrapeptide **I17** (252 mg, 89%) was obtained from dipeptide **I5** (244 mg, 0.4 mmol) and was used in the next step without further purification. Note that partial deprotection (~3%) of the Fmoc group was observed using $^1$H NMR spectroscopy.

White amorphous solid; $^1$**H NMR** (600 MHz, DMSO-*d*$_6$): $\delta$ = 8.22 (t, *J* = 5.5 Hz, 1H), 8.16 (t, *J* = 5.8 Hz, 1H), 7.94 (d, *J* = 7.5 Hz, 1H), 7.89 (d, *J* = 7.6 Hz, 2H), 7.73-7.70 (app.t, 2H), 7.51 (d, *J* = 8.3 Hz, 1H), 7.41 (t, *J* = 7.4 Hz, 2H), 7.33-7.30 (m, 2H), 6.71 (t, *J* = 5.3 Hz, 1H), 4.33-4.30 (m, 1H), 4.27-4.25 (m, 1H), 4.23-4.18 (m, 2H), 4.08-4.04 (m, 1H), 3.87 (ddd, *J* = 17.5, 5.6, 2.5 Hz, 1H), 3.84 (ddd, *J* = 17.5, 5.6, 2.5 Hz, 1H), 3.70 (dd, *J* = 16.8, 6.1 Hz, 1H), 3.65 (dd, *J* = 16.7, 5.7 Hz, 1H), 3.09 (t, *J* = 2.5 Hz, 1H), 2.88-2.84 (app.q, 2H), 1.68-1.57 (m, 2H), 1.54-1.40 (m, 3H), 1.39-1.29 (m, 11H), 1.27-1.16 (m, 2H), 0.88 (d, *J* = 6.6 Hz, 3H), 0.85 ppm (d, *J* = 6.5 Hz, 3H); $^{13}$**C NMR** (150 MHz, DMSO-*d*$_6$): $\delta$ = 172.4, 171.8, 168.5, 155.9, 155.5, 143.9, 143.7, 140.7 (2C), 127.6 (2C), 127.0 (2C), 125.3 (2C), 120.1, 120.0(7), 81.0, 77.3, 73.0, 65.5, 53.0, 52.6, 46.7, 41.8, 40.6, 39.5 (determined by HSQC), 31.5, 29.2, 28.3 (3C), 27.9, 24.1, 23.1, 22.5, 21.4 ppm; **IR** (solid): ṽ = 3292, 3067, 2930, 2866, 1708, 1681, 1632, 1522, 1464, 1450, 1392, 1366, 1335, 1280, 1254, 1233, 1172, 1121, 1081, 1044 cm$^{-1}$; **HRMS** (ESI): *m/z* calculated for $C_{37}H_{50}O_7N_5$ [*M*+H]$^+$: 676.3705, found: 676.3697; $[\alpha]_D^{25}$ = –17.4 (c = 0.25, MeOH).

**(9*H*-Fluoren-9-yl)methyl ((*R*)-1-(((*S*)-2,2-dimethyl-4,11,14-trioxo-3-oxa-5,12,15-triazaoctadec-17-yn-10-yl)amino)-4-methyl-1-oxopentan-2-yl)carbamate (I18)**

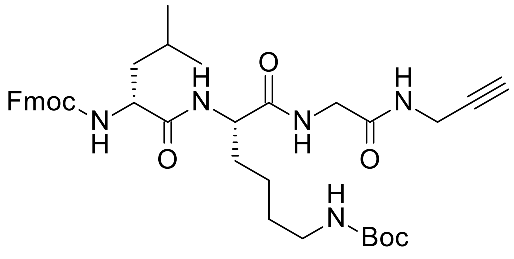


According to General Procedure E, tetrapeptide **I18** (496 mg, 94%) along with minor impurities was obtained from dipeptide **I6** (454 mg, 0.8 mmol) and was used in the following step without further purification.

White amorphous solid; $^1$**H NMR** (400 MHz, DMSO-*d*$_6$): $\delta$ = 8.18-8.11 (m, 3H), 7.89 (d, *J* = 7.6 Hz, 2H), 7.73-7.70 (app.t, 2H), 7.53 (d, *J* = 7.9 Hz, 1H), 7.41 (t, *J* =

7.4 Hz, 2H), 7.32 (tt, *J* = 7.4, 1.4 Hz, 2H), 6.71 (t, *J* = 5.3 Hz, 1H), 4.32-4.15 (m, 4H), 4.11-4.05 (m, 1H), 3.86 (ddd, *J* = 17.5, 5.7, 2.5 Hz, 1H), 3.80 (ddd, *J* = 17.5, 5.6, 2.5 Hz, 1H), 3.70 (dd, *J* = 16.4, 5.6 Hz, 1H), 3.65 (dd, *J* =16.5, 5.7 Hz, 1H), 3.07 (t, *J* = 2.5 Hz, 1H), 2.90-2.81 (m, 2H), 1.70-1.42 (m, 5H), 1.41-1.29 (m, 11H), 1.27- 1.14 (m, 2H), 0.89 (d, *J* = 6.5 Hz, 3H), 0.86 ppm (d, *J* = 6.5 Hz, 3H); **$^{13}$C NMR** (150 MHz, DMSO-$d_6$): $\delta$ = 172.6, 171.9, 168.4, 156.1, 155.5, 143.9, 143.7, 140.7 (2C), 127.6 (2C), 127.1 (2C), 125.3 (2C), 120.1, 120.0(8), 80.9, 77.3, 73.0, 65.6, 53.2, 52.6, 46.7, 41.8, 40.7, 39.5 (determined by HSQC), 31.3, 29.0, 28.3 (3C), 27.8, 24.2, 22.9, 22.5, 21.6 ppm; **IR** (solid): ṽ = 3295, 3066, 2924, 2853, 1685, 1638, 1534, 1478, 1449, 1404, 1391, 1366, 1322, 1277, 1250, 1235, 1172, 1123, 1105, 1083, 1040 cm$^{-1}$; **HRMS** (ESI): *m/z* calculated for $C_{37}H_{50}O_7N_5$ $[M+H]^+$: 676.3705, found: 676.3707; $[\alpha]_D^{25}$ = +11.4 (c = 0.50, MeOH).

**(9*H*-Fluoren-9-yl)methyl ((*S*)-3-cyclohexyl-1-(((*S*)-2,2-dimethyl-4,11,14-trioxo-3-oxa-5,12,15-triazaoctadec-17-yn-10-yl)amino)-1-oxopropan-2-yl)carbamate (I19)**

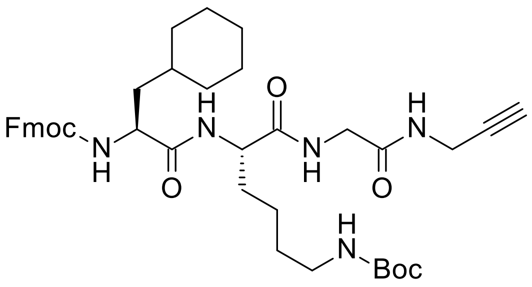


According to General Procedure E, tetrapeptide **I19** (536 mg, 97%) along with minor impurities was obtained from dipeptide **I7** (482 mg, 0.8 mmol) and was used in the following step without further purification. Note that partial deprotection (~10%) of the Fmoc group was observed by $^1$H NMR and $^{13}$C NMR spectroscopy.

White amorphous solid; **$^1$H NMR** (600 MHz, DMSO-$d_6$): $\delta$ = 8.23 (t, *J* = 5.5 Hz, 1H), 8.15 (t, *J* = 5.8 Hz, 1H), 7.92 (d, *J* = 7.6 Hz, 1H), 7.89 (d, *J* = 7.5 Hz, 2H), 7.72-7.70 (app.t, 2H), 7.52 (d, *J* = 8.3 Hz, 1H), 7.42 (t, *J* = 7.4 Hz, 2H), 7.33-7.31 (m, 2H), 6.71 (t, *J* = 5.3 Hz, 1H), 4.34-4.30 (m, 1H), 4.26-4.18 (m, 3H), 4.10-4.06 (m, 1H), 3.87 (ddd, *J* = 17.5, 5.5, 2.5 Hz, 1H), 3.84 (ddd, *J* = 17.4, 5.5, 2.5 Hz, 1H), 3.71 (dd, *J* = 16.7, 6.1 Hz, 1H), 3.65 (dd, *J* = 16.6, 5.7 Hz, 1H), 3.09 (t, *J* = 2.4 Hz, 1H), 2.87-2.84 (m, 2H), 1.72-1.56 (m, 6H), 1.55-1.41 (m, 3H), 1.39-1.28 (m, 12H), 1.27-1.16 (m, 3H), 1.15-1.07 (m, 2H), 0.93-0.78 ppm (m, 2H); **$^{13}$C NMR** (150 MHz, DMSO-$d_6$): $\delta$ = 172.5, 171.8, 168.5, 155.9, 155.5, 143.9, 143.7, 140.7(2), 140.6(9), 127.6 (2C), 127.1, 127.0, 125.2(8), 125.2(6), 120.1(2), 120.0(9), 81.0, 77.3, 73.0, 65.6, 52.6, 52.4, 46.7, 41.7, 39.6 (determined by HSQC), 39.0 (determined by HSQC), 33.5, 33.3, 31.7, 31.5, 29.2, 28.3 (3C), 27.9, 26.1, 25.9, 25.7, 22.5 ppm; **IR** (solid): ṽ = 3296, 3068, 2924, 2853, 1686, 1638, 1534, 1448, 1404, 1366, 1311, 1276, 1250, 1229, 1172, 1134, 1104, 1087, 1042 cm$^{-1}$; **HRMS**

(ESI): *m/z* calculated for $C_{40}H_{54}O_7N_5$ $[M+H]^+$: 716.4018, found: 716.4020; $[\alpha]_D^{25}$ = –14.1 (c = 0.50, MeOH).

**_tert_-Butyl ((_S_)-5-((_S_)-2-acetamido-4-methylpentanamido)-6-((2-((cyanomethyl)amino)-2-oxoethyl)amino)-6-oxohexyl)carbamate (I20)**

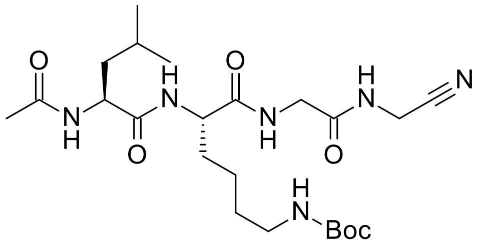


According to General Procedure F, *N*-acetylated peptide **I20** (255 mg, 35%) was obtained from *N*-Fmoc-protected peptide **I13** (1.00 g, 1.5 mmol), following column chromatography (25 g Sfär; 80 mL/min; initially, 100% ethyl acetate (8 CV), followed by a linear gradient (20 CV): $0\%_{v/v}$ → $100\%_{v/v}$ acetone in ethyl acetate).

White amorphous solid; $^{1}$**H NMR** (600 MHz, DMSO-$d_6$): $\delta$ = 8.48 (t, *J* = 5.6 Hz, 1H), 8.20 (t, *J* = 5.9 Hz, 1H), 7.99 (d, *J* = 7.9 Hz, 1H), 7.98 (d, *J* =7.3 Hz, 1H), 6.75 (t, *J* = 5.4 Hz, 1H), 4.30-4.27, (m, 1H), 4.17-4.11 (m, 3H), 3.76 (dd, *J* = 16.8, 6.1 Hz, 1H), 3.69 (dd, 16.8, 5.6 Hz, 1H), 2.89-2.85 (app.q, 2H), 1.84 (s, 3H), 1.68-1.62 (m, 1H), 1.61-1.57 (m, 1H), 1.56-1.49 (m, 2H), 1.47-1.39 (m, 2H), 1.39-1.29 (m, 11H), 1.29-1.16 (m, 2H), 0.88 (d, *J* = 6.6 Hz, 3H), 0.84 ppm (d, *J* = 6.6 Hz, 3H); $^{13}$**C NMR** (150 MHz, DMSO-$d_6$): $\delta$ = 172.5, 172.0, 169.6, 169.4, 155.5, 117.5, 77.3, 52.7, 51.0, 41.7, 40.5, 39.6 (determined by HSQC), 31.3, 29.2, 28.3 (3C), 27.0, 24.2, 23.1, 22.6, 22.5, 21.6 ppm; **IR** (solid): ṽ = 3285, 3067, 2957, 2936, 2870, 1684, 1631, 1539, 1457, 1393, 1367, 1279, 1251, 1172, 1035 cm$^{-1}$; **HRMS** (ESI): *m/z* calculated for $C_{23}H_{41}O_6N_6$ $[M+H]^+$: 497.3082, found: 497.3077; $[\alpha]_D^{25}$ = –21.0 (c = 0.50, MeOH).

**_tert_-Butyl ((_S_)-5-((_R_)-2-acetamido-4-methylpentanamido)-6-((2-((cyanomethyl)amino)-2-oxoethyl)amino)-6-oxohexyl)carbamate (I21)**

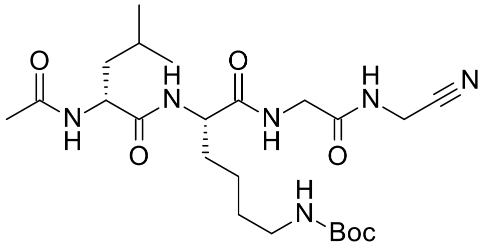


According to General Procedure F, *N*-acetylated peptide **I21** (86 mg, 43%) was obtained from *N*-Fmoc-protected peptide **I14** (357 mg, 0.5 mmol), following column chromatography (10 g Sfär; 35 mL/min; initially, 100% ethyl acetate (7 CV), followed by a linear gradient (15 CV): $0\%_{v/v}$ → $100\%_{v/v}$ acetone in ethyl acetate).

Off-white amorphous solid; $^{1}$**H NMR** (600 MHz, DMSO-$d_6$): $\delta$ = 8.40 (t, *J* = 5.6 Hz, 1H), 8.31 (d, *J* = 7.7 Hz, 1H), 8.22 (t, *J* = 6.0 Hz, 1H), 8.04 (t, *J* = 7.3 Hz, 1H), 6.73 (t, *J* = 5.4 Hz, 1H), 4.28 (q, *J* = 7.5 Hz, 1H), 4.18-4.11 (m, 3H), 3.74 (dd, *J* =16.8, 6.0 Hz, 1H), 3.70 (dd, *J* = 16.7, 6.0 Hz, 1H), 2.91-2.83 (m, 2H),

1.83 (s, 3H), 1.70-1.64 (m, 1H), 1.59-1.49 (m, 2H), 1.42 (t, $J$ = 7.3 Hz, 2H), 1.39-1.29 (m, 11H), 1.28-1.16 (m, 2H, ), 0.90 (d, $J$ = 6.5 Hz, 3H), 0.85 ppm (d, $J$ = 6.5 Hz, 3H); **$^{13}$C NMR** (150 MHz, DMSO-$d_6$): $\delta$ = 172.7, 172.1, 169.6, 169.5, 155.5, 117.5, 77.3, 52.7, 51.4, 41.7, 40.8, 39.4 (determined by HSQC), 30.9, 29.0, 28.3 (3C), 27.0, 24.2, 22.7, 22.6, 22.4, 21.9 ppm; **IR** (solid): ṽ = 3284, 3067, 2930, 2853, 1656, 1630, 1535, 1439, 1406, 1391, 1367, 1278, 1250, 1172, 1105, 1038, 1017 cm$^{-1}$; **HRMS** (ESI): *m/z* calculated for $C_{23}H_{39}O_6N_6$ [$M$-H]$^-$: 495.2937, found: 495.2937; $[\alpha]_D^{25}$ = +1.2 (c = 0.25, MeOH).

***tert*-Butyl ((*S*)-5-((*S*)-2-acetamido-3-cyclohexylpropanamido)-6-((2-((cyanomethyl)amino)-2-oxoethyl)amino)-6-oxohexyl)carbamate (I22)**

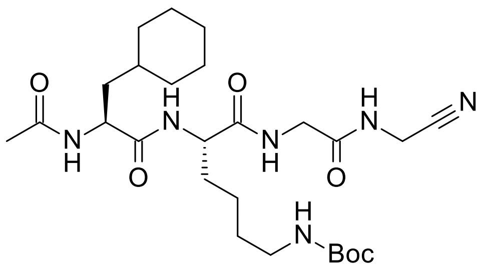


According to General Procedure F, *N*-acetylated peptide **I22** (178 mg, 46%) was obtained from *N*-Fmoc-protected peptide **I15** (717 mg, 0.7 mmol), following column chromatography (25 g Sfär; 80 mL/min; initially, 100% ethyl acetate (7 CV), followed by a linear gradient (15 CV): 0%$_{v/v}$ → 100%$_{v/v}$ acetone in ethyl acetate).

White amorphous solid; **$^1$H NMR** (600 MHz, DMSO-$d_6$): $\delta$ = 8.48 (t, $J$ = 5.6 Hz, 1H), 8.18 (t, $J$ = 5.9 Hz, 1H), 7.99 (d, $J$ = 7.9 Hz, 1H), 7.96 (d, $J$ = 7.5 Hz, 1H), 6.74 (t, $J$ = 5.4 Hz, 1H), 4.29 (ddd, $J$ = 9.7, 7.9, 5.1 Hz, 1H), 4.17-4.13 (m, 3H), 3.76 (dd, $J$ = 16.8, 6.1 Hz, 1H), 3.69 (dd, $J$ =16.8, 5.6 Hz, 1H), 2.89-2.85 (app.q, 2H), 1.84 (s, 3H), 1.70-1.57 (m, 6H), 1.55-1.50 (m, 1H), 1.49-1.44 (m, 1H), 1.42-1.32 (m, 12H), 1.29-1.23 (m, 2H), 1.22-1.15 (m, 2H), 1.15-1.07 (m, 2H), 1.14-1.07 ppm (m, 2H); **$^{13}$C NMR** (150 MHz, DMSO-$d_6$): $\delta$ = 172.6, 172.0, 169.6, 169.4, 155.5, 117.5, 77.3, 52.7, 50.4, 41.7, 39.5 (determined by HSQC), 39.1 (determined by HSQC), 33.5, 33.2, 31.9, 31.3, 29.2, 28.3 (3C), 27.0, 26.1, 25.8, 25.6, 22.6, 22.5 ppm; **IR** (solid): ṽ = 3285, 3075, 2924, 2852, 1679, 1631, 1536, 1445, 1393, 1367, 1279, 1251, 1229, 1173, 1038, 1016 cm$^{-1}$; **HRMS** (ESI): *m/z* calculated for $C_{26}H_{45}O_6N_6$ [$M$+H]$^+$: 537.3395, found: 537.3391; $[\alpha]_D^{25}$ = –10.3 (c = 0.50, MeOH).

***tert*-Butyl ((*S*)-5-((*S*)-2-acetamido-2-cyclohexylacetamido)-6-((2-((cyanomethyl)amino)-2-oxoethyl)amino)-6-oxohexyl)carbamate (I23)**

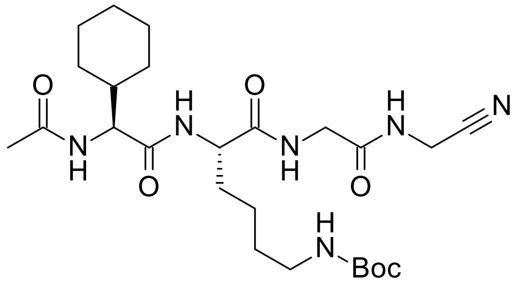

According to a modification of General Procedure F using 2 equiv. of both DMAP (44 mg, 0.36 mmol) and acetic anhydride (34 µL, 0.36 mmol), *N*-acetylated peptide **I23** (22 mg, 23%) along with minor impurities was obtained from *N*-Fmoc-protected peptide **I16** (125 mg, 0.18 mmol), following column chromatography (5 g Sfär; 18 mL/min; initially, 100% ethyl acetate (7 CV), followed by a linear gradient (15 CV): $0\%_{v/v} \rightarrow 100\%_{v/v}$ acetone in ethyl acetate).

Orange amorphous solid; $^1$**H NMR** (600 MHz, DMSO-$d_6$): $\delta$ = 8.50 (t, $J$ = 5.6 Hz, 1H), 8.21 (t, $J$ = 5.8 Hz, 1H), 8.05 (d, $J$ = 7.0 Hz, 1H), 7.87 (d, $J$ = 8.5 Hz, 1H), 6.74 (t, $J$ = 5.5 Hz, 1H), 4.17-4.10 (m, 4H), 3.77 (dd, $J$ = 16.8, 6.1 Hz, 1H), 3.68 (dd, $J$ = 16.8, 5.4 Hz, 1H), 2.88-2.85 (app.q, 2H), 1.86 (s, 3H), 1.67-1.49 (m, 8H), 1.40-1.31 (m, 11H), 1.30-1.17 (m, 2H, overlap with an impurity signal at 1.23 ppm), 1.16-1.03 (m, 3H), 1.01-0.92 ppm (m, 2H); $^{13}$**C NMR** (150 MHz, DMSO-$d_6$): $\delta$ = 172.0, 171.4, 169.6, 169.4, 155.5, 117.5, 77.3, 57.2, 52.9, 41.8, 39.6 (determined by HSQC), 39.5 (determined by HSQC), 31.1, 29.2, 29.0, 28.3 (3C), 28.1, 27.0, 25.8, 25.6 (2C), 22.6, 22.5 ppm; **IR** (solid): $\tilde{\nu}$ = 3284, 3079, 2926, 2853, 1684, 1656, 1629, 1536, 1440, 1392, 1367, 1279, 1251, 1205, 1173, 1145, 1107, 1037, 1017 cm$^{-1}$; **HRMS** (ESI): *m/z* calculated for $C_{25}H_{42}O_6N_6Na$ [$M$+Na]$^+$: 545.3058, found: 545.3054; $[\alpha]_D^{25}$ = –11.0 (c = 0.125, MeOH).

***tert*-Butyl ((*S*)-5-((*S*)-2-acetamido-4-methylpentanamido)-6-oxo-6-((2-oxo-2-(prop-2-yn-1-ylamino)ethyl)amino)hexyl)carbamate (I24)**

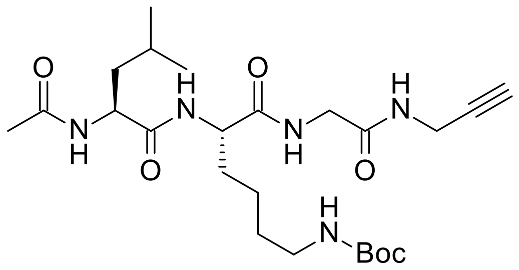

According to General Procedure F, *N*-acetylated peptide **I24** (59 mg, 40%) was obtained from *N*-Fmoc-protected peptide **I17** (200 mg, 0.3 mmol), following column chromatography (10 g Sfär; 35 mL/min; initially, 100% ethyl acetate (10 CV), followed by a linear gradient (20 CV): $0\%_{v/v} \rightarrow 100\%_{v/v}$ acetone in ethyl acetate).

White amorphous solid; $^1$**H NMR** (600 MHz, DMSO-$d_6$): $\delta$ = 8.19 (t, $J$ = 5.6 Hz, 1H), 8.10 (t, $J$ = 5.9 Hz, 1H), 7.99 (d, $J$ = 8.0 Hz, 1H), 7.94 (d, $J$ = 7.5 Hz, 1H), 6.74 (t, $J$ = 5.8 Hz, 1H), 4.29-4.26 (m, 1H), 4.17-4.14 (m, 1H), 3.88 (ddd, $J$ = 17.5, 5.6, 2.6 Hz, 1H), 3.84 (ddd, $J$ = 17.5, 5.6, 2.5 Hz, 1H), 3.70 (dd, $J$ = 16.6, 6.1 Hz, 1H), 3.64 (dd, $J$ = 16.7, 5.7 Hz, 1H), 3.10 (t, $J$ = 2.5 Hz, 1H), 2.89-2.85 (app.q, 2H), 1.84 (s, 3H), 1.68-1.62 (m, 1H), 1.61-1.56 (m, 1H), 1.54-1.46 (m, 1H), 1.45-1.40 (m, 2H), 1.39-1.29 (m,

11H), 1.28-1.16 (m, 2H), 0.88 (d, $J$ = 6.6 Hz, 3H), 0.84 ppm (d, $J$ = 6.6 Hz, 3H); **$^{13}$C NMR** (150 MHz, DMSO-$d_6$): $\delta$ = 172.4, 171.8, 169.4, 168.5, 155.5, 81.0, 77.3, 73.0, 52.7, 51.0, 41.8, 40.5, 39.5 (determined by HSQC), 31.4, 29.2, 28.3, 27.9, 24.1, 23.1, 22.6, 22.5, 21.6 ppm; **IR** (solid): ṽ = 3289, 3078, 2957, 2934, 2870, 1632, 1532, 1460, 1440, 1391, 1391, 1367, 1277, 1250, 1171, 1097, 1040, 1016 cm$^{-1}$; **HRMS** (ESI): $m/z$ calculated for $C_{24}H_{42}O_6N_5$ [$M$+H]$^+$: 496.3130, found: 496.3127; $[\alpha]_D^{25}$ = –20.1 (c = 0.25, MeOH).

***tert*-Butyl ((*S*)-5-((*R*)-2-acetamido-4-methylpentanamido)-6-oxo-6-((2-oxo-2-(prop-2-yn-1-ylamino)ethyl)amino)hexyl)carbamate (I25)**

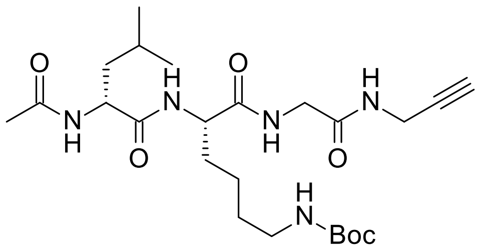


According to General Procedure F, *N*-acetylated peptide **I25** (128 mg, 38%) was obtained from *N*-Fmoc-protected peptide **I18** (456 mg, 0.7 mmol) following column chromatography (25 g Sfär; 80 mL/min; initially, 100% ethyl acetate (7 CV), followed by a linear gradient (15 CV): 0%$_{v/v}$ → 100%$_{v/v}$ acetone in ethyl acetate).

White amorphous solid; **$^1$H NMR** (600 MHz, DMSO-$d_6$): $\delta$ = 8.28 (d, $J$ = 7.7 Hz, 1H), 8.13 (t, $J$ = 6.0 Hz, 1H), 8.11 (t, $J$ = 5.6 Hz, 1H), 8.05 (d, $J$ = 7.3 Hz, 1H), 6.73 (t, $J$ = 5.4 Hz, 1H), 4.27 (q, $J$ = 7.5 Hz, 1H), 4.15-4.12 (m, 1H), 3.88 (ddd, $J$ = 17.6, 5.6, 2.5 Hz, 1H), 3.84 (ddd, $J$ = 17.5, 5.6, 2.6 Hz, 1H), 3.68 (dd, $J$ = 16.6, 5.9 Hz, 1H), 3.65 (dd, $J$ = 16.6, 6.0 Hz, 1H), 3.10 (t, $J$ = 2.5 Hz, 1H), 2.91-2.82 (m, 2H), 1.83 (s, 3H), 1.70-1.64 (m, 1H), 1.59-1.47 (m, 2H), 1.43-1.41 (m, 2H), 1.40- 1.29 (m, 11H), 1.28-1.16 (m, 2H), 0.90 (d, $J$ = 6.6 Hz, 3H), 0.85 ppm (d, $J$ = 6.5 Hz, 3H); **$^{13}$C NMR** (150 MHz, DMSO-$d_6$): $\delta$ = 172.7, 172.0, 169.6, 168.5, 155.5, 81.0, 77.3, 73.0, 52.7, 51.5, 41.8, 40.7, 39.5 (determined by HSQC), 31.0, 29.0, 28.3 (3C), 27.9, 24.2, 22.7, 22.6, 22.4, 21.9 ppm; **IR** (solid): ṽ = 3294, 3079, 2956, 2934, 2869, 1683, 1655, 1628, 1537, 1452, 1390, 1367, 1279, 1252, 1218, 1174, 1147, 1096, 1040, 1016 cm$^{-1}$; **HRMS** (ESI): $m/z$ calculated for $C_{24}H_{42}O_6N_5$ [$M$+H]$^+$: 496.3130, found: 496.3127; $[\alpha]_D^{25}$ = –1.8 (c = 0.25, MeOH).

***tert*-Butyl ((*S*)-5-((*S*)-2-acetamido-3-cyclohexylpropanamido)-6-oxo-6-((2-oxo-2-(prop-2-yn-1-ylamino)ethyl)amino)hexyl)carbamate (I26)**

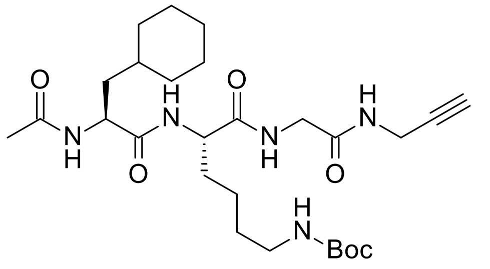

According to General Procedure F, *N*-acetylated peptide **I26** (224 mg, 61%) was obtained from *N*-Fmoc-protected peptide **I19** (491 mg, 0.7 mmol), following column chromatography (25 g Sfär; 80 mL/min; initially, 100% ethyl acetate (7 CV), followed by a linear gradient (15 CV): 0%$_{v/v}$ → 100%$_{v/v}$ acetone in ethyl acetate).

White amorphous solid; $^{1}$**H NMR** (600 MHz, DMSO-*d$_6$*): $\delta$ = 8.19 (t, *J* = 5.5 Hz, 1H), 8.08 (t, *J* = 5.8 Hz, 1H), 7.99 (d, *J* = 7.9 Hz, 1H), 7.92 (d, *J* = 7.6 Hz, 1H), 6.74 (t, *J* = 5.4 Hz, 1H), 4.28 (ddd, *J* = 9.7, 7.9, 5.2 Hz, 1H), 4.18-4.14 (m, 1H), 3.88 (ddd, *J* = 17.3, 5.4, 2.5 Hz, 1H), 3.84 (ddd, *J* = 17.3, 5.3, 2.5 Hz), 3.70 (dd, *J* = 16.7, 6.0 Hz, 1H), 3.64 (dd, *J* = 16.6, 5.6 Hz, 1H), 3.10 (t, *J* = 2.5 Hz, 1H), 2.90-2.83 (m, 2H), 1.84 (s, 3H), 1.69-1.58 (m, 6H), 1.54-1.44 (m, 2H), 1.42-1.30 (m, 12H), 1.29-1.15 (m, 4H), 1.14-1.07 (m, 2H), 0.92-0.79 ppm (m, 2H); $^{13}$**C NMR** (150 MHz, DMSO-*d$_6$*): $\delta$ = 172.4, 171.8, 169.4, 168.5, 155.5, 80.9, 77.3, 73.0, 52.6, 50.4, 41.8, 39.6 (determined by HSQC), 39.0 (determined by HSQC), 33.5, 33.2, 31.9, 31.3, 29.2, 28.3 (3C), 27.9, 26.1, 25.8, 25.6, 22.6, 22.5 ppm; **IR** (solid): ṽ = 3340, 3302, 3275, 3070, 2930, 2853, 1702, 1676, 1624, 1534, 1446, 1417, 1403, 1391, 1365, 1287, 1248, 1227, 1170, 1129, 1045, 1020, 1004 cm$^{-1}$; **HRMS** (ESI): *m/z* calculated for $C_{27}H_{46}O_6N_5$ [*M*+H]$^+$: 536.3443, found: 536.3435; $[\alpha]_D^{25}$ = –13.6 (c = 0.25, MeOH).

**(*S*)-5-((*S*)-2-Acetamido-4-methylpentanamido)-6-oxo-6-((2-oxo-2-(prop-2-yn-1-ylamino)ethyl)amino)hexan-1-aminium chloride (25)**

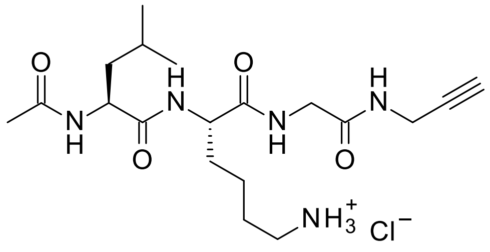

According to General Procedure H, tetrapeptide salt **25** (19 mg, 78%) was obtained from *N*-Boc-protected tetrapeptide **I24** (28 mg, 0.06 mmol).

Off-white amorphous solid; $^{1}$**H NMR** (600 MHz, DMSO-*d$_6$*): $\delta$ = 8.26 (t, *J* = 5.5 Hz, 1H), 8.11 (t, *J* = 5.8 Hz, 1H), 8.05 (d, *J* = 7.8 Hz, 1H), 8.02 (d, *J* = 7.8 Hz, 1H), 7.80 (br, 3H), 4.26 (q, *J* = 7.6 Hz, 1H), 4.22-4.19 (m, 1H), 3.88 (ddd, *J* = 17.5, 5.6, 2.5 Hz, 1H), 3.84 (ddd, *J* = 17.5, 5.6, 2.5 Hz, 1H), 3.70 (dd, *J* = 16.6, 6.0 Hz, 1H), 3.66 (dd, *J* = 16.6, 5.9 Hz, 1H), 3.11 (t, *J* = 2.5 Hz, 1H), 2.78-2.70 (m, 2H), 1.84 (s, 3H), 1.75 – 1.66 (m, 1H), 1.64 – 1.58 (m, 1H), 1.57 – 1.50 (m, 3H), 1.43 (t, *J* = 7.3 Hz, 2H), 1.37 – 1.23 (m, 2H), 0.88 (d, *J* = 6.6 Hz, 3H), 0.84 ppm (d, *J* = 6.5 Hz, 3H); $^{13}$**C NMR** (150 MHz, DMSO-*d$_6$*): $\delta$ = 172.5, 171.7, 169.5, 168.5, 81.0, 73.1, 52.3, 51.2, 41.8, 40.5, 38.6, 31.0, 27.9, 26.5, 24.2, 23.1, 22.5, 22.1, 21.5 ppm; **IR** (solid): ṽ = 3277, 3058, 2980, 2960, 2873, 1633,

1542, 1470, 1438, 1372, 1343, 1290, 1244, 1161, 1129, 1090, 1030 $cm^{-1}$; **HRMS** (ESI): *m/z* calculated for $C_{19}H_{34}O_4N_5$ $[M+H]^+$: 396.2605, found: 396.2603; $[\alpha]_D^{25}$ = –37.4 (c = 0.25, MeOH).

**(*S*)-5-((*S*)-2-Acetamido-3-cyclohexylpropanamido)-6-oxo-6-((2-oxo-2-(prop-2-yn-1-ylamino)ethyl)amino)hexan-1-aminium chloride (26)**

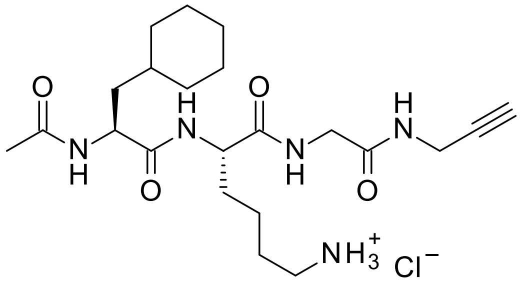

According to General Procedure H, tetrapeptide salt **26** (40 mg, apparent quant.) was obtained from *N*-Boc-protected tetrapeptide **I26** (40 mg, 0.08 mmol).

White amorphous solid; **$^1$H NMR** (600 MHz, DMSO-$d_6$): $\delta$ = 8.26 (t, *J* = 5.5 Hz, 1H), 8.09 (t, *J* = 5.8 Hz, 1H), 8.03 (d, *J* = 7.7 Hz, 1H), 8.00 (d, *J* = 7.8 Hz, 1H), 7.75 (br, 3H), 4.27 (ddd, *J* = 9.7, 7.7, 5.2 Hz, 1H), 4.23-4.19 (m, 1H), 3.88 (ddd, *J* = 17.5, 5.6, 2.5 Hz, 1H), 3.83 (ddd, *J* = 17.5, 5.6, 2.5 Hz, 1H), 3.70 (dd, *J* = 16.6, 5.8 Hz, 1H), 3.67 (dd, *J* = 16.6, 5.9 Hz, 1H), 3.11 (t, *J* = 2.5 Hz, 1H), 2.77 – 2.71 (m, 2H), 1.84 (s, 3H), 1.73 – 1.57 (m, 6H), 1.56 – 1.49 (m, 3H), 1.49 – 1.39 (m, 2H), 1.37 – 1.23 (m, 3H), 1.22 – 1.07 (m, 3H), 0.92 – 0.80 ppm (m, 2H); **$^{13}$C NMR** (150 MHz, DMSO-$d_6$): $\delta$ = 172.5, 171.7, 169.5, 168.5, 80.9, 73.1, 52.3, 50.5, 41.7, 39.0 (determined by HSQC), 38.6, 33.5, 33.2, 31.9, 31.0, 27.9, 26.5, 26.1, 25.8, 25.6, 22.5, 22.1 ppm; **IR** (solid): ṽ = 3279, 3055, 2926, 2855, 1641, 1538, 1468, 1441, 1372, 1345, 1289, 1256, 1244, 1228, 1160, 1092, 1039 $cm^{-1}$; **HRMS** (ESI): *m/z* calculated for $C_{22}H_{38}O_4N_5$ $[M+H]^+$: 436.2918, found: 436.2917; $[\alpha]_D^{25}$ = –24.1 (c = 0.25, MeOH).

**(*S*)-5-((*R*)-2-Acetamido-4-methylpentanamido)-6-oxo-6-((2-oxo-2-(prop-2-yn-1-ylamino)ethyl)amino)hexan-1-aminium chloride (27)**

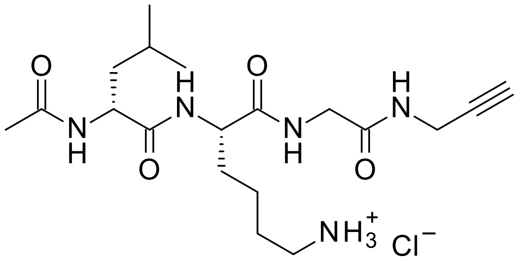

According to General Procedure H, tetrapeptide salt **27** (39 mg, apparent quant.) was obtained from *N*-Boc-protected tetrapeptide **I25** (40 mg, 0.08 mmol).

Pale yellow amorphous solid; **$^1$H NMR** (600 MHz, DMSO-$d_6$): $\delta$ = 8.29 (d, *J* = 7.9 Hz, 1H), 8.18 – 8.15 (m, 2H), 8.09 (d, *J* = 7.2 Hz, 1H), 7.75 (br, 3H), 4.25 (q, *J* = 7.5 Hz, 1H), 4.19-4.15 (m, 1H), 3.88 (ddd, *J* = 17.5, 5.6, 2.6 Hz, 1H) 3.84 (ddd, *J* = 17.5, 5.6, 2.6 Hz, 1H), 3.71 – 3.64 (m, 2H), 3.11 (t, *J* = 2.5 Hz, 1H), 2.78 – 2.69 (m, 2H), 1.84 (s, 3H), 1.74 – 1.68 (m, 1H), 1.60 – 1.48 (m, 4H), 1.43 (t, *J* = 7.3 Hz, 2H), 1.36 – 1.23 (m, 2H), 0.89 (d, *J* = 6.6 Hz, 3H), 0.85 ppm (d, *J* = 6.6 Hz, 3H); **$^{13}$C NMR** (150 MHz, DMSO-$d_6$): $\delta$ = 172.6, 171.8, 169.7, 168.4, 81.0, 73.1, 52.4, 51.6, 41.8, 40.6, 38.6, 30.7, 27.9, 26.4, 24.2, 22.8, 22.4, 22.1, 21.9 ppm; **IR** (solid): ṽ = 3278, 3054, 2980,

2962, 2934, 1651, 1633, 1542, 1471, 1457, 1437, 1374, 1340, 1283, 1251, 1224, 1160, 1125, 1089, 1030 cm$^{-1}$; **HRMS** (ESI): *m/z* calculated for $C_{19}H_{34}O_4N_5$ [*M*+H]$^+$: 396.2605, found: 396.2607; $[\alpha]_D^{25}$ = –7.3 (c = 0.25, MeOH).

**(*S*)-5-((*S*)-2-Acetamido-4-methylpentanamido)-6-((2-((cyanomethyl)amino)-2-oxoethyl)amino)-6-oxohexan-1-aminium chloride (28)**

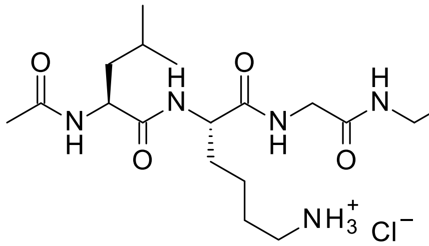


According to General Procedure G, tetrapeptide salt **28** (22 mg, apparent quant.) was obtained from N-Boc-protected tetrapeptide **I20** (25 mg, 0.05 mmol).

White amorphous solid; **$^1$H NMR** (600 MHz, DMSO-*d$_6$*): $\delta$ = 8.58 (t, *J* = 5.6 Hz, 1H), 8.21 (t, *J* = 5.9 Hz, 1H), 8.06 (d, *J* = 7.6 Hz, 1H), 8.05 (d, *J* = 7.8 Hz, 1H), 7.81 (br, 3H), 4.27 (q, *J* = 7.6 Hz, 1H), 4.23-4.19 (m, 1H), 4.14 (d, *J* = 5.6 Hz, 2H), 3.76 (dd, *J* = 16.8, 6.0 Hz, 1H), 3.72 (dd, *J* = 16.9, 5.8 Hz, 1H), 2.78 – 2.71 (m, 2H), 1.85 (s, 3H), 1.74 – 1.67 (m, 1H), 1.64 – 1.50 (m, 4H), 1.46-1.41 (m, 2H), 1.38 – 1.25 (m, 2H), 0.88 (d, *J* = 6.6 Hz, 3H), 0.84 ppm (d, *J* = 6.6 Hz, 3H); **$^{13}$C NMR** (150 MHz, DMSO-*d$_6$*): $\delta$ = 172.5, 171.9, 169.6, 169.5, 117.5, 52.3, 51.1, 41.7, 40.5, 38.6, 30.9, 27.0, 26.5, 24.2, 23.1, 22.5, 22.1, 21.5 ppm; **IR** (solid): ṽ = 3277, 3052, 2980, 2959, 2931, 2872, 1666, 1635, 1537, 1471, 1445, 1415, 1390, 1373, 1339, 1283, 1236, 1162, 1099, 1028 cm$^{-1}$; **HR-MS** (ESI): *m/z* calculated for $C_{18}H_{33}O_4N_6$ [*M*+H]$^+$: 397.2558, found: 397.2554; $[\alpha]_D^{25}$ = -36.8 (c = 0.25, MeOH).

**(*S*)-5-((*S*)-2-Acetamido-2-cyclohexylacetamido)-6-((2-((cyanomethyl)amino)-2-oxoethyl)amino)-6-oxohexan-1-aminium chloride (29)**

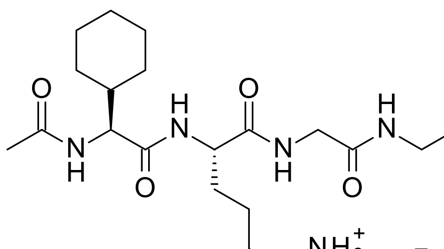


According to General Procedure G, tetrapeptide salt **29** (12 mg, 66%) was obtained from *N*-Boc-protected tetrapeptide **I23** (20 mg, 0.038 mmol).

Orange amorphous solid; **$^1$H NMR** (600 MHz, DMSO-*d$_6$*): $\delta$ = 8.59 (t, *J* = 5.6 Hz, 1H), 8.22 (t, *J* = 5.9 Hz, 1H), 8.11 (d, *J* = 7.3 Hz, 1H), 7.92 (d, *J* = 8.3 Hz, 1H), 7.80 (br, 3H), 4.21-4.17 (m, 1H), 4.14-4.12 (m, 3H), 3.76 (dd, *J* = 16.8, 6.1 Hz, 1H), 3.71 (dd, *J* = 16.8, 5.7 Hz, 1H), 2.75-2.72 (app.t, 2H), 1.86 (s, 3H), 1.71-1.51 (m, 10H), 1.36-1.27 (m, 2H), 1.15-1.06 (m, 3H), 1.02-0.94 ppm (m, 2H); **$^{13}$C NMR** (150 MHz, DMSO-*d$_6$*): $\delta$ = 171.9, 171.4, 169.6, 169.5, 117.5, 57.4, 52.6, 41.7, 39.6 (determined by HSQC), 38.6, 30.8, 29.0, 28.2, 27.0, 26.6, 25.8, 25.5(8), 25.5(6), 22.5, 22.1 ppm; **IR** (solid): ṽ = 3279, 3057, 2927, 2854, 1632,

1546, 1468, 1443, 1403, 1373, 1344, 1291, 1256, 1242, 1161, 1130, 1101, 1032 cm$^{-1}$; **HRMS** (ESI): *m/z* calculated for $C_{20}H_{35}O_4N_6$ [*M*+H]$^+$: 423.2714, found: 423.2709; $[\alpha]_D^{25}$ = –32.0 (c = 0.125, MeOH).

**(*S*)-5-((*R*)-2-Acetamido-4-methylpentanamido)-6-((2-((cyanomethyl)amino)-2-oxoethyl)amino)-6-oxohexan-1-aminium chloride (30)**

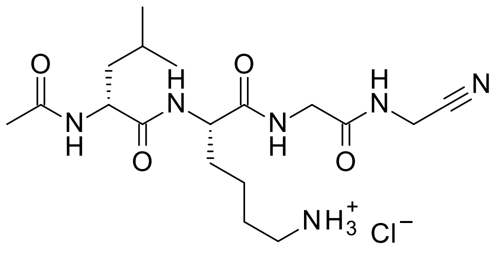


According to General Procedure G, tetrapeptide salt **30** (22 mg, apparent quant.) was obtained from *N*-Boc-protected tetrapeptide **I21** (25 mg, 0.05 mmol).

Pale yellow amorphous solid; **$^1$H NMR** (600 MHz, DMSO-*d$_6$*): *δ* = 8.52 (t, *J* = 5.6 Hz, 1H), 8.37 (d, *J* = 7.8 Hz, 1H), 8.27 (t, *J* = 6.0 Hz, 1H), 8.11 (d, *J* = 7.2 Hz, 1H), 7.86 (br, 3H), 4.26 (q, *J* = 7.4 Hz, 1H), 4.20-4.16 (m, 1H), 4.14-4.13 (m, 2H), 3.76 (dd, J = 16.7, 5.9 Hz, 1H), 3.70 (dd, *J* = 16.7, 5.9 Hz, 1H), 2.77– 2.69 (m, 2H), 1.84 (s, 3H), 1.75 – 1.69 (m, 1H), 1.60 – 1.49 (m, 3H), 1.44 (t, *J* = 7.3 Hz, 2H), 1.36 – 1.23 (m, 2H), 0.90 (d, *J* = 6.6 Hz, 3H), 0.85 ppm (d, *J* = 6.6 Hz, 3H); **$^{13}$C NMR** (150 MHz, DMSO-*d$_6$*): *δ* = 172.7, 171.9, 169.7, 169.5, 117.5, 52.4, 51.6, 41.7, 40.6, 38.5, 30.6, 27.0, 26.4, 24.2, 22.8, 22.4, 22.1, 21.9 ppm; **IR** (solid): ṽ = 3271, 3051, 2955, 2930, 2872, 1657, 1650, 1546, 1535, 1468, 1443, 1412, 1372, 1344, 1284, 1244, 1164, 1100, 1029 cm$^{-1}$; **HRMS** (ESI): *m/z* calculated for $C_{18}H_{33}O_4N_6$ [*M*+H]$^+$: 397.2558, found: 397.2555; $[\alpha]_D^{25}$ = –8.1 (c = 0.25, MeOH).

**(*S*)-5-((*S*)-2-Acetamido-3-cyclohexylpropanamido)-6-((2-((cyanomethyl)amino)-2-oxoethyl)amino)-6-oxohexan-1-aminium chloride (31)**

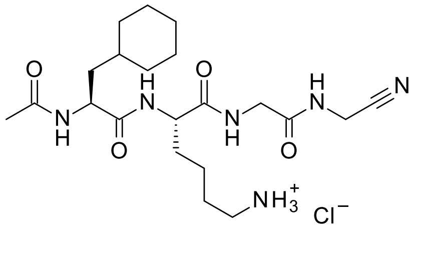


According to General Procedure G, tetrapeptide salt **31** (20 mg, 88%) was obtained from *N*-Boc-protected tetrapeptide **I22** (25 mg, 0.05 mmol).

Off-white amorphous solid; **$^1$H NMR** (600 MHz, DMSO-*d$_6$*): *δ* = 8.58 (t, *J* = 5.6 Hz, 1H), 8.21 (t, *J* = 5.9 Hz, 1H), 8.06 – 8.04 (m, 2H), 7.83 (br, 3H), 4.27 (ddd, *J* = 9.8, 7.8, 5.1 Hz, 1H), 4.23-4.20 (m, 1H), 4.14 (d, *J* = 5.7 Hz, 2H), 3.76 (dd, *J* = 16.9, 6.1 Hz, 1H), 3.72 (dd, *J* = 17.0, 5.9 Hz, 1H), 2.76 – 2.71 (m, 2H), 1.85 (s, 3H), 1.74 – 1.50 (m, 9H), 1.50 – 1.39 (m, 2H), 1.37 – 1.23 (m, 3H), 1.21 – 1.07 (m, 3H), 0.92 – 0.80 ppm (m, 2H); **$^{13}$C NMR** (150 MHz, DMSO-*d$_6$*): *δ* = 172.6, 171.9, 169.5(4), 169.5(2), 117.5, 52.3, 50.5, 41.7, 39.0 (determined by HSQC), 38.6, 33.5, 33.2, 31.8, 30.9, 27.0, 26.5, 26.1, 25.8, 25.6, 22.5, 22.1 ppm; **IR** (solid): ṽ = 3274, 3060, 2981, 2924, 2852, 1666, 1635, 1540, 1474, 1448, 1417, 1393, 1375, 1354, 1251, 1232, 1179, 1152, 1119, 1074, 1023 cm$^{-1}$;

**HRMS** (ESI): *m/z* calculated for $C_{21}H_{37}O_4N_6$ [*M*+H]$^+$: 437.2871, found: 437.2871; $[\alpha]_D^{25}$ = –24.1 (c = 0.25, MeOH).

**2,5-Dioxopyrrolidin-1-yl-(((9*H*-fluoren-9-yl)methoxy)carbonyl)-*L*-leucinate (I27)**

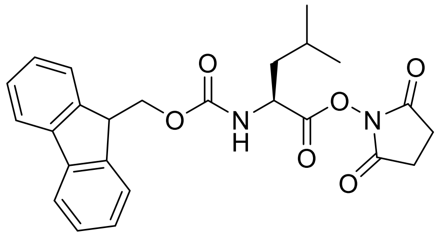

According to General Procedure A, ester **I27** (1.68 g, 65%) was obtained from commercially sourced (((9*H*-fluoren-9-yl)methoxy)carbonyl)-*L*-leucine (2.00 g, 5.7 mmol).

Characteristic analytical data: **$^1$H NMR** (400 MHz, DMSO-*d$_6$*): δ = 8.11 (d, *J* = 8.0 Hz, 1H), 7.89 (d, *J* = 7.6 Hz, 2H), 7.70 (dd, *J* = 7.5 Hz, 2H), 7.42 (dd, *J* = 7.9, 6.8 Hz, 2H), 7.32 (tt, *J* = 7.4 Hz, 2H), 4.42 (ddd, *J* = 10.3, 8.0, 4.5 Hz, 1H), 4.36 (d, *J* = 7.0 Hz, 2H), 4.24 (t, *J* = 7.0 Hz, 1H), 2.80 (s, 4H), 1.76 (td, *J* = 10.4, 4.4 Hz, 2H), 1.62 (td, *J* = 10.4, 6.0 Hz, 1H), 0.90 ppm (dd, *J* = 6.2 Hz, 6H); **$^{13}$C NMR** (101 MHz, DMSO-*d$_6$*): δ = 170.0, 169.1, 156.0, 143.8, 143.6, 140.7, 127.7, 127.1, 125.2, 120.1, 65.8, 50.5, 46.6, 38.9, 25.5, 24.1, 22.7, 21.0 ppm; **HRMS** (ESI): *m/z* calculated for $C_{25}H_{27}N_2O_6$ [*M*+H]$^+$: 451.1864, found: 451.1874.

***N*²-((((9*H*-Fluoren-9-yl)methoxy)carbonyl)-*L*-leucyl)-*N*⁶-(*tert*-butoxycarbonyl)-*L*-lysine (I28)**

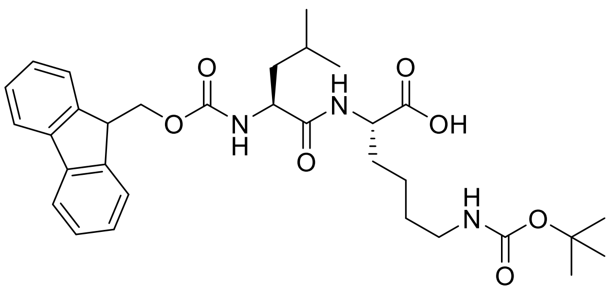

According to General Procedure B, dipeptide **I28** (3.77 g, 97%) was obtained from *N*-hydroxysuccinimide ester **I27** (3.00 g, 6.7 mmol) and commercially sourced L-Lys(Boc)-OH (2.47 g, 10.0 mmol).

Characteristic analytical data: **$^1$H NMR** (400 MHz, DMSO-*d$_6$*): δ = δ 12.48 (s, 1H), 8.02 (d, *J* = 7.6 Hz, 1H), 7.88 (d, *J* = 7.5 Hz, 2H), 7.72 (t, *J* = 6.7 Hz, 2H), 7.47 (d, *J* = 8.5 Hz, 1H), 7.41 (td, *J* = 7.5, 1.2 Hz, 2H), 7.31 (tdd, *J* = 7.4, 1.2 Hz, 2H), 6.72 (t, *J* = 5.4 Hz, 1H), 4.34 – 4.17 (m, 3H), 4.09 (m, 2H), 2.87 (q, *J* = 6.5 Hz, 2H), 1.71 – 1.49 (m, 3H), 1.44 (m, 2H), 1.34 (s, 9H), 1.33 – 1.27 (m, 4H), 0.87 ppm (dd, *J* = 6.5 Hz, 6H); **$^{13}$C NMR** (101 MHz, DMSO-*d$_6$*): δ = 173.5, 172.4, 155.9, 155.5, 144.0, 143.7, 140.7, 127.6, 127.0, 125.3, 120.1, 77.3, 65.5, 52.8, 51.8, 46.7, 40.8, 39.2, 30.7, 29.1, 28.3, 24.1, 23.1, 22.8, 21.5 ppm; **HRMS** (ESI): *m/z* calculated for $C_{32}H_{44}N_3O_7$ [*M*+H]$^+$: 582.3174, found: 582.3164.

***tert*-Butyl-(2-oxo-2-(prop-2-yn-1-ylamino)ethyl)carbamate (I29)**

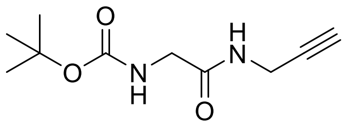

According to General Procedure C, alkyne **I29** (0.94 g, 80%) was obtained from commercially sourced *N*-Boc-glycine (1.1 g, 6.1 mmol)

and commercially sourced propargylamine hydrochloride (0.50 g, 5.5 mmol). The characteristic analytical data of **I29** are consistent with those reported[15–18].

Characteristic analytical data: **$^1$H NMR** (400 MHz, DMSO-*d$_6$*): δ = 8.21 (t, *J* = 5.6 Hz, 1H), 6.94 (t, *J* = 6.2 Hz, 1H), 3.85 (dd, *J* = 5.6, 2.5 Hz, 2H), 3.51 (d, *J* = 5.9 Hz, 2H), 3.09 (t, *J* = 2.5 Hz, 1H), 1.38 ppm (s, 9H); **$^{13}$C NMR** (101 MHz, DMSO-*d$_6$*): δ = 169.2, 155.8, 81.1, 78.0, 72.9, 43.0, 28.2, 27.8 ppm; **HRMS** (ESI): *m/z* calculated for $C_{10}H_{16}N_2O_3Na$ [*M*+Na]$^+$: 235.1053, found: 235.1049.

***tert*-Butyl-(2-(but-2-yn-1-ylamino)-2-oxoethyl)carbamate (I30)**

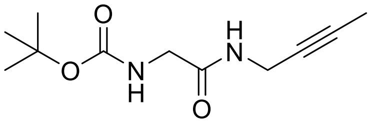


According to General Procedure C, alkyne **I30** (231 mg, 54%) was obtained from commercially sourced *N*-Boc-glycine (365 mg, 2.09 mmol) and commercially sourced but-2-yn-1-amine hydrochloride (200 mg, 1.90 mmol).

Characteristic analytical data: **$^1$H NMR** (400 MHz, DMSO-*d$_6$*): δ = 8.13 (t, *J* = 5.5 Hz, 1H), 6.92 (t, *J* = 6.2 Hz, 1H), 3.81 (dq, *J* = 4.9, 2.5 Hz, 2H), 3.50 (d, *J* = 6.2 Hz, 2H), 1.76 (t, *J* = 2.5 Hz, 3H), 1.38 ppm (s, 9H); **$^{13}$C NMR** (151 MHz, DMSO-*d$_6$*): δ = 169.0, 155.8, 78.0, 77.9, 76.4, 43.0, 28.2, 28.1, 3.0 ppm; **HRMS** (ESI): *m/z* calculated for $C_{11}H_{18}N_2O_3Na$ [*M*+Na]$^+$: 249.1210, found: 249.1209.

***tert*-Butyl-(2-((1-ethynylcyclopropyl)amino)-2-oxoethyl)carbamate (I31)**

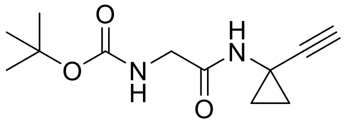


According to General Procedure C, alkyne **I31** (247 mg, 58%) was obtained from commercially sourced *N*-Boc-glycine (345 mg, 1.97 mmol) and commercially sourced 1-ethynylcyclopropanamine hydrochloride (209 mg, 1.78 mmol).

Characteristic analytical data: **$^1$H NMR** (400 MHz, DMSO-*d$_6$*): δ = 8.42 (s, 1H), 6.84 (t, *J* = 6.1 Hz, 1H), 3.44 (d, *J* = 5.6 Hz, 2H), 2.93 (s, 1H), 1.37 (s, 9H), 1.06 (q, *J* = 4.7 Hz, 2H), 0.91 ppm (q, *J* = 4.6 Hz, 2H); **$^{13}$C NMR** (151 MHz, DMSO-*d$_6$*): δ = 169.6, 155.8, 86.4, 78.0, 68.5, 42.9, 28.2, 21.8, 16.8 ppm; **HRMS** (ESI): *m/z* calculated for $C_{12}H_{19}N_2O_3$ [*M*+H]$^+$: 239.1390, found: 239.1397.

***tert*-Butyl-(2-oxo-2-((3-phenylprop-2-yn-1-yl)amino)ethyl)carbamate (I32)**

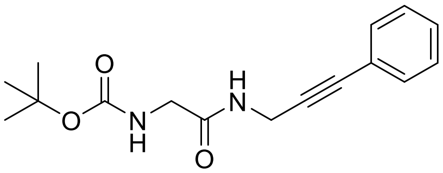


According to General Procedure C, alkyne **I32** (118 mg, 85%) was obtained from commercially sourced *N*-Boc-glycine (94 mg, 0.53 mmol) and commercially sourced 3-phenyl-2-propyn-1-

amine hydrochloride (80 mg, 0.48 mmol). The characteristic analytical data of **I32** are consistent with those reported[19].

Characteristic analytical data: **$^1$H NMR** (400 MHz, DMSO-$d_6$): δ = 8.32 (t, *J* = 5.5 Hz, 1H), 7.45 – 7.33 (m, 5H), 6.97 (t, *J* = 6.2 Hz, 1H), 4.13 (d, *J* = 5.5 Hz, 2H), 3.55 (d, *J* = 6.2 Hz, 2H), 1.38 ppm (s, 9H); **$^{13}$C NMR** (151 MHz, DMSO-$d_6$): δ = 169.3, 155.8, 131.4, 128.7, 128.6, 122.3, 87.1, 81.6, 78.1, 43.1, 28.6, 28.2 ppm; **HRMS** (ESI): *m/z* calculated for $C_{16}H_{20}N_2O_3Na$ [*M*+Na]$^+$: 311.1366, found: 311.1372.

**(9*H*-Fluoren-9-yl)methyl-((*S*)-1-(((*S*)-2,2-dimethyl-4,11,14-trioxo-3-oxa-5,12,15-triazaoctadec-17-yn-10-yl)amino)-4-methyl-1-oxopentan-2-yl)carbamate (I38)**

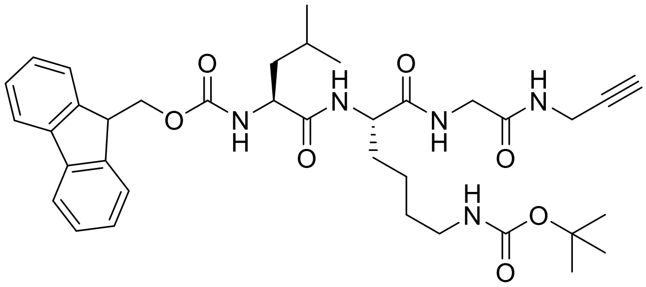


According to General Procedure D, amine hydrochloride **I33** was obtained *in situ* from *N*-Boc-protected peptide **I29** (0.10 mg, 0.48 mmol) and was directly used in the following reaction: According to General Procedure E, tetrapeptide **I38** (0.33 g, apparent quantitative) was obtained from dipeptide **I28** (0.27 g, 0.46 mmol) and amine hydrochloride **I33**.

Characteristic analytical data: **$^1$H NMR** (400 MHz, DMSO-$d_6$): δ = 8.24 (t, *J* = 5.6 Hz, 1H), 8.18 (t, *J* = 5.9 Hz, 1H), 7.96 (d, *J* = 7.5 Hz, 1H), 7.92 – 7.85 (m, 2H), 7.72 (t, *J* = 6.9 Hz, 2H), 7.52 (d, *J* = 8.4 Hz, 1H), 7.41 (td, *J* = 7.5, 1.1 Hz, 2H), 7.32 (tt, *J* = 7.4, 1.4 Hz, 2H), 6.71 (t, *J* = 5.7 Hz, 1H), 4.37 – 4.15 (m, 4H), 4.11 – 3.98 (m, 1H), 3.86 (dt, *J* = 6.0, 2.2 Hz, 2H), 3.77 – 3.60 (dd, *J* = 6.3 Hz, 2H), 3.09 (t, *J* = 2.5 Hz, 1H), 2.86 (q, *J* = 6.6 Hz, 2H), 1.67 – 1.55 (m, 2H), 1.55 – 1.37 (m, 3H), 1.41 – 1.13 (m, 4H), 1.34 (s, 9H), 0.87 ppm (dd, *J* = 13.1, 6.5 Hz, 6H); **$^{13}$C NMR** (101 MHz, DMSO-$d_6$): δ = 172.5, 171.9, 168.5, 155.9, 155.5, 144.0, 143.7, 140.7, 127.6, 127.1, 125.3, 120.1, 81.0, 77.3, 73.0, 65.5, 53.0, 52.7, 46.7, 41.8, 40.6, 39.5, 31.6, 29.2, 28.3, 27.9, 24.2, 23.1, 22.5, 21.4 ppm; **HRMS** (ESI): *m/z* calculated for $C_{37}H_{50}N_5O_7$ [*M*+H]$^+$: 676.3705, found: 676.3696.

**(9*H*-Fluoren-9-yl)methyl-((*S*)-1-(((*S*)-2,2-dimethyl-4,11,14-trioxo-3-oxa-5,12,15-triazanonadec-17-yn-10-yl)amino)-4-methyl-1-oxopentan-2-yl)carbamate (I39)**

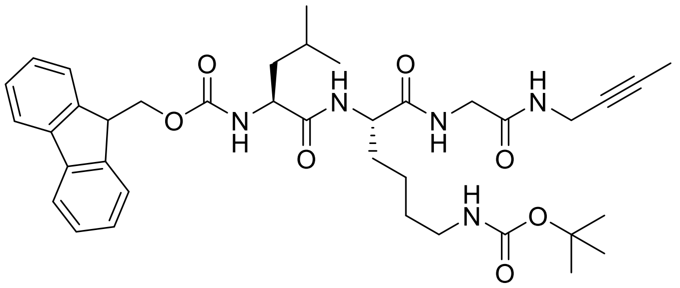

According to General Procedure D, amine hydrochloride **I34** was obtained *in situ* from *N*-Boc-protected peptide **I30** (0.23 g, 1.02 mmol) and was directly used in the following reaction: According to General Procedure E, tetrapeptide **I39** (0.76 g, apparent quantitative) was obtained from dipeptide **I28** (0.60 g, 1.03 mmol) and amine hydrochloride **I34**.

Characteristic analytical data: **$^1$H NMR** (400 MHz, DMSO-*d$_6$*): δ = 8.14 (s, 2H), 7.93 (d, *J* = 7.4 Hz, 1H), 7.89 (d, *J* = 7.6 Hz, 2H), 7.71 (t, *J* = 6.9 Hz, 2H), 7.50 (d, *J* = 8.5 Hz, 1H), 7.41 (t, *J* = 7.4 Hz, 2H), 7.32 (t, *J* = 7.5 Hz, 2H), 6.71 (t, *J* = 6.9 Hz, 1H), 4.36 – 4.19 (m, 4H), 4.10 – 4.00 (m, 1H), 3.81 (dt, *J* = 5.3, 2.6 Hz, 2H), 3.75 – 3.58 (m, 2H), 2.86 (d, *J* = 7.2 Hz, 2H), 1.75 (q, *J* = 2.4 Hz, 3H), 1.65 – 1.58 (m, 2H), 1.51 – 1.35 (m, 3H), 1.34 (s, 9H), 1.32 – 1.17 (m, 4H), 0.86 ppm (dd, *J* = 13.2, 6.5 Hz, 6H); **$^{13}$C NMR** (151 MHz, DMSO-*d$_6$*): δ = 172.4, 171.9, 168.4, 155.9, 155.6, 142.6, 139.4, 137.4, 128.9, 127.3, 121.4, 120.0, 109.8, 78.0, 77.3, 76.2, 65.5, 52.6, 52.2, 46.7, 43.7, 41.8, 40.5, 39.2, 31.8, 29.2, 28.7, 28.3, 24.0, 23.3, 23.1, 22.5, 21.7, 3.0 ppm; **HRMS** (ESI): *m/z* calculated for $C_{38}H_{52}N_5O_7$ [*M*+H]$^+$: 690.3861, found: 690.3833.

**(9*H*-Fluoren-9-yl)methyl-((*S*)-1-(((*S*)-6-((*tert*-butoxycarbonyl)amino)-1-((2-((1-ethynylcyclopropyl)amino)-2-oxoethyl)amino)-1-oxohexan-2-yl)amino)-4-methyl-1-oxopentan-2-yl)carbamate (I40)**

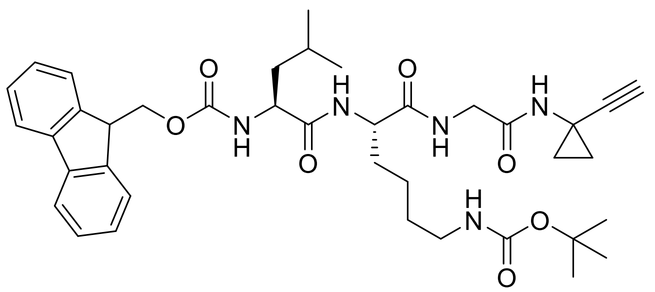

A solution of *N*-Boc-protected peptide **I31** (1.0 equiv., 0.25 g, 1.04 mmol) in commercially sourced formic acid (0.12 M) was stirred under an atmosphere of $N_2$ gas overnight (16 h). The reaction was concentrated under reduced pressure (30 °C) and the crude residue was redissolved in tetrahydrofuran/water (3:1 v/v, 0.13 M). The solution was basified with AmberLite™ HPR4800 OH (23-27 mesh) to pH 8 and filtered to afford amine hydrochloride **I35** which was directly used in the following reaction: According to General Procedure E, tetrapeptide **I40** (0.65 g, 90%) was obtained from dipeptide **I28** (0.61 g, 1.04 mmol) and amine hydrochloride **I35**.

Characteristic analytical data: **$^1$H NMR** (400 MHz, DMSO-$d_6$): δ = 8.35 (s, 1H), 8.11 (t, *J* = 5.8 Hz, 1H), 7.96 (d, *J* = 7.5 Hz, 1H), 7.89 (d, *J* = 7.6 Hz, 2H), 7.77 – 7.68 (m, 2H), 7.52 (d, *J* = 8.3 Hz, 1H), 7.41 (t, *J* = 7.4 Hz, 2H), 7.32 (t, *J* = 7.4 Hz, 2H), 6.71 (d, *J* = 5.8 Hz, 1H), 4.36 – 4.14 (m, 4H), 4.10 – 3.98 (m, 1H), 3.75 – 3.46 (dd, *J* = 5.5 Hz, 2H), 2.93 (s, 1H), 2.86 (q, *J* = 6.9 Hz, 2H), 1.72 – 1.56 (m, 2H), 1.55 – 1.40 (m, 3H), 1.38 – 1.12 (m, 4H), 1.34 (s, 9H), 1.06 (dq, *J* = 5.2, 2.7 Hz, 2H), 0.96 – 0.77 ppm (m, 8H); **$^{13}$C NMR** (151 MHz, DMSO-$d_6$): δ = 171.9, 168.8, 167.5, 155.6, 147.1, 143.9, 140.6, 140.5, 127.8, 127.3, 124.0, 120.7, 77.4, 73.2, 68.6, 65.1, 52.9, 52.7, 46.2, 41.8, 40.9, 39.5, 31.8, 28.9, 28.3, 24.0, 23.3, 22.4, 21.8, 21.7(5), 16.8 ppm; **HRMS** (ESI): *m/z* calculated for $C_{39}H_{52}N_5O_7$ [*M*+H]$^+$: 702.3861, found: 702.3864.

**(9*H*-Fluoren-9-yl)methyl-((*S*)-1-(((*S*)-6-((*tert*-butoxycarbonyl)amino)-1-((2-((1-(1-chlorovinyl)cyclopropyl)amino)-2-oxoethyl)amino)-1-oxohexan-2-yl)amino)-4-methyl-1-oxopentan-2-yl)carbamate (I41)**

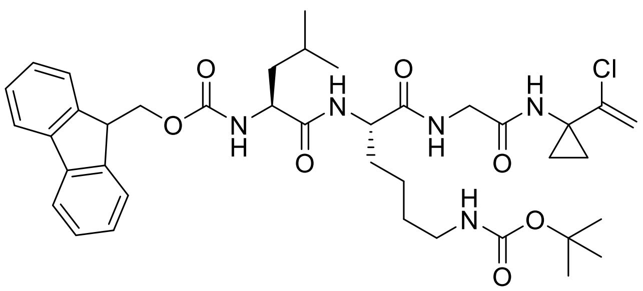

According to General Procedure D, amine hydrochloride **I36** was obtained *in situ* from *N*-Boc-protected peptide **I31** (82.9 mg, 0.348 mmol) and was directly used in the following reaction: According to General Procedure E, tetrapeptide **I41** (244 mg, apparent quantitative, 1:1 mixture of **I41** and **I40** as per $^1$H NMR analysis) was obtained from dipeptide **3** (205 mg, 0.35 mmol) and amine hydrochloride **I36** (mixture of **I36** and **I35**).

Characteristic analytical data for **I41**: **$^1$H NMR** (400 MHz, DMSO-$d_6$): δ = 8.35 (s, 1H), 8.17 (t, *J* = 5.7 Hz, 1H), 7.98 (d, *J* = 7.5 Hz, 1H), 7.89 (d, *J* = 7.6 Hz, 2H), 7.72 (t, *J* = 6.7 Hz, 2H), 7.50 (d, *J* = 7.2 Hz, 1H), 7.41 (t, *J* = 7.5 Hz, 2H), 7.36 – 7.28 (m, 2H), 6.71 (s, 1H), 5.43 (d, *J* = 1.8 Hz, 1H), 5.22 (d, *J* = 1.8 Hz, 1H), 4.36 – 4.13 (m, 4H), 4.11 – 4.00 (m, 1H), 3.70 – 3.55 (m, 2H), 2.86 (d, *J* = 7.7 Hz, 2H), 1.70 – 1.56 (m, 2H), 1.55 – 1.40 (m, 3H), 1.40 – 1.20 (m, 4H), 1.34 (s, 9H), 1.07 (d, *J* = 6.6 Hz, 2H), 0.92 (d, *J* = 6.6 Hz, 2H), 0.89 – 0.81 ppm (m, 6H); **$^{13}$C NMR** (151 MHz, DMSO-$d_6$): δ = 170.8, 170.4, 155.9, 155.5, 144.0, 143.7, 140.7, 139.4, 128.9, 127.3, 125.3, 120.1, 112.0, 77.3, 65.5, 53.0, 52.2, 46.7, 41.9, 40.6, 39.2, 36.0, 31.7, 29.2, 28.3, 24.1, 23.1, 22.6, 21.4, 16.8 ppm; **HRMS** (ESI): *m/z* calculated for $C_{39}H_{53}ClN_5O_7$ (**I41**) [*M*+H]$^+$: 738.3628, found: 738.3646; *m/z* calculated for $C_{39}H_{53}N_5O_7$ (**I40**) [*M*+H]$^+$: 702.3861, found: 702.3887.

**(9*H*-Fluoren-9-yl)methyl-((*S*)-1-(((*S*)-2,2-dimethyl-4,11,14-trioxo-18-phenyl-3-oxa-5,12,15-triazaoctadec-17-yn-10-yl)amino)-4-methyl-1-oxopentan-2-yl)carbamate (I42)**

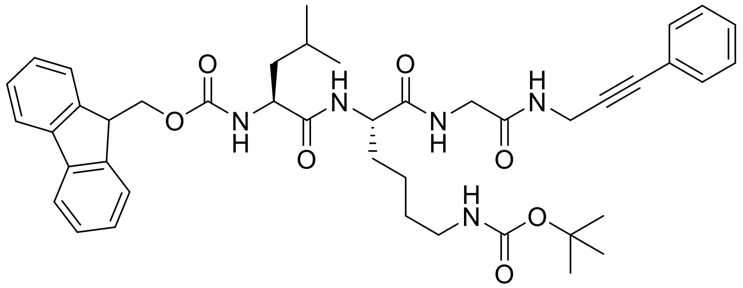

According to General Procedure D, alkyne hydrochloride salt **I37** was obtained *in situ* from *N*-Boc-protected peptide **I32** (0.25 g, 0.86 mmol) and was directly used in the following reaction: According to General Procedure E, tetrapeptide **I42** (0.61 g, 89%) was obtained from dipeptide **I28** (0.50 g, 0.86 mmol) and amine hydrochloride **I37**.

Characteristic analytical data: **$^1$H NMR** (400 MHz, DMSO-*d$_6$*): δ = 8.34 (t, *J* = 5.5 Hz, 1H), 8.23 – 8.12 (m, 1H), 7.94 (d, *J* = 7.7 Hz, 1H), 7.88 (d, *J* = 7.5 Hz, 2H), 7.71 (t, *J* = 6.8 Hz, 2H), 7.50 (d, *J* = 8.7 Hz, 1H), 7.45 – 7.27 (m, 9H), 6.71 (s, 1H), 4.34 – 4.19 (m, 4H), 4.13 (dd, *J* = 5.5, 1.9 Hz, 2H), 4.09 – 3.98 (m, 1H), 3.72 (t, *J* = 6.8 Hz, 2H), 2.86 (d, *J* = 7.6 Hz, 2H), 1.64 – 1.56 (m, 2H), 1.52 – 1.39 (m, 3H), 1.39 – 1.20 (m, 4H), 1.34 (s, 9H), 0.91 – 0.79 ppm (m, 6H); **$^{13}$C NMR** (151 MHz, DMSO-*d$_6$*): δ = 172.5, 171.9, 168.6, 156.0, 155.6, 144.0, 143.7, 140.7, 131.4, 128.7, 128.6, 127.7, 127.1, 125.3, 122.3, 120.1, 86.9, 81.6, 77.4, 65.5, 53.1, 52.7, 46.7, 41.9, 40.6, 39.2, 31.6, 29.2, 28.7, 28.3, 24.2, 23.1, 22.5, 21.4 ppm; **HRMS** (ESI): *m/z* calculated for $C_{43}H_{54}N_5O_7$ [*M*+H]$^+$: 752.4018, found: 752.3991.

***tert*-Butyl-((*S*)-5-((*S*)-2-acetamido-4-methylpentanamido)-6-oxo-6-((2-oxo-2-(prop-2-yn-1-ylamino)ethyl)amino)hexyl)carbamate (25d)**

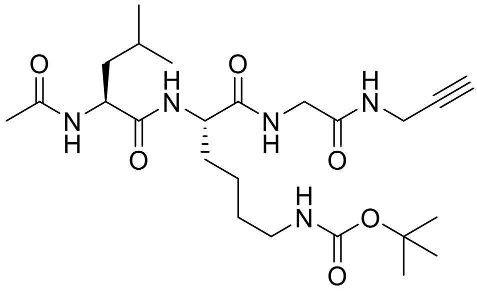

According to General Procedure F, *N*-acetyl tetrapeptide **25d** (20.3 mg, 8%) was obtained from *N*-Fmoc tetrapeptide **I38** (360 mg, 0.53 mmol) and commercially sourced acetic anhydride (53 µL, 0.56 mmol), following flash column chromatography (Biotage® Sfär Silica D Duo 5 g; 18 mL/min; 100%$_{v/v}$ cyclohexane (2 CV), linear gradient (30 CV): 0%$_{v/v}$ → 100%$_{v/v}$ acetone in cyclohexane, 100%$_{v/v}$ acetone (2 CV)), and preparative high-performance liquid chromatography (Avantor® ACE® 10 AQ 250 × 21.2 mm; 20 mL/min, linear gradient (30 min): 2%$_{v/v}$ → 98%$_{v/v}$ acetonitrile in water (each containing 0.1%$_{v/v}$ formic acid); $t_R$ = 15.4 min).

White amorphous solid ; **$^1$H NMR** (600 MHz, DMSO-*d$_6$*): δ = 8.19 (t, *J* = 5.6 Hz, 1H), 8.10 (t, *J* = 5.9 Hz, 1H), 7.99 (d, *J* = 8.0 Hz, 1H), 7.94 (d, *J* = 7.5 Hz, 1H), 6.74 (t, *J* = 5.7 Hz, 1H), 4.27 (td, *J* = 8.3, 6.1 Hz, 1H), 4.15 (td, *J* = 7.8, 5.1 Hz, 1H), 3.86 (td, *J* = 5.1, 2.4 Hz, 2H), 3.67 (dd, *J* = 21.0, 5.8 Hz, 2H), 3.10 (t,

$J$ = 2.5 Hz, 1H), 2.87 (q, $J$ = 6.7 Hz, 2H), 1.84 (s, 3H), 1.65 (ddt, $J$ = 14.4, 10.2, 5.5 Hz, 1H), 1.59 (dq, $J$ = 13.4, 6.7 Hz, 1H), 1.52 (dtd, $J$ = 13.6, 9.4, 5.0 Hz, 1H), 1.43 (ddd, $J$ = 9.1, 6.1, 2.3 Hz, 2H), 1.37 (s, 9H), 1.35 – 1.31 (m, 2H), 1.26 – 1.15 (m, 2H), 0.86 ppm (dd, $J$ = 24.6, 6.6 Hz, 6H); **$^{13}$C NMR** (176 MHz, DMSO-$d_6$): δ = 172.4, 171.8, 169.4, 168.5, 155.5, 80.9, 77.3, 73.0, 52.7, 51.0, 41.8, 40.5, 39.5, 31.3, 29.2, 28.3, 27.9, 24.1, 23.1, 22.6, 22.5, 21.6 ppm; **IR** (solid): ṽ = 3289, 3078, 2957, 2934, 2870, 1632, 1532, 1460met, 1440, 1391, 1367, 1277, 1250, 1171, 1097, 1040, 1016 cm$^{-1}$; **HRMS** (ESI): *m/z* calculated for $C_{24}H_{42}N_5O_6$ [*M*+H]$^+$: 496.3130, found: 496.3123.

**tert-Butyl-((S)-5-((S)-2-acetamido-4-methylpentanamido)-6-((2-(but-2-yn-1-ylamino)-2-oxoethyl)amino)-6-oxohexyl)carbamate (25e)**

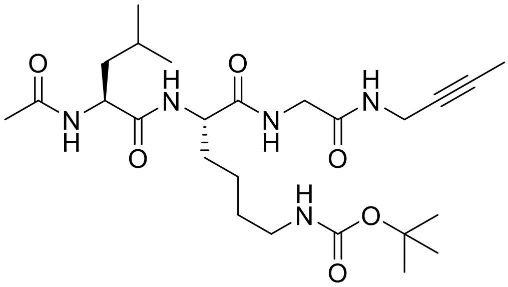

According to General Procedure F, *N*-acetyl tetrapeptide **25e** (5.8 mg, 1%) was obtained from *N*-Fmoc tetrapeptide **I39** (759 mg, 1.01 mmol) and commercially sourced acetic anhydride (0.11 mL, 1.10 mmol), following flash column chromatography (Biotage® Sfär Silica D Duo 10 g; 40 mL/min; 100%$_{v/v}$ cyclohexane (2 CV), linear gradient (8 CV): 0%$_{v/v}$ → 30%$_{v/v}$ acetone in cyclohexane, linear gradient (30 CV): 30%$_{v/v}$ → 100%$_{v/v}$ acetone in cyclohexane, 100%$_{v/v}$ acetone (10 CV)), and preparative high-performance liquid chromatography (Avantor® ACE® 10 AQ 250 × 21.2 mm; 20 mL/min, linear gradient (45 min): 2%$_{v/v}$ → 98%$_{v/v}$ acetonitrile in water (each containing 0.1%$_{v/v}$ formic acid); $t_R$ = 20.4 min).

White amorphous solid ; **$^1$H NMR** (400 MHz, DMSO-$d_6$): δ = 8.11 (t, $J$ = 5.5 Hz, 1H), 8.07 (t, $J$ = 5.9 Hz, 1H), 8.00 (d, $J$ = 7.9 Hz, 1H), 7.94 (d, $J$ = 7.6 Hz, 1H), 6.75 (t, $J$ = 5.6 Hz, 1H), 4.32 – 4.22 (m, 1H), 4.15 (q, $J$ = 7.6 Hz, 1H), 3.82 (dq, $J$ = 5.3, 2.8 Hz, 2H), 3.66 (qd, $J$ = 16.6, 5.6 Hz, 2H), 2.90 – 2.82 (m, 2H), 1.84 (s, 3H), 1.76 (t, $J$ = 2.4 Hz, 3H), 1.70 – 1.55 (m, 2H), 1.54 – 1.46 (m, 1H), 1.45 – 1.39 (m, 2H), 1.37 (s, 9H), 1.35 – 1.28 (m, 2H), 1.28 – 1.16 (m, 2H), 0.92 – 0.81 ppm (m, 6H); **$^{13}$C NMR** (151 MHz, DMSO-$d_6$): δ = 172.4, 171.8, 169.5, 168.4, 155.6, 78.0, 77.4, 76.3, 52.7, 51.1, 41.8, 40.5, 39.5, 31.4, 29.2, 28.3, 28.2, 24.2, 23.1, 22.6, 22.5, 21.6, 3.1 ppm; **IR** (film): ṽ = 3281 (br), 3081, 2956, 2933, 2869, 2361, 2252, 1779, 1689, 1628, 1538 cm$^{-1}$; **HRMS** (ESI): *m/z* calculated for $C_{25}H_{44}N_5O_6$ [*M*+H]$^+$: 510.3286, found: 510.3291.

***tert*-Butyl-((*S*)-5-((*S*)-2-acetamido-4-methylpentanamido)-6-((2-((1-ethynylcyclopropyl)amino)-2-oxoethyl)amino)-6-oxohexyl)carbamate (25f)**

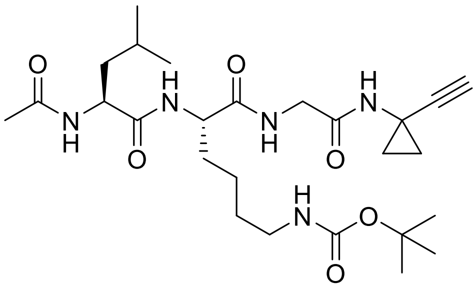

According to General Procedure F, *N*-acetyl tetrapeptide **25f** (56.6 mg, 13%) was obtained from *N*-Fmoc tetrapeptide **I40** (610 mg, 0.87 mmol) and commercially sourced acetic anhydride (87 µL, 0.92 mmol), following flash column chromatography (Biotage® Sfär Silica D Duo 10 g; 40 mL/min; 100%$_{v/v}$ cyclohexane (6 CV), linear gradient (8 CV): 0%$_{v/v}$ → 30%$_{v/v}$ acetone in cyclohexane, linear gradient (30 CV): 30%$_{v/v}$ → 100%$_{v/v}$ acetone in cyclohexane, 100%$_{v/v}$ acetone (10 CV)), and preparative high-performance liquid chromatography (Avantor® ACE® 10 AQ 250 × 21.2 mm; 20 mL/min, linear gradient (45 min): 2%$_{v/v}$ → 98%$_{v/v}$ acetonitrile in water (each containing 0.1%$_{v/v}$ formic acid); $t_R$ = 20.1 min).

White amorphous solid ; **$^1$H NMR** (400 MHz, DMSO-*d$_6$*): δ = 8.30 (s, 1H), 8.05 (t, *J* = 5.8 Hz, 1H), 7.98 (t, *J* = 8.1 Hz, 2H), 6.75 (t, *J* = 5.7 Hz, 1H), 4.27 (qd, *J* = 7.3, 4.3 Hz, 1H), 4.11 (td, *J* = 8.0, 5.2 Hz, 1H), 3.69 – 3.50 (m, 2H), 2.94 (s, 1H), 2.87 (q, *J* = 6.6 Hz, 2H), 1.84 (s, 3H), 1.68 – 1.55 (m, 2H), 1.54 – 1.46 (m, 1H), 1.45 – 1.40 (m, 2H), 1.36 (s, 9H), 1.35 – 1.29 (m, 2H), 1.28 – 1.14 (m, 2H), 1.07 (m, 2H), 1.08 – 0.83 (m, 2H), 0.86 ppm (dd, *J* = 16.9, 6.5 Hz, 6H); **$^{13}$C NMR** (151 MHz, DMSO-*d$_6$*): δ = 172.5, 171.8, 169.5, 169.0, 155.6, 86.2, 77.4, 68.6, 52.8, 51.1, 41.9, 40.5, 39.5, 31.3, 29.2, 28.3, 24.2, 23.1, 22.7, 22.5, 21.8, 21.6, 16.8 ppm (2C); **IR** (film): ṽ = 3307 (br), 2958, 2118, 1645, 1535, 1452, 1367, 1287, 1251, 1171 cm$^{-1}$; **HRMS** (ESI): *m/z* calculated for $C_{26}H_{44}N_5O_6$ [*M*+H]$^+$: 522.3286, found: 522.3288.

***tert*-Butyl-((*S*)-5-((*S*)-2-acetamido-4-methylpentanamido)-6-((2-((1-(1-chlorovinyl)cyclopropyl)amino)-2-oxoethyl)amino)-6-oxohexyl)carbamate (25c)**

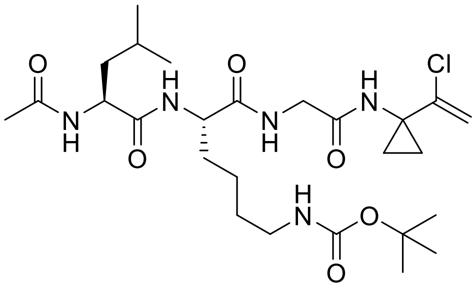

According to General Procedure F, *N*-acetyl tetrapeptide **25c** (10.03 mg, ~ 5%) was obtained from *N*-Fmoc tetrapeptide **I41** (244 mg, 1:1 mixture of **I41** and **I40** as per $^1$H NMR analysis) and commercially sourced acetic anhydride (34.5 µL, 0.37 mmol), following flash column chromatography (Biotage® Sfär Silica D Duo 10 g; 40 mL/min; 100%$_{v/v}$ cyclohexane (2 CV), linear gradient (8 CV): 0%$_{v/v}$ → 30%$_{v/v}$ acetone in cyclohexane, linear gradient (30 CV): 30%$_{v/v}$ → 100%$_{v/v}$ acetone in cyclohexane, 100%$_{v/v}$ acetone (10 CV)), and preparative high-performance liquid chromatography (Avantor® ACE® 10 AQ 250 × 21.2 mm; 20 mL/min, linear gradient (30 min): 2%$_{v/v}$ → 98%$_{v/v}$ acetonitrile in water (each containing 0.1%$_{v/v}$ formic acid); $t_R$ = 17.4 min).

White amorphous solid ; $^{1}$**H NMR** (600 MHz, DMSO-*d6*): δ = 8.32 (s, 1H), 8.12 (t, *J* = 5.8 Hz, 1H), 7.99 (t, *J* = 8.3 Hz, 2H), 6.74 (t, *J* = 5.7 Hz, 1H), 5.43 (d, *J* = 1.8 Hz, 1H), 5.23 (d, *J* = 1.8 Hz, 1H), 4.30 – 4.23 (m, 1H), 4.11 (q, *J* = 6.3 Hz, 1H), 3.69 – 3.55 (m, 2H), 2.87 (q, *J* = 6.7 Hz, 2H), 1.84 (s, 3H), 1.68 – 1.61 (m, 1H), 1.60 – 1.55 (m, 1H), 1.54 – 1.47 (m, 1H), 1.42 (m, 2H), 1.37 (s, 9H), 1.36 – 1.31 (m, 2H), 1.29 – 1.16 (m, 4H), 0.93 (m, 2H), 0.86 ppm (dd, *J* = 24.8, 6.6 Hz, 6H); $^{13}$**C NMR** (151 MHz, DMSO-*d6*): δ = 172.5, 172.0, 169.4, 169.1, 155.5, 141.2, 112.2, 77.3, 52.8, 51.0, 42.2, 40.5, 39.5, 35.9, 31.2, 29.2, 28.3, 24.1, 23.0, 22.7, 22.4, 21.6, 14.4, 14.2 ppm; **IR** (film): ṽ = 3281, 3067, 2954, 2936, 2868, 2247, 1685, 1630, 1532 cm$^{-1}$; **HRMS** (ESI): *m/z* calculated for $C_{26}H_{45}ClN_5O_6$ [*M*+H]$^+$: 558.3053, found: 558.3064.

***tert*-Butyl-((*S*)-5-((*S*)-2-acetamido-4-methylpentanamido)-6-oxo-6-((2-oxo-2-((3-phenylprop-2-yn-1-yl)amino)ethyl)amino)hexyl)carbamate (25g)**

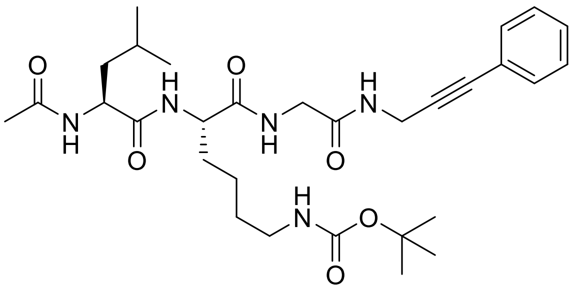

According to General Procedure F, *N*-acetyl tetrapeptide **25g** (93.0 mg, 21%) was obtained from *N*-Fmoc tetrapeptide **I42** (590 mg, 0.78 mmol) and commercially sourced acetic anhydride (80 µL, 0.85 mmol), following flash column chromatography (Biotage® Sfär Silica D Duo 10 g; 40 mL/min; 100%$_{v/v}$ cyclohexane (2 CV), linear gradient (8 CV): 0%$_{v/v}$ → 30%$_{v/v}$ acetone in cyclohexane, linear gradient (30 CV): 30%$_{v/v}$ → 100%$_{v/v}$ acetone in cyclohexane, 100%$_{v/v}$ acetone (10 CV)), and preparative high-performance liquid chromatography (Avantor® ACE® 10 AQ 250 × 21.2 mm; 20 mL/min, linear gradient (45 min): 2%$_{v/v}$ → 98%$_{v/v}$ acetonitrile in water (each containing 0.1%$_{v/v}$ formic acid); $t_R$ = 24.4 min).

White amorphous solid ; $^{1}$**H NMR** (400 MHz, DMSO-*d6*): δ = 8.31 (t, *J* = 5.5 Hz, 1H), 8.13 (t, *J* = 5.9 Hz, 1H), 8.00 (d, *J* = 7.9 Hz, 1H), 7.96 (d, *J* = 7.6 Hz, 1H), 7.45 – 7.31 (m, 5H), 6.75 (t, *J* = 5.6 Hz, 1H), 4.28 (td, *J* = 8.3, 6.3 Hz, 1H), 4.18 (dd, *J* = 8.0, 5.4 Hz, 1H), 4.14 (dd, *J* = 5.5, 3.2 Hz, 2H), 3.71 (dd, *J* = 11.7, 5.8 Hz, 2H), 2.87 (q, *J* = 6.6 Hz, 2H), 1.84 (s, 3H), 1.71 – 1.63 (m, 1H), 1.62 – 1.56 (m, 1H), 1.55 – 1.47 (m, 1H), 1.46 – 1.39 (m, 2H), 1.36 (s, 9H), 1.35 – 1.29 (m, 2H), 1.28 – 1.16 (m, 2H), 0.84 ppm (dd, *J* = 14.1, 6.6 Hz, 6H); $^{13}$**C NMR** (101 MHz, DMSO-*d6*): δ = 172.4, 171.9, 169.4, 168.6, 155.6, 131.4, 128.7, 128.6, 122.2, 86.9, 81.6, 77.3, 52.7, 51.1, 41.9, 40.5, 39.5, 31.4, 29.2, 28.6, 28.3, 24.2, 23.0, 22.6, 22.5, 21.6 ppm; **IR** (film): ṽ = 3287 (br), 3080, 2971, 2883, 2117, 1683, 1634, 1451, 1368, 1287, 1252, 1172 cm$^{-1}$; **HRMS** (ESI): *m/z* calculated for $C_{30}H_{46}N_5O_6$ [*M*+H]$^+$: 572.3443, found: 572.3440.

***tert*-Butyl-((*S*)-5-((*S*)-4-methyl-2-(2,2,2-trifluoroacetamido)pentanamido)-6-oxo-6-((2-oxo-2-(prop-2-yn-1-ylamino)ethyl)amino)hexyl)carbamate (25a)**

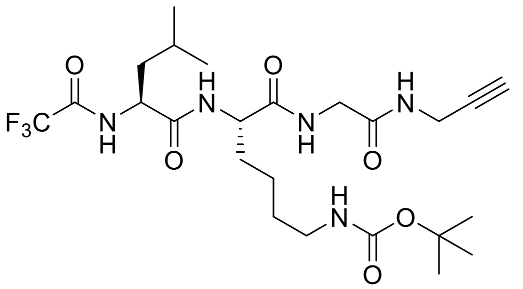

According to General Procedure F, *N*-acetyl tetrapeptide **25a** (28.3 mg, 11%) was obtained from *N*-Fmoc tetrapeptide **I38** (300 mg, 0.44 mmol) and commercially sourced trifluoroacetic anhydride (65 µL, 0.47 mmol), following flash column chromatography (Biotage® Sfär Silica D Duo 25 g; 80 mL/min; 100%$_{v/v}$ cyclohexane (2 CV), linear gradient (8 CV): 0%$_{v/v}$ → 30%$_{v/v}$ acetone in cyclohexane, linear gradient (30 CV): 30%$_{v/v}$ → 100%$_{v/v}$ acetone in cyclohexane, 100%$_{v/v}$ acetone (2 CV)), and preparative high-performance liquid chromatography (Avantor® ACE® 10 AQ 250 × 21.2 mm; 20 mL/min, linear gradient (45 min): 2%$_{v/v}$ → 98%$_{v/v}$ acetonitrile in water (each containing 0.1%$_{v/v}$ formic acid); $t_R$ = 22.9 min).

White amorphous solid ; **$^1$H NMR** (400 MHz, DMSO-*d$_6$*): δ = 9.43 (d, *J* = 8.2 Hz, 1H), 8.18 – 8.08 (m, 2H), 8.07 (t, *J* = 5.8 Hz, 1H), 6.63 (t, *J* = 5.7 Hz, 1H), 4.42 (ddd, *J* = 10.6, 8.1, 4.2 Hz, 1H), 4.20 (td, *J* = 8.0, 5.6 Hz, 1H), 3.87 (dt, *J* = 5.3, 2.5 Hz, 2H), 3.68 (d, *J* = 5.8 Hz, 2H), 3.04 (t, *J* = 2.5 Hz, 1H), 2.88 (q, *J* = 6.6 Hz, 2H), 1.71 – 1.61 (m, 2H), 1.60 – 1.47 (m, 3H), 1.37 (s, 9H), 1.36 – 1.32 (m, 2H), 1.31 – 1.17 (m, 2H), 0.94 – 0.79 ppm (m, 6H); **$^{13}$C NMR** (101 MHz, DMSO-*d$_6$*): δ = 171.8, 170.6, 168.5, 156.3 (q, *J* = 36.3 Hz), 155.6, 115.9 (t, *J* = 288.0 Hz), 81.0, 77.4, 73.1, 52.9, 51.7, 41.8, 39.2 (2C), 31.5, 29.2, 28.3, 27.9, 24.3, 23.0, 22.6, 21.1 ppm; **$^{19}$F NMR** (376 MHz, DMSO-*d$_6$*): δ = −73.9 ppm (s, 3F); **IR** (film): ṽ = 3288 (br), 2347, 2253, 1673 (br), 1542, 1368, 1180, 1167 cm$^{-1}$; **HRMS** (ESI): *m/z* calculated for $C_{24}H_{39}F_3N_5O_6$ [*M*+H]$^+$: 550.2847, found: 550.2847.

**(*S*)-5-((*S*)-2-Acetamido-4-methylpentanamido)-6-oxo-6-((2-oxo-2-(prop-2-yn-1-ylamino)ethyl)amino)hexan-1-aminium chloride (25)**

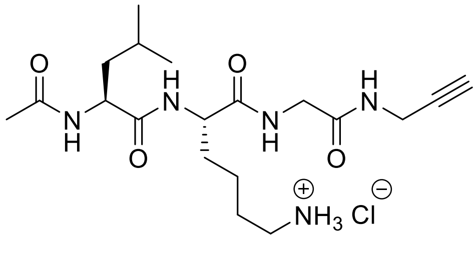

According to General Procedure G, tetrapeptide **25** (6.3 mg, 60%) was obtained from *N*-Boc tetrapeptide **25d** (12.2 mg, 0.025 mmol).

White amorphous solid ; **$^1$H NMR** (400 MHz, DMSO-*d$_6$*): δ = 8.26 (t, *J* = 5.5 Hz, 1H), 8.09 (t, *J* = 5.8 Hz, 1H), 8.01 (t, *J* = 7.3 Hz, 2H), 7.67 (s, 2H), 4.30 – 4.17 (m, 2H), 3.86 (ddd, *J* = 5.7, 2.6, 1.4 Hz, 2H), 3.69 (dd, *J* = 5.6, 1.5 Hz, 2H), 3.11 (t, *J* = 2.5 Hz, 1H), 2.75 (d, *J* = 8.4 Hz, 2H), 1.84 (s, 3H), 1.68 (dt, *J* = 13.8, 6.3 Hz, 1H), 1.59 (dt, *J* = 13.5, 6.8 Hz, 1H), 1.55 – 1.46 (m, 3H), 1.43 (dd, *J* = 8.3, 6.3 Hz, 2H), 1.37 – 1.26 (m, 2H), 0.86 ppm (dd, *J* = 16.4, 6.6 Hz, 6H); **$^{13}$C NMR** (151 MHz, DMSO-*d$_6$*): δ = 172.5, 171.7, 169.5, 168.5, 80.9,

73.1, 52.3, 51.1, 41.7, 40.4, 38.7, 31.0, 27.9, 26.5, 24.2, 23.1, 22.5, 22.1, 21.5 ppm; **IR** (film): ṽ = 3010, 2898, 2855, 2265, 1734, 1636, 1541, 1437, 1291 $cm^{-1}$; **HRMS** (ESI): *m/z* calculated for $C_{19}H_{34}N_5O_4$ [*M−Cl⁻*]⁺: 396.2605, found: 396.2601.

**(*S*)-5-((*S*)-2-Acetamido-4-methylpentanamido)-6-((2-(but-2-yn-1-ylamino)-2-oxoethyl)amino)-6-oxohexan-1-aminium chloride (25j)**

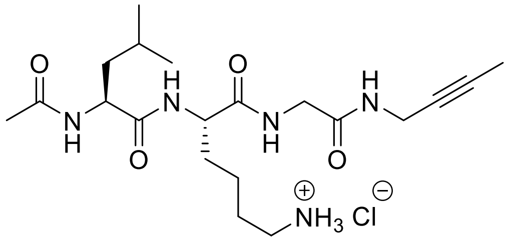


According to General Procedure G, tetrapeptide **25j** (2.0 mg, 50%) was obtained from *N*-Boc tetrapeptide **25e** (3.9 mg, 0.008 mmol).

White amorphous solid ; **¹H NMR** (400 MHz, DMSO-*d₆*): δ = 8.35 (s, 2H), 8.18 (t, *J* = 5.5 Hz, 1H), 8.10 (t, *J* = 5.9 Hz, 1H), 8.04 (dd, *J* = 10.4, 7.8 Hz, 2H), 4.30 – 4.23 (m, 1H), 4.22 – 4.16 (m, 1H), 3.82 (p, *J* = 2.5 Hz, 2H), 3.73 – 3.61 (m, 2H), 2.71 (t, *J* = 7.4 Hz, 2H), 1.84 (s, 3H), 1.76 (t, *J* = 2.5 Hz, 3H), 1.72 – 1.64 (m, 1H), 1.63 – 1.57 (m, 1H), 1.56 – 1.53 (m, 1H), 1.49 (m, 2H), 1.43 (t, *J* = 7.3 Hz, 2H), 1.34 – 1.21 (m, 2H), 0.86 ppm (dd, *J* = 16.5, 6.6 Hz, 6H); **¹³C NMR** (151 MHz, DMSO-*d₆*): δ = 172.4, 171.7, 169.5, 168.3, 78.0, 76.2, 52.3, 51.2, 41.7, 40.5, 38.8, 31.1, 28.2, 27.0, 24.2, 23.1, 22.5, 22.1, 21.5, 3.0 ppm; **IR** (film): ṽ = 3296 (br), 2957, 2925, 2856, 2361, 2270, 1652, 1547, 1469, 1237 $cm^{-1}$; **HRMS** (ESI): *m/z* calculated for $C_{20}H_{36}N_5O_4$ [*M−Cl⁻*]⁺: 410.2762, found: 410.2760.

**(*S*)-5-((*S*)-2-Acetamido-4-methylpentanamido)-6-((2-((1-ethynylcyclopropyl)amino)-2-oxoethyl)amino)-6-oxohexan-1-aminium chloride (25h)**

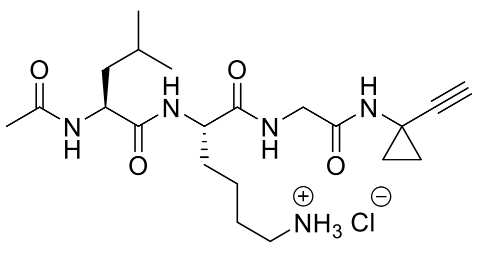


According to General Procedure G, tetrapeptide **25h** (7.6 mg, 77%) was obtained from *N*-Boc tetrapeptide **25f** (11.5 mg, 0.022 mmol).

White amorphous solid ; **¹H NMR** (400 MHz, DMSO-*d₆*): δ = 8.39 (s, 1H), 8.10 – 8.01 (m, 3H), 7.80 (s, 2H), 4.31 – 4.21 (m, 1H), 4.18 (td, *J* = 8.3, 5.2 Hz, 1H), 3.69 – 3.53 (m, 2H), 2.95 (s, 1H), 2.74 (s, 2H), 1.84 (s, 3H), 1.74 – 1.64 (m, 1H), 1.63 – 1.55 (m, 2H) 1.53 (m, 2H), 1.44 (t, *J* = 7.3 Hz, 2H), 1.37 – 1.25 (m, 2H), 1.10 – 1.05 (m, 2H), 0.96 – 0.91 (m, 2H), 0.87 ppm (dd, *J* = 16.7, 6.5 Hz, 6H); **¹³C NMR** (151 MHz, DMSO-*d₆*): δ = 172.5, 171.6, 169.5, 168.9, 86.2, 68.6, 52.4, 51.1, 41.8, 40.5, 38.6, 31.0, 26.5, 24.1, 23.1, 22.5, 22.1, 21.8, 21.5, 16.7 ppm (2C); **IR** (film): ṽ = 2928, 2264, 1733, 1644, 1548, 1439, 1292, 1109 $cm^{-1}$; **HRMS** (ESI): *m/z* calculated for $C_{21}H_{36}N_5O_4$ [*M−Cl⁻*]⁺: 422.2762, found: 422.2742.

**(*S*)-5-((*S*)-2-Acetamido-4-methylpentanamido)-6-oxo-6-((2-oxo-2-((3-phenylprop-2-yn-1-yl)amino)ethyl)amino)hexan-1-aminium chloride (25i)**

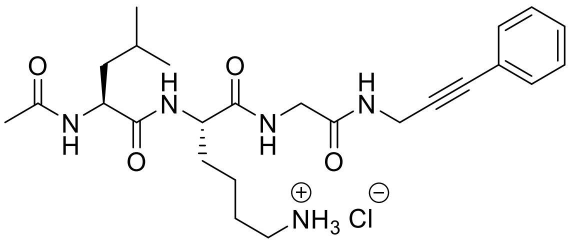

According to General Procedure G, tetrapeptide **25i** (4.1 mg, 89%) was obtained from *N*-Boc-protected tetrapeptide **25g** (5.2 mg, 0.009 mmol).

White amorphous solid ; **$^{1}$H NMR** (400 MHz, DMSO-*d$_6$*): δ = 8.37 (t, *J* = 5.5 Hz, 1H), 8.24 (s, 1H), 8.13 (t, *J* = 5.8 Hz, 1H), 8.03 (d, *J* = 7.8 Hz, 2H), 7.45 – 7.33 (m, 5H), 4.25 (dt, *J* = 14.9, 7.2 Hz, 2H), 4.17 – 4.11 (m, 2H), 3.74 – 3.70 (m, 2H), 2.74 (t, *J* = 7.5 Hz, 2H), 1.84 (s, 3H), 1.74 – 1.64 (m, 1H), 1.64 – 1.54 (m, 2H), 1.54 – 1.46 (m, 2H), 1.46 – 1.40 (m, 2H), 1.34 – 1.28 (m, 2H), 0.85 ppm (dd, *J* = 14.7, 6.6 Hz, 6H); **$^{13}$C NMR** (151 MHz, DMSO-*d$_6$*): δ = 172.5, 171.7, 169.5, 168.5, 131.3, 128.7, 128.6, 122.2, 86.8, 81.6, 52.3, 51.2, 41.8, 40.4, 38.7, 31.1, 28.6, 26.6, 24.2, 23.1, 22.5, 22.1, 21.5 ppm; **IR** (film): ṽ = 3270 (br), 2930, 2864, 2335, 2255, 1638, 1549 cm$^{-1}$; **HRMS** (ESI): *m/z* calculated for $C_{25}H_{38}N_5O_4$ [*M−Cl*$^-$]$^+$: 472.2918, found: 472.2916.

**(*S*)-5-((*S*)-4-Methyl-2-(2,2,2-trifluoroacetamido)pentanamido)-6-oxo-6-((2-oxo-2-(prop-2-yn-1-ylamino)ethyl)amino)hexan-1-aminium chloride (25k)**

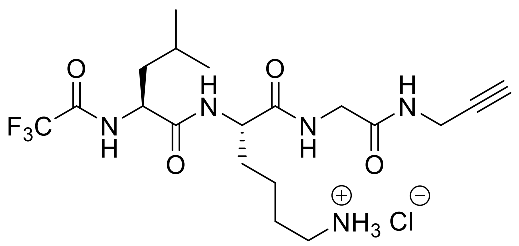

According to General Procedure G, tetrapeptide **25k** (5.5 mg, 73%) was obtained from *N*-Boc tetrapeptide **25a** (8.4 mg, 0.015 mmol).

White amorphous solid ; **$^{1}$H NMR** (400 MHz, DMSO-*d$_6$*): δ = 9.54 (d, *J* = 7.9 Hz, 1H), 8.29 (dd, *J* = 9.5, 6.5 Hz, 2H), 8.16 (t, *J* = 5.8 Hz, 1H), 7.64 (s, 3H), 4.46 – 4.35 (m, 1H), 4.25 (q, *J* = 8.0 Hz, 1H), 3.90 – 3.84 (m, 2H), 3.73 – 3.62 (m, 2H), 3.12 (t, *J* = 2.5 Hz, 1H), 2.74 (d, *J* = 7.1 Hz, 2H), 1.73 – 1.60 (m, 2H), 1.60 – 1.46 (m, 5H), 1.35 – 1.27 (m, 2H), 0.87 ppm (dd, *J* = 14.0, 6.3 Hz, 6H); **$^{13}$C NMR** (151 MHz, DMSO-*d$_6$*): δ = 171.6, 170.7, 168.5, 156.3 (d, *J* = 36.9 Hz), 115.9 (d, *J* = 288.1 Hz), 81.0, 73.1, 52.5, 51.7, 41.7, 39.2, 38.7, 31.2, 27.9, 26.6, 24.3, 23.0, 22.1, 21.1 ppm; **$^{19}$F NMR** (376 MHz, DMSO-*d$_6$*): δ = -73.8 ppm (m, 3F); **IR** (film): ṽ = 3289 (br), 2961 (br), 2253, 1719, 1648, 1541, 1389, 1261, 1215, 1185, 1073 cm$^{-1}$; **HRMS** (ESI): *m/z* calculated for $C_{19}H_{31}F_3N_5O_4$ [*M−Cl*$^-$]$^+$: 450.2323, found: 450.2325.

**(*S*)-2-((*S*)-4-Methyl-2-(2,2,2-trifluoroacetamido)pentanamido)-*N*-(2-oxo-2-(prop-2-yn-1-ylamino)ethyl)-6-(2,2,2-trifluoroacetamido)hexanamide (25b)**

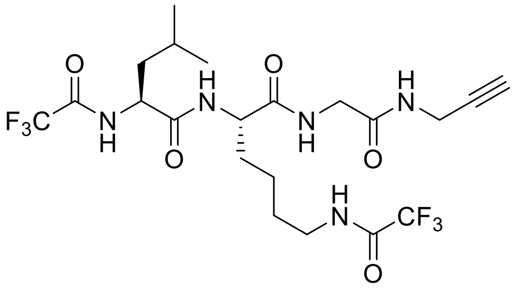

*N*-Boc tetrapeptide **25k** (60 mg, 0.110 mmol) was deprotected according to General Procedure G. Commercially sourced 4-(*N,N*-dimethylamino)pyridine (1.05 equiv.), anhydrous tetrahydrofuran (0.09 M) and commercially sourced trifluoroacetic anhydride (1.05 equiv.) were added, and the reaction stirred under reflux (1 h). After cooling to ambient temperature, the mixture was concentrated under reduced pressure, and the crude residue was redissolved in ethyl acetate. The mixture was washed with aqueous HCl (1 M), saturated aqueous $NaHCO_3$ and brine, then dried over anhydrous $Na_2SO_4$, filtered and concentrated under reduced pressure. The residue was purified via flash column chromatography (Biotage® Sfär Silica D Duo 5 g; 18 mL/min; $100\%_{v/v}$ cyclohexane (3 CV), linear gradient (15 CV): $0\%_{v/v}$ → $100\%_{v/v}$ acetone in cyclohexane, $100\%_{v/v}$ acetone (3 CV)) and high-performance liquid chromatography (Avantor® ACE® 10 AQ 250 × 21.2 mm; 20 mL/min, linear gradient (45 min): $2\%_{v/v}$ → $98\%_{v/v}$ acetonitrile in water (each containing $0.1\%_{v/v}$ formic acid); $t_R$ = 21.0 min), to give the desired peptide **25b** (0.43 mg, 1%).

White amorphous solid; **$^1$H NMR** (600 MHz, DMSO-*d₆*): δ = 9.53 (d, *J* = 7.9 Hz, 1H), 9.38 (s, 1H), 8.24 (t, *J* = 5.5 Hz, 2H), 8.18 (t, *J* = 5.8 Hz, 1H), 4.45 – 4.38 (m, 1H), 4.22 (td, *J* = 8.0, 5.3 Hz, 1H), 3.86 (ddd, *J* = 6.1, 3.8, 2.5 Hz, 2H), 3.68 (d, *J* = 5.9 Hz, 2H), 3.15 (q, *J* = 6.6 Hz, 2H), 3.14 – 3.08 (m, 1H), 1.72 – 1.57 (m, 2H), 1.59 – 1.43 (m, 5H), 1.28 (ddd, *J* = 24.2, 16.3, 8.9 Hz, 2H), 0.87 ppm (dd, *J* = 20.2, 6.4 Hz, 6H); **$^{13}$C NMR** (151 MHz, DMSO-*d₆*): δ = 171.7, 170.6, 168.4, 156.2 (d, *J* = 36.4 Hz), 156.1 (d, *J* = 35.8 Hz), 116.0 (d, *J* = 288.3 Hz), 115.9 (d, *J* = 287.7 Hz), 81.0, 73.0, 52.7, 51.7, 41.7, 39.5, 39.0, 31.4, 27.9, 27.8(6), 24.3, 23.0, 22.5, 21.1 ppm; **$^{19}$F NMR** (565 MHz, DMSO-*d₆*): δ = −73.9 (s, 3F), −74.4 ppm (s, 3F); **IR** (film): ṽ = 3293 (br), 3094, 2949, 2126, 1704, 1640, 1548, 1209, 1186, 1158 $cm^{-1}$; **HRMS** (ESI): *m/z* calculated for $C_{21}H_{30}F_6N_5O_5$ [*M*+H]$^+$: 546.2146, found: 546.2141.

#### $^1$H, $^{13}$C, and $^{19}$F NMR spectra of final compounds

Compound **25d**

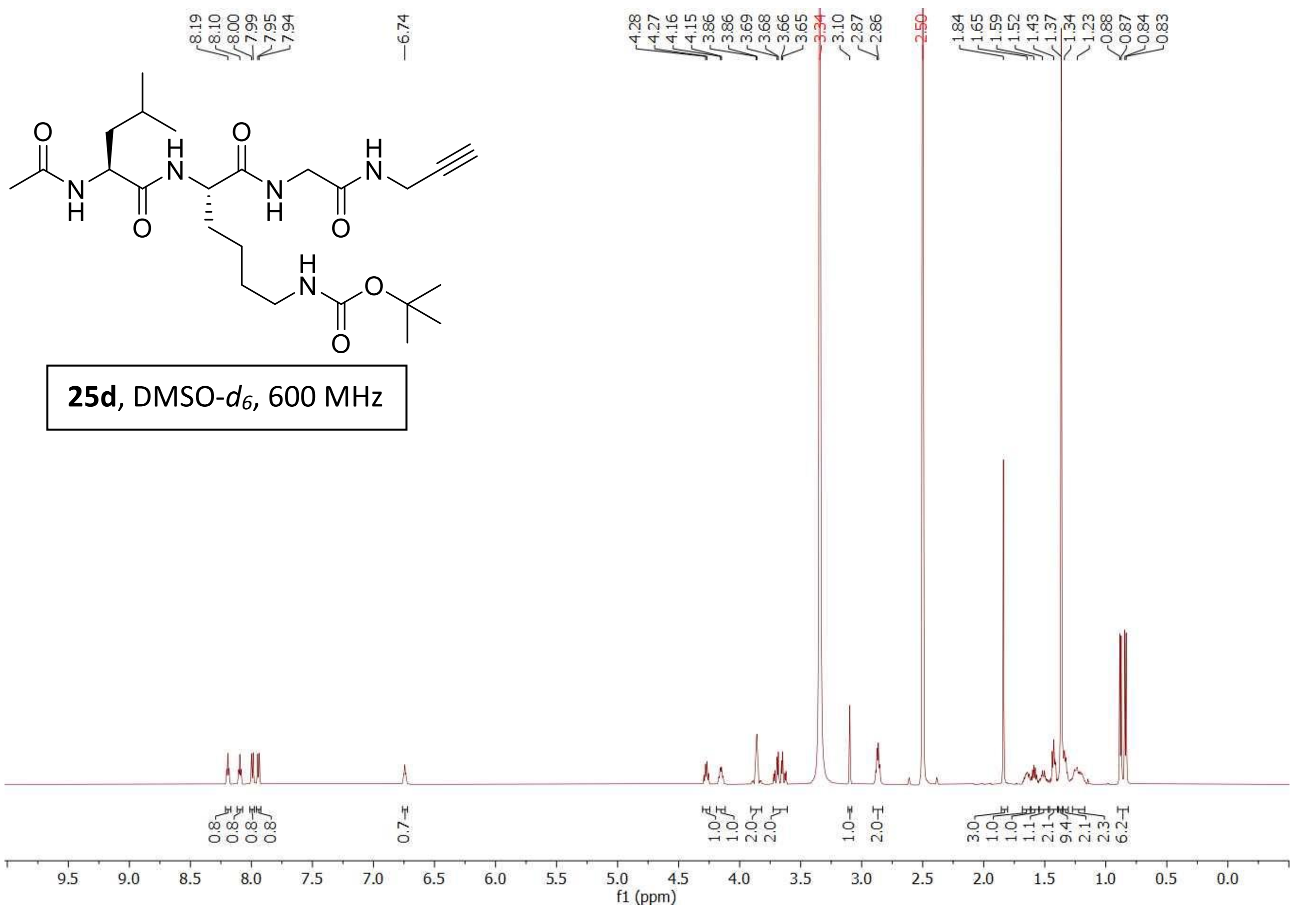


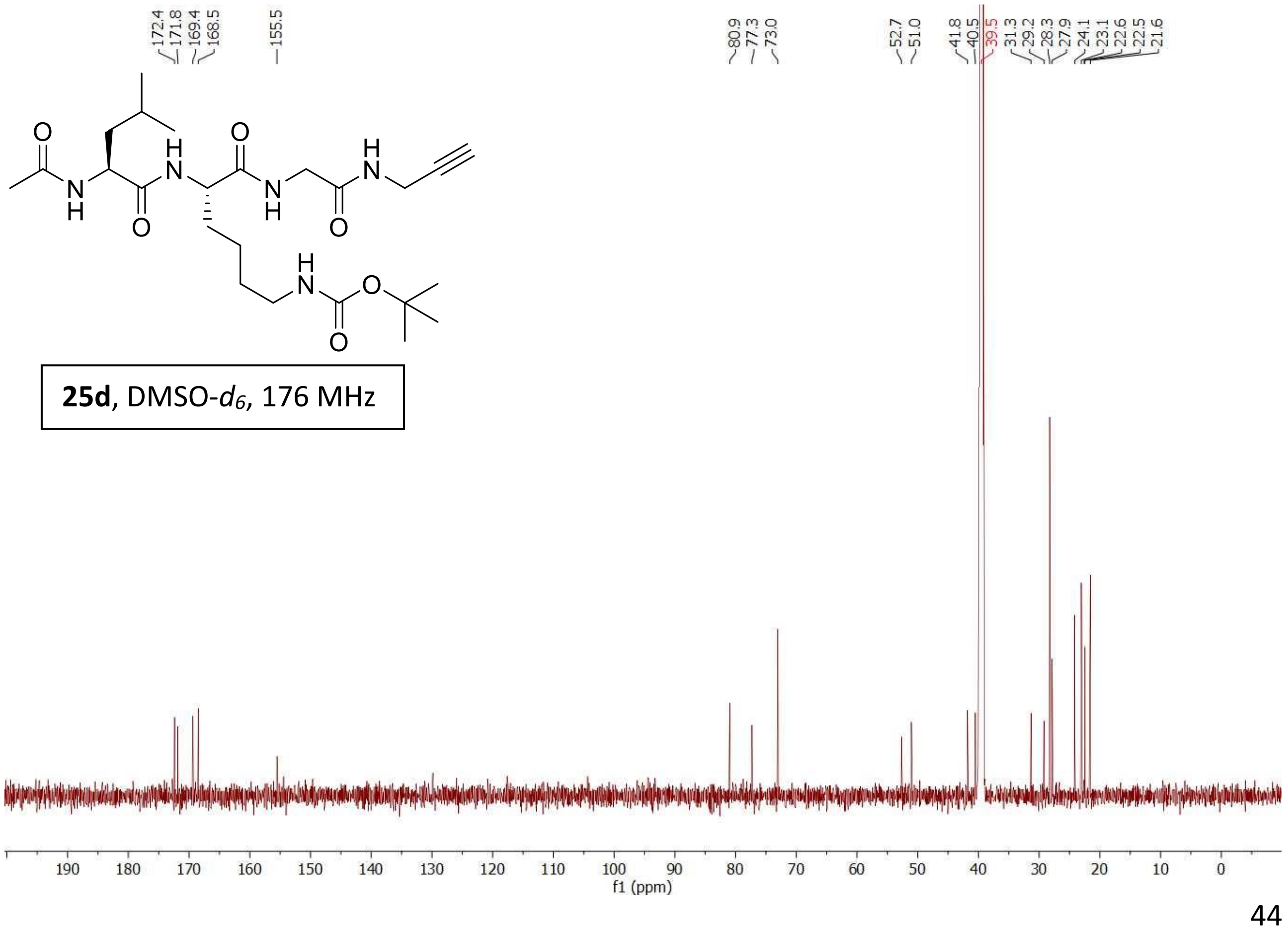

Compound **25e**

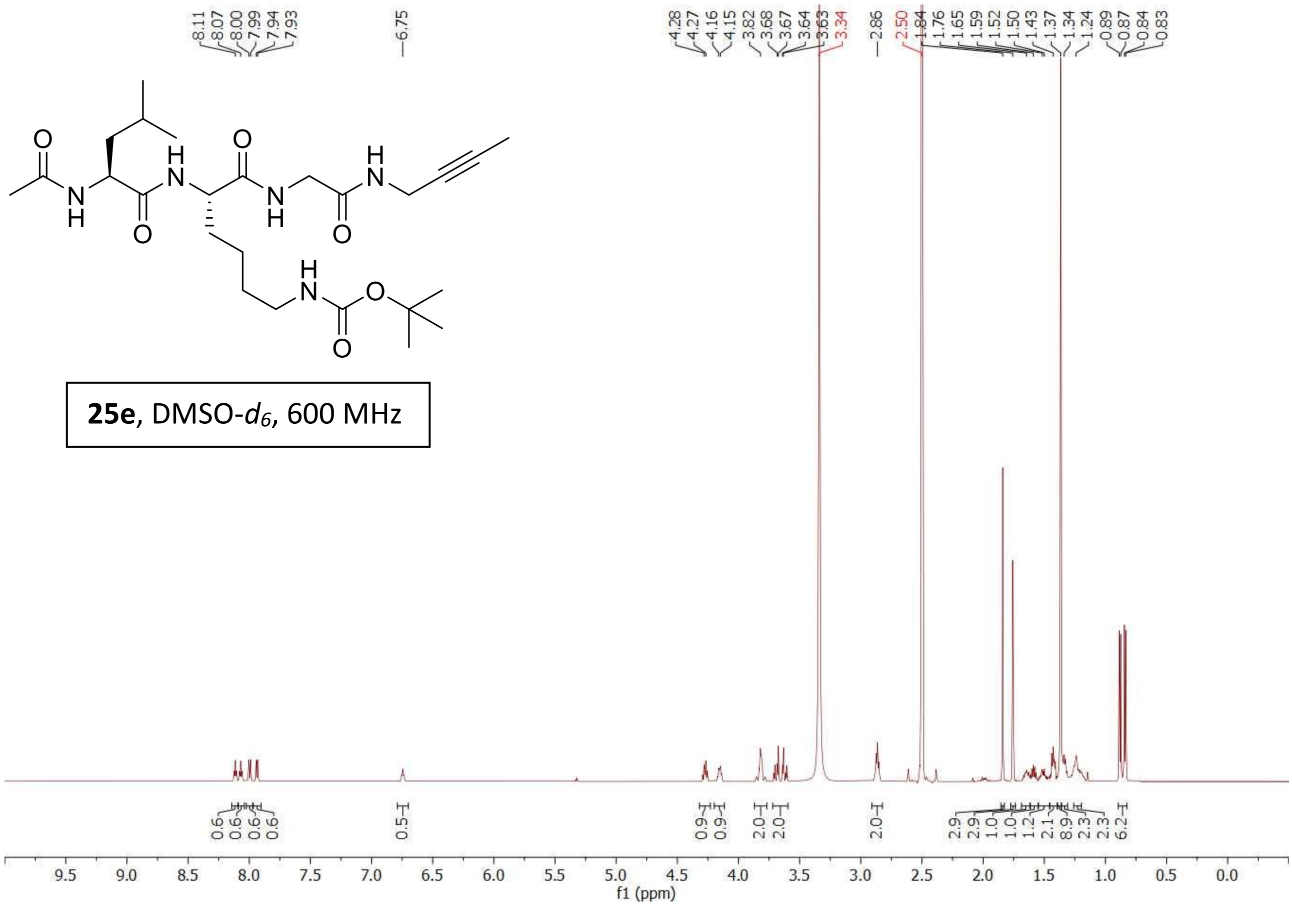


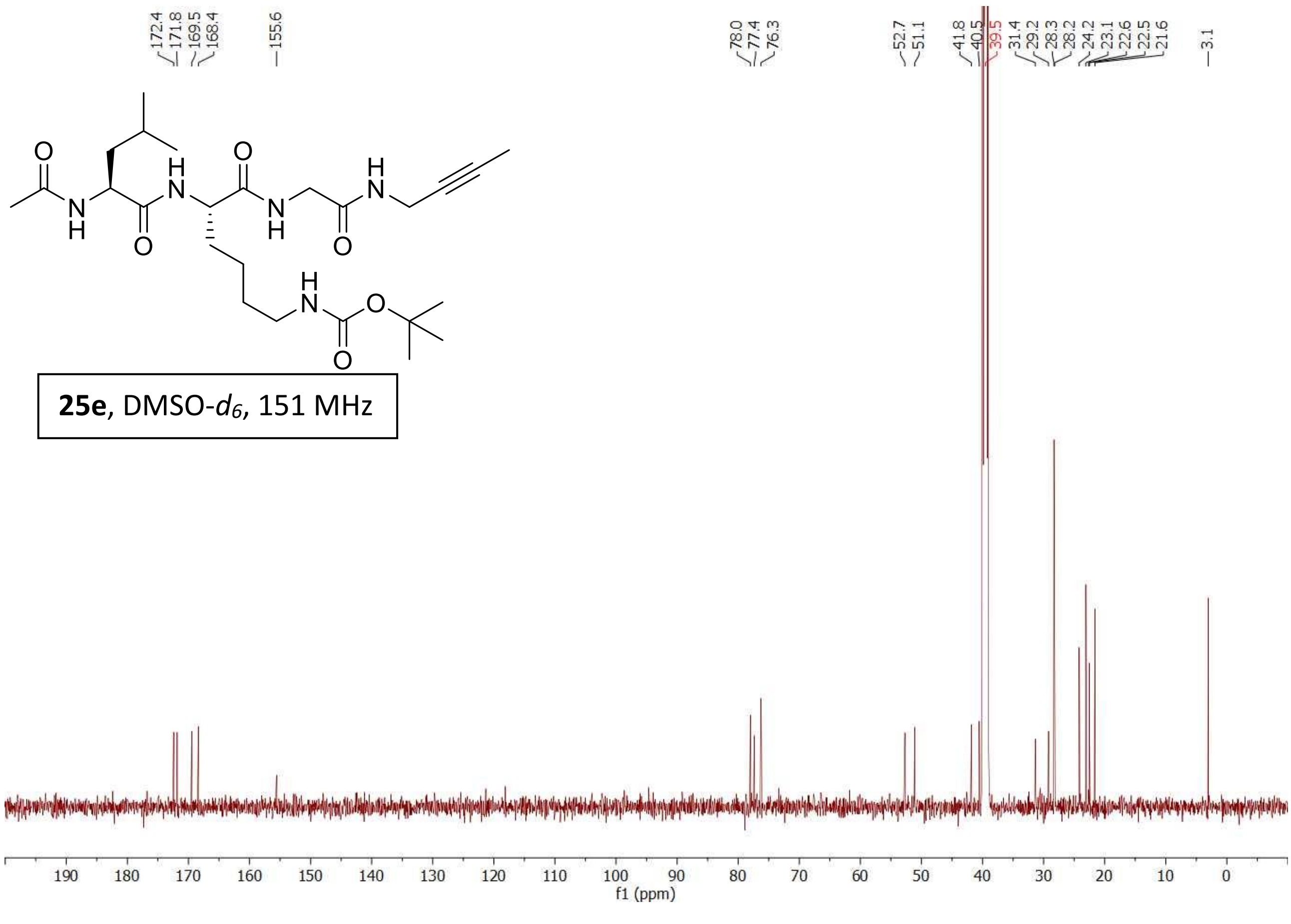

Compound **25f**

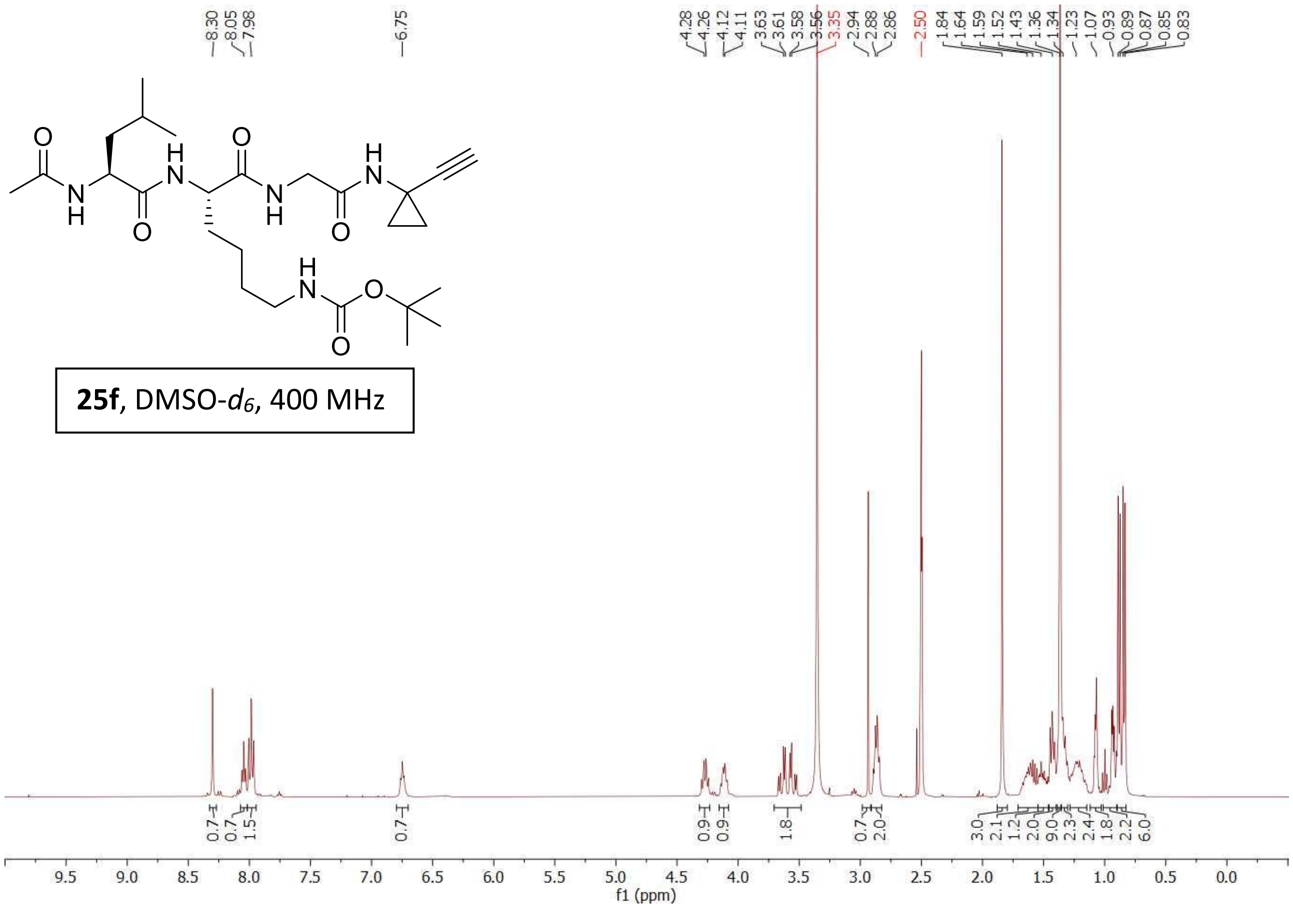


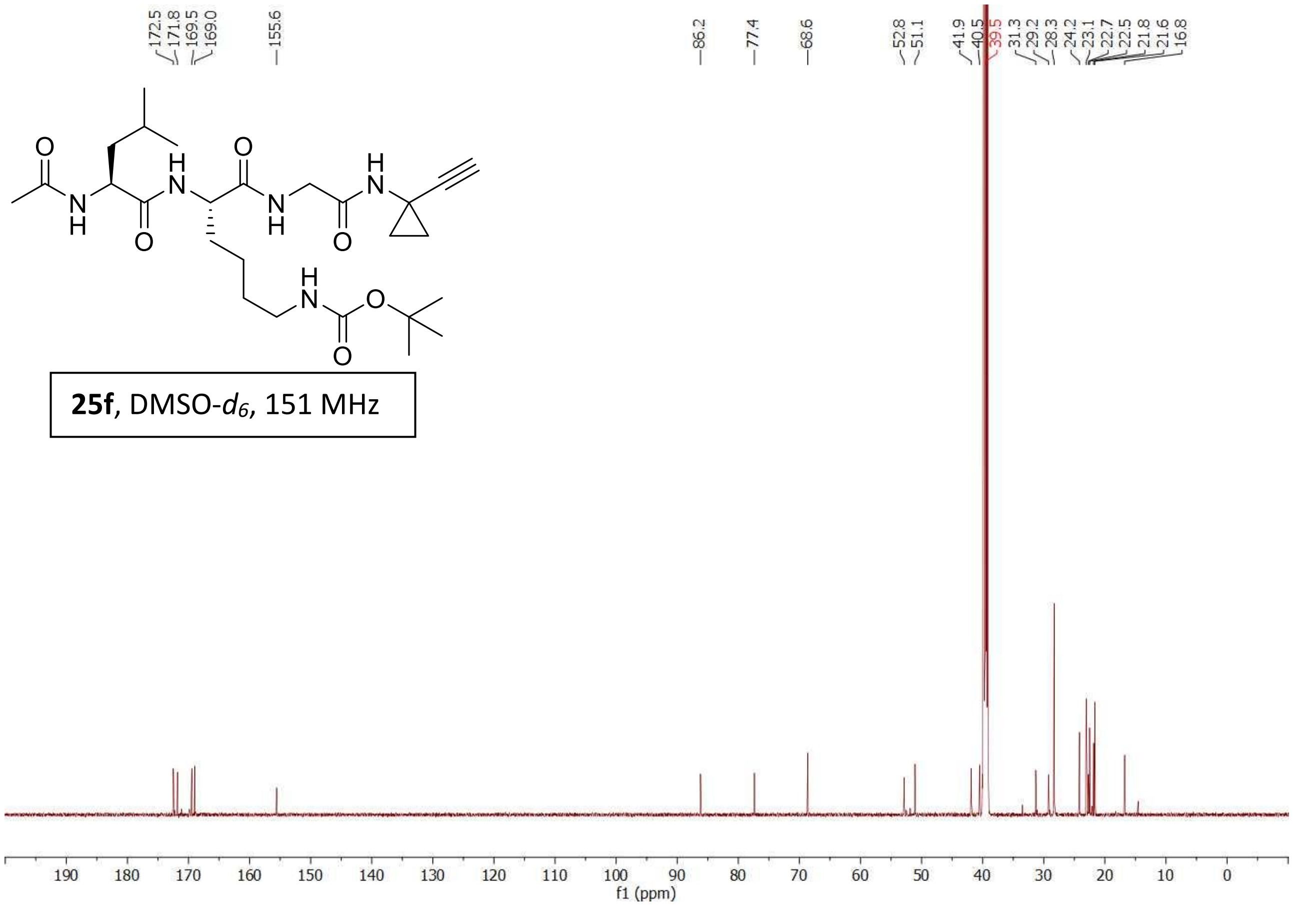

Compound **25c**

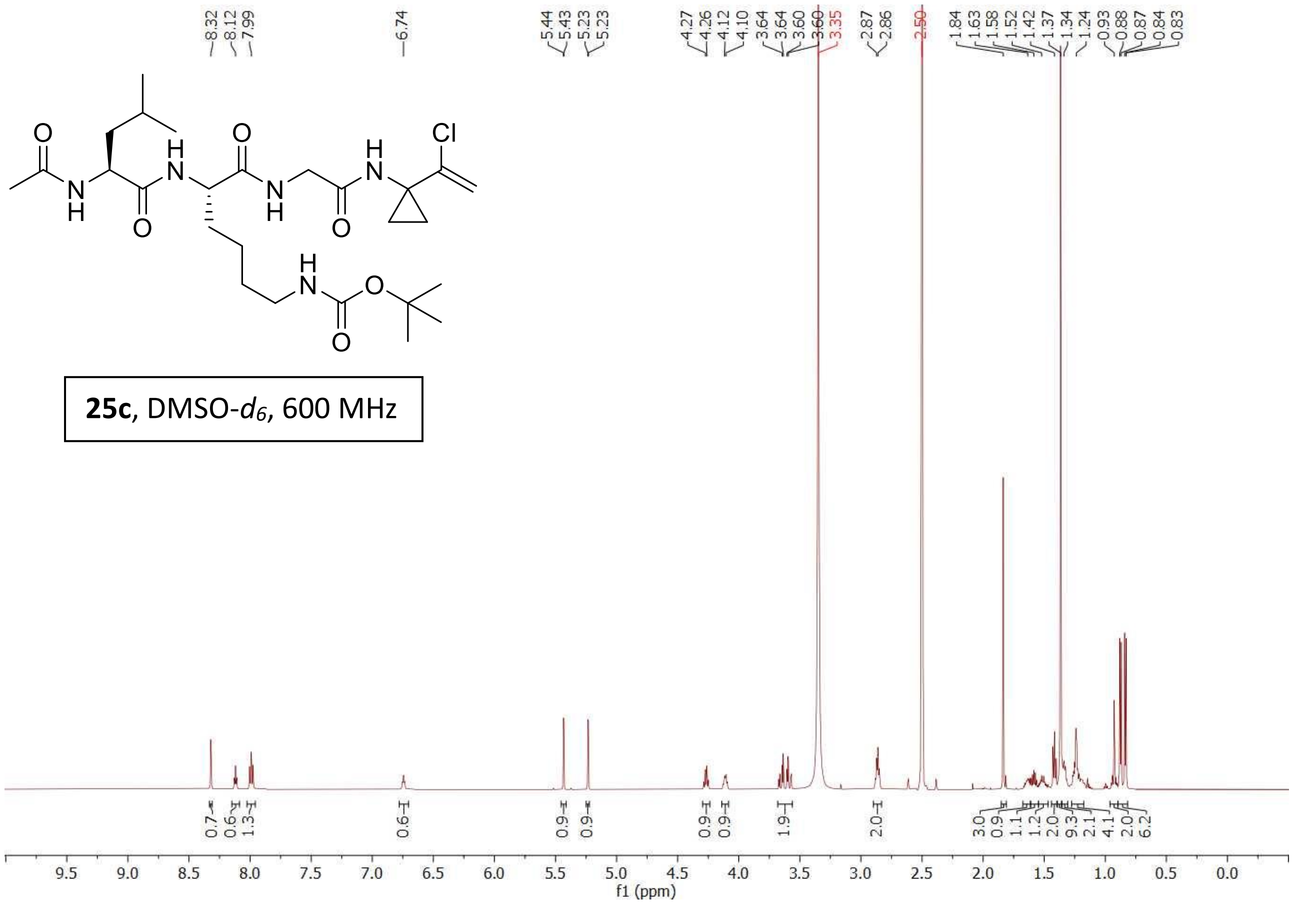


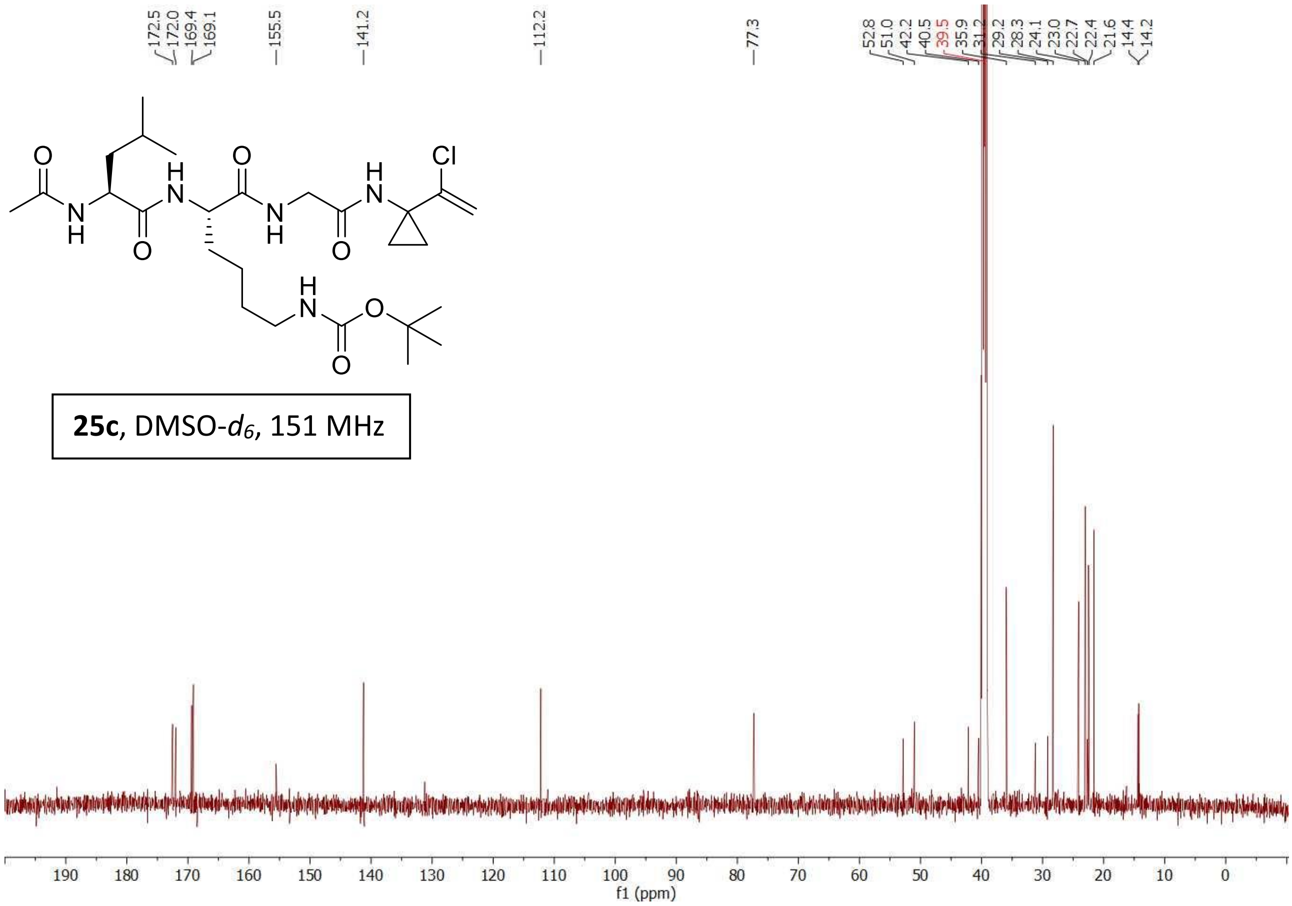

Compound **25g**

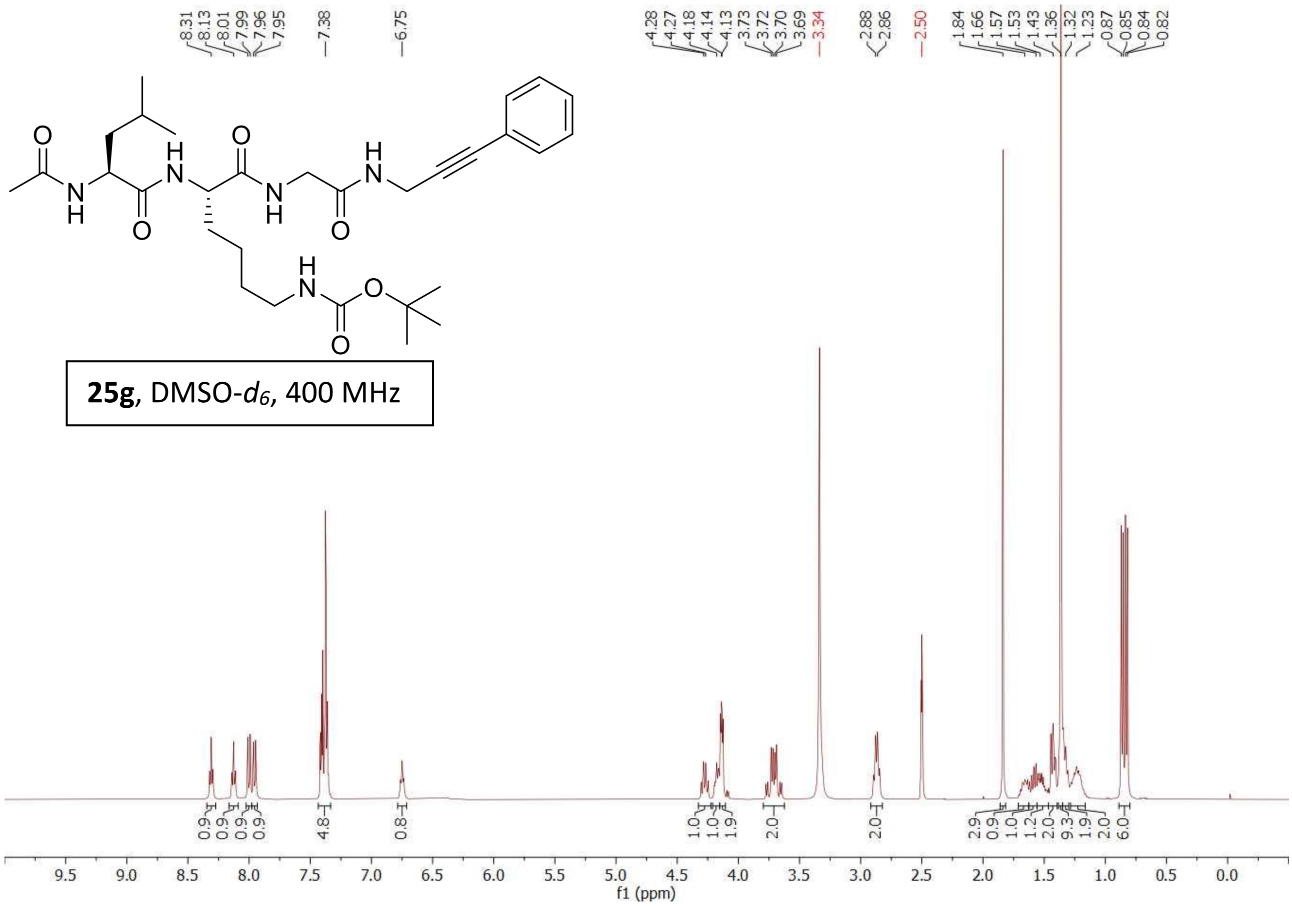


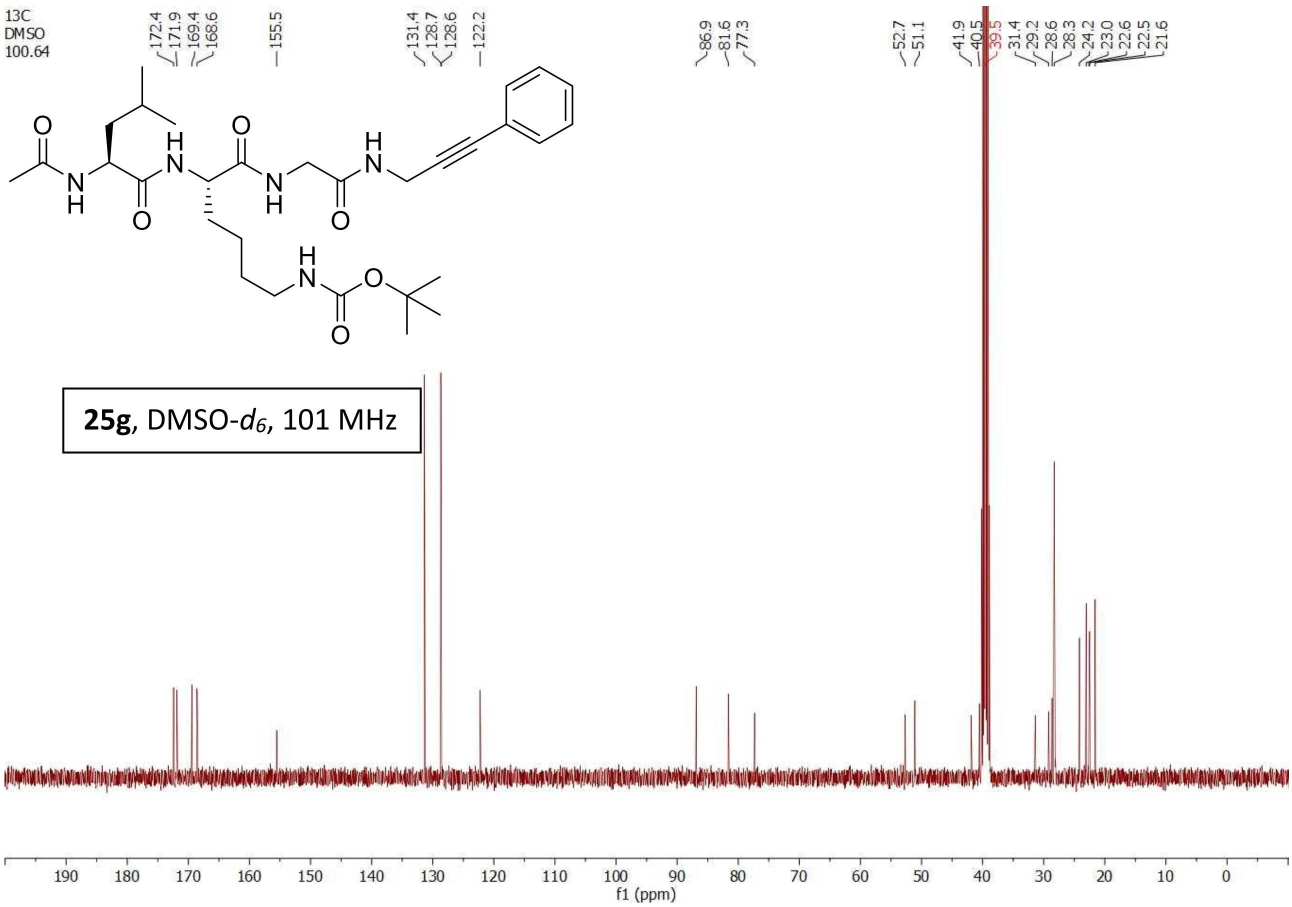

Compound **25a**

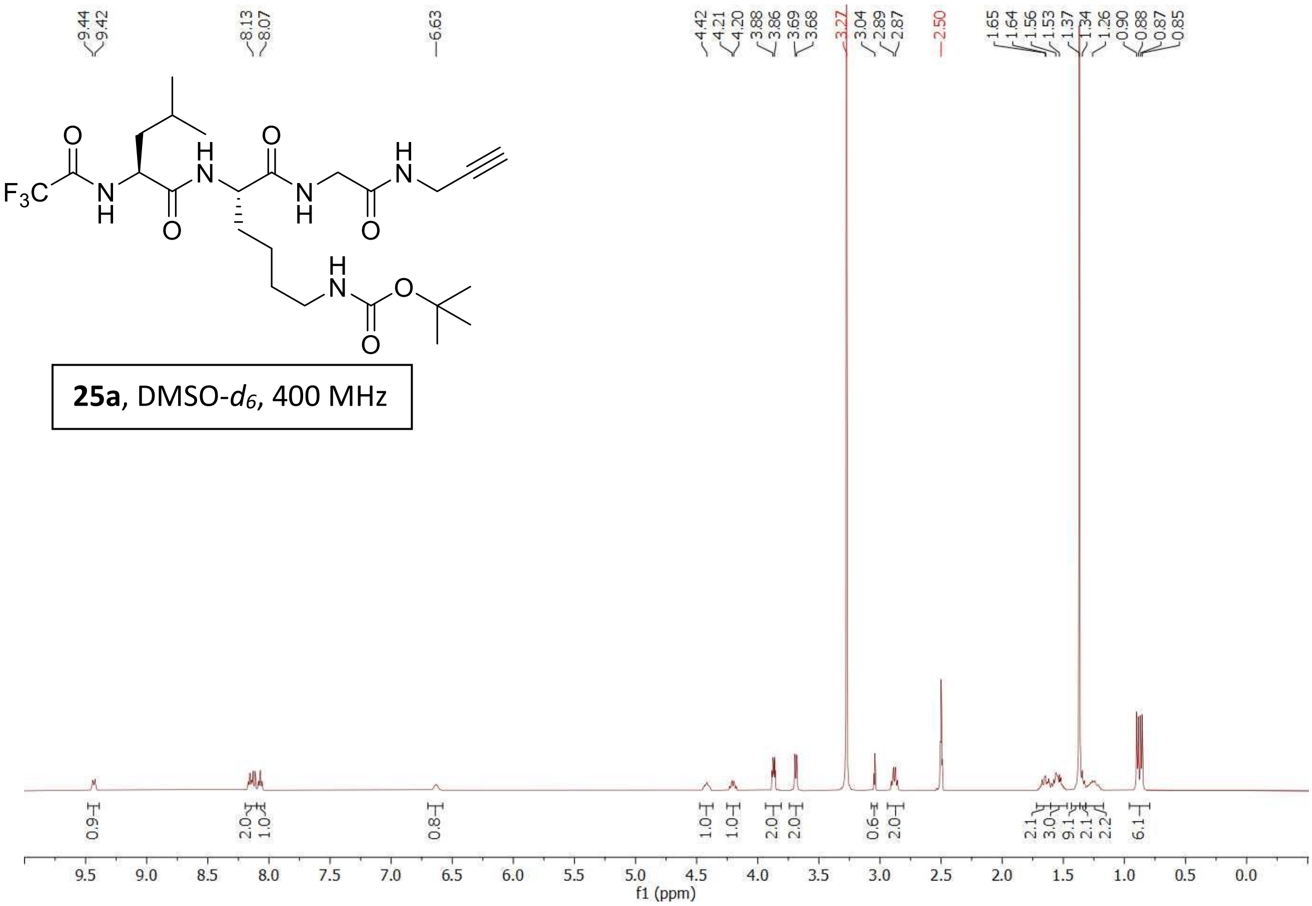


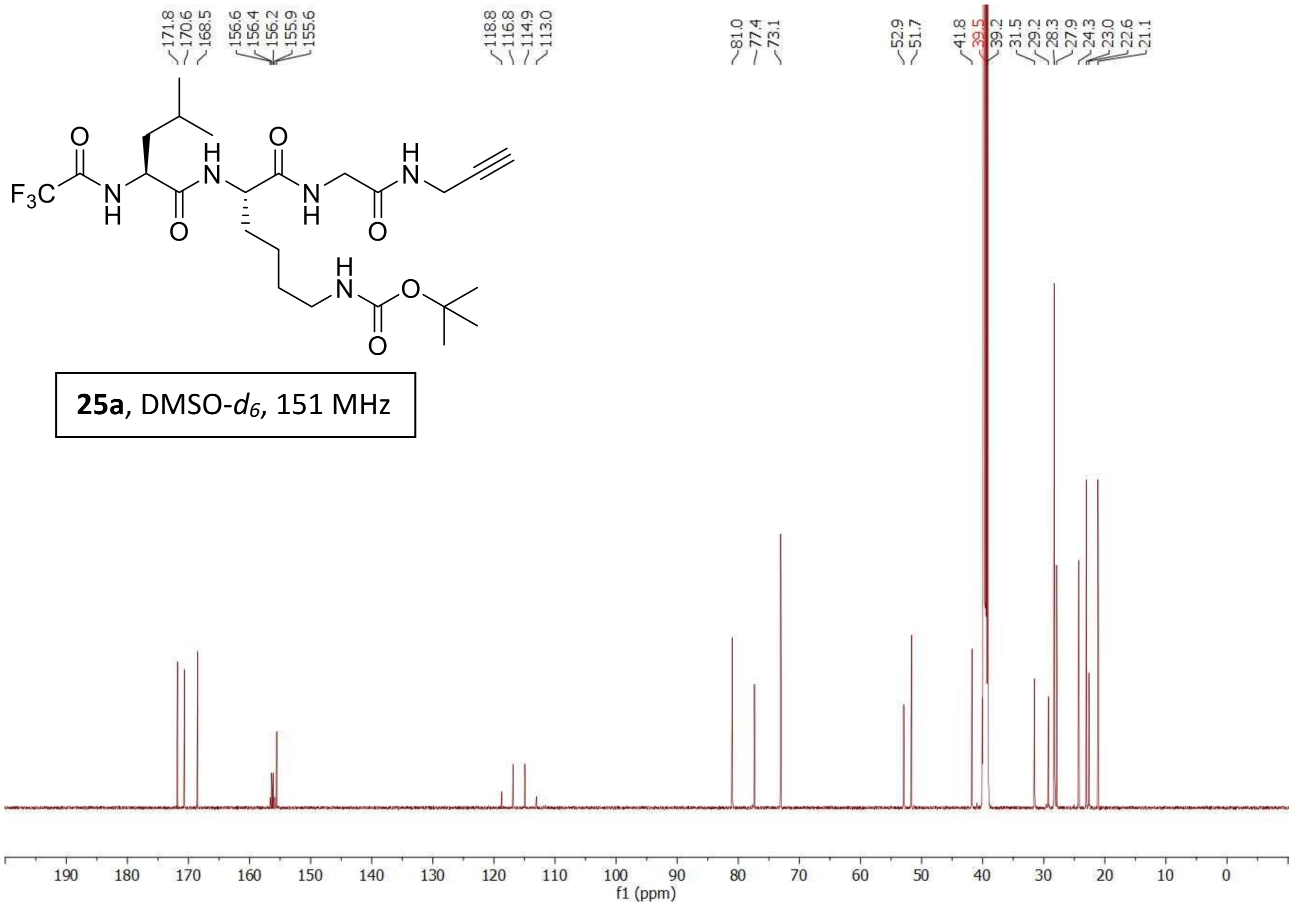

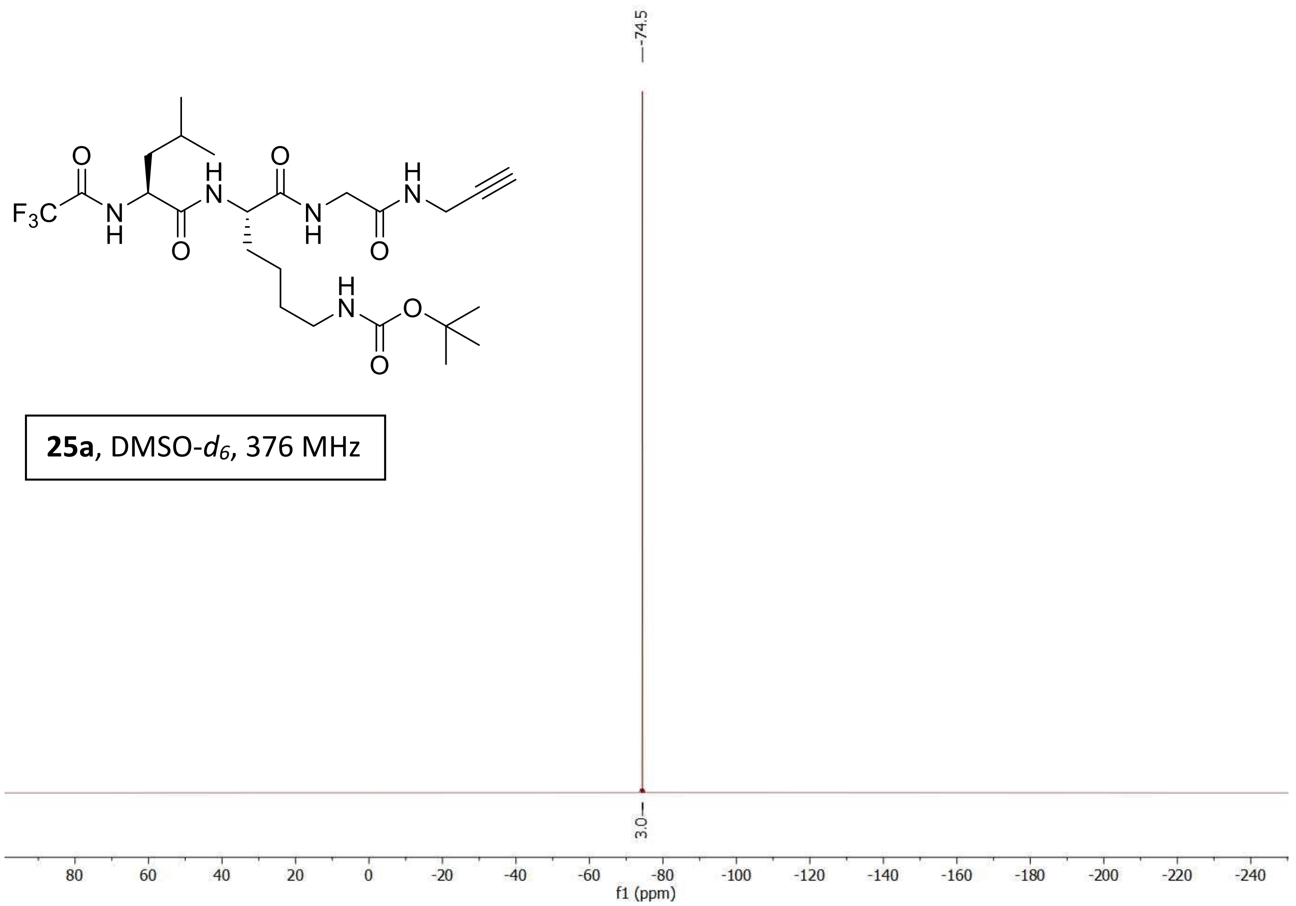


**25a**, DMSO-$d_6$, 376 MHz

Compound **25**

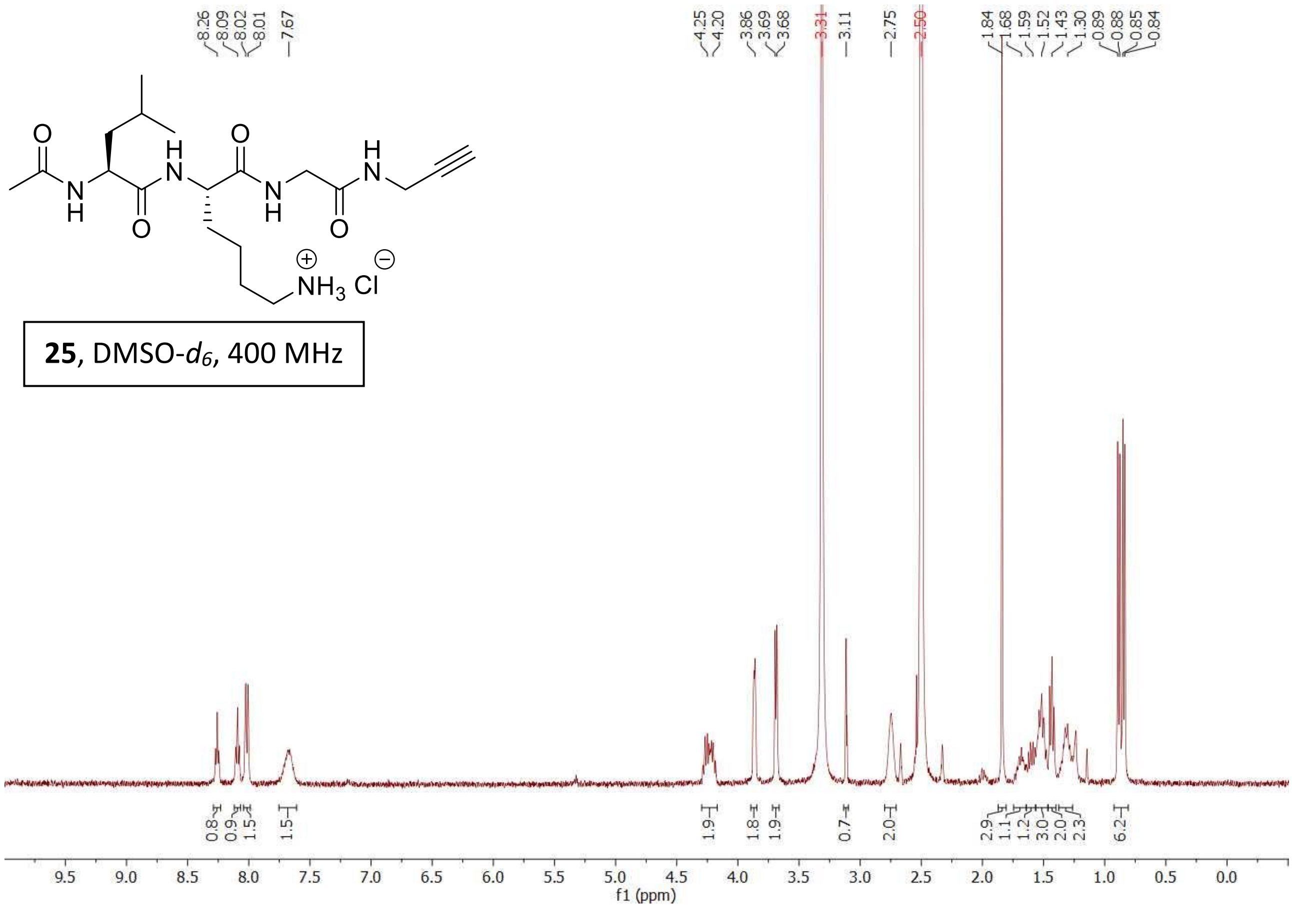


**25**, DMSO-$d_6$, 400 MHz

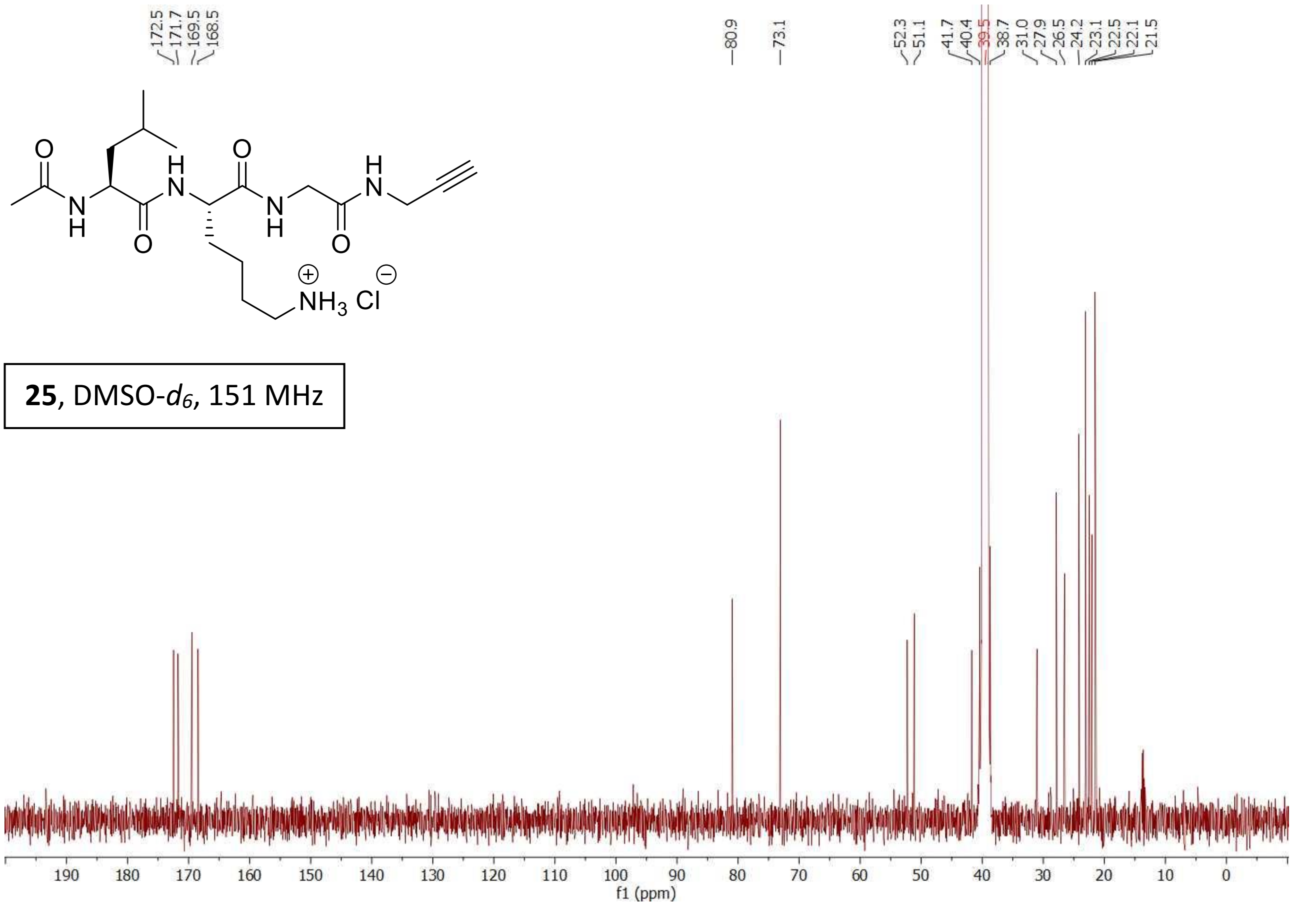


**25**, DMSO-$d_6$, 151 MHz

Compound **25j**

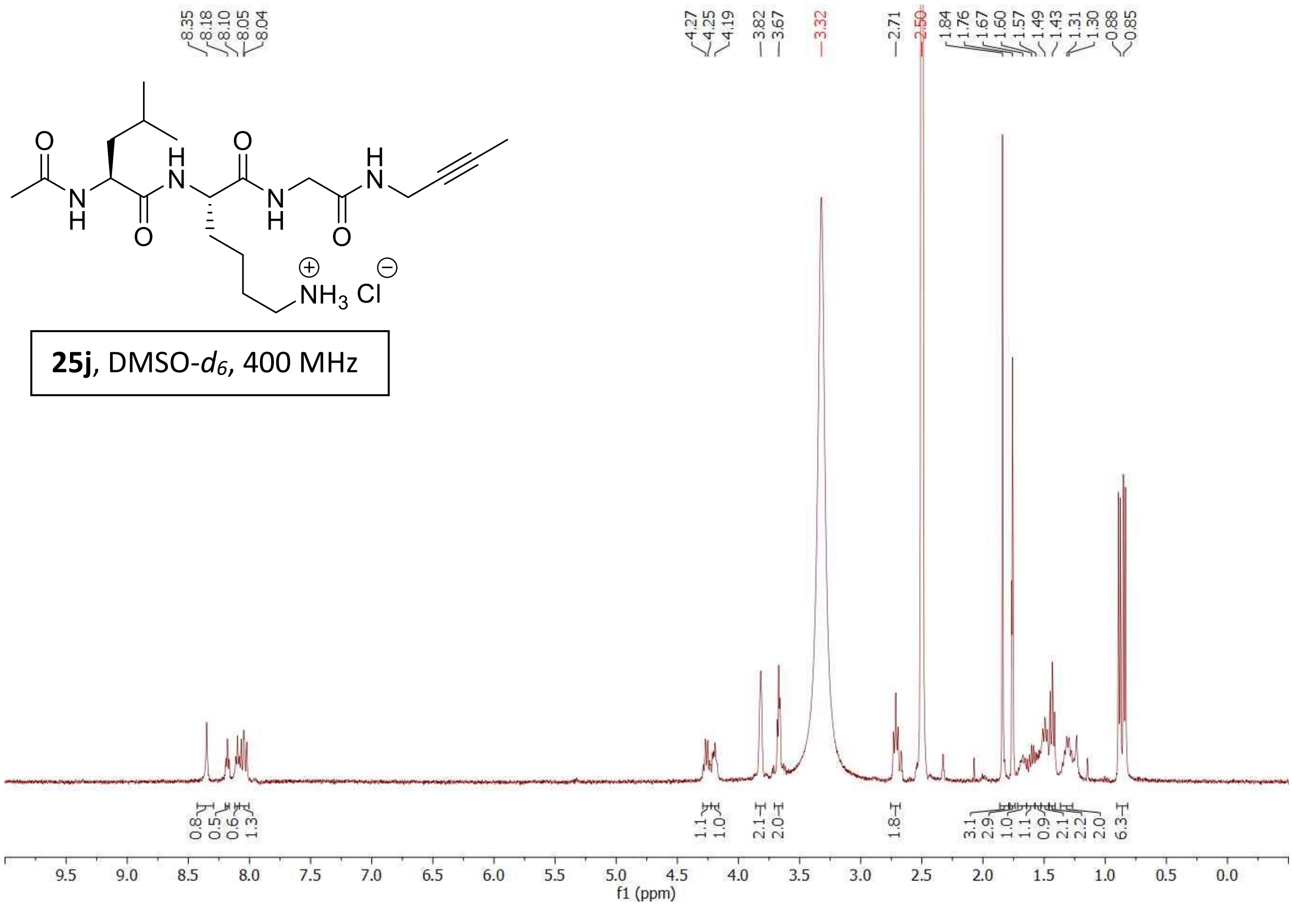


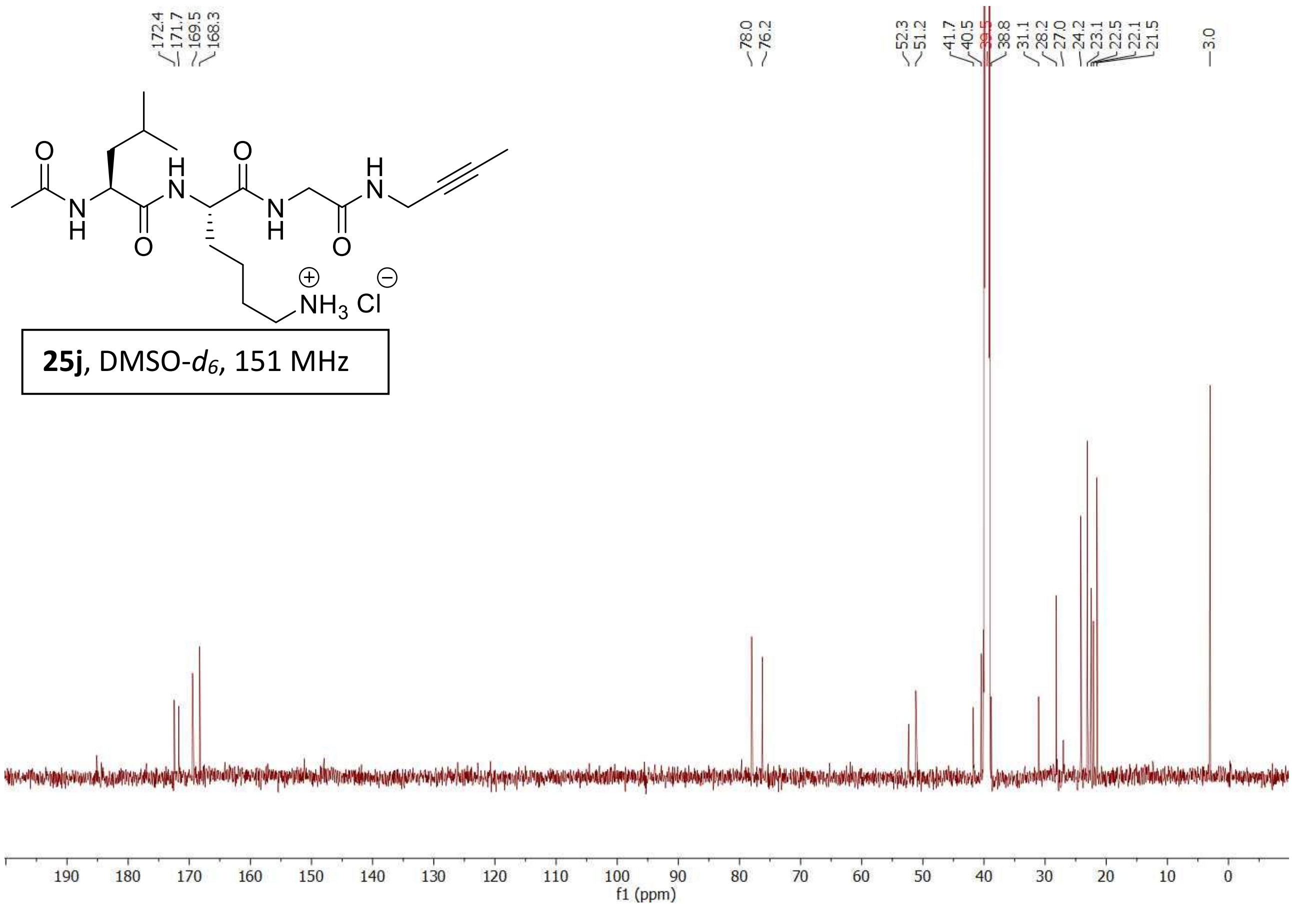

Compound **25h**

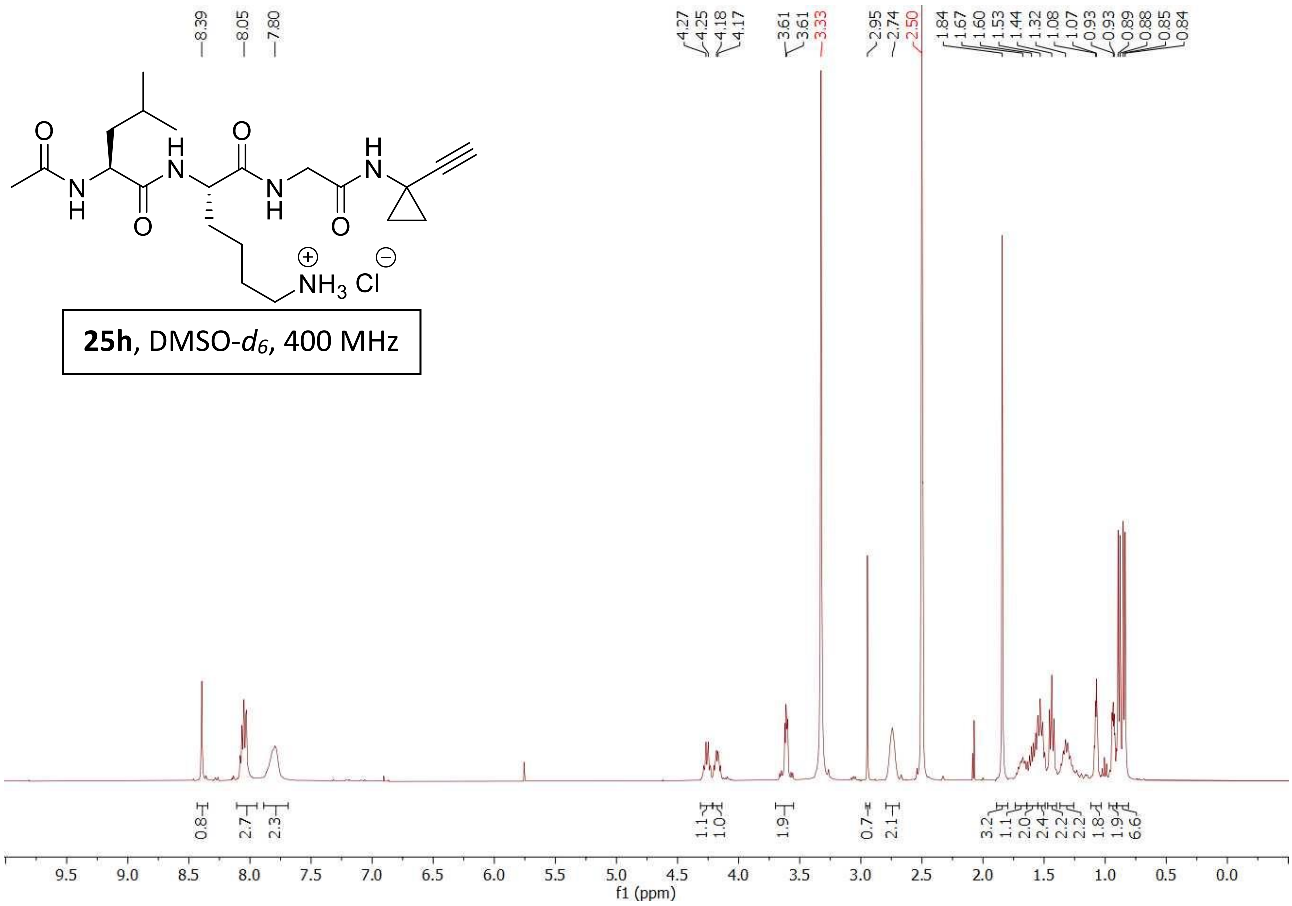


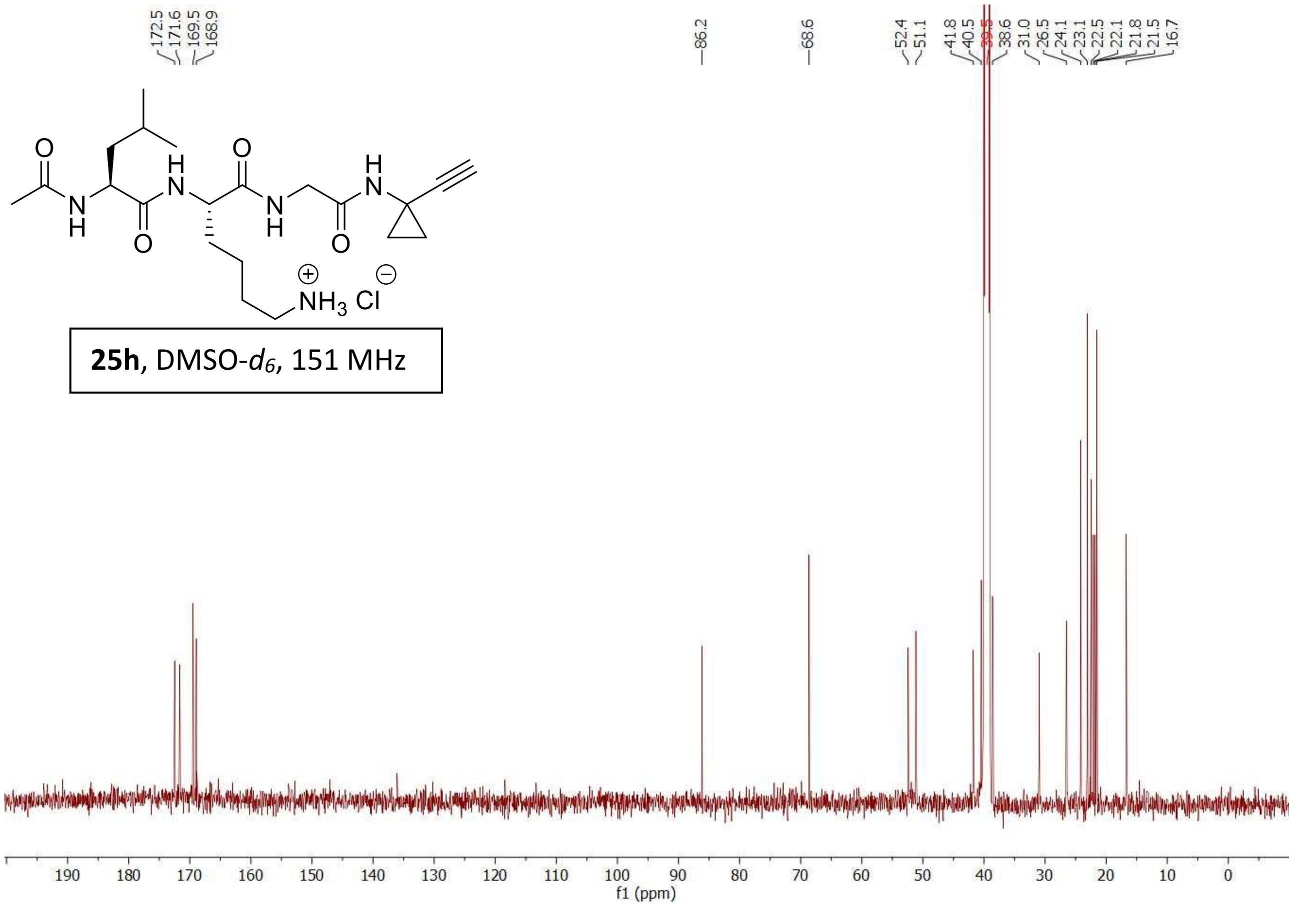

Compound **25i**

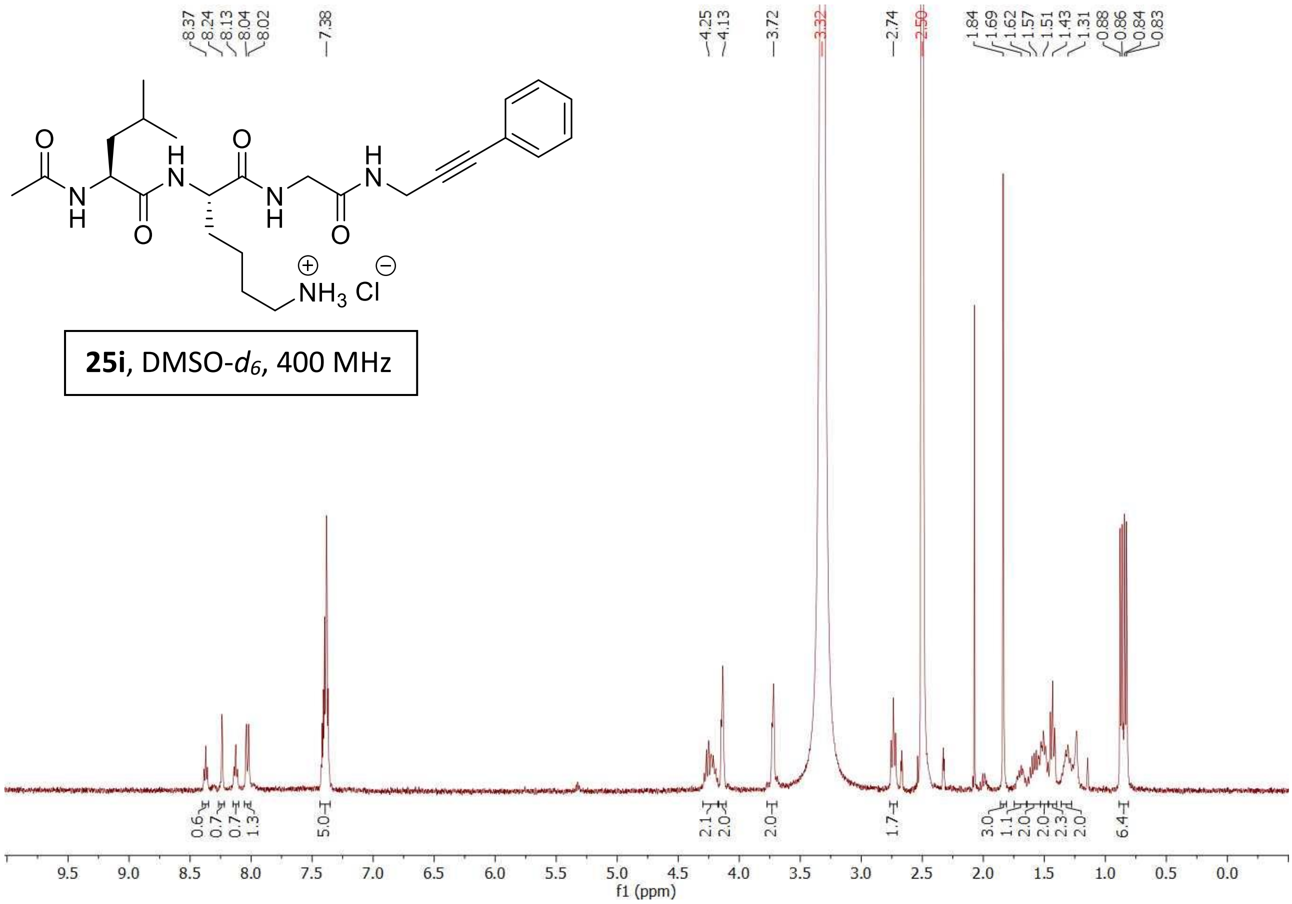


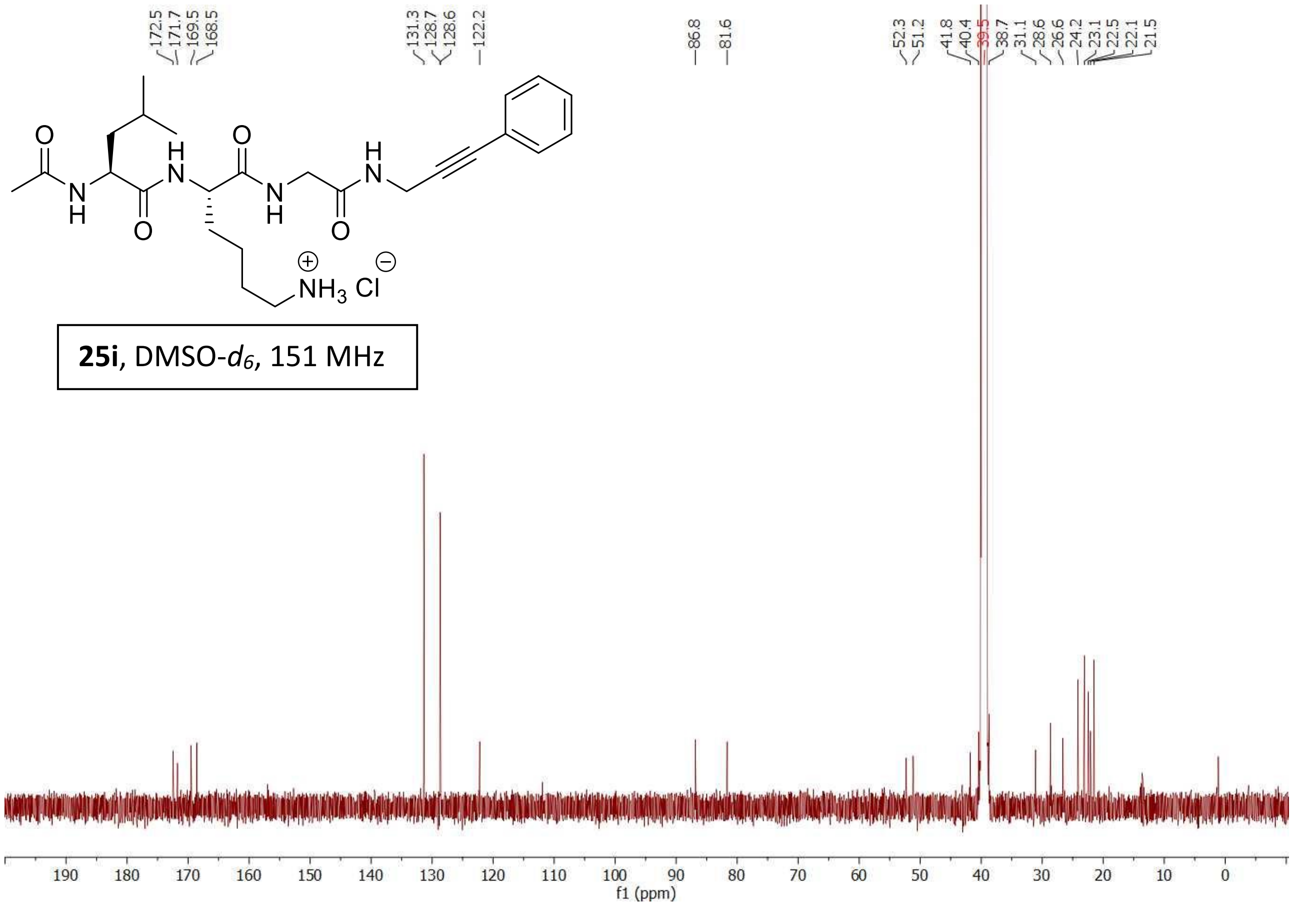

Compound **25k**

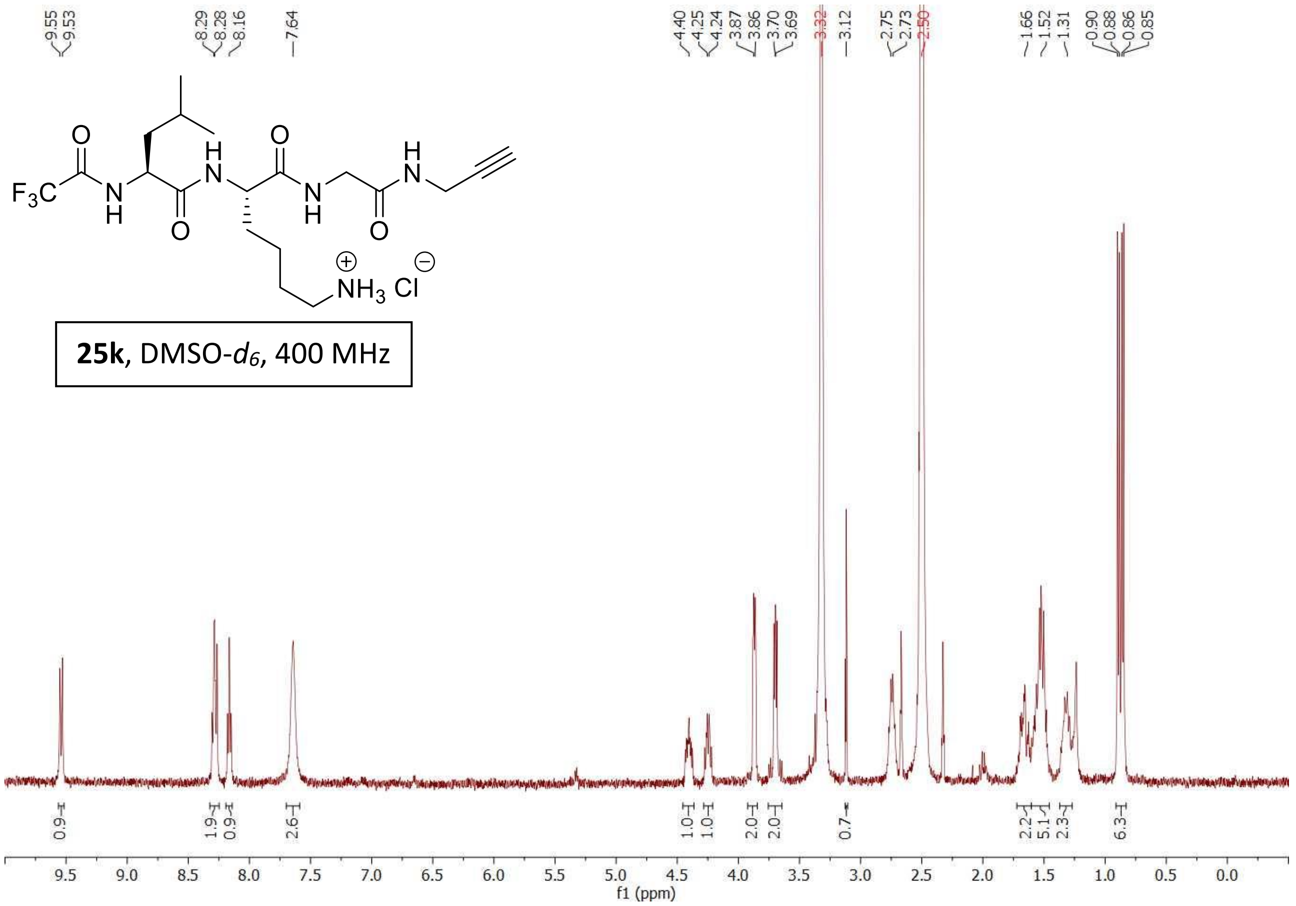


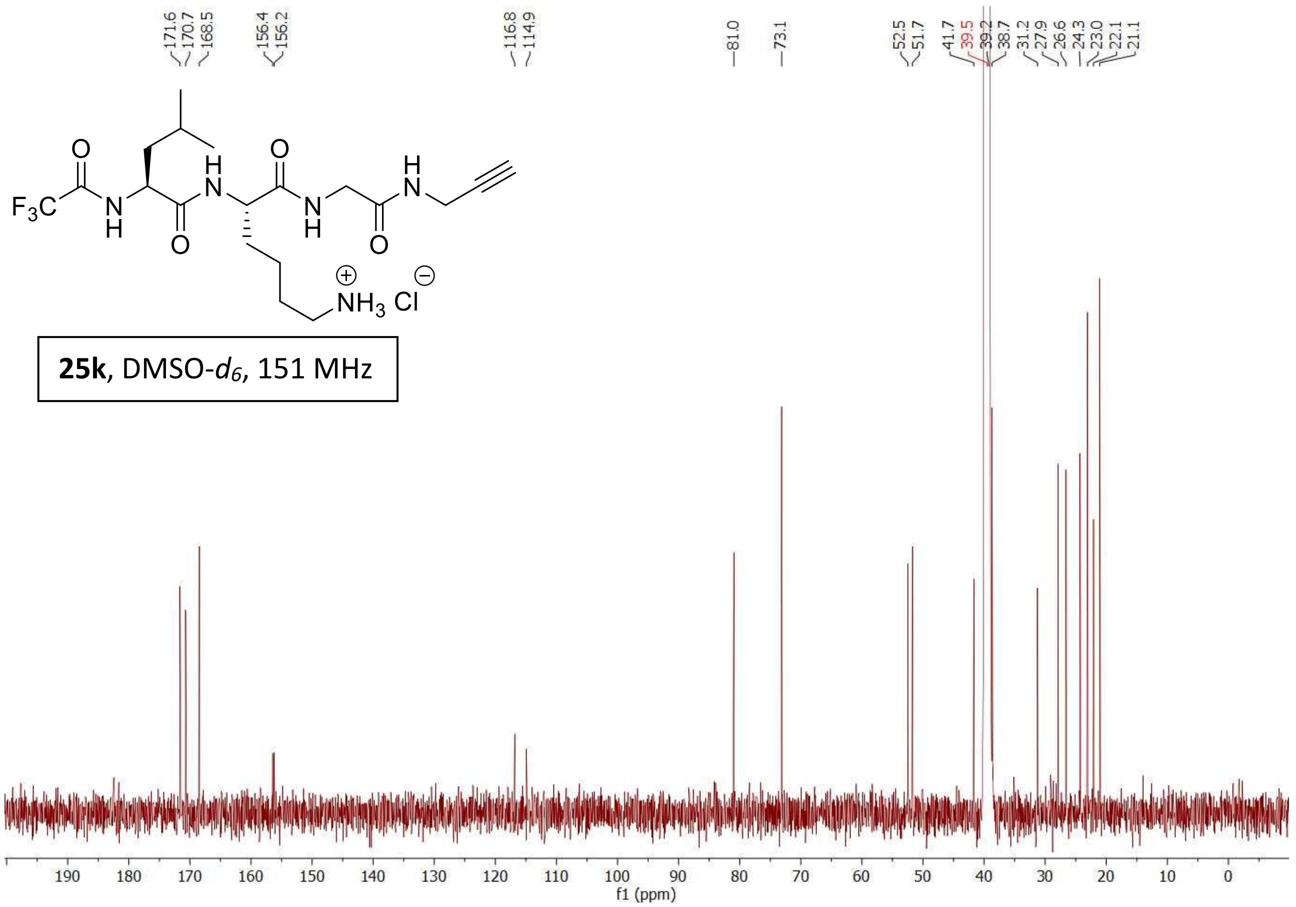

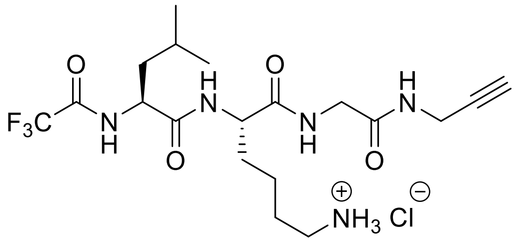

**25k**, DMSO-$d_6$, 376 MHz

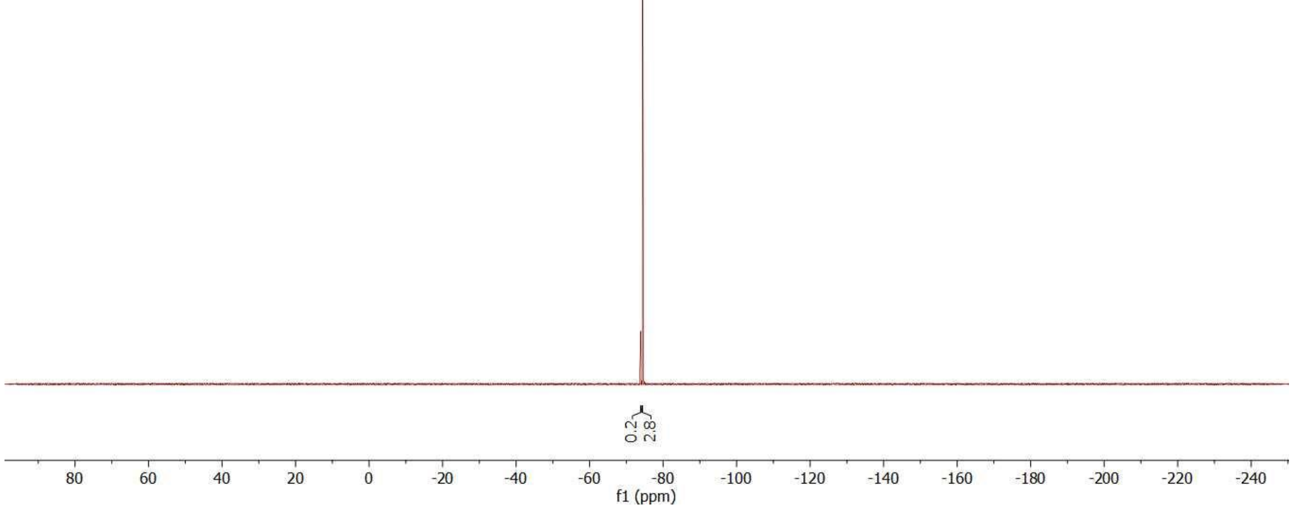

Compound **25b**

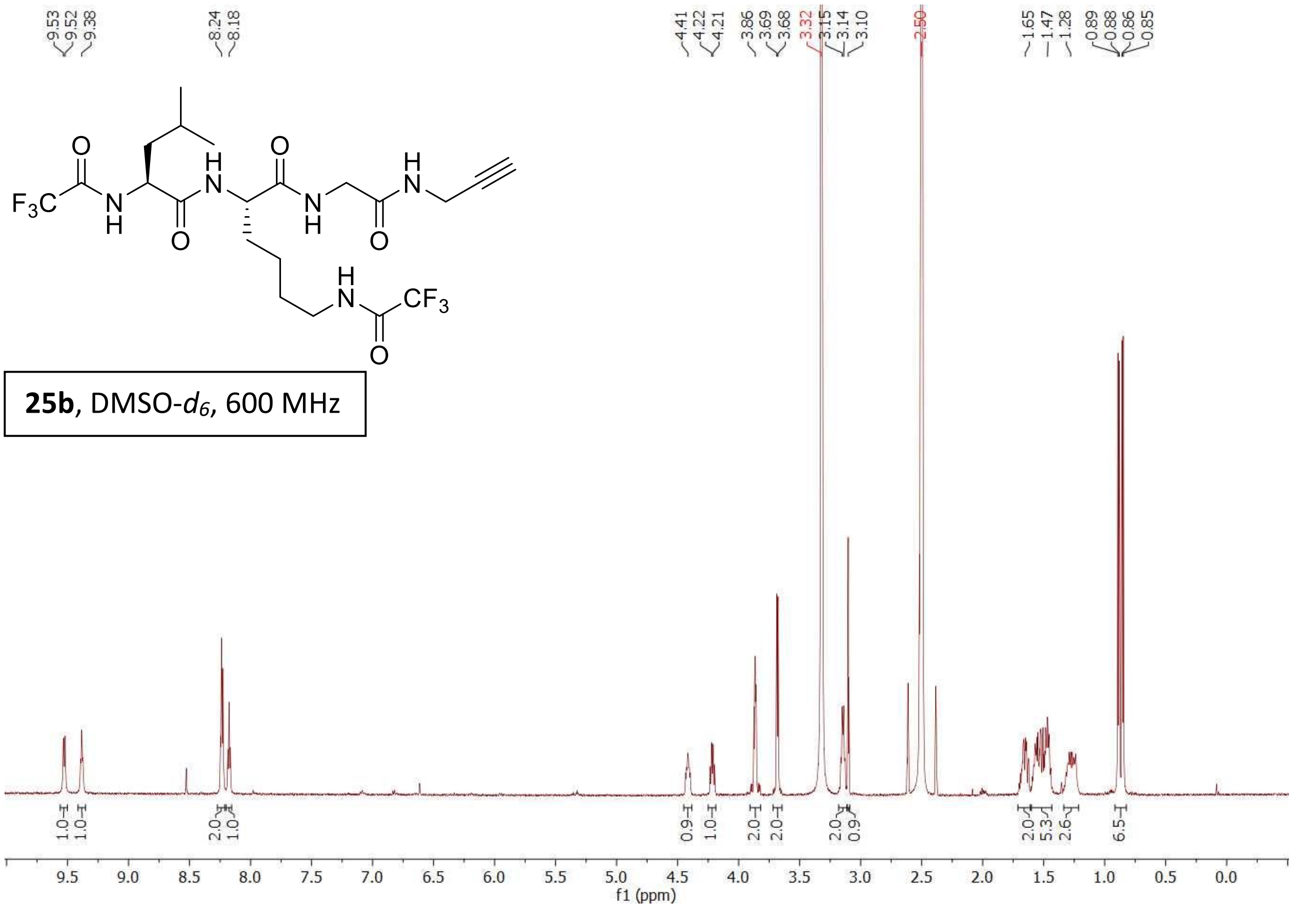


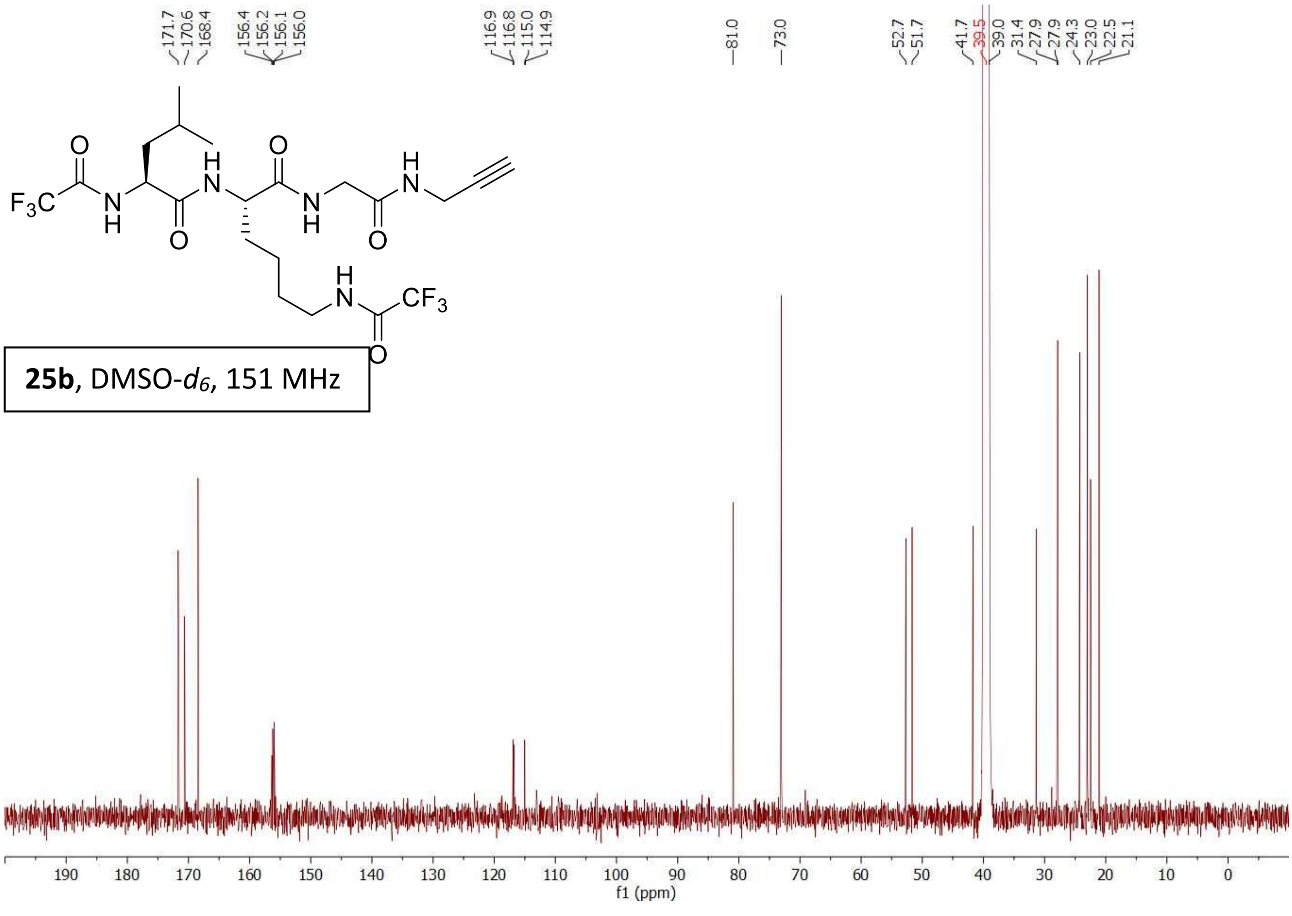

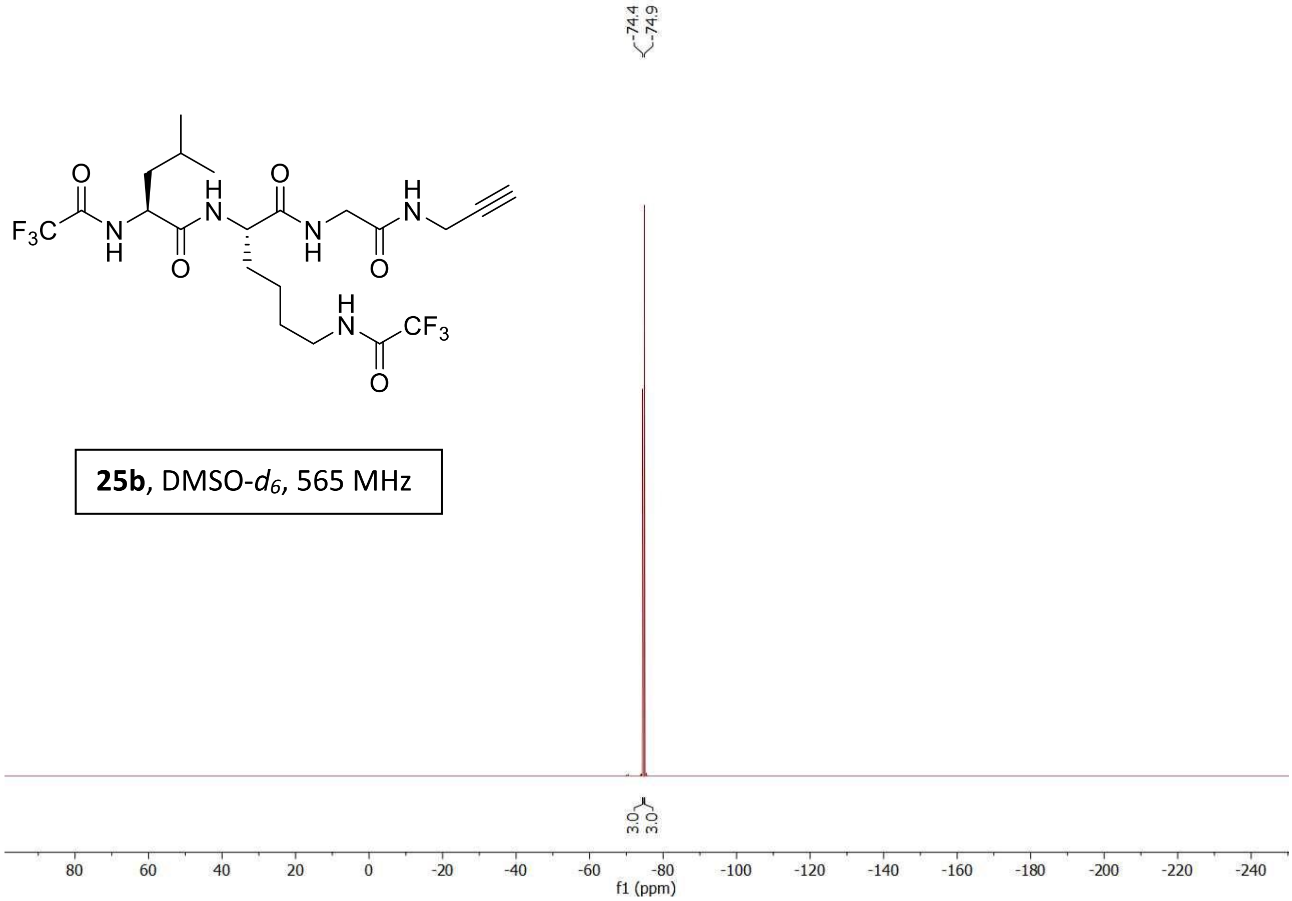


**25b**, DMSO-$d_6$, 565 MHz
